## Supplemental files for "Effective prediction of biosynthetic pathway genes involved in bioactive polyphyllins in Paris polyphylla"

**Supplementary file1**

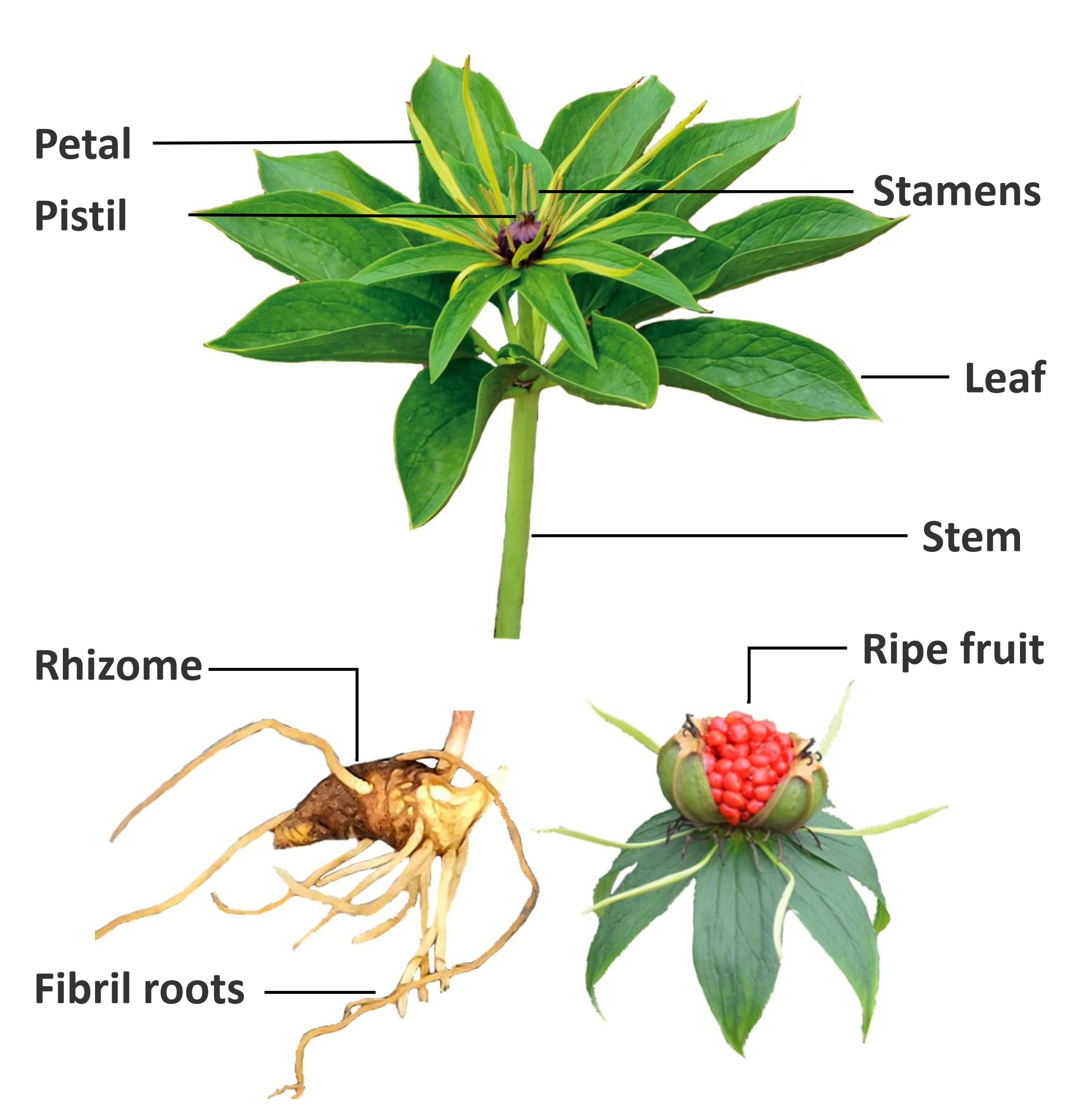

**Supplementary Figure 1.** Different organs of *P. polyphylla* var*. yunnanensis* used for transcriptomic and metabolomic analysis.

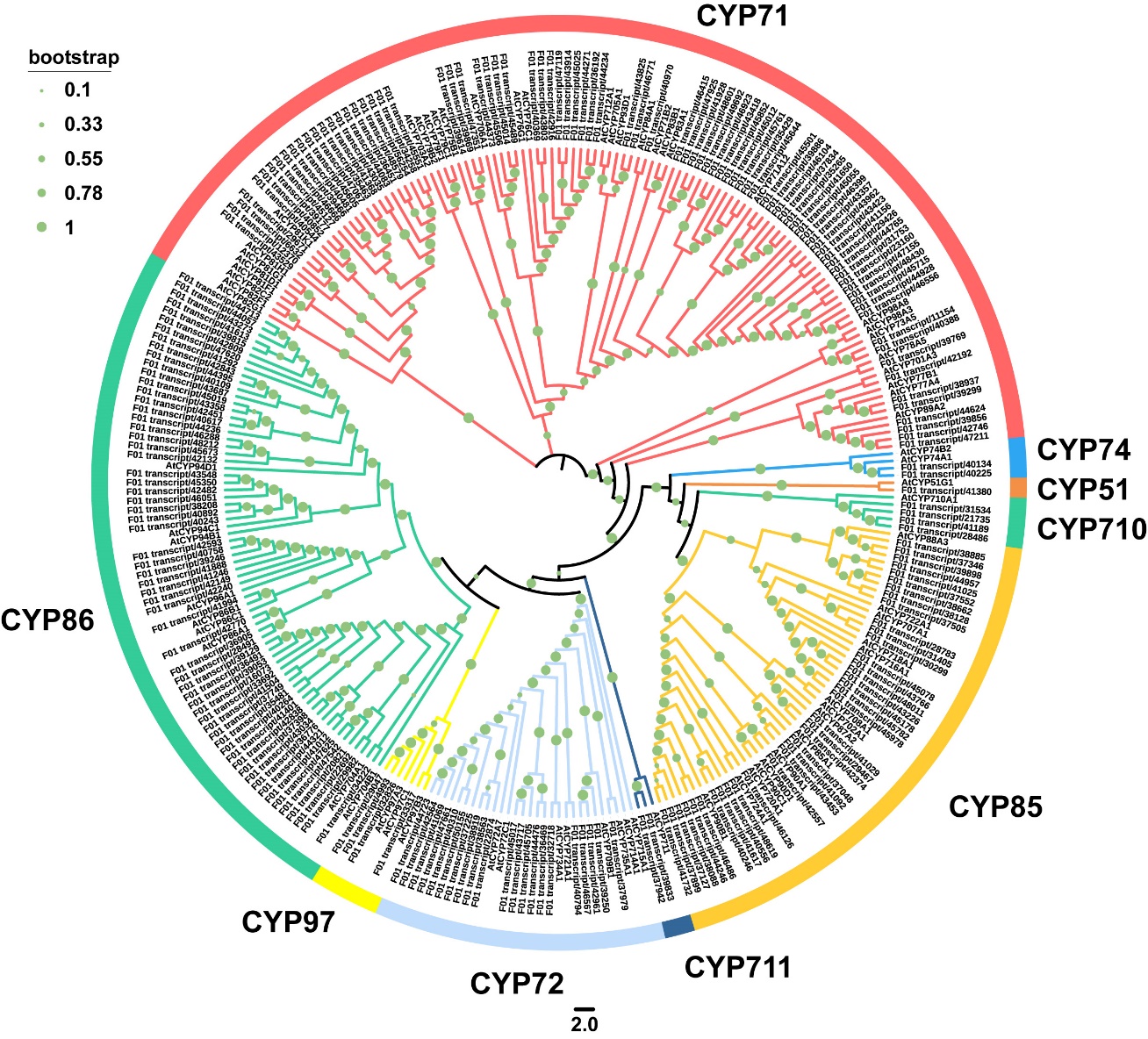

**Supplementary Figure** **2**. Phylogenetic tree of CYPs. Selected Arabidopsis CYPs were included in the analysis to define CYP families. The evolutionary history was inferred using the Neighbor-Joining method. The bootstrap consensus tree inferred from 1000 replicates is taken to represent the evolutionary history of the taxa analyzed. Evolutionary analyses were conducted in MEGA6. According to the reported family CYP genes of *Arabidopsis thaliana*, the phylogenetic tree is divided into CYP51, CYP71, CYP710, CYP711, CYP72, CYP74, CYP85, CYP86, and CYP94.

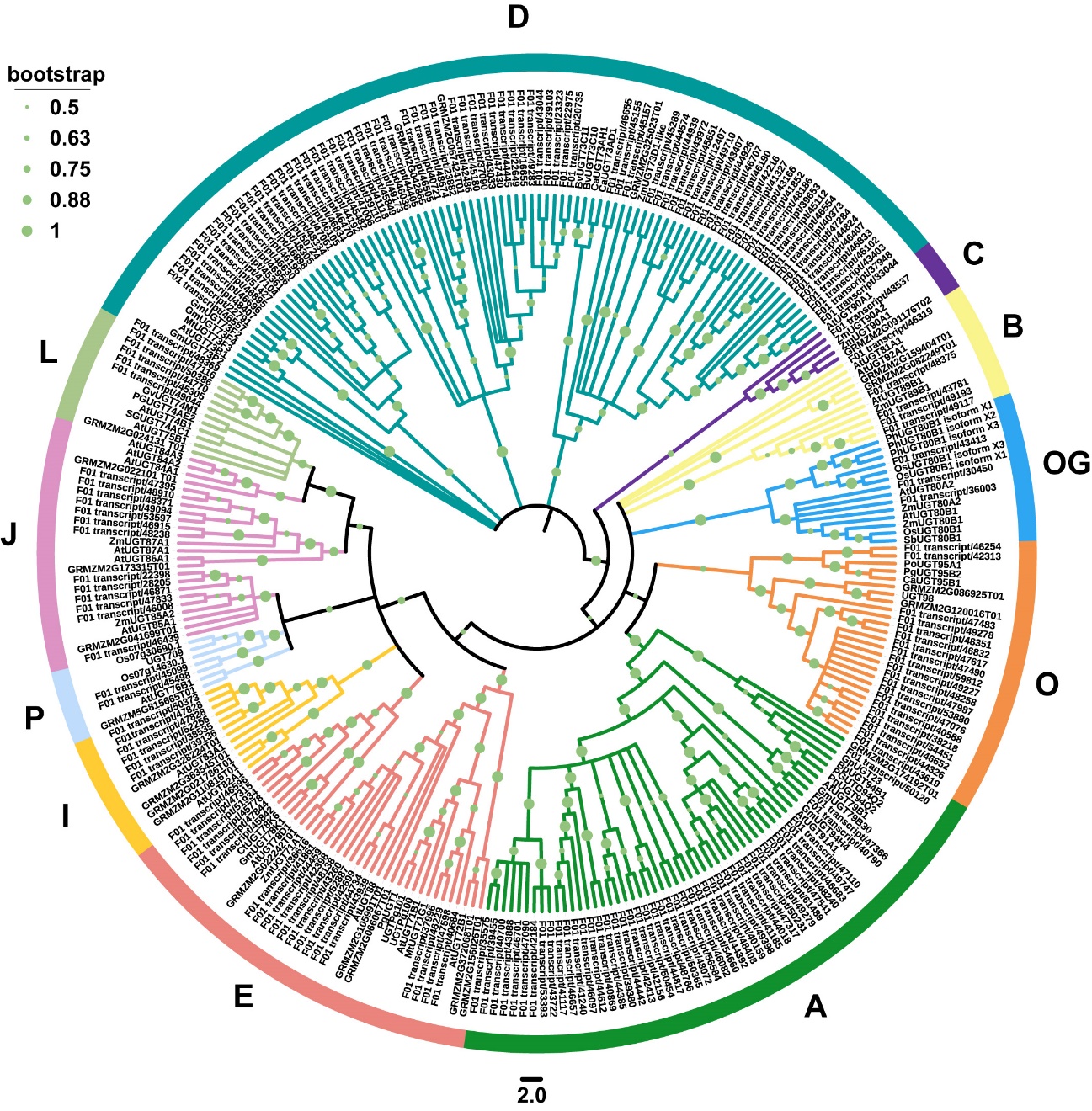
**Supplementary Figure 3.** Phylogenetic tree of UGTs. Predicted amino acid sequences of UGTs were aligned with selected UGTs from multiple species using MUSCLE. A neighbor-joining tree was constructed using MEGA6. At, *Arabidopsis thaliana*; Bp, *Bellis perennis*; Bv, *Barbarea vulgaris*; Ca, *Cicer arietinum*; Ct, Crystal structure; Gm, Glycine max; Gv, *Gypsophila vaccaria*; Mt, *Medicago truncatula*; Os, *Oryza sativa*; Pg, *Panax ginseng*; Ph, *Petunia maxim hybrid*; Po, *Pilosella officinarum*; Sg, *Siraitia grosvenorii*; Zm, *Zea mays*; GRMZM, UGT sequences from maize sequence database.

**Supplementary Figure**
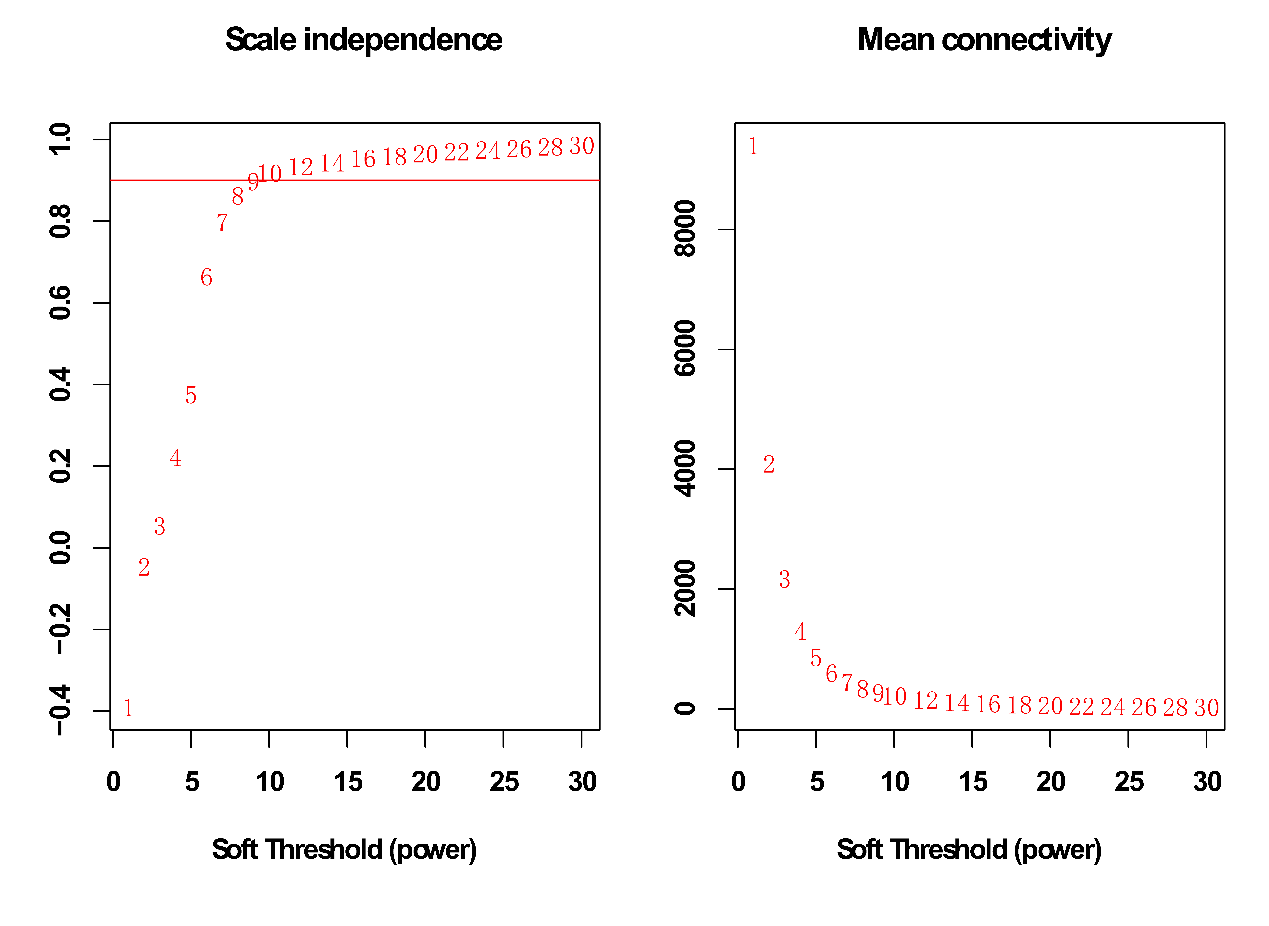
**4.** dendrogram of 8 tissue samples from *Paris polyphylla* var. *yunnanensis.* The sample clustering tree is constructed by calculating the correlation coefficient of the expression level of each sample, and outlier samples are checked and removed.

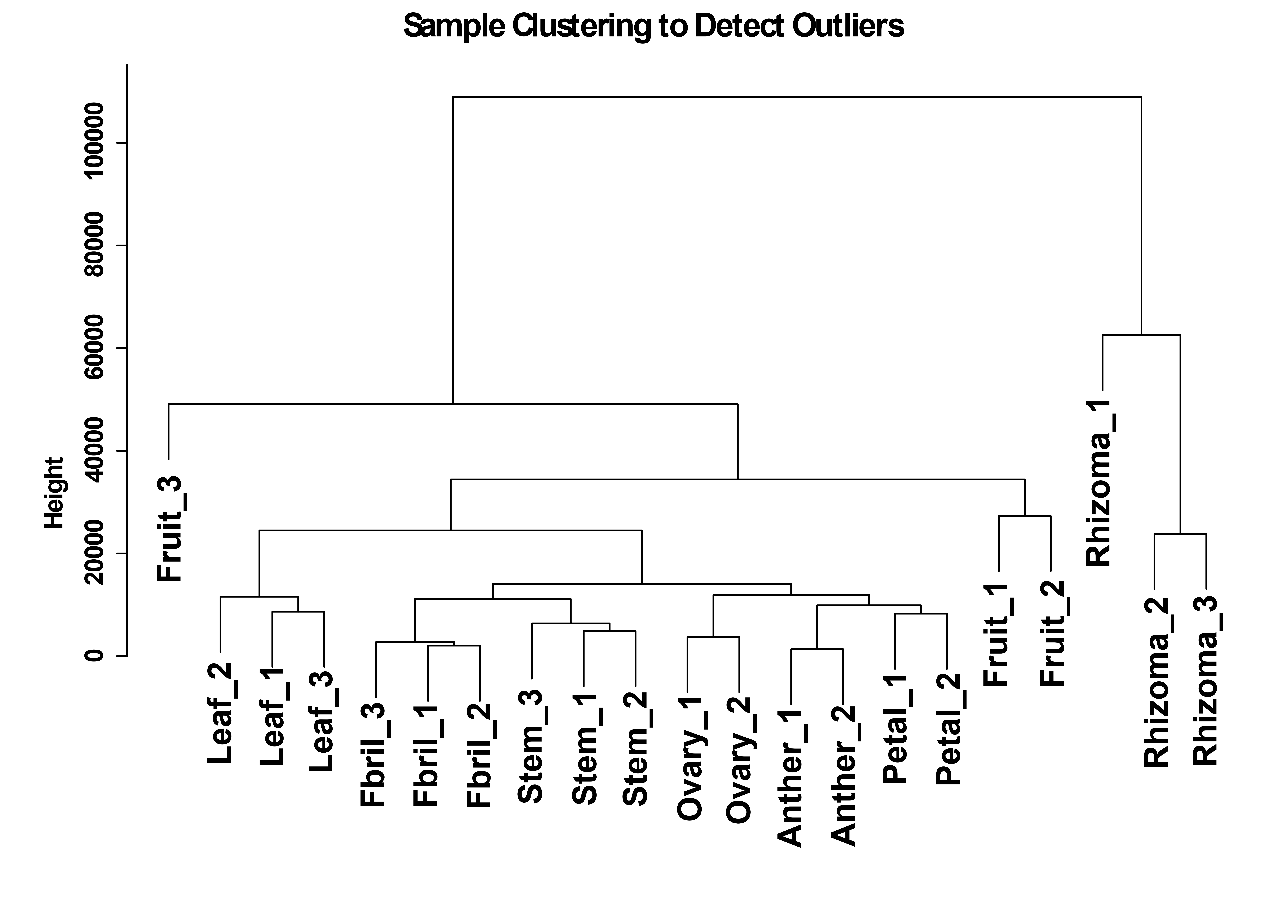
 **Supplementary Figure 5.** Soft threshold determination of *Paris polyphylla* var. *yunnanensis* genes co-expression network (A) The scale-free fit index (y-axis) as a function of the soft-thresholding power (x-axis) and (B) the mean connectivity (degree, y-axis) as a function of the soft-thresholding power (x-axis).

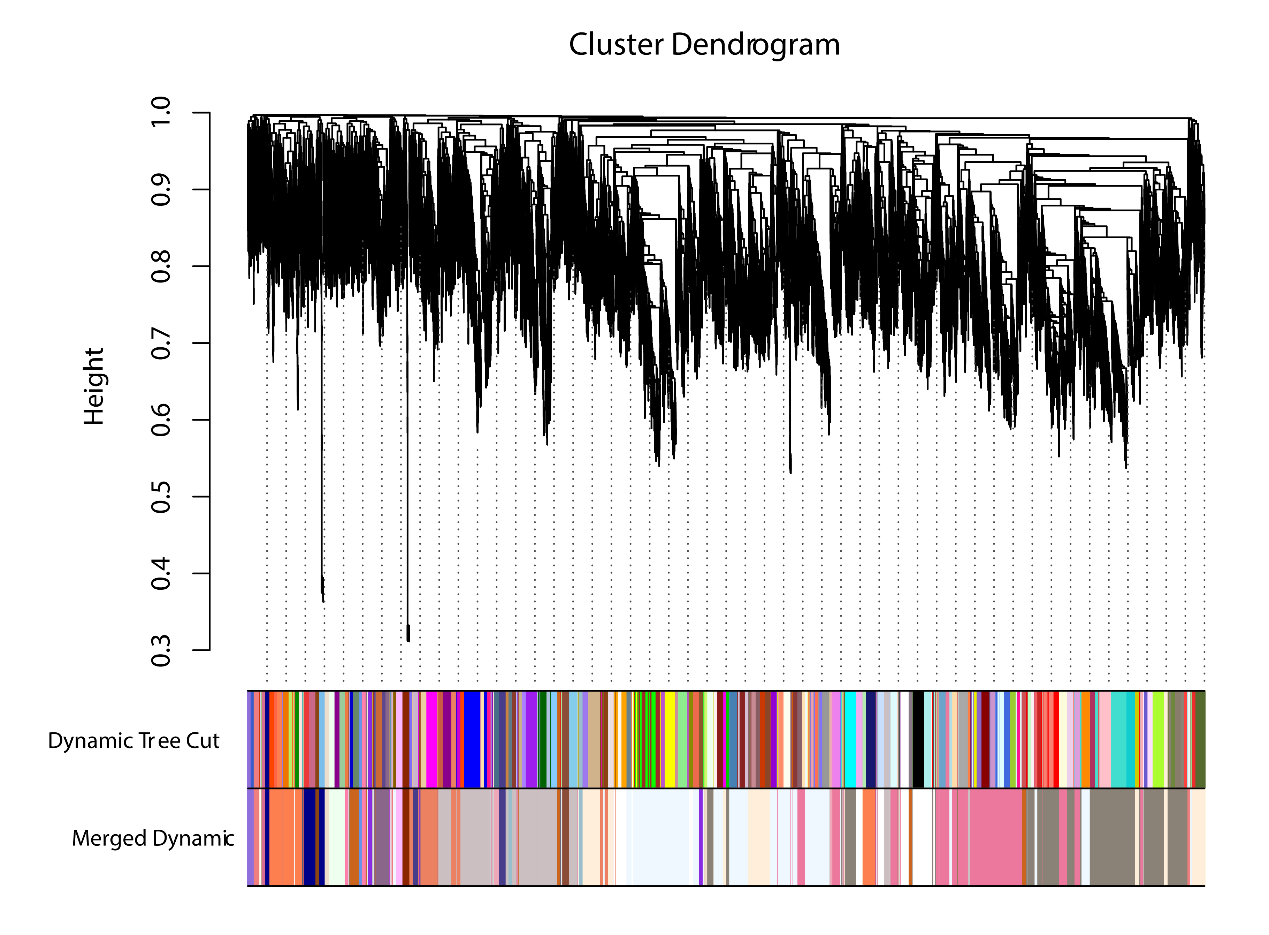
**Supplementary Figure 6.** Clustering dendrograms of *Paris polyphylla* var. *yunnanensis* genes. Each clade corresponds to a module and is marked with a different color.

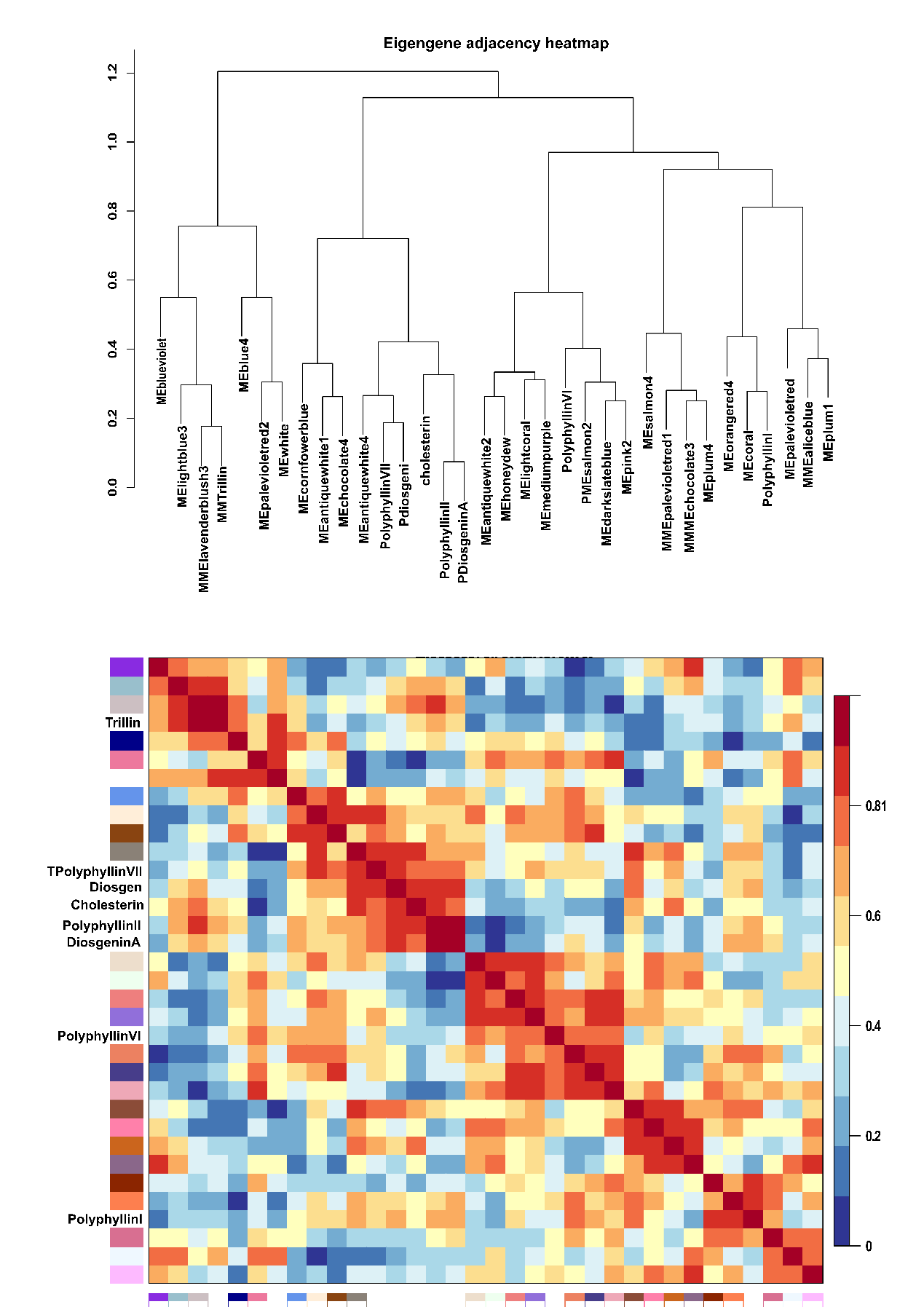
**Supplementary Figure 7.** Visualizing genes and metabolites co-expression network using a heatmap plot. The heatmap depicts the topological overlap matrix among all genes with the key metabolites in the biosynthetic pathway of polyphyllin. The lighter color represents the low correlation between the genes and metabolites biosynthesis, and the gradually darker color represents the strong correlation between the genes and metabolites biosynthesis.

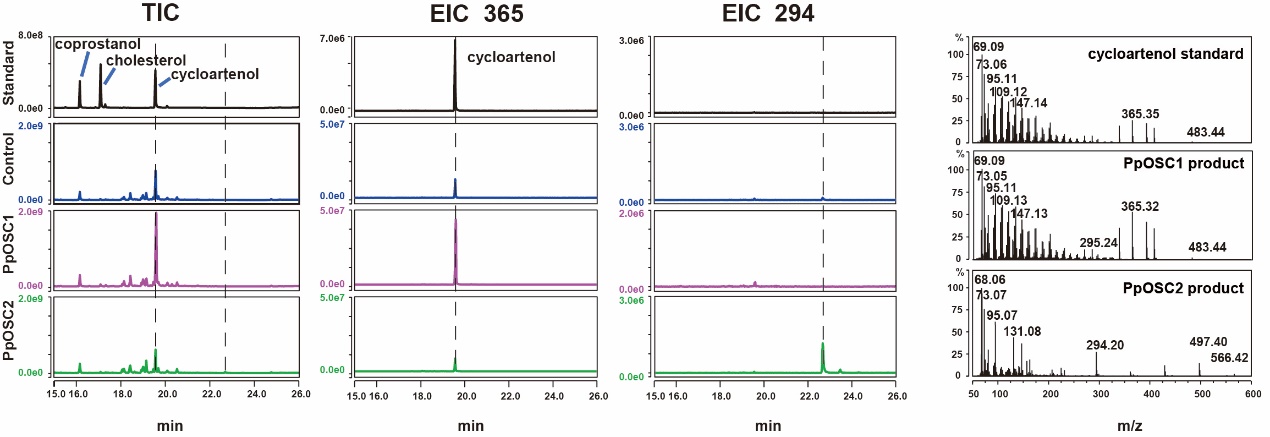

**Supplementary Figure 8.** Functional verification of *PpOSC* gene. Two OSC genes were identified in *P. polyphylla* var*. yunnanensis*, and the results of GC-MS showed that *PpOSC1* gene increased the yield of cycloartenol after being transferred to *N.benthamiana*.

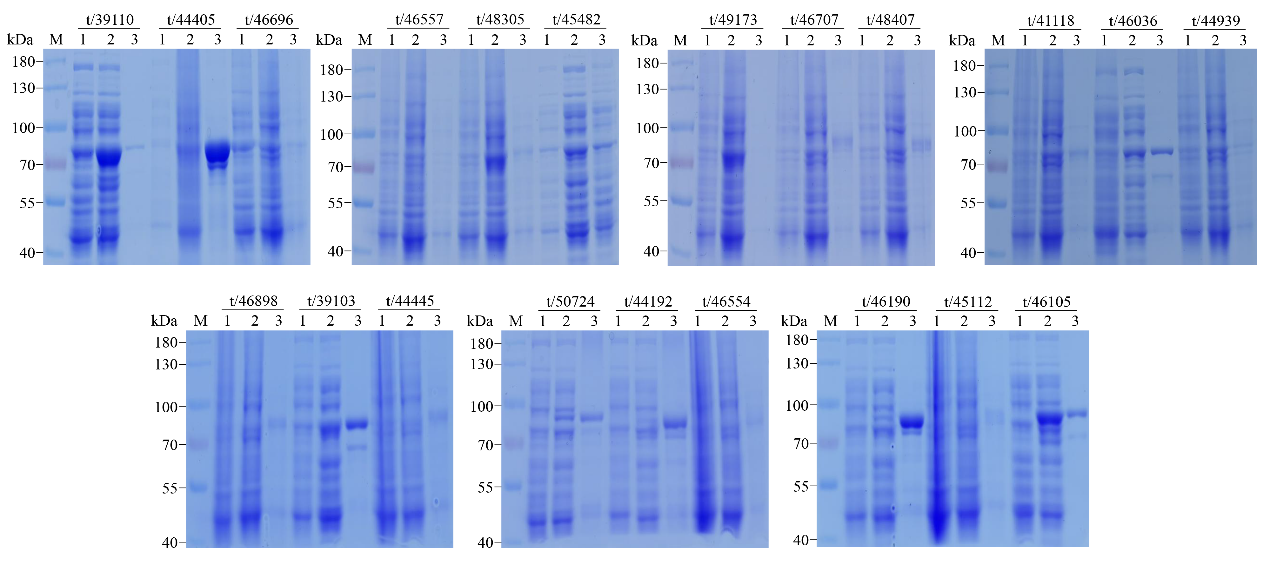

**Supplementary Figure 9.** The protein expression results of candidate UGTs. *Lanes*: M, protein molecular weight marker (Thermo fisher); 1, pGEX-*PpUGT* vectors transformed in *E. coli* Rosetta (DE3) cells without IPTG induction; 2, supernatant of pGEX-*PpUGT* vectors transformed in *E. coli* Rosetta (DE3) cells with IPTG induction; 3, pellet of pGEX-*PpUGT* vectors transformed in *E. coli* Rosetta (DE3) cells with IPTG induction.

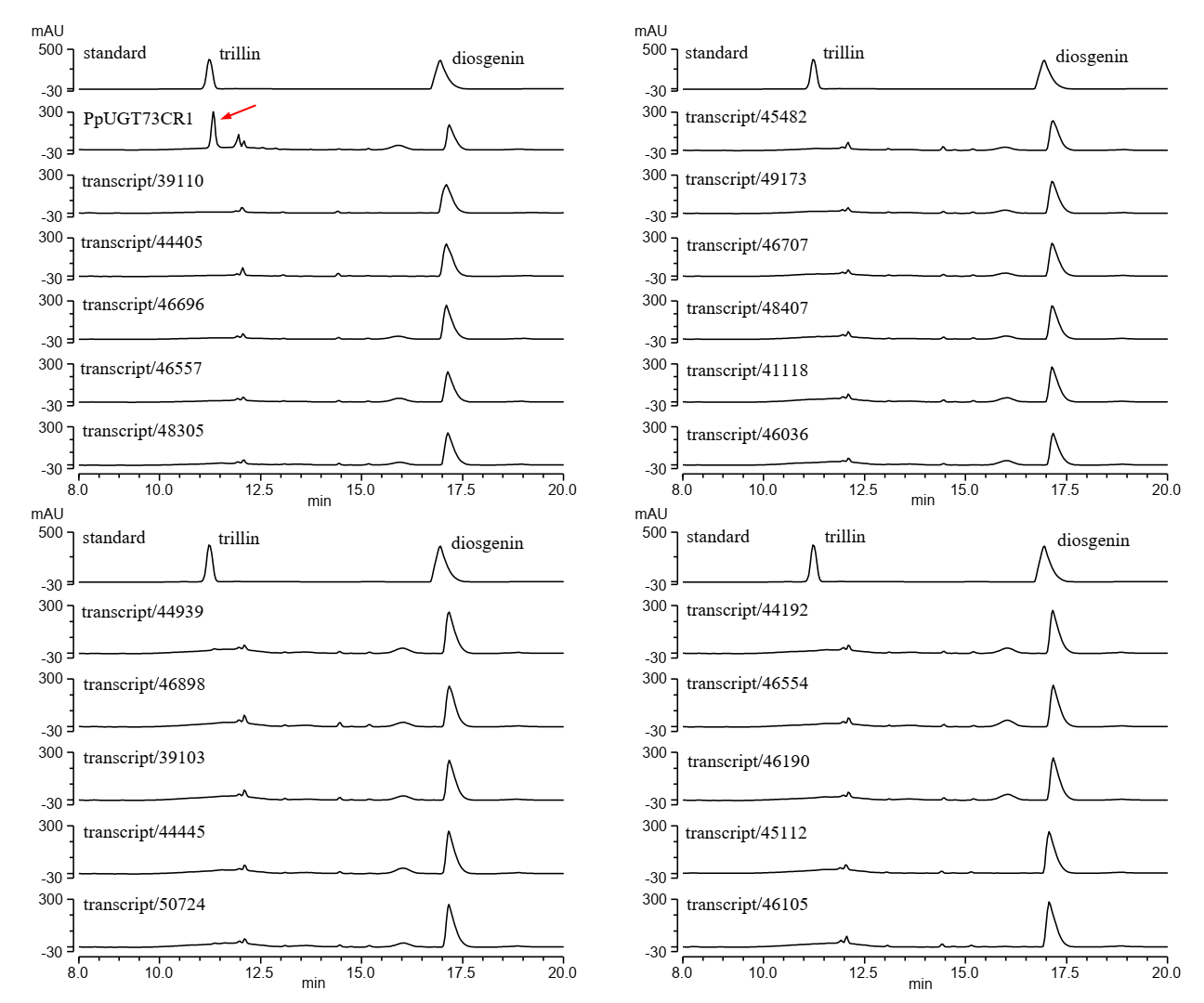

**Supplementary Figure 10.** HPLC analysis of diosgenin glycosylation catalyzed by candidate PpUGTs.

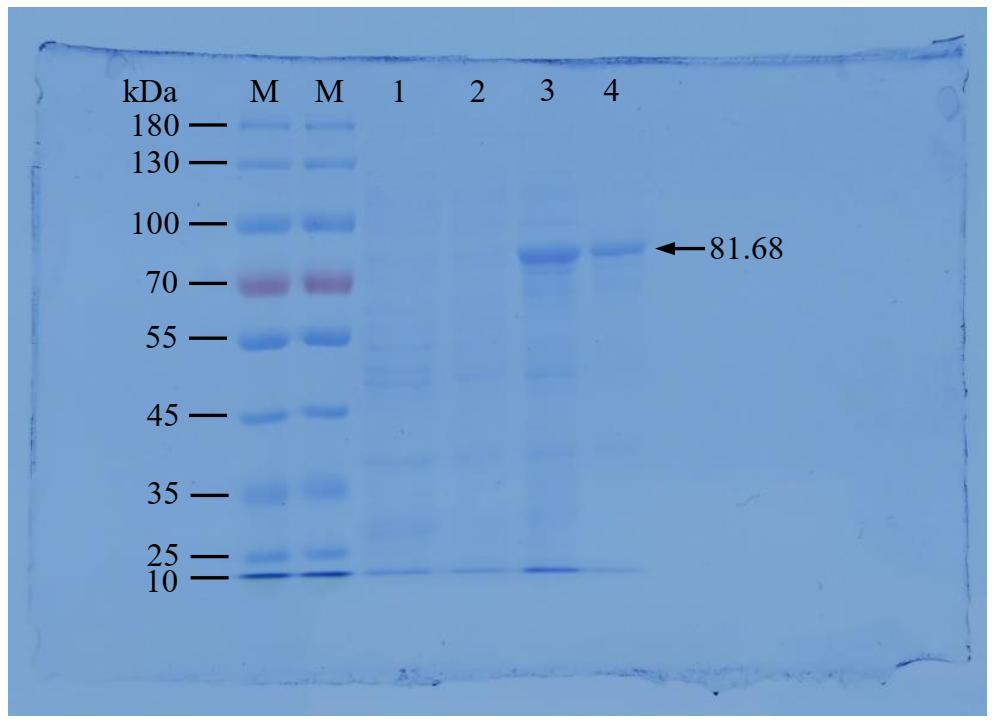

**Supplementary Figure 11.** SDS-PAGE analysis of expressed PpUGT73CR1 protein.

**Supplementary Table 1. OSC gene information of other plants in the OSC phylogenetic tree.**

| **No.** | **Species** | **GenBank ID** | **Product** |
| --- | --- | --- | --- |
| 1 | *Arabidopsis thaliana* | At3g45130 | Lanosterol |
| 2 | *Lotus japonicus* | AB244671 | Lanosterol |
| 3 | *Panax ginseng* | AB009031 | Lanosterol |
| 4 | *Abies magnifica* | AF216755 | Cycloartenol |
| 5 | *Adiantum capillus-veneris* | AB368375 | Cycloartenol |
| 6 | *Arabidopsis thaliana* | At2g07050 | Cycloartenol |
| 7 | *Avena strigosa* | AJ311790 | Cycloartenol |
| 8 | *Betula platyphylla* | AB055509 | Cycloartenol |
| 9 | *Betula platyphylla* | AB055510 | Cycloartenol |
| 10 | *Centella asiatica* | AY520819 | Cycloartenol |
| 11 | *Costus speciosus* | AB058507 | Cycloartenol |
| 12 | *Cucurbita pepo* | AB116237 | Cycloartenol |
| 13 | *Dioscorea zingiberensis* | AM697885 | Cycloartenol |
| 14 | *Eleutherococcus senticosus* | JQ400139 | Cycloartenol |
| 15 | *Glycyrrhiza glabra* | AB025968 | Cycloartenol |
| 16 | *Kalanchoe daigremontiana* | HM623872 | Cycloartenol |
| 17 | *Kandelia candel* | AB292609 | Cycloartenol |
| 18 | *Lotus japonicus* | AB181246 | Cycloartenol |
| 19 | *Luffa cylindrica* | AB033334 | Cycloartenol |
| 20 | *Maytenus ilicifolia* | KX147271 | Cycloartenol |
| 21 | *Oryza sativa* | AK121211 | Cycloartenol |
| 22 | *Panax ginseng* | AB009029 | Cycloartenol |
| 23 | *Pisum sativum* | D89619 | Cycloartenol |
| 24 | *Polypodiodes niponica* | AB530328 | Cycloartenol |
| 25 | *Rhizophora stylosa* | AB292608 | Cycloartenol |
| 26 | *Ricinus communis* | DQ268870 | Cycloartenol |
| 27 | *Cucurbita pepo* | AB116238 | Cucurbitadienol |
| 28 | *Siraitia grosvenorii* | HQ128567 | Cucurbitadienol |
| 29 | *Oryza sativa* | AK066327 | Parkeol |
| 30 | *Oryza sativa* | MG932696 | Orysatinol |
| 31 | *Catharanthus roseus* | JN991165 | α-Amyrin |
| 32 | *Eriobotrya japonica* | JX173279 | α-Amyrin |
| 33 | *Ilex asprella* | KM111167 | α-Amyrin |
| 34 | *Malus ×domestica* | FJ032006 | α-Amyrin |
| 35 | *Malus ×domestica* | FJ032008 | α-Amyrin |
| 36 | *Olea europaea* | AB291240 | α-Amyrin |
| 37 | *Arabidopsis thaliana* | At1g78950 | β-Amyrin |
| 38 | *Aralia elata* | HM219225 | β-Amyrin |
| 39 | *Artemisia annua* | EU330197 | β-Amyrin |
| 40 | *Aster sedifolius* | AY836006 | β-Amyrin |
| 41 | *Avena strigosa* | AJ311789 | β-Amyrin |
| 42 | *Betula platyphylla* | AB055512 | β-Amyrin |
| 43 | *Bruguiera gymnorrhiza* | AB289585 | β-Amyrin |
| 44 | *Bupleurum chinense* | EU400220 | β-Amyrin |
| 45 | *Euphorbia tirucalli* | AB206469 | β-Amyrin |
| 46 | *Gentiana straminea* | FJ790411 | β-Amyrin |
| 47 | *Glycyrrhiza glabra* | AB037203 | β-Amyrin |
| 48 | *Glycyrrhiza uralensis* | FJ627179 | β-Amyrin |
| 49 | *Hedera helix* | KU942522 | β-Amyrin |
| 50 | *Ilex asprella* | KM111168 | β-Amyrin |
| 51 | *Kalopanax septemlobus* | KT150523 | β-Amyrin |
| 52 | *Lotus japonicus* | AB181244 | β-Amyrin |
| 53 | *Medicago truncatula* | AJ430607 | β-Amyrin |
| 54 | *Nigella sativa* | FJ013228 | β-Amyrin |
| 55 | *Panax ginseng* | AB009030 | β-Amyrin |
| 56 | *Panax ginseng* | AB014057 | β-Amyrin |
| 57 | *Panax japonicus* | KP658156 | β-Amyrin |
| 58 | *Panax quinquefolius* | JX262290 | β-Amyrin |
| 59 | *Pisum sativum* | AB034802 | β-Amyrin |
| 60 | *Polygala tenuifolia* | EF107623 | β-Amyrin |
| 61 | *Solanum lycopersicum* | HQ266579 | β-Amyrin |
| 62 | *Theobroma cacao* | EOY26230 | β-Amyrin |
| 63 | *Tripterygium wilfordii* | KY885468 | β-Amyrin |
| 64 | *Vaccaria hispanica* | DQ915167 | β-Amyrin |
| 65 | *Betula platyphylla* | AB055511 | Lupeol |
| 66 | *Bruguiera gymnorrhiza* | AB289586 | Lupeol |
| 67 | *Cleome arabica* | KX897945 | Lupeol |
| 68 | *Glycyrrhiza glabra* | AB116228 | Lupeol |
| 69 | *Glycyrrhiza uralensis* | AB663343 | Lupeol |
| 70 | *Kalanchoe daigremontiana* | HM623871 | Lupeol |
| 71 | *Lotus japonicus* | AB181245 | Lupeol |
| 72 | *Malus ×domestica* | KT383436 | Lupeol |
| 73 | *Olea europaea* | AB025343 | Lupeol |
| 74 | *Ricinus communis* | DQ268869 | Lupeol |
| 75 | *Taraxacum officinale* | AB025345 | Lupeol |
| 76 | *Arabidopsis thaliana* | At5g42600 | Marneral |
| 77 | *Arabidopsis thaliana* | At5g48010 | Thalianol |
| 78 | *Aster tataricus* | AB609123 | Shionone |
| 79 | *Kalanchoe daigremontiana* | HM623869 | Glutinol |
| 80 | *Kalanchoe daigremontiana* | HM623870 | Friedelin |
| 81 | *Maytenus ilicifolia* | KX147270 | Friedelin |
| 82 | *Monteverdia ilicifolia* | MG677552 | Friedelin |
| 83 | *Monteverdia ilicifolia* | MG677553 | Friedelin |
| 84 | *Populus davidiana* | KY931453 | Friedelin |
| 85 | *Kalanchoe daigremontiana* | HM623868 | Taraxerol |
| 86 | *Luffa cylindrica* | AB058643 | Isomultiflorenol |
| 87 | *Panax ginseng* | AB265170 | Dammarenediol II |
| 88 | *Stevia rebaudiana* | AB455264 | Baccharis oxide |
| 89 | *Arabidopsis thaliana* | At4g15370 | Mixed products |
| 90 | *Arabidopsis thaliana* | At1g78955 | Mixed products |
| 91 | *Arabidopsis thaliana* | At1g78970 | Mixed products |
| 92 | *Arabidopsis thaliana* | At1g78960 | Mixed products |
| 93 | *Arabidopsis thaliana* | At1g66960 | Mixed products |
| 94 | *Arabidopsis thaliana* | At4g15340 | Mixed products |
| 95 | *Arabidopsis thaliana* | At5g36150 | Mixed products |
| 96 | *Arabidopsis thaliana* | At1g78500 | Mixed products |
| 97 | *Kandelia candel* | AB257507 | Mixed products |
| 98 | *Lotus japonicus* | AF478455 | Mixed products |
| 99 | *Malus ×domestica* | KT383435 | Mixed products |
| 100 | *Oryza sativa* | AK070534 | Mixed products |
| 101 | *Pisum sativum* | AB034803 | Mixed products |
| 102 | *Rhizophora stylosa* | AB263203 | Mixed products |
| 103 | *Rhizophora stylosa* | AB263204 | Mixed products |
| 104 | *Solanum lycopersicum* | HQ266580 | Mixed products |
| 105 | *Tripterygium wilfordii* | KY885467 | Mixed products |
| 106 | *Tripterygium wilfordii* | KY885469 | Mixed products |

**Supplementary Table 2. Experimental conditions of LC-MS-MS for determination of polyphyllin in *Paris polyphylla* var. *yunnanensis.***

|  | Parent Ion | Daughter Ion | Polarity | DP | EP | CE | CXP |
| --- | --- | --- | --- | --- | --- | --- | --- |
| Diosgenin | 413 | 269.1 | ESI- | -100 | -10 | -35 | -32 |
| Trillin | 575.2 | 119 | ESI- | -100 | -10 | -21 | -32 |
| Prosapogenin | 721.4 | 575.3 | ESI- | -100 | -10 | -10 | -32 |
| Polyphyllin I | 853.5 | 721.3 | ESI- | -100 | -10 | -40 | -32 |
| Polyphyllin II | 1013.9 | 721.6 | ESI- | -100 | -10 | -55 | -32 |
| Polyphyllin VI | 867.8 | 721.4 | ESI- | -100 | -10 | -30 | -32 |
| Polyphyllin VII | 736.8 | 119.1 | ESI- | -100 | -10 | -55 | -32 |
| Digitoxin | 763.5 | 503.3 | ESI- | -100 | -10 | -35 | -32 |

**Supplementary Table 3. Strains and plasmids used in functional verification of OSC gene.**

| **Plasmids and strains** | **Description** | **Reference** |
| --- | --- | --- |
| pδBLE2.0 | Yeast expression vector | Yuan et al.(2016) |
| pδHis | pδBLE2.0 derivative with HIS3 gene. | This study |
| pδHis-PpOSC1 | pδHis derivative with *PpOSC1* gene. | This study |
| pδHis-PpOSC2 | pδHis derivative with *PpOSC2* gene. | This study |
| OPC1 | One isolated variant derived from CEN.PK2-1C with an orthogonal cytosolic FPP biosynthetic pathway under the control of galactose-inducible promoter: *ERG10*, *ERG13*, *HMG1/2*, *ERG12*, *ERG8*, *ERG19*, *IDI1* and *ERG20* | Yuan et al.(2016) |
| BY-SQ1 | OPC1（gal80∆::ERG9） | Yuan et al. |
| SQ-PpOSC1 | One isolated variant derived from BY-SQ1 with a *PpOSC1* gene under the control of galactose-inducible promoter | This study |
| SQ-PpOSC2 | One isolated variant derived from BY-SQ1 with a *PpOSC2* gene under the control of galactose-inducible promoter | This study |

Jifeng Yuan and Chi-Bun Ching. Mitochondrial acetyl-CoA utilization pathway for terpenoid productions. Metabolic Engineering, 2016, 38, 303–309.

**Supplementary Table 4. Primer sequences for predicting UGT genes involved in the biosynthesis of polyphyllin.**

| **No.** | **Gene** | **Primer sequence** |
| --- | --- | --- |
| 1 | F01_transcript/39110 | F：GCCCCTGGGATCCCCGGAATTCATGGGCTCCGACGCGAAG  R：CGATGCGGCCGCTCGAGTCGACCTACTTTTCCTTCCGAAGCC |
| 2 | F01_transcript/44405 | F：GCCCCTGGGATCCCCGGAATTCATGGGCTCCGACGATCGTC  R：CGATGCGGCCGCTCGAGTCGACTCATCGTCGCTGGCCATGA |
| 3 | F01_transcript/46696 | F：GCCCCTGGGATCCCCGGAATTCATGGCGTCACAAACTCAACC  R：CGATGCGGCCGCTCGAGTCGACTTAGGATGAAACTGCTGGTAT |
| 4 | F01_transcript/46557 | F：GCCCCTGGGATCCCCGGAATTCATGGCGAAGCTCTTTGCCG  R：CGATGCGGCCGCTCGAGTCGACTTAGGGTGAAACTGCTGGTAT |
| 5 | F01_transcript/48305 | F：GCCCCTGGGATCCCCGGAATTCATGAGCTCCCTCCGCATCG  R：CGATGCGGCCGCTCGAGTCGACTCATTGATTTTTAGTGGCCAAA |
| 6 | F01_transcript/45482 | F：GCCCCTGGGATCCCCGGAATTCATGGATCTTCACGTCTTCTTC  R：CGATGCGGCCGCTCGAGTCGACTCAGACCTGGTCCTTGCTAG |
| 7 | F01_transcript/49173 | F：GCCCCTGGGATCCCCGGAATTCATGGGCTCCGTTCCGATAG  R：CGATGCGGCCGCTCGAGTCGACTTATTTGGGGTGATCGAACTG |
| 8 | F01_transcript/46707 | F：GCCCCTGGGATCCCCGGAATTCATGGGAGAGCAACCACTGAA  R：CGATGCGGCCGCTCGAGTCGACCTATACAAATTTTGTTGCCATA |
| 9 | F01_transcript/48407 | F：GCCCCTGGGATCCCCGGAATTCATGGCGTCACAAACTCAACC  R：CGATGCGGCCGCTCGAGTCGACTTAGGATGAAACTGCTGGTAT |
| 10 | F01_transcript/41118 | F：GCCCCTGGGATCCCCGGAATTCATGGATCTTCACGTCTTCTTC  R：CGATGCGGCCGCTCGAGTCGACTCACTTATTTGGGGTGATGGA |
| 11 | F01_transcript/43033 | F：GCCCCTGGGATCCCCGGAATTCATGCTGGCCCTCCTCCGC  R：CGATGCGGCCGCTCGAGTCGACTCAGTTGCGACCAGAACAAT |
| 12 | F01_transcript/46036 | F：GCCCCTGGGATCCCCGGAATTCATGCTACCGTCGTCACCAC  R：CGATGCGGCCGCTCGAGTCGACTTATTTGGGGTGATCGAACTG |
| 13 | F01_transcript/44939 | F：GCCCCTGGGATCCCCGGAATTCATGGCCCGCCTCCTCGCT |
|  |  | R：CGATGCGGCCGCTCGAGTCGACTTAGTCACCGCCACCATTC |
| 14 | F01_transcript/12407 | F：GCCCCTGGGATCCCCGGAATTCATGGGTGGCCTGAAGACGT |
|  |  | R：CGATGCGGCCGCTCGAGTCGACTTAGTCACCGCCACCATTCT |
| 15 | F01_transcript/46898 | F：GCCCCTGGGATCCCCGGAATTCATGGTGAGACACGCTAGTGG |
|  |  | R：CGATGCGGCCGCTCGAGTCGACTCATTGATTTTTAGTGGCCAA |
| 16 | F01_transcript/39103 | F：GCCCCTGGGATCCCCGGAATTCATGGAGTCCCAACCAGAGC |
|  |  | R：CGATGCGGCCGCTCGAGTCGACTCAATTGCGACCAGAACAATC |
| 17 | F01_transcript/44445 | F：GCCCCTGGGATCCCCGGAATTCATGGAGTCCCAACCAGAGC |
|  |  | R：CGATGCGGCCGCTCGAGTCGACTCAATTGCGACCAGAACAATC |
| 18 | F01_transcript/50724 | F：GCCCCTGGGATCCCCGGAATTCATGAGCTCCCTCCGCATCG |
|  |  | R：CGATGCGGCCGCTCGAGTCGACTCATTGATTTTTAGTGACCAAAT |
| 19 | F01_transcript/23323 | F：GCCCCTGGGATCCCCGGAATTCATGCTGGCCCTCCTCCGC |
|  |  | R：CGATGCGGCCGCTCGAGTCGACTCAATTGCGACCAGAACAATC |
| 20 | F01_transcript/44192 | F：GCCCCTGGGATCCCCGGAATTCATGGGTTCCGAAGCTCAACA |
|  |  | R：CGATGCGGCCGCTCGAGTCGACCTAAGTCAAATTTCGCATGTC |
| 21 | F01_transcript/46554 | F：GCCCCTGGGATCCCCGGAATTCATGGATGCCATTGGCCATCC |
|  |  | R：CGATGCGGCCGCTCGAGTCGACCTATTCTGGTAAGTTGTTATTCT |
| 22 | F01_transcript/46190 | F: GCCCCTGGGATCCCCGGAATTCATGCTGCCCGAATCCAGCA  R: CGATGCGGCCGCTCGAGTCGACCTATACAATCTTTGTTGCCATA |
| 23 | F01_transcript/48334 | F: GCCCCTGGGATCCCCGGAATTCATGCTCAAGTCCCAAATCCT  R: CGATGCGGCCGCTCGAGTCGACTCACCCTCCCCACCGC |
| 24 | F01_transcript/33044 | F: GCCCCTGGGATCCCCGGAATTCATGACCAACCAAACATCCATC |
|  |  | R: CGATGCGGCCGCTCGAGTCGACTTAACGCTCCATGACATATTG |
| 25 | F01_transcript/45112 | F: GCCCCTGGGATCCCCGGAATTCATGACCAAACAAACACATCCAA |
|  |  | R: CGATGCGGCCGCTCGAGTCGACCTATTCTGGTAAGTTGTTATTCT |
| 26 | F01_transcript/46105 | F: GCCCCTGGGATCCCCGGAATTCATGGGTTCCGAATCTCAGCC |
|  |  | R: CGATGCGGCCGCTCGAGTCGACTCAGGCTTCAGTTTTCGTAGT |

**Supplementary Table 5. Statistical table of transcriptome clean data of tissues in *Paris polyphylla* var. *yunnanensis*.**

| **Samples** | **Read Number** | **Base Number** | **GC Content** | **≥Q30** |
| --- | --- | --- | --- | --- |
| Fibr1YD | 49,181,509 | 14,677,368,556 | 51.43% | 94.36% |
| Fibr2YD | 48,722,996 | 14,530,314,100 | 50.98% | 93.85% |
| Fibr3YD | 47,968,347 | 14,279,858,192 | 51.23% | 94.41% |
| Frui1YD | 41,058,810 | 12,253,675,238 | 50.91% | 94.51% |
| Frui2YD | 41,408,373 | 12,360,642,172 | 50.45% | 94.35% |
| Frui3YD | 41,559,959 | 12,402,733,534 | 50.84% | 94.12% |
| Leaf1YD | 41,759,276 | 12,462,943,898 | 50.98% | 94.51% |
| Leaf2YD | 41,293,042 | 12,318,843,690 | 50.90% | 94.42% |
| Leaf3YD | 49,031,482 | 14,628,544,714 | 50.74% | 94.65% |
| Rhiz1YD | 40,972,867 | 12,240,818,562 | 51.59% | 94.10% |
| Rhiz2YD | 41,581,302 | 12,411,327,902 | 51.39% | 94.60% |
| Rhiz3YD | 41,955,499 | 12,528,751,962 | 51.80% | 94.63% |
| Stem1YD | 41,926,855 | 12,492,177,446 | 50.19% | 94.08% |
| Stem2YD | 40,316,576 | 12,039,632,420 | 50.13% | 94.03% |
| Stem3YD | 43,876,436 | 13,108,713,088 | 50.95% | 94.16% |
| AB-2 | 41,208,394 | 12,316,381,062 | 51.71% | 94.15% |
| AB-3 | 62,353,157 | 18,617,620,552 | 52.02% | 94.43% |
| AY-2 | 58,423,460 | 17,456,235,974 | 52.19% | 94.27% |
| AY-3 | 61,158,841 | 18,269,204,240 | 52.37% | 94.27% |
| AZF-2 | 56,419,925 | 16,873,584,024 | 52.47% | 94.15% |
| AZF-3 | 48,235,606 | 14,415,014,318 | 51.52% | 94.38% |

Read Number：Total number of pair-end Reads in Clean Data；Base Number：The total number of bases in Clean Data；GC Content：The percentage of G and C bases of total bases in Clean Data；≥Q30：The percentage of bases with a quality value of ≥ 30

**Supplementary Table 6. The number of genes involved in polyphyllin biosynthesis pathway in WGCAN module.**

| **Modules** | **AACT** | **HMGS** | **HMGR** | **MVK** | **PMK** | **MPDC** | **Ipl** | **GPPS** | **FPS** | **TPS** | **SQS** | **SQE** | **CAS** | **SSR2** | **SMO3** | **CPI** | **CYP51** | **C14-R** | **8,7 SI** | **SMO4** | **C5-SD2** | **7-DR2** | **Totle** |
| --- | --- | --- | --- | --- | --- | --- | --- | --- | --- | --- | --- | --- | --- | --- | --- | --- | --- | --- | --- | --- | --- | --- | --- |
| **aliceblue** | 0 | 1 | 1 | 0 | 1 | 0 | 1 | 0 | 0 | 3 | 3 | 0 | 2 | 1 | 2 | 0 | 0 | 1 | 2 | 1 | 1 | 0 | **20** |
| **palevioletred2** | 0 | 4 | 0 | 0 | 0 | 1 | 1 | 0 | 0 | 5 | 1 | 1 | 2 | 0 | 1 | 0 | 0 | 0 | 0 | 1 | 0 | 0 | **17** |
| **coral** | 0 | 1 | 1 | 0 | 0 | 0 | 0 | 0 | 0 | 0 | 0 | 2 | 0 | 7 | 1 | 0 | 0 | 1 | 0 | 1 | 0 | 0 | **14** |
| **antiquewhite4** | 2 | 3 | 3 | 1 | 0 | 1 | 0 | 0 | 0 | 2 | 0 | 2 | 1 | 3 | 2 | 0 | 0 | 0 | 0 | 1 | 0 | 0 | **21** |
| **lavenderblush3** | 0 | 1 | 1 | 0 | 0 | 0 | 0 | 0 | 2 | 4 | 1 | 0 | 0 | 0 | 0 | 0 | 0 | 0 | 0 | 0 | 0 | 2 | **11** |
| **antiquewhite1** | 0 | 0 | 0 | 0 | 0 | 1 | 0 | 0 | 0 | 1 | 0 | 0 | 0 | 1 | 1 | 0 | 0 | 0 | 1 | 0 | 0 | 1 | **6** |
| **white** | 0 | 0 | 0 | 1 | 0 | 0 | 0 | 0 | 0 | 1 | 0 | 1 | 0 | 1 | 0 | 0 | 0 | 0 | 0 | 0 | 0 | 0 | **4** |
| **salmon2** | 0 | 1 | 0 | 0 | 0 | 0 | 0 | 0 | 0 | 1 | 1 | 0 | 0 | 1 | 0 | 0 | 0 | 0 | 0 | 2 | 0 | 0 | **6** |
| **honeydew** | 1 | 1 | 0 | 0 | 2 | 0 | 0 | 0 | 0 | 1 | 0 | 0 | 0 | 0 | 0 | 0 | 0 | 0 | 1 | 0 | 2 | 1 | **9** |
| **lightcoral** | 0 | 0 | 0 | 0 | 0 | 1 | 0 | 0 | 0 | 0 | 0 | 0 | 0 | 0 | 0 | 0 | 0 | 0 | 0 | 0 | 0 | 0 | **1** |
| **blue4** | 0 | 0 | 0 | 0 | 0 | 0 | 0 | 0 | 0 | 0 | 1 | 0 | 0 | 0 | 0 | 0 | 0 | 0 | 0 | 0 | 0 | 0 | **1** |
| **darkslateblue** | 0 | 0 | 0 | 0 | 0 | 0 | 0 | 0 | 1 | 0 | 0 | 0 | 0 | 0 | 0 | 0 | 0 | 0 | 0 | 0 | 0 | 0 | **1** |
| **plum4** | 0 | 0 | 0 | 0 | 0 | 0 | 0 | 0 | 0 | 0 | 0 | 0 | 0 | 0 | 0 | 0 | 0 | 0 | 0 | 0 | 0 | 0 | **0** |
| **antiquewhite2** | 0 | 0 | 0 | 0 | 0 | 0 | 0 | 0 | 0 | 2 | 1 | 0 | 0 | 0 | 0 | 0 | 0 | 0 | 1 | 0 | 0 | 0 | **4** |
| **mediumpurple** | 0 | 0 | 0 | 0 | 0 | 0 | 0 | 0 | 0 | 3 | 0 | 0 | 0 | 0 | 0 | 0 | 0 | 0 | 0 | 0 | 0 | 0 | **3** |
| **lightblue3** | 0 | 0 | 0 | 0 | 0 | 0 | 0 | 0 | 0 | 0 | 0 | 0 | 0 | 0 | 0 | 0 | 0 | 0 | 0 | 0 | 0 | 0 | **0** |
| **chocolate3** | 0 | 1 | 0 | 0 | 0 | 0 | 0 | 0 | 0 | 0 | 0 | 0 | 1 | 0 | 0 | 0 | 0 | 0 | 0 | 0 | 0 | 0 | **2** |
| **palevioletred** | 0 | 0 | 0 | 0 | 0 | 0 | 0 | 0 | 0 | 0 | 0 | 0 | 0 | 0 | 0 | 0 | 0 | 0 | 0 | 0 | 0 | 0 | **0** |
| **plum1** | 0 | 0 | 0 | 0 | 0 | 0 | 0 | 0 | 0 | 0 | 0 | 0 | 0 | 0 | 0 | 0 | 0 | 0 | 0 | 0 | 0 | 0 | **0** |
| **cornflowerblue** | 0 | 1 | 0 | 0 | 0 | 0 | 0 | 0 | 0 | 0 | 0 | 0 | 1 | 0 | 0 | 0 | 0 | 0 | 0 | 0 | 0 | 0 | **2** |
| **pink2** | 0 | 0 | 0 | 0 | 0 | 0 | 1 | 0 | 0 | 0 | 0 | 0 | 0 | 0 | 0 | 0 | 0 | 0 | 0 | 0 | 0 | 0 | **1** |
| **orangered4** | 0 | 0 | 0 | 0 | 0 | 0 | 0 | 0 | 0 | 0 | 0 | 0 | 0 | 0 | 0 | 0 | 0 | 0 | 0 | 0 | 0 | 0 | **0** |
| **blueviolet** | 0 | 0 | 0 | 0 | 0 | 0 | 0 | 0 | 0 | 0 | 0 | 0 | 0 | 0 | 0 | 0 | 0 | 0 | 0 | 0 | 0 | 0 | **0** |
| **salmon4** | 0 | 0 | 0 | 0 | 0 | 0 | 0 | 0 | 0 | 0 | 0 | 0 | 0 | 0 | 0 | 0 | 0 | 0 | 0 | 0 | 1 | 0 | **1** |

**Supplementary file 2**

OSC:

>F01_transcript/16440 full_length_coverage=3;length=2651;num_subreads=46(OSC1)

GTGTTCGGTTCCCCTCACTCTTTGACAGCACTGAAAACAGAGAGAGGGAGCGCTCCAATACCCTAAAGAACCCTAATTCAGATTCTCTATTTTGACATCCTCTCATTGAATTTGGAATCTTTAAGCTATCCCTCTTGGCGCGCTAATGTGGAAGCTGAAGATCGCGAATGGCGGCAGCCAGTGGCTGCGGACAAACAACAAACACGTTGGGAGGCAGACGTGGGAGTTCGATCCCAACCTCGGGACGCCGGAGGAGATCGAAGCAGTCGAGAAGGCCCGCCGGGAATTCACGGAACACCGCTTCGAGAAGAAGCACAGCGCTGATTTGCTCATGCGGATGCAGTTTGCGAAGGAGAATCCGGTTGAAATAAATCTTCCTCATATCAAGCTCGAAGAGCACGAAGATGTTACGGAGGAGGCCGTATTGACAACATTAAGAAGGGCCATCACCCGCCACTCTACTCTCCAAGCTCATGATGGACACTGGCCCGGGGATTATGGAGGTCCTATGTTCCTTATGCCTGGCTTGATTATAGCTTTATATGTTACGGGGGCAATAAATAGTGTTCTTTCATCGGAGCACCGGAGAGAGATTCGCCGATACCTCTACAACCATCAGAACAAGGATGGCGGCTGGGGTTTGCATATTGAGGGCCAAAGCACCATGTTTGGTTCATCCTTGACCTATGTGACACTCAGGTTGCTTGGAGAAGGAGCTGAATGTGGTGACGGGGCTATGCAGAAAGGGCGGAAGTGGATTTTAGACCATGGTGGAGCAACTGCTATAACATCGTGGGGAAAATTTTGGCTCTCGGTACTTGGAGTATTTGACTGGTCTGGCAACAATCCATTACCCCCAGAAATGTGGTTACTTCCATATTTTCTCCCAGTGCATCCAGGAAGAATGTGGTGCCATTGCCGTATGGTTTATTTGCCCATGTCATATTTATACGGCAAGAGATTTGTGGGTCCAATTACGCCAACAATTTTAGAGTTGAGGAAGGAGATATTTATTCAACCATATAACAAAATTGATTGGAATCTGGCTCGTAATGAATGTGCAAAGGAAGACCTGTACTACCCTCATCCCATAATACAGGATATCCTTTGGGCTTCCCTTCATAAGGTTGCTGAGCCTATTCTGTTGCATTGGCCTGGTAGTAAGCTGAGAGAGAAGGCTCTGACTACTGCCATACAGCACATTCATTATGAAGACGAGAACACTCGATATGTTTGCATTGGTCCTGTGAACAAGGTACTAAATATGCTTTGCTGCTGGGCGGAGGATCCAAATTCTGAGGCATTCAAGCTGCACCTTCCACGAATCTTTGATTATTTATGGCTCGCTGAAGATGGCATGAAGATGCAGGGATATAATGGCAGCCAATTGTGGGACACAGCTTTTACTGTTCAAGCTATTATTGCAACTAATCTTTCTGAAGAATTTGGGCCTACTCTTAGAAATGCACATGAATACATAAAGAATACACAGGTTCTTGACAACTGTCCTGGCGACCTTTCTTTCTGGTATCGTCACATCTCTAAAGGTGCGTGGCCCTTTTCAACTGCGGATCATGGATGGCCAATTTCCGATTGCACAGCAGAAGGATTAAAGGCAGCACTTCTGTTATCGAGAATTTCCCCAAAAAGTGTTGGTGATCCAATTGATGCAAAGTGGCTTTACGAGGCTGTGAATGTCATTCTTTCCTTAATGAACAAGGACGGTGGTTTTGCTACATATGAACTTACAAGATCATATGCATGGTTGGAGCTTATCAATCCTGCTGAAACTTTTGGGGACATTGTTATCGACTATCCATATGTCGAATGCACTTCAGCATCAATTCAAGCATTGACATCATTTAAAAAATTATACCCTGGACATCGCAGAGAAGAAATTGAAAGCTGCATAAAAAAGGCAGCTCATTTCATTGAAAGTATACAACGACCTGATGGTTCATGGTATGGCTCTTGGGCTGTTTGCTTCACATATGGCATATGGTTTGGAGTCCTGGGACTGATTGCTGCAGGAAGGACGTACAAGAATAGCTCTACCATTCGTAAAGCATGTGATTTTCTGTTGTCAAAACAGTTGCCTTCTGGTGGTTGGGGAGAGAGCTATCTCTCATGTCAAGATAAGGTTTACACAAATCTTGAAGGCAACCGGTATCATGCTGTGAATACTGGGTGGGCTATGTTAACTCTAATTGATGCTGGACAGGCTGATAGAGATCCAAAGCCTCTGCACCGTGCAGCAAAGGCATTGATTAACTTGCAAATGGAGAATGGTGAATATCCGCAACAAGAAATCATGGGAGTGTTCAACAGGAACTGCATGATAAGCTACTCAGCATATCGCAATATTTTCCCGATATGGGCTCTCGGAGAATATCGCAATCGGGTCCTTTGCCCTTCCCAGAAACAGTGAGCTCGCTCTTCAGCATTTTATGCCCTTTTTACACATTGGTCCATCTGTTCCAACAACCACTGTCATCTCTCTGCGCCAAACCTTGAACAGATGAAGAAAAAAAAGGCATGAAGGGTCACCAGTCCATTGGGCTTACATTTTCACTTTCTCTCAAGTAGATTAAAAAGACTTACGTCCTGACCTTGAGAATTCTTGTGGCTGGTGCTTATTTATTGTTTTTTAC

>F01_transcript/12661 full_length_coverage=2;length=2831;num_subreads=59(OSC2)

GGGTTCCCCTCACTCTTTGACAGCACTGAAAACAGAGAGAGAGAGAGAGGGAGCGCTCCAATACCCTAAAGAACCCTAATTCAGATTCTCTATTTTGACATCCTCTCATTGAATTTGGAATCTTTAAGCTATCCCTCTTGGCGCGCTAATGTGGAAGCTGAAGATCGCGAATGGCGGCAGCCAGTGGCTGCGGACAAACAACAAACACGTTGGGAGGCAGACGTGGGAGTTCGATCCCAACCTCGGGACGCCGGAGGAGATCGAAGCAGTCGAGAAGGCCCGCCGGGAATTCACGGAACACCGCTTCGAGAAGAAGCACAGCGCTGATTTGCTCATGCGGATGCAGTTTGCGAAGGAGAATCCGGTTGAAATAAGTCTTCCTCATATCAAGCTCGAAGAGCACGAAGATGTTACGGAGGAGGCCGTATTGACAACATTAAGAAGGGCCATCACCCGCCACTCTACTCTCCAAGCTCATGATGGACACTGGCCCGGGGATTATGGAGGTCCTATGTTCCTTATGCCTGGCTTGATTATAGCTTTATATGTTACGGGGGCAATAAATAGTGTTCTTTCATCGGAGCACCGGAGAGAGATTCGCCGATACCTCTACAACCATCAGAACAAGGATGGCGGCTGGGGTTTGCATATCGAGGGCCAAAGCACCATGTTTGGTTCATCCTTGACCTATGTGACACTCAGGTTGCTTGGAGAAGGAGCTGAATGTGGTGACGGGGCTATGCAGAAAGGGCGGAAGTGGATTTTAGACCATGGTGGGGCAACTGCTATAACATCGTGGGGAAAATTTTGGCTCTCGGTACTTGGAGTATTTGACTGGTCTGGCAACAATCCATTACCCCCAGAAATGTGGTTACTTCCATATTTTCTCCCAGTGCATCCAGGAAGAATGTGGTGCCATTGCCGTATGGTTTATTTGCCCATGTCATATTTATACGGCAAGAGATTTGTGGGTCCAATTACGCCGACAATTTTAGAGTTGAGGAAGGAGATATTTATTCAACCATATAACAAAATTGATTGGAATCTGGCTCGTAATGAATGTGCAAAGGAAGACCTGTACTACCCTCATCCCATAATACAGGATATCCTTTGGGCTTCCCTTCATAAGGTTGCTGAGCCTATTCTGTTACATTGGCCTGGTAGTAAGCTGAGAGAGAAGGCTCTGACTACTGCCATACAGCACATTCATTATGAAGACGAGAACACTCGATATGTTTGCATTGGTCCTGTGAACAAGGTGCTAAATATGCTTTGCTGCTGGGCGGAGGATCCAAATTCTGAGGCATTCAAGCTGCACCTTCCACGAATCTTTGATTATTTATGGCTCGCTGAAGATGGCATGAAGATGCAGGGATATAATGGCAGCCAATTGTGGGACACAGCTTTTACTGTTCAAGCTATTATTGCAACTAATCTTTCTGAAGAATTTGGGCCTACTCTTAGAAATGCACATGAATACATAAAGAATACACAGGTTCTTGACAACTGTCCTGGCGACCTTTCTTTCTGGTATCGTCACATCTCTAAAGGTGCGTGGCCCTTTTCAACTGCGGATCATGGATGGCCAATTTCCGATTGCACATCAGAAGGATTAAAGGCAGCACTTCTGTTATCAAGAATTTCCCCAAAAAGTGTTGGTGATCCAATTGATGCAAAGTGGCTTTACGAGGCTGTGAATGTCATTCTTTCCTTAATGAACAAGGACGGTGGTTTTGCTACATATGAACTTACAAGATCATATGCATGGTTGGAGCTTATCAATCCTGCTGAAACTTTTGGGGACATTGTTATAGACTATCCATATGTCGAATGCACTTCAGCATCAATTCAAGCATTGACATCATTTAAAAAATTATACCCTGGACATCGCAGAGAAGAAATTGAAAGCTGCATAAAAAAGGCAGCTCATTTCATTGAAAGTATACAACGACCTGATGGTTCATGGTATGGCTCTTGGGCTGTTTGCTTCACATATGGCATATGGTTTGGAGTCCTGGGACTGATAGCTGCAGGAAGGACGTACAAGAGTAGCTCTACCATTCGTAAAGCATGTGATTTTCTGTTGTCAAAACAGTTGCCTTCTGGTGGTTGGGGAGAGAGCTATCTCTCATGTCAAGATAAGGTTTACACAAATCTTGAAGGCAACCGGTATCATGCTGTGAATACTGGGTGGGCTATGTTAACTCTAATTGATGCTGGACAGGCTGATAGAGATCCAAAGCCTCTGCACCGTGCAGCAAAGGCATTGATTAACTTGCAAATGGAGAATGGTGAATATCCGCAACAAGAAATCATGGGAGTGTTCAACAGGAACTGCATGATAAGCTACTCAGCATATCGCAATATTTTCCCGATATGGGCTCTGGGAGAATATCGCAATCGGGTCCTTTGCCCTTCCCACAAACAGTGAGCTCGCTCTTCAGCATTTTATGCCCTTTTTACACATTCTTCCATCTGTTCCAACAACCACTGTCATCTCTCTGCGCCAAACCTTGAACAGATTAAGAAAAAAAAGGCATGAAGGGTCACCAGTCCATTGGGCTTACATTCACTTTCTCTCAAGTAGATTAAAAGGACTTTTACGTCCTGACCTTGAGAATTCTTGTGGCTGGTGCTTATTTATTGTTATATATATATATATATTGTGTTTATTTTGCTGCGAAGACGAATTCAATTTCATCTAATAGACCTAGCTTGAGCTGTAATTTAGTTGTCAATACTGAGTAATTATATTTGGAATGCAACAGTTTGGTTAAATGAGTAAAATCTGCACATTGCTATAAATCGGACAGTACAATTTAAAAGCACTC

CYP:

>F01_transcript/29047 full_length_coverage=2;length=2212;num_subreads=60

GGCTGTTCGACACTCCAAACTGTCCAAACCCAACGTTGATGGCGTTGAGGCTTCATCTCCTCGCCTTCCTCTCATTCCTCTTCTCCCCCATTCTGCTGCAACTCATCAATGGCGTCCGCTGCCGTCACCGCCGCCGTCTCCCCTTCGCCCTGCCCCCCTCAGCTCCCGTCCAAATCCCATCTCCGGCGAGCCGCCGCCCTCTCCATCCTCTCTCGCGGCGGCGCCGTACGCTGCTCCGCCTCATCCAACGGCCGAAACCCTATCCCCGGCGGCGATCAGTCCTCCAAGGACGCCGATCGCCTCATCGAGAAGAAGCGCCGCTCCGACCTCGCCGACCGCATCGCCTCCGACGAGTTCACCGTCCAGCAATCCAGGTTGGTTTCTCTGTTGCGGAGGCTTGGACCGCCGGGGGAATTCTTGGCGGAGCTTCTGTCGAGGTCGGAGATTCCTCAGGCGAGGGGAGAGATCAGCTCCGTTGGCCGGACGGCCTTCTTCATCCCCCTTTACGAGCTTTTCCTCAGATACGGGGGGATCTTCCGCCTCACCTTCGGCCCCAAGTCCTTTCTGATTGTTTCCGATCCTGCTATCGCCAAGCACATACTCAAGGACAACTCCAAGGCTTATTCTAAGGGTATCCTAGCAGAAATTCTCGAGTTTGTTATGGGAAAGGGTTTGATCCCAGCCGATGGTGAAATCTGGCGTGTCCGAAGACGGGCGATTGTCCCGGCATTGCATCAGAAGTACGTGTCTGCCATGATCGGCCTCTTTGGAAAAGCTTCATATCGGCTATGTGAGAAGTTAGATGCCGCAGCAACCGATGGAGAGGATGTCGAGATGGAATCCCTCTTCTCGCGGTTGACACTTGATATCATTGGCAAGGCCGTCTTTAATTATGATTTTGATTCCTTATCACATGATAATGGAATAGTCGAGGCAGTTTACACTGTATTGCGGGAGGCAGAGCAACGGAGTACTTCTCCAATACCAACTTGGGAAATTCCTTTATGGAAGGATATATCTCCAAGGCAGAAGAAGGTCAATGTAGCTCTTAAGTTGATAAATGACACCCTTGATGATTTGATTGCTATCTGCAAGAGAATAGTAGATCAAGAGGAGTTGCAATTTCATGAAGAGTACATGAATGAGCAAGACCCAAGCATTCTTCACTTTTTATTGGCATCAGGGGATGACGTTTCTAGCAAGCAGCTTCGTGATGATTTGATGACTATGCTTATAGCCGGCCATGAAACATCTGCAGCGGTGCTAACATGGACCTTTTATCTTCTTTCTAAGGAACCAAGAGTCATGGCCAAGCTCCAAGATGAGGTTGACTCTGTTCTAGGAGACAGAGTTCCAACCATTGAAGATGTCAAGAAACTGAAGTATACTACTCGAGTGATCAACGAATCACTGAGACTCTACCCACAACCACCAGTCTTAATTCGCCGTTCTCTTGAGAATGATGTGCTCGGGAAGTACCCTATCAAAAGGGGTGAAGATATTTTCATTTCTTTATGGAACCTCCATCGTTGTCCTGATCATTGGGTTGATGCCGAGGGTTTCAACCCTGAAAGATGGCCTCTAGATGGACCAAATCCAAATGAGACTAATCAGAATTTCAGCTATTTACCTTTCGGTGGTGGGCCAAGGAAATGCGTTGGAGACATGTTTGCTTCATTTGAGACCGTGGTGGCGACATCGATGTTGGTGAGACGATTCAACTTCCAATTAGCCATCACGGCGCCTCCTAAGAGGAAAAAAAAAAAGAAACCAATATTGGTTTCTCCTTGAGAAGCGGATTATACAGGAAAACAGCTGGGACTAGCTTCTGCTCGTAGCTTTTACATGAAGCAGCTGCTATAACAGCTTAATTGACTAGAGATGTGTTGAGTGATGCAGGTGGAAATGACGACAGGAGCAACCATTCACACGACGGAGGGGTTGGTGATGACAGTAACACCAAGGATGCAGCCTCCGATAATTCCCTCGCTGGGGTTGAAGCGAGTGGTAACAGTAGAAAGTGATCAGCCTCAAAACTGGTCTCCCCCAGCTCCATTTCGAATCGGATAGCAGCCGCCTCATTTGATGCACTTGCCATCCATCTCTGGCAATGTACATTTTCTCGCAACTTGTCTTTGTTATATTTCATGGAGAAATTAGCAAGCATGATATGATCAATTCCACTTCAGAGAAATTTCTTTCTAATCCTCAC

>F01_transcript/33317 full_length_coverage=8;length=2144;num_subreads=60

GGGAAAAAAAAATAAACTGTTTGAATTTTTTTTAAAAAAAGTTTCCTATAGTGGTCTCTCTCTCTCTCTCTCTCTCTCTCTCTCTTTTCTCGCTCTCTTATCTCTTCCGGACCTCTCCACTAACTCTCCTTCCTATTGTGTTTTGTGTTGGGTTGGCACAAAAGAGCTCTGGTATCGGCCATGCATTCCGTGGCTCTGTTGCAGCTCCCAGCGGCCACCAATGGACTCAATTTCCCCCAAATTTGTTCCTCTAAAACCTCAACTCCCTTCACTGTTGGCTCCTCACATCCTTTTTCAACAACGAGACGCTCGAAAACCAGATGCCAGTCGGCTAATACTGAAGGACCAAAGACAAAGTGGAATATGCTCGACAATGCAAGCAACATTCTTACTAATTTTTTAAGTGGGGGTAGCCTTGGTTCAATGCCGACTGCTGAAGGTGCTGTCTCTGATCTGTTTGGCCGGCCTCTTTTCTTTTCTCTTTATGATTGGTTTTTAGAGCATGGTTCTGTTTACAAGCTTGCTTTTGGACCGAAAGCGTTTGTTGTCGTATCGGATCCTGTTGTTGCGAGACATATTCTTCGAGAAAACTCATATTCTTATGACAAGGGAGTTCTTGCTGATATCCTAGAACCAATAATGGGGAAAGGACTCATACCTGCAGATCTTGAAACTTGGAAGCTCCGGAGGAGAGTAATTGCACCTGGGTTTCATTCTTTGTTCCTGGAAGCAATGGTCAACGTTTTCTCCAATTGTTCTGACAGAACTGTTTCAAAGTTTGAGATGCTTATTGAAAGGGAGAACAATGATGAAAAATCAATTGAGTTGGATCTTGAAGCAGAATTTTCAAGTTTGGCACTTGACATTATTGGGCTGGGTGTATTTAACTATGACTTTGGATCTGTCACTAAGGAATCTCCTGTGATCAAGGCTGTCTATGGGACTCTTTTTGAAGCTGAACATCGTTCAACCTTTTACATTCCTTATTGGAAGCTTCCTTTCTCAAGATGGTTAGTTCCAAGGCAACGTAAGTTTCACGATGATCTCAAGGTTATCAATGACTGCCTTGATGGGCTTATTAGAAATGCAAAAGAGACTAGGCAGGAAACTGATGTTGAAAAACTTCAACAAAGAGATTATTCAAGCCTGAAGGATGCAAGTTTACTGCGTTTTTTTGTTGATATGAGAGGTGCTGATGTTGATGACCGCCAGCTTAGGGATGACCTCATGACCATGCTTATTGCTGGACATGAAACAACTGCTGCTGTTCTGACATGGGCTATTTTCCTACTTGCCCAGAACCCCTCCAAAATGAAAAAGGCTCAGGCAGAGGTTGATTCTGTTTTTGATCAACAGAGGATAACTTTGGAGTGCATTAAAAAATTGAAGTACATACGTCTTATCATTGTTGAAGCTCTCCGTCTATATCCACAACCTCCCATACTTATTAGACGTGGACTTAAAGCTGATATACTTCCAGGAGGATATAAAGGTCATAGAGATGGATATAAAATTCCTGCTGGGACTGATATCTTTATCTCTGTTTACAACCTTCACAGATCACCGTATTTTTGGGACCGGCCTCATGAATTTGAGCCAGAGAGATTCTTAGTTTCGAAGAAGAGCGAAGGGATTGAAGGATGGGCGGGATTTGATCCGAGCCGGAGTCCTGGTTCCTTGTATCCTAATGAGATAGTCTCGGACTTTTGCCTTCCTGCCGTTTGGTGGGGGACCTCGCAAGTGCGTTGGAGACCAGTTTGCGCTCTTGGAATCGACCGTAGCATTGGCCGTGCTGATTCAGAAGTTCGATGTGGAGTTGAGAGGATCTGCAGATGATGTCGAGCTCGTCACGGGGGCCACCATTCATACCAAGAATGGTCTGTGGTGCAAAGTGAAGAAAAGGACTCGAAAGGAACAAAGTGAGCCTAAATCTTGAATTGATGAAGGTTATGGACTGTGAAGTAGCTTCTCTTTCAGGTTCAACACTGAAAGAATTGAGAAATGTTGTTTGCAAAATCATTCACCAACCCCCAGGGTTTTTTTTTTTTTTTTTTTCCAGACAGTTTCATTGTGTAAATGTATCAAGCTTTACTTAAGTTTACAGGAGAATGGTTCCCTATTATTTGGCCACACTAATTTCCACC

>F01_transcript/34222 full_length_coverage=6;length=1841;num_subreads=60

ACTCCAGCAAAAACCTTGGAGAGAGAGAGAGAGAGTCCAATGGAGCTGCAGTATGGTCTTGATTGGTTTTCAGGGATTCTTGTTCTTCTGGGGCTCTACTGTATCGGATGGAGCTGGATGGGAAAGAGCAAGAAGCAGTATCCACCAATTGCCGGCAGTATCTTCCACCAGTTCTTGAACCTCCCAAGGCTCCACGACTACCACACCAAACTCTCTCACAAGTACAGAACCTTCAGGCTGTATGCTCCCTTTCGGCACCACATTTACACTGTGGATCCAACCAACATCGAGCACATCCTCCAGACCAACTTCAACAACTATGGCAAGGGGCAGTACAATTACGTGAACACTGCGGAACTATTTGGAGATGGGATTTTCGCGGTGGACGGCGAGAAGTGGCGCCATCAGCGGAAGCTTGCGAGCTATGAGTTCTCCACCAAAGTTTGGAGGGATTTCAGTGCCACTGTCTTCAAAACAACTGCAAGCAAACTCGCTCAAATCATTTCTGCTGCTGCATCATCCAATCAAACAATAGATATTCAGGACTTGTTCATGAAATCGACAATGGACTCTATTTTCAAAGTTGGATTTGGCACAGAATTGGATTGTCTTGGAGGATCAGACGAGGGGAGCAAATTTGCAAAGGCATTCGATACCTCGAGCGAACTTATTACTCGGCGTTATGTTCACGCCTTTTGGAAGATCGAAAGGTTTCTAAAGTTCGGTCCGGAGGCCGAACTGAGGAAGAACATTAAAGACATCAATGACTTTGTCTACAAGCTGATTAGTACAAAAACGGAACAAATACCTATCGAAAGAAGTGATTCGGGGAAAAAAGAGGACATCTTGTCTAGGTTTCTTGTGGAAATGGATAAAGATCCGGAGAACATGACACACAAATACTTAAGGGACATTGTACTCAATTTCCTAATCGCCGGCAAAGACACCACAGCAGGAACCCTAGCATGGTTCATCTACATGCTCTGCAAGCACCCTTCAGTACAAGAAAAGGTCGCCAAAGAAGTCAAGGAATCGATTGAATCCAACGAGGGCATAACTACCAACGACTTCTCCAAATGTATCACCGACGAAGCCCTCAATAAGATGCAATACCTCCATGCAGCCTTGTCTGAAACTCTGAGGTTGTACCCTGCTGTTCCCTTGGATAATAAGGTGTGTTTAGCAGATGACACCTTGCCAGATGGCTCCAATGTGAGGAAAGGAGACATTGTATTCTACCAACCTTATGCTATGGGTAGAATGGAATTCTTGTGGGGTCCTGATGCCGAAATCTTTCGGCCAGAACGGTGGATCGACGATCAAGGGTTTTTTCACCCCGAAAGCCCTTTTAAGTTCACAACTTTTCAGGCTGGGCCACGGATTTGTTTAGGTAAGGAGTTTGCCTACAGGCAAATGAAGATCTTTGCCGCCATTCTCCTCCGATTCTTTGTATTCAAACTCGCAGACGAGACAGCGGACATAAACTATCGGACCATGATAACTCTTCACATCGATCAAGGCCTTCACATCCATGCTTTCCACAGATAGAAAAAGACAACATGGATGTTACCTCTAATCCTCCTTGAAATTGCACCGTTATAGCAATGTATTTTGTGGGCGATATATCTGTTATGTTGTCGAGTTCTATAAGCTAAATAACGATTTGGAAGGTAACCGGTGTGAATTTGATAGACTATTCACAACCATAATATGTACTAAATGCCTTGTACTTTAAACAAAATAAGACAAGATCTCAAGAACTCGAGGCACGAGTACCGAGTAATGTGATGGGCAGTAAGTAAAAATTCCCTTGAATAAGTAAACTAGCTAAACTTTTGGCT

>F01_transcript/35265 full_length_coverage=2;length=2001;num_subreads=45

GAACGACACCAAGAGAGAAAGATGAACTCTAAAGATCTCTGGCTAGATATCCATCAACATGCTGCTCTAATCGAATACCTCGCCCTCCTCGCCTTGTTATTCACTCTCTACAAGCTTTGGCGCCGCAACAAACCATCCAAGAAATCAAACCTACCGCCTTCTCCTCCAAAGCTCCCGGTCATCGGAAACCTCCACCAGCTCGGCTCCATCCCTCACCACACCCTCCGATCCCTCTCCGACAAGTATGGCCGCTGATGCTCTTCCACTTTGGTCGCGTCCCCACCATCATCGTCTCGTCCGCCGCCATGGCCCAAGAGGTGATGAAGACCCACGACCTTCCCTTCTCGAGCCGTCCCTCGTCCAAAATATTTAGCACGTTTTCCTACAACCACAGAGGCGTCGTATTTACGCCCTATGGCGAGTATTGGAGGCAGGTGAGGAGGATATGTGTCCACCACCTCCTGAGCCCCAAGAGGGTCCAATCTTTTCGATCCGTGAGGGAGGAAGAGATCAAGATCATGGTTGATAAAATACGAAGCCGCTCGGATTTGGGGCCGATCGACTTGAGCGAAATAATGATCACGTTTACAAGTGATACAACATGCAGGGTCGCTTTTGGGAGGAAGTACTTGAGCAAGGAGGGAGGAGGCAGCGGAGCCCGAGAGCTTCTGGATGAGTTCGTGGCATTGTTAGGGAAGTTCCCAGTTGGGGACTTCATACCGGGATTCTCGTGGTTTGATCGGTGGAGCGGGTTGGATGCCAGAGTGAGGAAGACCTTCAAAGAATTCGATACTTTCATTAATAAAGTGGTGGAGGATCATATCCATGGTGAGAGGTCTCATGACGGTGAGGACTTTGTTGATGTTCTTCTCTCACGGGATGAGGGATCACATGACACTACTGGTTTCTCTCTCAGTAGAGAGAGCATTAAGGCCCTAATTCTGGACATGTTTGCTGCTGGAACAGAAGACATCTTCACCGTCCTTGAATGGATAATGGCGGAGCTAATCAGACACCCGGAATCAATGAAGAGAGTACAAGAGGAAGCACGCAGTATTGATGTGTTAAAAGAAGAGGACTTAGATAACATGAGATATCTAAAAGCAGTAATTAAGGAAGCTCTGAGGCTGCACCCTCCGATTCCCTTACTTGTTCCTCGAGAATCCACCGAAGATGTCCGATTATGTGGTTATGATATCCCGGCGAAGACTAGGATCATGGTCAATGCTTGGGCGATCGCAAGGGATCCCGGATTATGGGATCGACCGGATGAGTTCTTGCCGGATCGATTCATGGACAGTTCGATTGATTTCAAAGGACATGACTTCAAATATATACCGTTCGGTGCTGGTCGAAGAGGCTGCCCAGGAATTGGGTATTCCATTCCCACTATTGAACTCGCGCTTGCCAATCTTCTGCGGCATTTTGATTGGGCTCTGCCTAATGGGGAGATGATGGATATCAGTGAAACATCGGGCATGGCAACTCACAAGAAATCTAGCCTTGTTCTTGTGGCCAAACCGTTACTTTGATTATAGCACGAGTGCATGCACTAGATTGGATTAAGTGTACTAAAGTAAAAGTACGAGATGCAATAAGACTTTGACCTATCTTAAGGATTTATAGGGTAAATGTTTGGTTGGTGCTTGAAAACTAATGGTGGGATGAACCAGGAGGCTTCCTACTGAAGATATCGATTGGAGCTCTGCCCCAATTTTTCATCGACTGAATATCTGGTGTTCTGAAGATTTTTCACTAAGACCTCCCCTATATCATGATCCTCTCCTAAGGATACAAACACACAACTGAGAATACGGCATTGGGGAGAACAGGATTATTTTATTCCAGATAAATAATGAATAGGGAGAAGACATTAGTTATTGTATCGTGAGAAAAGTATTATTCATTTGGAGATCATCAATTTTGCTTCAAACTATTATTATACGATGTAATCTTCTGAACACAATTTTACTGTAGGATTGCTATCACTATAGGCCAGAG

>F01_transcript/35481 full_length_coverage=2;length=2005;num_subreads=60

GATATTCAATTGAATTTCAGCCGTTGGATTATAGATTTGGCCGGGGATGGCAAGATGCTCTCTCATTGTAAAACCCCCCTTGGCAATTGCAGGTTTTTAAGAAGCAAAAAAAAAAAAAAAAAATCATGGCTTCTCATCTCACTGCCATTGCCTTCCTCGCTCTCTTTCTCCTCTTCGCTGCCATCATCGCCATTTTTCTCTACATTGCCGGCTCCATCGCAGCCTGGGCTTTCTTCTACTTGAAGGGATCCATTCAGAAATCCGGCCGCCCCCCCATGGCTGGCAGCATGTTTGACGTGCTCCTCTATTTCGACACCTTCTTCGACTTCCAAACCACCCTCGCCCGCCGCCACCGTACCTTTCGGCTATCACACAACGAGATCTTCACCACTGACCCCGCCAACATCGAGCACATACTCAAGACTAACTTCTCCAAATATGTCAAGGGCGAGCCCCATAGTAGCGTCCTGATGGATCTCTTAGGTGAAGGGATATTCGCGGTGGATGGGGAGAAGTGGCAACATCAGAGAAAGCTTGCGAGCTACGAGTTCTCCACTAGAATCCTTCGAGGATACAGCAGTGTCGTGTTTCGCTCGAATGCTGCCAAGCTTGCGATGAAGATTGCTGGTTCGGCGAGCACTGGAGCTGTGATAGACATGCAGGAACTTTTTATGAAATCAACATTGGATTCGATATTTAAGGTCGGATTCGGGGTTGAGCTTGACACACTTTCTGGGTCCGAAGATGTCGGAACTCGATTCAGCAAAGCCTTTGACAACGCAAACTGGATCACTTGCCGGAGGTATTTCGATATGTTCTGGAGACTCAAAAGGCGTCTCAACATTGGGTTAGAGGCCAAACTGAAGGAGGACATCAAGCTGATTGATGACTTTGTCTTTCAATTGATCCGCCGCAAGAGGGAGCAGATGAAAAATTGGATTGAAGAAAATGACAAAGAAGATATCTTGTCGAGATTTATAGTCGCGAGCAAGGAGGATCCAGAGCACATAACCGATCGATATCTGCGAGACATAATCCTCAATTTCCTCATCGCCGGGAAGGACACAACAGCAAACACTCTAACATGGTTCTTCTACTTGCTCTGCAAGAACCCCATAGTGCAGGAAAAGATCGTCTGCGAGATCAACGACGCTGTGACATCGGAAGGCCAAAGTGTGGACATAATAGACTTTGCGTCCCAATTAACACAACCAGTGCTTGACAAGATGCACTACTTCCACGCTGCGCTCACAGAGACTCTAAGGATCTATCCCGCTGTTCCATCTGATGGAAAGACTGCGGAGGAGGACGATGTCCTTCCGGATGGATTTAAGGTGAAGAAAGGAGATGGGGTGAGTTATATGCCATACGCCATGGGGAGAATGACATATCTCTGGGGCGAGGATGCCGAGGAATTCAGGCCGGAGCGGTGGCTTCAGAATGGGACCTTCCAGCCGGAGAGCACTTTCAAGTTCATCACGTTTAATGCTGGGCCTCGTATATGCTTAGGGAAGGATTTTGCTTACAAGCAAATGAAGATTTTGGCCGCCGTCCTCCTCCATTTCTTCAAATTCAAACTTGAGGATGAGTCAAAGACAGCTGCATATAGAACCATGTTGACCCTACATATAGATAAAGGCCTTCCTCTTATTGCATCTTGTCGATGATATTAATATCGAAAGTCATAATGTATTTCTGTTTCTCCATGAGGAGATAGCAACTAGCATCTTCAACTTTCTCTTCTCATGTGGTAGCAGACTAGCAGGGGAAGATCAACAACATTGGAAGATCAGTTTTTTGTGTTTCTGATTTTGTATCAAGCTTCTCAATTCCTGTGTATATGAGTAGTAATGAACCTGTGGTTATGTAATATGAAGGGGGATCAACTGATTCTCTAGTATCTATGTAATATCAAACTTTGTTCTTGTGTACACTATGGTGTACACTGTTCTGATTATAATATGGGCATCTTGCTTTGCATAGGTGGAGCAAAGTCATTACTTCG

>F01_transcript/37127 full_length_coverage=5;length=1942;num_subreads=60

GACCCTTTACAGAGAGAGATTAAGAGAGAGAGAGAGAGAGAGAGAGAGAGAGAGAGAGAGAGAGAGAGAGGTAAGTCTCTCAAAGGGGAGTGCTCCTACTCCCTACCATGGCGTTGGAGCTGATCCTGGTTCTGTCTTCCTTGATCGTCATCCTCATCATCTTCTTCAGCTTCAAGAGTAATGGGAAGAGTGAGAACAAATTGGCCAAGCTCCCACCGGGCCAAATGGGCTGGCCCTTCATCGGCCAAACCATCCCCTTCATGCAGCCACACTCCTCCGCCTCCCTCGGCCTCTTCATGGACCAAAACATAGCCAAGTATGGGAGGATCTTCCGGACGAACTTGCTGGCGAAGCCGACCATCGTGTCAGCGGACCCGGACTTCAACCGGTACATCCTGCAGAACGAAGGGCGGCTGTTCGAGAACAGCTGCCCCACGAGCATCAAGGAGATCATGGGGCCGTGGTCGATGCTCGCCCTCGCGGGCGACATCCACCGCGAGATGCGATCCATCGCGGTGAATTTCATGAGCAACGTCAAGCTCCGAACCTACTTCCTCCCTGACATCGAGCAGCAGGCCATAAAGGTCCTCGCCAGTTGGGAGAACACCCCCGAGGCCTTCTCGGCCCAAGAACAAGGAAAGAAGTTTGCGTTCAATCTGATGGTGAAGCATCTGATGAGCATGGATCCCGGGATGCCGGAGACTGAGAAGCTGCGGACGGAGTACCACGCATTCATGAAGGGAATGGCTTCAATTCCTTTGAATTTGCCCGGAACGGCTTATAGAAAGGCCTTGCAGTCGAGGTCAATAATTTTGAAGATCATGGGGCAAAAGCTAGACGAAAGAATCCGACAGGTACGTGACGGGTGCGAGGGATTGGAGCAGGATGATCTTCTGGCGTCCGTCTCCAAGCACCCACACCTCACAAAGGAGCAGATTCTCGATCTCATACTCAGCATGCTCTTCGCCGGTCACGAGACTTCCTCGGCTGCCATCGCCCTAGCTATCTATTTTCTCGATTCGTGCCCTAAAGCCGCCCAACAATTGAGGGAGGAGCACGTGGAGATCGCGCGACAAAAGGCTGAGCGAGGCGAGACGGGGCTCAATTGGGACGACTACAAGCAAATGGAGTTCACCCATTGTGTCATAAATGAAACCCTAAGGCTCGGAAACATTGTCAAGTTCTTGCATAGGAAAACTCTTAAAGATGTCCAATTCAAAGGGTATGACATTCCATGTGGGTGGGAGGTGGTGACGATCATCTCAGCCGCTCATTTGGATCCGTCCGTGTATGATGAACCTCAGCGTTACAATCCATGGAGATGGCAGAATATCTCGGCGACTGCATCAAAGAACAACAGCATAATGTCGTTCAGTGGCGGCCCTCGCTTGTGCCCTGGTGCTGAGCTGGCGAAGCTGGAGATGGCAGTTTTTCTGCACCATCTCGTCCGAAAGTTCCATTGGGAGCTGGCAGAGCATGACTACCCCGTATCTTTCCCCTTCCTCGGATTCCCCAAGGGCCTGCCAATCAAGGTTCGTCCCCTCGAGAAGTCAAAAATCTGAGGTCTAATGATGTTATGAACTCTTGAGAGGCTGGTATGGTATGTGGTCTGTTTCAACTATTCAAGTGTGGTTGTAGTTTGTTTAGTAGTAAAGGACGTAGCATTCTGTGTGGACAAAAAAATTAAGGTTTCTGGAAAATAAAAGAGTGTGATAAGTTAGTATCTGATTGTTAAGAGTATTTTCATTTCAAGATTTATCATATGGCCGGGTATGTTTGGCTAGGAGTGGACAGTTTTTAGGGAATGGTTTTTATATAGAAAGATCTGTTTGTTTATGGATTATTTTTATTATTTCTCTTGTTTTTATATAAGATTTAAAAGAATGTTCCATCGAAGATAATTTGAAAGAAATAAAACTGAAGTAACAATATGTTGGCACCC

>F01_transcript/37505 full_length_coverage=5;length=1954;num_subreads=60

GGGATGATAGTGAAATTCCTAACTCCAAATAGTCCAAATACAAATTATTCTGTCAAGCAGAATATTTGGGCAATGAATATTGCGCTGTCCTCACCTCCTTCCTTATAAAAATGGCGCCGGCGATCTGGGGGCAATGAGCGAGTTGAAGCAAGTTCCGGCGATGGCGACGGGAATCGCAGTCGCCGCCGGCTTCGTGATCTTCTCCTCCCTTGCATGGATCCTGCGGCGATTCAACGACTGGTGGTTCCTCAGAGGGATCGAGGCCGAGGCCGAGGCCGGCGGCCCAAAGATCCCTCCGGGATCGCTCGGATGGCCGGTGATCGGAGAGATCATCGACTTCCTTTGGTGCTTCAAGTTCTTGGGGAGGCCGGATGATTTCATCACCAAGCGGAAATTGAGATATGGAGACACTGGTATTTATAGGACTCATTTATTTGGACTTCCGGCTATAGTCACATGTTCACCAGAGTTCAACAAAAAAATCCTTGCTAGCAGCATTGAAGATGGAGATTTTAAGACAGGTTGGCCTTCCAATCAGCTGTTAGGGAGTTCTTCCGTATCTGTTATTGATGGATTATTGCACAAGAGGATGAGGAGGCATTTGCTGGAAGGTTTTAATAGTCCGAGGGCCCTTAGTTCCCATCTAGCTACTGCACAACCTTCCATCATCTCAGGTCTAGAAGATTGGGTGACTAAGAGGAAGATAGTCGCATATAAGGAAACCAAAGCTATGATGTTCCGCAACATTTGTGATATCCTTGTTAGCTTAAAGTCAGAGGCACTTCTAAATAAAATGGAAGTTCTGTATAGGGGACTCATGGCAGGGTTCAGATCAATGACGATCAACATCCCAGGAACAGCATTCCACCATGCTCTCAAGTGCAGGAAAAAGCTAACCATCATTTTATCTGATGAAGTGCTTGAGAGGAAAAAGCAAGGCATTCAAAAGCATGATTTCATGCAATTGCTGATGGATTCGGTCGACGAAAATGGAAATAAGCTGAATGATATGGAAGTTGTTGAGAATGTAGTTTCATTAATATTAGGTGGTTATGAATCGACATCCGATGTCATGTCATGGGGTCTTTACTATTTGGCTAAATACCCAGAGATACTCGAAAAGCTCAAGGATGAAACTCGTTCCATCAGGATGCAGAAGTCCAAGGATGAGCTTCTTACAGCTGAAGATATCAAAAAATTCAAGTATACCTCCAAGGTGGCAGAAGAACTGATTCGTCTTTCAAATATTTCACCTTTTCTATTCAGAAGAGTTGCGAAGGATAATGTAGTCATTAATGGGTACAAGTTTCCCAAGAATTGGAAGGTTATCGTGTGGATTCGTGCAATTCACCTTGACCCTGAATACTATTACCGTCCATTGGAATTCAAGCCGGACAGATGGGATGATTTCAAGCCAAAACCTGGAACCTACTCTGTTTTCGGAGCAGGTACGAGATATTGTCCAGGCAACAGCTTTGCAAAGCTCCAAGTGATGATGTTTCTACATCATTTGTGCCTCAAGTATAGGTGGGAGCTTCTGAATCCTGACGCTGGCATCACATATCTACCACACCCGAGACCAGCGGATGGAGCTGAAATGATTTTCGGTCGAGCAATTTGACAAATGAAAAGGTATTATAGTGGGGCAATGGCGTATTCCAATTAAGGAGATCAAAGACCTGAGCATTAATCATTAAAGATCTTCAAAAATTAGAAGCGTCGGTGTGCTTGAAGTTTTTTGTGGATCTTTCCATGATTATCTTTCGATTCTAGTCCTTCTTTTGTTGTTTTTTGATTATATGAATTTTATGTATGGTTGTAAGACTTTTATTCTTGGCTCATCTTTATGAGCTGTTTATTAAATATTAATAATATTGGAACATTCCATCTCTCTTCCATAATGGAAGAGAGATTATTTCTATCAACAACAAAAAATTACACTGGAAACGTGTGC

>F01_transcript/38919 full_length_coverage=3;length=1900;num_subreads=60

GACTGACTCAATGAGTCAACAAAACGGGCGACGTAATCGCTTAGCTCATCTCTGCCATCTTTCTGCTCTTATGTGATGGCTTCAGATCACCCGTCTCTCTAACATTTCGATCCCAATGGATTCACGGGTCTTAGGGGCGCTGGCGGCGCTGCTGGCGGCGGCGGCCGCGTGGGTTATGAGAGCGGCGGCCGAGTGGTTGTGGTGGCGGCCGCGTCGGCTGGAGCGTGCCCTCCGGTCGCAAGGAGTTGAGGGAAATCCCTATCGTTTCCTCAATGGAGACTTGAAGGAGACAGTACGGCTCACCAAGGAAGCCCGGGCTCAACCCATCTCCCCACCATCTCACCGCTTCCTCCATCGTATCAACCCCCTCCTCTTACGCGCCATCTCCCATCACGGTAAATTTGCATTGACTTGGATAGGGCCCACTCCCCGAGTGAGCATTATGGACCCAGAGTTGGTACGAGAAGTTCTATCAAACAAGTTTGGTCATTTTGCGAAACCTAAGGCTGCCCCGATTGTCAAGTTGTTAGCTACAGGACTCGCAAACTATGAAGGTGAAAAATGGGTCAGGCACAGGAGAATTATTAACCCTGCTTTCCATCTCGAGAAGTTGAAGCGCATGTTGCCAGCATTTTTTTCTTGTTGCAGTGAATTGATAGGCAGGTGGGAGAGTTTGGTTGGCTGTGATGAATCCCGTGAAGTAGATGTTTGGCCTGAATTACAAAACCTCACAGGAGATGTCATATCTAGGACTGCATTCGGCAGCAGCTATGCCGAAGGGCGACGGATATTTCAACTCCAATCGGAGCAAGCTGAACTTCTTATTCAAGCCGTTCAAACTGTATACATACCGGGTTACTGGTTTTTGCCGACTCCGAAGAATATTAGAAGAACAAAAATCGACAAAGAGGTTCGAGCACTCTTGCGAAGCATAATTGAGAAGAGAGAAAATGCCATGAAAATGGGGGATGTACATGATGACTTGTTGGGTTTGTTAATGGAATACAATTTGAAAGAATCAGAACATGGCAACTCTAAGAACATTGGGATGACTACTGAAGACGTGATCGAAGAATGCAAGTTGTTCTACTTTGCAGGGCAAGAGACGACATCAGTTTTGCTCACATGGACGATGATTTTGTTGGGTATGCATCCAAGCTGGCAGGATCGTGCAAGAGAAGAGGTTTCACAAGTATTTGGAAAGAGCAAACCAGACTTCGATGGCTTAAGCCGCTTGAAGACTGTGACCATGATCCTATACGAGGTTCTCAGGCTGTACTCCCCTATTATTTTCCTCACCCGGAGAACATACAAGCCGATGAAGCTCGGAGGCATCACTTTTCCACCTGAAGTACAGCTCGCCCTGCCAATAATTTTCATTCATCATGATCCAGAATTCTGGGGTGAAGACGCCGAAGAATTCAATCCCGACAGATTTGCTGATGGGGTCTCAAAGGCATCGAAAAATCAGATGGCCTTCTTTCCGTTTGGTTGGGGTCCTCGTATCTGCATCGGCCAGGGCTTTGCAATGCTAGAAGCTAAGATGGGGCTGAGCATGATTCTTCAACGGTTCTCCTTCGAGCTCTCGCCAAACTACAGTCATGCTCCTCACACTAAGATTACTCTCCAGCCCCAGCATGGAGCTCAAATGGTGCTGCATAGGCTCTAAAACCTGCTGTTGATTGTTCCATGACTTTGAATTTGAAGACAGAATTGTGAAATCTGTATAGTTTGTGCAGTGTATTTTACTATGTAGTGATAATTCAAATGAGTATTGGCAGTGTTCGCATTTCTGGACCTCGAACCCTCCCAGTCCGACGGTCCAGAGGCCTTAGTATTTTCTGACAATAAATAAAAAATACCTGAGACTTGGGAGGCGGATGCTCTCCATAGTTGGATT

>F01_transcript/38937 full_length_coverage=5;length=1908;num_subreads=60

GCCCTTCATCAGTAATGGCTTCTTCTCTCTCCTCAACCTCCATGACAACCATCTTCACCTCTCAGCTCTTCTTCACATCGCTCGCCTTGGCGCTCTTGCTCCTCATCCTCCTACTCTTCAACAGCAACAAGTTGAAGGGGAACGGGACGTCGAAGCGGCTCAAACTCCCCCCCGGTCCACCGGGATGGCCGGTGGTTGGAAACCTCTTCCAGTTCTTTCGCTCGGGCAAACAATTCATCCACTACACCCGCGACCTCCGCGCTATCTACGGTCCCATCTTCACCCTCCGGATGGGGTCCCGCACCCTCATCGTCGTCAGCAGCGCCGAGCTTCGCCCATGAGGCCCTCATCACCAAGGGCCAGCTCTTCGCCAGCCGGCCCGCCGAGAACCCGACACGGACCATCTTCAGCTGCGACAAGTTCACGGTCAACTCTGCGGTCTACGGCCCCGAGTGGCGCTCCCTCCGACGCAACATGGTCTCCGGCATGCTCTCCGCCTCCCGCATCCGCGAGTACCGCGGCCACCGGCAGTCGGCCATGGACCTCTTCATCTCCCGCATCCGTGCCGAGGCTGATGCTGCCGAGGGCGCCGTCTACGTTCTCAAGAACGCGCGGTTCGCTGTTTTCTGCATCCTCCTCTCGATGACCTTCGGGCTGGAGCTCGACGAGGAGGTCATCGAGCAGATCGACCAGCTCATGAAGAAGGTGCTCATAAACCTCATTCCGAGGATCGACGACTACCTCCCCCTCCTCGCCCCCTTCTTCTCGAAACAGAGGAAGGAGACGCTCGAGATCCGAAAAGAGCAGATGGAAACCCTTGTTCCGCTCATCGAGCAGCGCCAGTCCTTGCTCCAAAACCCTAAACCAAATACCGCGCCCTTCTCGTACCTCGACTCCCTCTTCGACCTAAAGGTCGAAGGTCGGAAATCGGCGCCGACGCCCGCCGAGCTGGTCACCCTGTGCTCCGAGTTCATCAATGGCGGCACCGACACCACCGCCACCGCCATCGAGTGGGGCATGGCCCGCCTCATCGAGAACCTCGACATGCAGTCGAGGATTTACGAGGAGATTGTAGGAGTGGTCGGGGATGAAAGAGCGGTGGACGAGGCCGATACCGAGAAGATGGTGTACTTGCAGGTGTTCGTGAAGGAGCTGCTGCGGAAGCACCCGCCCACATACTTCTCGCTGACCCACGCGGCGATCCAGCCGGCGAAACTCGGGGGGTACGACATCCCGACCGACTGCCAACTTAGACATTTTCCTCCCGACCATCTCGGAGGACCCGAAGCTGTGGATCGACCCGGCCCGGTTCAACCCGGATCGGTTCCTCGGGGGCGGGGAGACGGCGGACATGACGGGGGTGACGGGGATCAAGATGATCCCGTTCGGGGCGGGGCGGAGGATTTGCCCGGGGCTCGGGATGGCGATGACGCACATCACGCTGATGATGGCGAGGATGGTGCAGGCGTTCGAGTGGCGGGCCCACCCGTCGCAGACAGTGCTTGGACTTCGACGACAAGCTTGAGTTCACCGTCGTCATGCGCCGGCCACTGCTGGCCGTGGTGCGGCCGAGGAAGTAGATTGGTGGTGGGGGGTTTGCATGGTTGGGCTGGTAGTACGGAACGTTTCTGCTTTTACTTTGTGCTTCCATTTGGTTCTTATTAATAATTGGTATGGTAGTTGAGAGAGGGAAAGTATGGTACGTGGGTGGGTGAATATTTAAGAGGGAAGTTATTAGAGAGTGCTTTTGAACCGAATGTTGAAATAACAGTGTTTCAATGTTGAAGCATTGAGCAGCCCAGATTTAATATGTCCGACAATCCACTTCGAATTTCATTTTTTTTGGCTTGCAATTCTGTACCAGCTTGGATATTTATATTATTATACTAAGGCGGTTAATTTTGTCAG

>F01_transcript/39299 full_length_coverage=2;length=1885;num_subreads=55

CCCTTCATCAGTAATGGCTTCTTCTCTCTCCTCAACCTCCATGACAACCATCTTCACCTCTCAGCTCTTCTTCACATCGCTCGCCTTGGCGCTCTTGCTCCTCATCCTCCTACTCTTCAACAGCAACAAGTTGAAGGGGAACGGGACGTCGAAGCGGCTCAAACTCCCCCCCGGTCCACCGGGATGGCCGGTGATTGGAAACCTCTTCCAGTTCTTTCGCTCGGGCAAACAATTCATCCACTACACCCGCGACCTCCGCGCTATCTACGGTCCCATCTTCACCCTCCGGATGGGGTCCCGCACCCTCATCGTCGTCAGCAGCGCCGAGCTCGCCCATGAGGCCCTCATCACCAAGGGCCAGCTCTTCGCCAGCCGGCCCGCCGAGAACCCGACACGGACCATCTTCAGCTGCGACAAGTTCACGGTAAACTCTGCGGTCTACGGCCCCGAGTGGCGCTCCCTCCGACGCAACATGGTCTCCGGCATGCTCTCCGCCTCCCGCATCCGCGAGTACCGCTGCCACCGGCAGTCGGCCATGGACCTCTTCATCTCCCGCATCCGTGCCGAGGCTGATGCTGCTGAGGGCGCCGTCTACGTTCTCAAGAACGCGCGGTTCGCTGTTTTCTGCATCCTCCTCTCGATGACCTTCGGGCTGGAGCTCGACGAGGAGGTCATCGAGCAGATCGACCAGCTCATGAAGAAGGTGCTCATAAACCTCATTCCGAGGATCGACGACTACCTCCCCCTCCTCGCCCCCTTCTTCTCGAAACAGAGGAAGGAGACGCTCGAGATCCGAAAAGAGCAGATGGAAACCCTTGTTCCGCTCATCGAGCAGCGCCAGTCCTTGCTCCAAAACCCTAAACCAAATACCGCGCCCTTCTCGTACCTCGACTCGCTCTTCGACCTAAAGGTCGAAGGGCGGAAATCGGCGCCGACGCCTGCCGAGCTGGTCACCCTGTGCTCCGAGTTCATCAATGGCGGCACCGACACCACCGCCACCGCCATCGAGTGGGGCATGGCCCGCCTCATCGAGAACCTCGACATGCAGTCGAGGATTTACGAGGAGATTGTAGGAGTGGTCGGGGATGAAAGAGCGGTGGACGAGGCCGATACCGAGAAGATGGTGTACTTGCAGGCGTTCGTGAAGGAGCTGCTGCGGAAGCACCCGCCCACATACTTCTCGCTGACCCACGCGGCGATCCAGCCGGCGAAACTCGGGGGGTACGACATCCCGACCGACGCCAACTTAGACATTTTCCTCCCGACCATCTCGGAGGACCCGAAGCTGTGGATCGACCCGGCCCGGTTCAACCCGGATCGGTTCCTCGGGGGCGGGGAGACGGCGGACATGACGGGGGTGACGGGGATCAAGATGATCCCGTTCGGGGCGGGGCGGAGGATTTGCCCGGGGCTCGGGATGGCGATGACGCACATCACGCTGATGATGGCGAGGATGGTGCAGGCGTTCGAGTGGCGTGCCCACCCGTCGCAGACAGTGCTGGACTTCGACGACAAGCTTGAGTTCACCGTCGTCATGCGCCGGCCACTGCTGGCCGTGGTGCGGCCGAGGAAGTAGGTTGGTGGGGGGGGTTTGCATGGTTGGGCTGCTAGTACGGAACGTTTCTGCTTTTACTTTGTGCTTCCATATGGTTCTTATTAATAATTAGTATGGTAGTTGAGAGAGGGAAAGTATGGTACGTGGGTGGGTGAATATTTAAGAGGGAAGTTATTAGAGAGTGCTTTTGAACCGAATGTTGAAATAACAGTGTTTCAATGTTCAAGCATTGAGCAGCCCAGATTTAAGATGTCGACAATGCACTTCGGCTTGCAATTCTGTACAAGCTTGGATTTTTATATTATTATACTAAGGCGGTTAATTTTGCCT

>F01_transcript/40246 full_length_coverage=4;length=1853;num_subreads=51

GACTCTCACCACCACCGCTCCTGATTAGTGTGTGCGCTGGTTATAAAGAGAGAAAGAAGATAGAGGGAGAGAGAGAAGAGAGAGAATGGCACCAGTGGTGATTCTCTTCTTCCTCTTCCCCACACTACTGGTCCTAGTGGTTGCAGTCTTGGGTTTGAGAGGAAGCGATGACAGCTGGAAGAAGAGAGGGCTCAAGGTTCCCCCGGGGAGCATGGGCTGGCCTCTCCTCGGCGAGACCATCGCCTTCCGACGCCTCCACCCTTGCACTTCCCTCGGCGAATACATGGAAGAACATGTCAACAAGTACGGGAAGATCTACCGGTCGAACCTCTTCGGGGCACCGACGATCGTGTCGGCGGATGCTGAGCTGAACCGATTCGTGCTGATGAACGACGAGCGGCTTTTCGAGCCGTGCTTCCCGAAGAGCGTGGCCGACATCCTCGGCCACACATCGATGCTGGTGCTCACCGGCGAGATGCACCGCTACATGAAGTCGCTCTCCGTCAACTTCATGGGCATCGCCCGCCTTCGAAACAACTTCCTCGGGGACTCCGAGCTTTACATCACCCAGAACTTCAACAGATGGAAGGAGAACATCCCATTCCCTGCCAAAGAGGAAGCTTGCAAGGTGACCTTCAATCTAATGGTGAAGAATATACTGAGCTTAAATCCCGGGGAGCCGGAGAGCGAACACCTCCGAAAGCTATACATGTCCTTCATGAAGGGTGTCGTCGCAATTCCTCTCAATCTCCCGGGCACTGCGTACAAAAAGGCCATACAATCGAGAGCGACGATACTTAAGATGATCGAGAAGTTGATGGAGGAAAGGATCCGAAATAAAAAGGCGGGGACCGACGAGATCGGAGAAGCCGATCTCCTCGGATTCATCCTCGAGCAGTCGAACCTCGATGCCGAGCAGTTTGGTGACCTTCTGTTAGGGTTGTTGTTCGGTGGCCACGAGACCTCCGCTACGGCCATTACTCTTGTCATCTACTTCCTTTACGACTGCCCCAAAGCCGTCGACCATCTAAGGGAGGAGCATCTCGGAATAGTAAGGGCTAAGAAAGCGAGGGGGGAGCCTCCAGCACTGACTTGGGATGACTACAAACAGATGGAGTTCAGCCAGTGTGTGAGTTTCTCTCTCTTCCCCTTCTTTTTTTATTATTCTCTCTCTATTATTATTCTCTCTCTCTGAAAGAGAAATATGAGACGTCCCATGTGAAGGTGGTAAGCGAGACTCTTCGTCTAGGGAACATCATCAAATTTGTCCACAGGAAAGCCAAAACAGACGTTCAGTTCAAAGGGTACGATATTCCCAAGGGCTGGAGTGTTATTCCAGTGTTTGCCGCTGCCCATCTCGATCCGTCGGTGTATGAAAACCCTCAGAAGTTCGATCCATGGAGATGGCAGACCATCTCGACCGGCACAGCACGAATAGACAACTACATGCCTTTTGGACAGGGCCTCCGCAACTGTGCAGGGTTGGAGCTTGCCAAGATGGAGATCGTCGTCTTCCTCCATCACCTCACCCTCAACTTCGACTGGGAGATGGCAGAGCCCGACCACCCGCTTGCCTACGCCTTCCCCGATTTTCCCAAGGGACTCCCAATCAAGGTCCGCCGTCTCGCGCTCAAATGAAGCTAAGGCTCCAACCATATTCGATGCTCCGTGGCCGTTGGATTGATGCTTGAGGGTTTAGAGATTTGGGTTTTAATGGTAAACTACATTTGAATTGTCGTGTTGCGAGTTATGTTAGCTTCTTCCACATATCTATGTGACCGTTGTTTTTTTTTGTATCAAAACTGCGTTACAAATAAATTGGTAAGGATGCTTGTGTTGCATCTTTTATTAGG

>F01_transcript/40556 full_length_coverage=5;length=1841;num_subreads=60

ACTCTCACCACCACCGCTCCTGATTAGTGCGTGCGCTGGTTATAAAGAGAGAAAGAAGAAAGAGGGAGAGAGAGAAGAGAGAGAATGGCACCAGTGGTGATTCTCTTCTTCCTCTTCCCCACACTACTGGTCCTAGTGGTTGCAGTCTTGGGCTTGAGAGGAGGCGATGACAGCTGGAAGAAGAGAGGGCTCAAGGTTCCCCCGGGGAGCATGGGCTGGCCTCTCCTCGGCGAGACCATCGCCTTCCGACGCCTCCACCCTTGCACTTCCCTCGGCGAATACATGGAAGAACATGTCAACAAGTACGGGAAGATCTACCGGTCGAACCTCTTCGGGGCACCCACGATCGTGTCGGCGGATGCTGAACTGAACCGGTTCGTGCTGATGAACGACGAGCGGCTTTTCGAGCCGTGCTTCCCGAAGAGCGTGGCCGACATCCTCGGCCACACATCGATGCTGGTGCTCACCGGCGAGATGCACCGCTACATGAAGTCGCTCTCCGTCAACTTCATGGGCATCGCCCGCCTTCGAAACAACTTCCTCGGGGACTCCGAGCTTTACATCACCCAGAACTTCAACAGATGGAAGGAGAACATCCCATTCCCTGCCAAAGAGGAAGCTTGCAAGGTGACCTTCAATCTAATGGTGAAGAATATACTGAGCTTAAATCCCGGGGAGCCGGAGAGCGAACACCTCCGAAAGCTATACATGTCCTTCATGAAGGGTGTCGTCGCAATTCCTCTCAATCTCCCGGGCACTGCGTACAAAAAGGCCATACAATCGAGAGCGACGATACTTAAGATGATCGAGAAGTTGATGGAGGAAAGGATCCGAAATAAAAAGGCGGGGACCGACGAGATCGGAGAAGCCGATCTCCTCGGATTCATCCTCGAGCAGTCGAACCTCGATGCCGAGCAGTTTGGTGACCTTCTGTTAGGGTTGTTGTTCGGTGGCCACGAGACCTCCGCTACGGCCATTACTCTTGTCATCTACTTTCTTTACGACTGCCCCAAAGCCGTCGACCATCTAAGGGAGGAGCATCTTGGAATAGTAAGGGCGAAGAAAGCGAGGGGGGAGCCTCCAGCATTGACTTGGGATGACTACAAACAGATGGAGTTCAGCCAGTGTGTGGTAAGCGAGACTCTTCGTCTAGGGAACATCATCAAATTTGTCCACAGGAAAGCCAAAACAGACGTTCAGTTCAAAGGGTACGATATTCCCAAGGGCTGGAGTGTTATTCCAGTGTTTGCCGCTGCCCATCTCGATCCGTCGGTGTATGAAAACCCTCAGAAGTTCGATCCATGGAGATGGCAGACCATCTCGACCGGCACAGCACGAATAGACAACTACATGCCTTTTGGACAGGGCCTCCGCAACTGTGCAGGGTTGGAGCTTGCCAAGATGGAGATCGTCGTCTTCCTCCATCACCTCACCCTCAACTTCGACTGGGAGATGGCAGAGCCCGACCACCCGCTTGCCTACGCCTTCCCCGATTTTCCCAAGGGACTCCCAATCAAGGTCCGCCGTCTCGCGCTCAAATGAAGCTAAGGCTCGAACCATATTCGATGCTCTGTGGCCGTTGGATTGATGCTTCAGGGTTTAGAGATTTGGGTTTTAATGGTAAACTACATTTGAATTGCTGTGTTGTGAGTTATGTAAGCTTCTTCCACAGATCTATGTGACCGTTGTTTGTTTTTGTATCAAGACTGCGTTATAAATAAATTGGTAGGGAGTACTGGATGCTTGTGTCATGGAGATTCACATGACAAGTCAAATAATGGAAGAAGGGGTTCTAGTGGCCATTATTTCTATTATCTATTACTAACACAGATCTCCAAACG

>F01_transcript/40617 full_length_coverage=5;length=1855;num_subreads=60

GAAAATAAAAAAAACAGTGGAGAATAAGAGAGCCCCCCACTTGAAACAAAAAAAAAACGCCGAAGGCAGTTGTGCCGTCTTTTCGACCCGATGGATCTCTCCCTCTCTTCTCTTATCCTCCCCGCCACCGCCCTCGTCATCGCCGCCCTTCTCTCCTTCGTCTTCTTCTCCTCCAAACAATCCCAACCCCAAAACCCCATCATCATCAAGGGCCACCCCATCATCGGCAACCTCCTCCTCTTCGTCAAGAACGAACACCGACTCATCGATTGGGTCGCCGAGCTCGTCGCCGCCAGCCCCACCTGCACCGCCACCGTCGTCCCCATCGTCTTCACCGCCAACCCTTCCAACGTGGAGCACCTCACCAAGTCCCGCTTCGACAACTACCCCAAGAGCTCCACCGTCACCGCCTGCCTCCACGACTTCCTCGGCCGAGGCATCATCAACGTCGACGGCGACAGCTGGCGATCCCAGCGCAAGTCCGCCAGCTTCGAGTTCAACACCCGATCCATCCGCTCCTTCATCCTCGACACCGTCAGCTTCGAAGTCTCCGCCCGACTTCTCCCCTTTCTGTCCGCCGCTGCCGCTTCAACAGATACCGAACGAACCATCGATCTGCAGGACGTGCTCGAAAGATTCGCTTTCGACAACATCTGCCGCCTCATTTTCGACAAGGACCCCAAATGCCTCGGCGGCGGCAGCGACGATGGCGAGCGTTTCTACCACGCTTTCGACGAGGCCGCAACCCTCAGCATCTTCAGATGGAGATACCCTTTCCACTGTATGTGGAAGCTCCTCAGATTCCTCGACGTCGGATCGGAAAAACGGCTGAGAGCTTCCATGACTGTCGTCCATGAATCCGTTTGTCGCTGGATCCGGAGCCGCCGTGCTAGCGGCGGCAGCAGCAAGAGCGACTTTTTGTCGAGGTTGGCCGACGATGCAGCCCACTCAGATGAGTTCGTAAGGGACATATTGATCAACTTCGTGCTCGCCGGTCGGGACACCACCCCATCGGCGCTCACTTGGTTCTTCTGGATCACATCGTCCAGACCCGATGTCTTGCTGAAGATCATCAACGAGATTAAGTCGATTCGTTCGAATGACAATCACGAAGCCTTCACTCTGGACGAGCTGAGGGAGATGCAGTATCTTCATGCGGCGATATCGGAGTCCATGAGGCTGTATCCGCCGGTGGCATTGCAGGCGAGGGAATCACTAAAGGACGACGTGTTGCCGGATGGGACGGAGGTGAGGAAAGGGTGGACGTTGATGTACAGCGCGTATGCGATGGGGAGGATGGAGGGAATTTGGGGGAAGGATTGCTTGGAGTTCAGGCCGGAGAGGTGGTTAGACGAGGTGGGTGTTTTCAAGCCGACGGATCCGTTTCGGTACCCGGTGTTTCACGCTGGGCCCCGGCTTTGTTTGGGGAAGGATTTCGCGTATATACAGATGAAGGCAGTGGCGGCGAGTATTCTTGAGAGGTTTGAATTAGAGGTGGCCGAGGAGCGAGGGAGGTATCAGCAACTGCTCACTCTGCGGATGGCGGAAGGATTGCCGGTGAGAGTGAAGGAGAGGAAGAGAGATGCTTAGATGTTCAACGTTATGCTTCTTGGTCTCCAGGATATTTGGGTGTTCCACGATATGCCTCAGAGAATGATGGGCTTCACCTTTTTTTTTTTTTTGCAATCACCCTTCATCTCACTCTATCTAATTAATGTAAGATTACTCGCCCCAATTACATTGTCATTTCTGCTCGAGAACCACTATGATGGTCAAGTTGGCCTTCCCAAACATCCTAATGAGTAGCACCATGTCCTCAGAATTTAGTGTTAGTTGATTTGGGAACTCGCACCTT

>F01_transcript/41092 full_length_coverage=3;length=1826;num_subreads=60

GTGATGAGGCCTATAAATACACATAGAATCCATTGTTGAGGCCAGAGAGCAATAGGAGCAAACCCATCTCATTCGTTGGAATCCAAGGGTGAGAAAATGGTGCTCTTTGGAGTAGTGCTTGGGTTGGGCCTTGGATTTCTTGGAATTTGTTGTTGTTTGCTAAGATGGAATGAGGTGAGGTACAAGATGAAGGGCTTACCTCCAGGGACTATGGGGTGGCCGGTCTTTGGAGAAACTACAGAGTTTCTGAAGCAAGGCCCAATCTTTATGAAAACCCAGAGGGCAAGGTATGGGAGTTTGTTCAGGTCTCATATATTGGGATGCCCTACAGTTGTGTGCATGGATCCAGAGCTCAACAGGTTCATCCTCATGAATGAAGGGAAGGGATTTATTCCTGGTTATCCAAAATCCATGTTAGATATATTGGGTGAGTGCAACATTGCAGCTGTGCATGGCCCACTCCACAAGACCATGAGGAGTGCAATGCTTGGTCTCATTAATCCTCCCATGATCAGGGATCAACTCCTCCACAAGATTGATGAGTTCATGAGATCCCACCTTAGCAACTGGGATGGAAAAGTCATAGATATTCAAGAGAAAACAAAGGAGATGGCCTTGCAGTCCTCCCTGAAGCAAATTGCTGGAATCGAAGCCAGTCCGATTTCTGATGCCCTCAAATCGGAGATCTTCAAACTTGTTCTAGGAACTCTCTCACTGCCCATCAATCTTCCAGGGACAAACTATCGTCGTGGATTCCAGGCTAGGCAAAAAGTTTTAGGCATGCTGAGAGAGATCATTGAGGAGAGGAGGAGTTCCCAGTGTTCCCAAAATGACATGCTTGATTCTCTTTTGAAGACTGAAGACAACACAAAAGTAAATCTCACTGATGAACAGATTATTGATCTGATCATCGCTCTCGTGTATTCGGGTTACGAAACCGTTTCCACGACTTCAATGATGGCTGTCAAATATCTTCATGACCACCCAAGAGCTCTTGAAGAACTCAGGAACGAACATCTGACGATCAGGAAAAGAAAGTTGGATTGTGATGGGATTGATTGGAATGACTACAAGACAATGAATTTCACTCGTGCGGTGATATTTGAGACTTTGAGATTGGCCACAATTGTAAATGGAGTGCTGAGGAAAACAACCCAAGATATGGAATTGAAAGGGTTCATAATTCCAAAAGGGTGGAGAATCTATGTTTACACAAGGGAGATCAACTATGACCCTTTTCTGTATCCCGAACCATTTACCTTCAATCCATGGAGATGGCTGGACAAGAACTTGGAGTCACACCAGAATCTCATGATGTTCGGTGGAGGAAGTAGGATGTGCCCGGGAAAAGAATTAGGGACAGCAGAGATCGCGACATTTCTCCACTACTTTGTAACTAGATACAGATGGGAGGAAGTTGGAAGCGATCCAATAGTGAAATTTCCAAGAGTTGAGGCGCCCAACGGGTTACACATCCGGGTTCGAGCTTACTGAGTTCACAAGGACATTAGATAGGCTTACTAACATCAGTTCTTTTCTGAATTTACTTCTAGTCCTTTTTTTTGTAAATGAGATTAGACTAGTATCTGAGAGTATAGATGGAGTCATATTGATAGATTTACTTACAAAGGAGTCCATGCTGTAGTGAGTTGCTGAATCAATAGAGGGATGGCACATTTTGTCATTTCAAAGGTTGCTAATCTAATCCTGAAGTTTTTCTTTTGCCAAAGGATAATTCAGTTTGTGCCTTAACCTGCAACTTGTACAACCGTTTATTGTATTCACGGAGAACTAAATTAATCCAATATGGTTCGTTCTCATAAGTT

>F01_transcript/41380 full_length_coverage=6;length=1818;num_subreads=60

GGCAATGGTTTGGGTTCCATTTGTGATTTTAATAAAACTGAGGAACTTTCAGCAGTCTTACTGATAACCATCCCCTCTAATCTCATCTTAGGGTTTCACCACAGATCGGAGACGATCCTCGCATTCAGGCAAATCGCCATGGATCTGAAGTTGTTGAAGGCAGGATTGTTGTTCGTGGCGACCATAGTGGTGGCGAGTCTCCTCTCGGGGTTTCTGGCGGCGCGTAGCAAATCGAAAAAAACGGCTCCCACCGAGAATAAAGGCGTTTCCGATCATCGGAGGACTTCTGCGGTTCATGAAGGGGCCGATCATAATGATCAGGGAGGAGTATCAGAAACTGGGGAGCGTGTTCACGGTCAACGTGCTCAACTGGCCCATTACCTTCTTCATCGGCCCCGAGGTGTCAGCACACTTCTTCAAGGCCCCCGAGGCCGATCTCAGCCAGCAGGAGGTGTACCAGTTCAACGTCCCCACCTTCGGACCGGGGGTCGTCTTCGACGTCGACTACTCCGTGCGCCAGGAGCAGTTCCGATTCTTCACCGAGGCTTTGAGGGTCACTAAGCTCAGGAGCTATGTTGATCAGATGGTCGTCGAGGCACAGGACTATTTTTCAAAATGGGGAGAGAGCGGGACGGTGGATCTAAAATACGAGCTGGAGCATCTCATCATCTTGACGGCAAGCCGGTGTTTGTTAGGACGGGAGGTGCGTGACAAGCTCTTTGACGATGTCTCGGCTCTCTTCCACGACCTCGACAACGGGATGCGTCCCATCAGCGTAATCTTCCCTTACCTCCCGATCCCCGCGCACCGGAAGCGAGACCGGGCCCGGGCCCGCATCGCCGAGATCTTCACGACCATCATCAATTCACGGAAATCCTCCGGCAAGACCGAGGACGACATGCTCCAATGCTTTATCGAGTCGAAATACAAGAACGGGCGGCCCACCACCGAGGGCGAGATCACCGGCCTCCTCATCGCGGCGCTCTTCGCCGGGCAGCACACCAGCTCCATTACCTCCACCTGGACCGGGGCTTACATCCTCACGTTCCGGAAGTTTCTATCGTTGGCCGTCGACGAGCAGAAGGTTCTGTTGAGGAAGCACGGGGAGAAGGTGGATCACGACGTCCTATCGGAGATGGACGTCCTGTATCGCTGCATCAAGGAGGCGCTGAGGTTGCATCCGCCGTTGATCATGCTGCTGCGACGCTGCCATTCGGAGTTCACCGTGAGGACGCGGGAGGGGGACGAGTACAAGATTCCCAAGGGCCATATTGTCGCGACCTCGCCGGCGTTTGCGAACCGGCTCCCGCATGTTTTCAAGGACCCAGATTCGTACGACCCGGACCGGTTCGCGGTTGGGAGGGAAGAGGATAAGGCGGCGGGGGCGTTCTCGTACATTTCTTTCGGGGGCGGGAGGCACGGATGCCTCGGGGAACCCTTTGCTTATCTCCAGATTAAGGCGATTTGGAGCCATTTGCTGAGGAATTTCGAGTTCGAGCTCATCTCGCCTTTCCCGGAGAACGACTGGAATGCAATGGTGGTCGGGGTGAAGGGAGAGGTGATGGTTCGCTACAAGCGAAGGAAGCTGTCGGCTGAGATATGATCCATTGTTTTATGTTAGTCTTCTGTGAGCTATGTTATCATCAGAAACTGCTTCAACTTTCTTTAGCTGGTTTTTCTAATCGTAAGTGTATGCTTTTGAGTCCAAGTGTGTATTGTTGTTCAATGCAACTGCATGTATGCCCCCTATATACATGCACACGGTATTCTAGTATATAAATGTCATTTAGCACAGCTAGACGAACCCTTAGATTTACC

>F01_transcript/41617 full_length_coverage=8;length=1801;num_subreads=60

GACTCTCACCACCACCGCTCCTGATTAGTGCGTGCGCTGGTTATAAAGAGAGAAAGAAGATAGAGGGAGAGAGAGAAGAGAGAGAATGGCACCAGTGGTGATTCTCTTCTTCCTCTTCCCCACACTACTGGTCCTAGTGGTTGCAGTCTTGGGCTTGAGAGGAGGCGATGACAGCTGGAAGAAGAGAGGGCTCAAGGTTCCCCCGGGGAGCATGGGCTGGCCTCTCCTCGGCGAGACCATCGCCTTCCGACGCCTCCACCCTTGCACTTCCCTCGGCGAATACATGGAAGAACATGTCAACAAGTACGGGAAGATCTACCGGTCGAACCTCTTCGGGGCACCCGACGATCGTGTCGGCGGATGCTGAGCTGAACCGGTTCGTGCTGATGAACGACGAGCGGCTTTTCGAGCCGTGCTTCCCGAAGAGCGTGGCCGACATCCTCGGCCACACATCGATGCTGGTGCTCACCGGCGAGATGCACCGCTACATGAAGTCGCTCTCCGTCAACTTCATGGGCATCGCCCGCCTTCGAAACAACTTCCTCGGGGACTCCGAGCTTTACATCACCCAGAACTTCAACAGATGGAAGGAGAACATCCCATTCCCTGCCAAAGAGGAAGCTTGCAAGGTGACCTTCAATCTAATGGTGAAGAATATACTGAGCTTAAATCCCGGGGAGCCGGAGAGCGAACACCTCCGAAAGCTATACATGTCCTTCATGAAGGGTGTCGTCGCAATTCCTCTCAATCTCCCGGGCACTGCGTACAAAAAGGCCATACAATCGAGAGCGACGATACTTAAGATGATCGAGAAGTTGATGGAGGAAAGGATCCGAAATAAAAAGGCGGGGACCGACGAGATCGGAGAAGCCGATCTCCTCGGATTCATCCTCGAGCAGTCGAACCTCGATGCCGAGCAGTTTGGTGACCTTCTGTTAGGGTTGTTGTTCGGTGGCCACGAGACCTCGGCTACGGCCATTACTCTTGTCATCTACTTTCTTTACGACTGCCCCAAAGCCGTCGACCATCTAAGGGAGGAGCATCTTGGAATAGTAAGGGCGAAGAAAGCGAGGGGGGAGCCTCCAGCATTGACTTGGGATGACTACAAACAGATGGAGTTCAGCCAGTGTGTGGTAAGCGAGACTCTTCGTCTAGGGAACATCATCAAATTTGTCCACAGGAAAGCCAAAACAGACGTTCAGTTCAAAGGGTACGATATTCCCAAGGGCTGGAGTGTTATTCCAGTGTTTGCCGCTGCCCATCTCGATCCGTCGGTGTATGAAAACCCTCAGAAGTTCGATCCATGGAGATGGCAGACCATCTCGACCGGCACAGCACGAATAGACAACTACATGCCTTTTGGACAGGGCCTCCGCAACTGTGCAGGGTTGGAGCTTGCCAAGATGGAGATCGTCGTCTTCCTCCATCACCTCACCCTCAACTTCGACTGGGAGATGGCAGAGCCCGACCACCCGCTTGCCTACGCCTTCCCCGATTTTCCCAAGGGACTCCCAATCAAGGTCCGCCGTCTCGCGCTCAAATGAAGCTAAGGCTCGTACCATATTCGATGCTCCGTGGCCGTTGGATTGATGCTTCAGGGTTTAGAGATTTGGATTTTAATGGTAAACTACATTTGAATTGTCGTGTTGCGAGTTATGTAAGCTTCTTCCACATATCTATGTGACCGTTGTTTTTTTTTGTATCAAAACTGCGTTACAAATAAATTGGTAAGGATGCTTGTGTTGCATCTTTTATTAGGAAATCTTTGATTATAGCGATCCTAATTATTTTCTGGGTTTTT

>F01_transcript/41732 full_length_coverage=6;length=1814;num_subreads=60

GACCCTTTACAGAGAGAGATTGAGAGAGAGAGAGAGAGAGAGAGAGAGAGAGAGAGAGAGAGAGAGAGGTAAGTGTGTCAAAGGGGAGTGCTCCTACTCCCACCATGGCGTTGGAGCTGATCCTGGTTCTGTCTTCCTTGATCGTCATCCTCATCATCTTCTTCAGCTTCAAGAGTAATGGGAAGAGTGAGAGCAAATTGGCCAAGCTGCCACCGGGCCAAATGGGCTGGCCCTTCATCGGCCAAACCATCCCCTTCATGCAGCCACACTCCTCCGCCTCCCTCGGCCTCTTCATGGACCAAAACATAGCCAAGTATGGGAGGATCTTCCGGACGAACTTGCTGGCTAAGCCGACCATCGTGTCAGCGGACCCGGACTTCAACCGGTACATCCTGCAGAACGAAGGGCGGCTGTTCGAGAACAGCTGCCCCACGAGCATCAAGGAGATCATGGGGCCGTGGTCGATGCTCGCCCTCGCGGGCGACATCCACCGCGAGATGCGATCCATCGCGGTGAATTTCATGAGCAACGTCAAGCTCCGAACCTACTTCCTCCCTGACATCGAGCAGCAGGCCATAAAGGTCCTCGCCAGTTGGGAGAACACCCCCGAGGCCTTCTCGGCCCAAGAACAAGGGAAGAAGTTTGCGTTCAATCTGATGGTGAAGCATCTGATGAGCATGGATCCCGGGATGCCGGAGACTGAGAAGCTGCGGACGGAGTACCACGCATTCATGAAGGGAATGGCTTCAATTCCTTTGAATTTGCCCGGAACGGCTTATAGAAAGGCCTTGCAGTCGAGGTCGATAATCTTAAAGATCATGGGGCAAAAGTTAGACGAGAGAATCCGACAGGTACGCGACGGGTGCGAGGGATTAGAGCAGGATGATCTTCCTGGCGTCCGTCTCCAAACACCCACACCTCACAAAGGAGCAGATTCTCGACCTCATACTCAGCATGCTCTTCGCCGGTCACGAGACTTCCTCGGCTGCCATCGCCCTAGCTATCTATTTTCTCGATTCGTGCCCTAAAGCCGCCCAACAATTGAGGGAGGAGCACGTGGAGATCGCGCGACAAAAGGCCGAGCGAGGCGAGACGGGGCTCAACTGGGACGACTACAAGCAAATGGAGTTCACCCATTGTGTCATAAATGAAACCCTAAGGCTCGGAAACATTGTCAAGTTCTTGCATAGGAAAACTCTTAAAGATGTCCAATTTAAAGGGTATGACATTCCATGTGGGTGGGAGGTGGTGACGATCATCTCAGCCGCTCATTTGGATCCGTCCGTGTATGATGAACCTCAGCGTTACAATCCATGGAGATGGCAGAATATCTCGGCGACTGCATCAAAGAACAACAGCATAATGTCGTTCAGTGGCGGCCCTCGCTTGTGCCCTGGTGCTGAGCTGGCCGAAGCTGGAGATGGCAGTTTTTCTGCACCATCTCGTCCGAAAGTTCCACTGGGAGCTGGCAGAGCATGACTACCCCGTATCTTTCCCCTTCCTCGGATTCCCCAAGGGCCTGCCAATCAAGGTTCGTCCCCTCGAGAAGTCAGAAGCCTGAGGTTTAATGATGTTATATATGAACTCTTGAGAGGCTGGTATGGTATGTGGTATGTTTCAACTATTCAAGTGTGGTTGTAGTTTGTTTAGTAGTAAAGGACGTAGCATTCTGTGTGGACAAAAAATTAAGGTTTCTGAAAAATAAAAGAGGGAGAAGGTGGTATCTGATTGTTAAGAGTATTCTCATTTCAAGATTTATCATATGACAGTTTTTAGGGAATGGTTTTTATATAGAAAGATCTATTTGTTTATGG

>F01_transcript/42374 full_length_coverage=6;length=1794;num_subreads=60

GATATTCTCACACTCTCTCTCTCTCTAATCCCATGTGCACCCATATCTGATATCGCATCCCACCTCTATGCCAGTTCCCACTTCTATTGAACTATATCACCAGAGGCTAGCACTTAGTAATCACCTATAAATATGGTGCCCTATGTTGCACTCTTTGGTGGGAGTTTTCTCTTCTTATGGGTGATCCATTGGGTGTACAGATGGATGCACCCAAGATGCAGTGGGCGGCTCCCGCCTGGTTCCATGGGGCTCCCCTTGATTGGTGAGAGTCTCCAATTCTTCGCCCCGAACAACTCCTTCGATATCCCGCCTTTTGTCAGAGAGAGAATCGAGAGATATGGACCGGTATTCAAGACAAGTTTAGTTGGAAAGCTTATTGTGGTATCCACCGACCCGGAGTTTAACCACTTTGTGTTCCAGCAAGAAGGGACGTTGTTTCAGAGTTGGTATCCTGAAACATTCACGGAGATCTTCGGACGTGATAATGTGGGCACGTTACACGGGTTCATGTACAAGTATCTCAAGAGCATGATTCTCAAGCTCTTTGGCACCGAGAGCCTTAAACAGATGATGCTTGATGTGGAGAAGTCGGCTTGTAGCAACCTTAGAATGTGGTCAAACTCGTCCACCGTCGAGCTGAAGGAAGGCATAGCAACTATGATATTTGGTCTCACCGCAAAGAAGTTGATAAGTTATGAGCCCTCGAGTTCTTCGGACAATTTGAGGAACAACTTCGTTGCTTTCATACAAGGATTGATTTCCTTCCCGCTGAACATACCCGGCACTGCTTACCACCGATGCTTACAGGGGCGAAAGAAGGCGATGAAGGTGCTCAAGGACATGCTAACAGAGAGACGGAATTCACCGAAAAGGCAATGCATTGACTTCTTTGACTACGTCGTCGAAGAGCTGAAGAAAGAGCGGACATTTCTAACAGAAAAGATTGCGCTGGATTTGATGTTTGCGCTTCTATTTGCAAGCTTTGAGACTATATCATTGGCTTTGACAGCAGCTGTTAAATTTCTCACCGAGAACCCTAGAGTGCTGGATGAACTAACGGAGGAGCATGAAGCGATAGTAACGAACCGGGAGGACCCGGATTCAGGAATCACATGGGATGAATATAAATCGATGGCCTTCACATTTCAGGTGATCAATGAAACAGTAAGGTTAGCAAACATTGCGCCAGGAATCTTCAGAAAAACTTTGAAAGATATAAAGATAAATGGATATACAATCCCAGCAGGTTGGGGTGTCATGGTCTCTCCTCCAGCTGTTCACTTGAATACTGAGCTATACAAGGATCCACTCACCTTCAACCCAGGGAGATGGAAGGGTAGGCCGGAAATGAGCGGTGGATCAAAGCATTTCATGGCATTCGGAGGGGGGATTAGATTTTGTGTGGGAACCGAGTTCGCCAAGATGCAGATGGCTATCTTCCTCCACTGCTTAGTTGCAAAATACAGATGGAGGGCAATCAAAGGAGGGAACATAATTCGAACTCCCGGGTTAAGCTTTCCAGATGGATACCACATCCAACTCCTCTCCAAAGATTAAAGCTGAGCTGGCCATCTTTCTTCGTGCACCACAAGAGTTGTAGCTGTTAGAATATCTATCCCAATGTATTAGAATATTTACTATAATGTCATTAGGGTATATATTTTCTTTACTTGTTCTTTCCTTTAAGTGTTATTTTCTTTACTCGCACTCTCATGTAGTCCTATATAATGTTGCCCATTGTGCCTAATAATATTGAGTAAACCCTTAGGAAATTTTCCCCACGAGTTGATCTCT

>F01_transcript/42557 full_length_coverage=2;length=1792;num_subreads=60

GAAACAGACAGACAGACAGACTCCATCTAGATTCCAAGCCCCCCCGTGCGGTGGTGACCCCCACAGTAGCGAACTGAACCCACATCATTAATGTTGTAAAGATCAATCCAACTCATCCCTTTATTACTCCCTCGCACCGCCATCGCTCTTTTCCATCCCCTTTCCTCTTTCCTCTCTGTCTGGATTGAGAAAGACTCAATGGCATTTTTACTCCTCGTGATCACCATCATTGTTGGTGTGATGATTGTATCCCTGCTGATCCGCCGTTCGTCGGCGGTTTATCCGCCGGGGAGCACGGGGATGGTGCCGGTGGTGGGGGAGACGCTGCGTCTGGTGGCGGCGTACAAGACGGAGGACCCTGAGACCTTCATCGACGATCGCGTCCGCCGCCACGGCCCAATCTTCACCACCCACGTCTTCGGGGAGCGCACCGTCTTCTCCACCGACCCGGAATTCAATCGGATGGCGCTCGCGAACGAGGGCCGCTCCTTCCGGGGAAGCTACCCCACCTCCATCGCCACTCTGCTGGGGACCCACTCCGTCCTCCTCACCCCCGTGGGCCCCCGACACAAGCGCATGCACGCCCTCATCCTCTCCACCTTCGCCTCCACCGCCACCCTTCGAGGATCGCCATACCTCCTCCCCGACATCGACCGCCTCGTCCGATTCAACCTCGACGGGTGGAGCACCCACAATGCCGTGCGGCTGCTGGACCAGACCAAGAAGATCACCTTCGAGCTCACGGTGAAGCAGCTGATGAGCCAGGACCCCGGCGAGTGGACCCAGGGTTTGCGACACCATTACCTCCGCCTCATCGACGGCTTCTTCTCCATCCCCCTCCCCTCCTTCATCCCGTTCACCACCTACTCCCGTGCGCTCAAGGCCCGGAAAGAAGTGGCGGAGGCCATACGAGGCCTGGTGAGGAGGAGGAAGGGGATGATGGGAACGGCGGCAGCAGGTGGGAGAAAGGATCTGTTGGAGGAGCTCTTGGAAGCAGAGGCGACGGAGGACGAGATGGTGGATTTCATGGTGTCGTTGCTGGTGGCCGGGTACGAGACCACCTCCACCATCATGACCCTCGCCGTCAAGTTCCTCACCGACACCCCACTCGCCCTCTTCACCCTTCGGCGAGAACATGAGGAAATAGTGAGGGCGAGGAAGAAGCAGCCATCGGAGCCTTTGGATTGGAGCGATTACAAGTCCATGCCTTTCACCCAATGTGTAATCAACGAGACGCTCCGAGTGGCTAATGTGATTAGTGGAGTGTTCAGGCGAGCGGTCACTGACATTCACTACAAAGGGTACACGATCCCAAAAGGGTGCAAGGTGTTCACATCCTTTAGAGCTGTCCACCTGGATCACCAATACTACAAGGATGCTCGGACATTCAACCCATGGAGATGGCAGGAAAAGGATGCACCCCAACAAACAGGAGGAGCTGGAGGCATATTCACTCCCTTTGGTGGGGGACCACGCATGTGCCCCGGCTACGAGCTTGCCAGGGTCGAGGTCTCCGTTTTCCTCCACTACCTTCTTACCCGTTTTAGTTGGGATGCTGCTGAGGAAGACAAGCTAGTTTTCTTCCCCACCACCCGGACCCTGAAAGGCTATCCCATCAATGTCCGCCATCGCCAGTTGAAACAGCGGTCCCTCTGACTGGATATACTTGAGCATAGATGATGATGATTCATCAAGATCTGCTACTATATCTAATCATATAGAAAGTATAATTTTAGTGCACTGTATCCTGGGATAATAACATTACTCATATAGAATAAGCTGGAGTTGTACT

>F01_transcript/42843 full_length_coverage=3;length=1771;num_subreads=60

GCTCTCTGAGCCCCAAGTCAAACAGCTGGAAGGAGGGAAATGGATCCCCCCTCACTGCTCTCTCTCGCCCTCCTCCTCCTCGCCGCCGCCGCCGCCGCCGCCTTTGTCCTCTTCCCCCGCCGGAGATCGCCAGAGAAAAAACCCAGCTCCGGGAACCCCGGCTCGCTGTCGGAGCTGATCAAGAACGGGCACCGGATTCTCGACTGGATGGTCGAGATCCTCGCGGCGAGCCCCACGAACACGGTCGCGACGTACATGGGAATCGTGACCGCGAACCCGGCCAACGTTGAGCACATGCTGAAGACCAAGTTCGAGAACTACCCGAAGGGCGACCGGTTCGTTACGTTATTGGAGGATTTCCTCGGTCGAGGGATCTTCAACTCCGATGGAGATCACTGGAAGCTGCAGCGGAAGACGGCGAGCTTGGAGTTCAATACCAAGACTATTCGGACTTTCGTCATGGAGAATGTTCGGGTATCCGTTGTGGACCGCCTTATTCCCATCTTCGCTCGTGCTGCGGCTTCCGGGGAGACGATAGATCTTCAGGAAACCCTGGAGCGCTTCGCCTTCGACAACGTCTGCAAGGTGTCCTTCAACGAGGACACCGGCCGCCTCTCGGGCGACGACACTGTTGAGGGCCGCGAGTTCGCCCGCGCATTCGAGCAAGCCAGCGAGCTAATCGTCGGCCGCTACAAGCACCCCTTCCTCCTCTCCTGGAAGCTCATGCGTTTTTTTAACATCGGCGACGAGCGCCGCCTCAAAGAGAAAATCGCGACCGTCCACCGGTTCGCCACCTCGGTCATCCGCCGCCGCAAGTCCGCAGCGTCCCTGGGCGACGACCTCCTATCCCGGTTCATCGCGGAGGCGGACTACCCGGACGAGTTCCTCCGCGACATCATCATCAGCTTCGTGCTGGCCGGGCGGGACACCACGTCGGCCACGCTGACGTGGTTCTTCTGGTTGGCCTCGACTCGGCCGGAGGTTCTTGCCCGGGTCGAGGCCGAGGTGAACGCCGTCCGCCGCAAGAACGGGACCCGTGCGGGGGAGATGTTCACCTTGGAGGAAGTCCGGGAGATGGACTTCCTCCACGCTGCGCTTTCCGAAGCGCTGCGTCTATATCCGCCAGTGCCGCTGCAGACGCGAGCGTGCCACGAGGCGGACGAGTTCCCGGACGGAACCAAAGTCCGTCCGGGAACTACAGTGATGTATAACTCGTACGCGATGGGACGGATGAAGAGCATATGGGGGGAGGACTACACGGAGTTTCGTCCGGAACGGTGGTTGGACAAGGGGGGTGGGTTCCAGCCGAAGAGTCCGTTCCGGTTCCCGGTGTTTCACGCGGGGCCGAGGATGTGTTTGGGGAAGGAGATGGCGTATATTCAGATGAAGGCGGTGGCGGCGAGCGTGGTGGAGAGGTTCGAGGTGGAGGTGGTGGACAAGGAGAGAGTGAGGGAGAAGGACTACACTATGATTCTTAGGGTCAAAGGGGGACTTCCGGTCCGGCTCAAGGAGAAATCGGTTGCCGCAGGTTAATTGGGGTTCGGTTCGGTTTGTTGAACCCGATTTGGCTTTTATTTTGCAGTTGACGTTTTCACGTTTTGTTTGTCTTTCTGATATTTTCAATAAACATCATGAAAGTTGGCTAAGTTCGGTCACTCACATACTAATAAGAGTTGGGGGTTGTTTGGCTGGCTTCTCATTGGGCAGGAATCTCATTTCTGTTTATGTGCGGGAAGTAAAAACCAGAAGCAAAAAATTTGTGAGCCACCAC

>F01_transcript/43034 full_length_coverage=2;length=1789;num_subreads=60

GGGTGACATGGTGATCCGGGCTCGCTGATCGTTGACGAGAAGTAATCTAAAAAAGATGCTTCTCTCTTGAGGAACCTAAAACCCCCTTACAATTGCAGGGTTCGTTTTCTTCTACTTGAAGGAATCCATTCAGAAAACCGGACGTCCCCCAATCGCTGGCACCGTGTTCCACATGCTCATCAATTTCGACACCATCTACGACTTCCAAACAACCCTCGCCCGCCGCCACCGCACCTTCCGGCTCATCAAGCAATCACACAGCGAGATCTTCACCGCCGACCCTGCCAACATCGAGCACATCCTCAGGACCAACTTCTCCAAATATGCCAAGGGCGAGCACCATATTAGCGTCCTGATGGATCTCTTCGGCGAAGGGATATTCGCGGTGGATGGGGAAAAGTGGCAACATCAGAGAAAGCTTGCGAGCTACGAGTTCTCCACTAGAATCCTTCGAGAATTCAGCAGTGTGGTGTTTCGCTCGAATGCTGCTAAGCTTGCAGAGAAGATTGCAGGTCCTGCGAGCACTGGAGCTGTAATAGACATGCAGGAACTTTTTATGAAATCGACATTGGATTCGATATTTAAGGTCGGATTCGGGGTCGAGCTTGACACACTTTCTGGTTCCGAAGATGTCGGAACTCGATTCAGCAAAGCCTTTGACGATGCAAACTATATCACTTGCCGGAGGTATCCCGATATATTCTGGAGACTCAAAAGACGTCTGAACATTGGGTATGAGGCCAAACTGAAGGAGGACATCAAGGTGATTGATGACTTTGTCTTCCAATTGATCCGCCTCAAGAGGGAGCAGATGAAAAATGCGATGGGAGAAAATAACAAAGAAGATATCTTGTCGAGGTTTATAGTCGCTGGTGAGGAGGACCCAGAGCGTATAACCGATCGATATCTACGAGACATAATCCTCAATTTCTTGATCGCCGGAAAGGATACATCAGCAAACACTCTAACTTGGTTCTTCTACTTGCTCTGCAAGAACCCTATAGTGCAGGAAAAGATCGTCTGCGAGATCAGAGATGCTGTGACATTGGAAGACCAAAGTGCTGACATTATTGAATTTGCGTCCGGATTAACACAACAAGTGCTTGACAAGATGCACTACCTCCACGCTGCGCTCACAGAGACTCTAAGGATCTATCCCGCTGTTCCATCGGATGGGAAGACTGCGGAGGAGGACGATGTCCTTCCGGATGGATTTAAGGTCAAGAAAGGAGATGGTGTGGGTTATATGCCATACGCCATGGGGAGGATGACATATCTTTGGGGCGAGGATGCCGAGGAATTCCGGCCGGAGCGGTGGCTTGAGAATGGGACCTTCCGGCCAGAGAGCCCTTTCAAATTCATCGCGTTTAATGCGGGGCCTCGTATATGCTTAGGGAAGGATTTTGCTTATAAGCAAATGAAGATTTTGGCCGCCGTCCTCCTCCATTTTTTCAAATTCAAACTTGAGGATGAGTCCAAGACAGCTACATATAGAACCATGTTGACTCTACATATAGACAAAGGCCTTCCTCTTATTGCATCTTGTCGATGAAATTTATATCTAAAGTCATAATGTATTTATGTTCTTTTTGTTCTTGTTCTTCTTGCTTCGGATTTTAGAGGAGGGCGACTGTCACCTACAATAATTATAGCCGGTCCTCGCACGGCATTTTGATGTGTTTATCTATTCTAAACTCTATATTATTTTGGTATTGTAATATTGGAAAACTGTTGTACAAAAATATCAATGTATTTCTGTTAAGCTATCTTAACTTGATGAGCAAATATT

>F01_transcript/43717 full_length_coverage=6;length=1765;num_subreads=60

GATCTTACCCTTCCCTAAAGTGAGAGAGAGAGAGAGAGAGAGAGAGAGAGAGAGAGAGGGAGAGATGTATATGTTTGTGCTGGTAGCTCCCATTGTGGTGTACTACGCATGGTGGCAGTGGTTGAGGCCGTTGATGCTGGAGAGGAAGCTGAGACGGCAAGGGATCGAGGGGAACACCTACCGCTTCCTCCTATGGGGACATGATGGAAGAGGCTCAGCTGATCCAGAGAGCATGGTCCAGCCCAATGGATCTCAACCATCGGATCGTGCCACGTGTCGCTCCATTCACTCATCAGATGGTGCAGAAATACGGCAAGACATGTTTCTGGTGGTTTGGAGCAACTCCAAGTTTGATCATATGTGATCCAGAGATGGCAAAGGAGGTTTATTTAGACAGATCGGGTCACTTCCAGAAGCCACCACAGAATCCTCACACCAAGCTCCTAACCCAAGGGGTCTCCAGCCTCGAAGGAGAAGAATGGGCCAACCGTCGAAGGATTATCTCTCCGGCTTTCCATGTTGAGAAACTCAAGAGAATGATGCCTTCCTTCACGAATAGCTGCCTTGATCTGATAAAGAGATGGGAGAAGTCAGTCGGTGCGAAAGGATCATGCGAGCTGGATGTATGGATAGAATTCCAGGATCTAACAGGTGATGTCATCTCTCGCGCAGCATTCGGAAGCACCTACGAAGAAGGAAAACAAATTTTCGAACTCCAGAAGGAACATATTGTTCTCGTCATGGAAGCTGCTCAATCTCCATATATTCCAGGTTTCAGATTCCTACCCACCGCAAAGAAGAAGAGAAGAATGCATGTGGACAAGGAGATCAAGACCAAGTTACGGGAAATGATCAACAAGAAGTTGAACTCTATGAACAGAGGAGAATCTGGTAGCGACGACTTACTTGGATTGCTACTACAATCCGTGAACCTCACAGATCAGGATAAGCACGGAATCACGACCGACGAAGTGATCGAAGAATGCAAGCTGTTCTACTTTGCTGGCCAAGAGACAACCTCGTTGTGGCTAACGTGGACATTTGTTCTGTTAGCTATGCACCCGATTTGGCAGCAAAGAGCGAGGGGCGAAATTCCTTCAAAGATGTGGGAAGAATACACCTGATTTCGAATCCATAAGCCAACTTAAGATTGTGACCATGATTCTGTACGAGGTTCTGCGGCTATATCCGCCAGTGCCGTCGCTCTTCCGGCACACACATGAACCAACGAAATTAGGAGACATCTCTCTCCCAGCTGGCATTGATCTCTTGCTACCGACGATCCTCATCCATCATGATCGACAACTTTGGGGCAAAGATGCAGAAGAATTCAATCCGAATAGATTTTCAGAAGGGATATCAATGGCATCATTGGAGGGTCAACATGCATTCTTTTCCCTTCGGATGGGGGCCAAGGATTTGTCTGGGCCAGAGCTTTGCCATGATCGAAGCAAAGATGGCTCTGACTATAATTCTCCAACATTTCTCTTTTGACCTCTCACCCTCCTATGCTCATGCTCCTTACAATGTAGAAACTCTTCAGCCACAACATGGAGCTCATATCATCATACATCAACTTTGAGATTGTAATGAAGCTATTCTTGATTATATCATTGTCTAGGTGATTAAATCTATTGATACAATTCGAAATTTTATTTTTTTGGAGTGGACAATTACTTAATTTGGTGTTGGAATGATAACTCTAGTTTTGTTAAAATAATCAATGAGAGGCTGTCCCTTTTCTGGACTAATTTTGCCTTTAGCC

>F01_transcript/43914 full_length_coverage=11;length=1742;num_subreads=60

GACACTCATGACATCCAAATGGATAGCACAAGGTTTCTCTGTTGATTTTGGGGTATGGAACTACTAGCGCAATGGAGTCTATTATTATATATTTCGCTGGGCGCCATAGCTCTCTATACTCTCCTCATGCAAGCTAGAGTATTAAGATCCACCAACAGACTTCCACCTGGCCCTCGGCCACTCCCCATTGTCGGAAACCTGTTCTTGCTCGGCCAAACCCCCCACCGCTCCCTTGCCCGACTTGCTGACACATTCGGCCCCTTAATGTCCCTCAGGCTCGGCCAAGTCATCACTGTCGTCGTCTCATCACCCGAAATGGCCCGTGAAGTCCTCCACAATAGGGACCAAGCCTTTTCATCCCGCTCGGTCCCCGACGCCGTCCGGATCGACGGCTACCACGAGGTATCCATCGCGTGGCTGCCTGTTTCACCACTATGGCGGAAGCTCCGTAAGATCTACGCTTTCCACCTCTTCTCCCCTCGGAAGATGGAGCTCAACGAAAGCCTCCGGATCGAGAAGTTGAAGGAGCTGGTGAGTTACGTGCGCCAACGCTGCGGTGCTCCAGTGGACATTGGGCAAGCCGCGTTTATGACTTCCCTGAATTTGATATGGAATATGCTGTTCTCCATTGATGTCATCGATTCAGTGGTTGGATATTCAAATTTTAGAGGCGTGGTGTGGGGGGTAGTGAAAGAGGCCGGGAGGCCAAACTTGTCTGATTTCTTCCCGGCGATGGCGGGGCTCGACCTCCAAGGCAGGAGGGGCCGGGTTGGCAAACTCTCTAAAAATATGGATGGGGTGTTTGAGGAATTTATAGCACAGAGATTGAAGAGTTCCACAACATATAATGACTTCTTGGATTGGCTCATCAAGTGTCAACACGAAGAAGATGGATTGGAGCTGCATGGAGGCATTGTCAAATCTTTCCTTAAGGATATTTTCATCGCTGGAAGCGAGTCGACCTCTAGCACAGTGGTATGGGCAATGGCAGAGCTGCTCCGCAACCCGAGATCGATGGCAAAGGCACGCTCCGAGCTTATCGATGCCATCGGGGACGGGAACGAAATAGAGGAGCCCATCATCTTGCATCTTCGGTACCTCAAAGCAGTCGTGAAAGAGACGCTCCGTCTCCACCCCCCGCTCCCCTTCCTTCTACCGCGTAGGGCCGAGACTGACGTCGAGCTAAATGGCTATATCATACCCAAACATTCTCAGGTACTGGTGAATGTTTGGGCTATGGGCCGAGACCATCGAGTTTGGGACGATCCCGACAAGTTCTTGCCTGAAAGGTTCCTCGATAAGGATGTGAACTTCAATGGTGGAAATTTTGAGCTGATACCATTTGGTGCGGGGCGGAGGATTTGTCCTGCGTTAGGAATGGGGTCTCGGATGGTTCATCTTATGTTGGCTTCGCTGCTATATCCATTCGAGTGGAAGGTGCCGGACGGGGTCCATGTTGATATGAAGGAGAAGTTCGGCCTCACATTGGCTAAGGCTGTGCCTCTTCGAGCTGTCCCCATCTAGAGAGCGTCCTCATCATAAAAACGGCCTTTCTGGCACCTTCATTATTACTTTGTATAATAGCCAACTCAGAAAAGGATTGGTATCAATAATTGCTTATTACGAGGGCCCTATACGAGCACTCGGAATGGGAACTCTTGGTAATAAAAACTTAGTCAAATTTAAAATTTTGCTTGTAAAATTTTATTGTCGTGAGCTATCATTTACTTTTATTTTTGTGC

>F01_transcript/43962 full_length_coverage=18;length=1728;num_subreads=60

GATAATGCAATAGATTGGAAACATAGTAGCGGTGTACCACAATGTCGCCAACAATCCTCCAATGCTTCTTCCACCAATATTTTCTCACTCTTCTCTTCTTGTTGTTCCCTCTCTTTTTACTTATTAGTACTATAATAAGGAGCTCCTCCACTAAGAGCAATTTTCCTCCATCCCCACCCAAGCTTCCCATAATCGGGAATCTACACCAGCTCGGCCCACTCCCCCACCGCTCCCTCCGTGACCTCTCGGAGAAGCATGGGAGCCTCATGCTTCTCCACCTTGGTCGTGTACCGATGCTGGTGGCCTCGTCCGCAGACGGGGCCCGTGAGATCATGAAAACTCATGATCTCACCTTCGCCAGCCGGCCTTCCTCCAAAACTGCCAGGATCCTGTTCTTCGGTGCCTCGGATATGGCCTTGGCGCCTTATGGCGAGCACTGGAGGCAAATGAGGAAAGTCACCATCACCAACCTCTTAAGCACCAAGATGGTTCAGTCCTTCCGGCTCGCTAGGGAAGAGGAAGTAGGCTTCATGATTGAAAAGGTCTCATCCCAAGCGCCGTCGGTCGTGGATATGAGTGAAGTCTTGTATTCCTTCATGAACGACCTCATCTGCAGAGTGGTATCGGGGAAGTTCTTTAGGGAAGAGGGAAGGAATGAGACGTTCGAGAGATTAATAAAGGAGAACATTCTTTTGTTCGGTAAGTTCCATGTGGGTGACTACTTCCCATGGTTTTCATGGCTTGATGATCTCTTCTTCGGCTTGGATGCGAGGGCTAAGAGGAACGCCAAGAAATGGGACGATGTTCTTGAAGCCACTCTTAGTGACCATGCCCATCGATTGAGGGGAGAGGAGCACAATGACGACCACTTTGTGGATGTGTTGCTCTCTCTTCAAAAGGACCCCAACATGGAGTATATCCTCACCAGGGAGTACATCAAAGGACTCTTAGTAGATATGTTTGCTGCGGGGACCGATACATCATTTGTAGCCCTCGAATGGGCGGTGGCTGAGCTCCTTCGGAACCCAGAAGCAATGAAGAAACTGCGAGACGAGGTGCAACGCGTATCAGGCGGAAAATCCATAATCAGAGAAGAAGATTTGAGTGAAATGAGCTACCTAAAAGCTGTAATAAAAGAAGTCCTGCGACTGCACCCTCCGGCTGCATTGCTGCTTCCGAGAGAATCGATGGAGGAATGCGAGATACAAGGCTATAAGATCCCCAAGGGAGCAAGAGTCGCCGTCAATGCTTGGGCCATTCGGAAGGGATCCCAAAATTTGGGAATTACCTGAGGAATTTCAGCCTGAAAGGTTCCTCTGTACCGGCTCGCCAGACTTCAAAGGAAATGATTTCGAGTTCATTCCATTTGGGGCAGGCCGACGGATTTGTCCGGCTATTAATTTTGCGGTTCCCACTGGAGAACTTGCGCTGGCGAATCTCGTGCATCGGTTCGACTGGGAGCTGCCGGCGGGCACAACCAGTGAGGGGCTCGACATGGCCGAGGGCGAAGGGATAATAACCAGCAAGAAACAGAGAATTCACCTCATTATGAATCTTCGTTAACTAGCTCATTTCACTCAGATCTTCCTCTCAGATTATGGTTATGGAGGAAACCTAAGGCTAGTTTGATGCCAAGTATCCAATATAAGTGTTGGGATGTATTGTTTACTTAAGGTCGTACAAGTCGCCGAAGATAATAATATGAATAATATGTTGCTTTTTCTGACG

>F01_transcript/44246 full_length_coverage=2;length=1716;num_subreads=14

CGACCCTTTACAGAGAGAGATTAAGAGAGAGAGAGAGAGAGAGAGCGAGAGAGAGAGAGAGAGGTAAGTCTCTCAAAGGGGAGTGCTCCTACTCCTACCATGGCGTTGGAGCTGATCCTGGTTCTGTCTTCCTTGATCGTCATCCTCATCATCTTCTTCAGCTTCAAGAGTAATGGGAAGAGTGAGAACAAATTGGCCAAGCTCCCACCGGGCCAAATGGGCTGGCCCTTCATCGGCCAAACCATCCCCTTCATGCAGCCACACTCCTCCGCCTCCCTCGGCCTCTTCATGGACCAAAACATAGCCAAGTATGGGAGGATCTTCCGGACGAACTTGCTGGCGAAGCCGACCATCGTGTCAGCGGACCCGGACTTCAACCGGTACATCCTGCAGAACGAAGGGCGGCTGTTCGAGAACAGCTGCCCCACGAGCATCAAGGAGATCATGGGGCCGTGGTCGATGCTCGCCCTCGCGGGCGACATCCACCGCGAGATGCGATCCATCGCGGTGAATTTCATGAGCAACGTCAAGCTCCGAACCTACTTCCTCCCTGACATCGAGCAGCAGGCCATAAAGGTCCTCGCCAGTTGGGAGAACACCCCCGAGGCCTTCTCGGCCCAAGAACAAGGGAAGAAGTTTGCGTTCAATCTGATGGTGAAGCATCTGATGAGCATGGATCCCGGGATGCCGGAGACTGAGAAGCTGCGGACGGAGTACCACGCATTCATGAAGGGGATGGCTTCAATTCCTTTGAATTTGCCCGGAACGGCTTATAGAAAGGCCTTGCAGTCGAGGTCGATAATATTAAAGATCATGGGGCAAAAGTTAGACGAGAGAATTCGACAGGTACGCGACGGGTGCGAGGGATTGGAGCAGGATGATCTTCTGGCGTCCGTCTCCAAGCACCCACACCTCACAAAGGAGCAGATTCTCGACCTCATACTCAGCATGCTCTTCGCCGGTCACGAGACTTCCTCGGCTGCCATCGCCCTAGCTATCTATTTCCTCGATTCGTGCCCTAAAGCCGCCCAACAATTGAGGGAGGAGCACGTGGAGATCGCGCGACAGAAGGCTGAGCGAGGCGAGACGGGGCTCAATTGGGACGACTACAAGCAAATGGAGTTCACCCATTGTGTCATAAATGAAACCCTAAGGCTCGGAAACATTGTCAAGTTCTTGCATAGGAAAACTCTTAAAGATGTCCAATTCAAAGGGTATGACATTCCATGTGGGTGGGAGGTGGTGACGATCATCTCAGCCGCTCATTTGGATCCGTCCGTGTATGATGAACCTCAGCGTTACAATCCATGGAGATGGCAGAATATCTCGGCGACTGCATCAAAGAACAACAGCATAATGTCGTTCAGTGGCGGCCCTCGCTTGTGCCCTGGTGCTGAGCTGGCGAAGCTGGAGATGGCAGTTTTTCTGCACCATCTCGTCCGAAAGTTCCACTGGGAGCTGGCAGAGCATGACTACCCCGTATCTTTCCCCTTCCTCGGATTCCCCAAGGGCCTGCCAATCAAGGTTCGTCCCCTCGAGAAGTCAGAAGCCTGAGGTCTAATGATGTTATATATGAACTCTTGAGAGGCTGGTATGGTATGTGGTCTGTTTCAACTATTCAAGTGTGGTTGTAGTTTGTTTAGTAGTAAAGGACGTAGCATTCTGTGTGGACAAAATATTAAGGTTTTCTGAAAAATAAAAGAGGGAGAAGGTGGC

>F01_transcript/44395 full_length_coverage=3;length=1690;num_subreads=17

GTCTGAGCCCCAACTCAAACAGTTGGAAGGAGGGAAATGGATCCCCCTCACTGCTCTCCCTCGCCCTCCTCCTCCTCGCCGCTGCCGCCGCCGCCGCCTTTGTCCTCTTCCCCCGCCGGCGATCGCCGGAGAAAAAACCCAGCTCCGGGAACCCCGGCTCGCTGTCGGAGCTGATCAAGAACGGGCACCGGATTCTCGACTGGATGGTCGAGATCCTCGCGGCGAGCCCCTACGAACACGGTCGCGACGTACATGGGAATCGTGACCGCGAACCCGGCCAACGTTGAGCACATGCTGAAGACCAAGTTCGAGAACTACCCGAAGGGCGACCGGTTCGTTACGTTATTGGAGGATTTCCTCGGTCGAGGGATCTTCAACTCCGATGGGGATCACTGGAAGCTGCAGCGGAAGACGGCGAGCTTGGAGTTCAATACCAAGACTATTCGGACTTTCGTCATGGAGAATGTTCGGGTATCCGTTGTGGACCGCCTTATTCCCATCTTCGCTCGTGCTGCGGCATCCGGAGAGACGATAGATCTTCAGGAAACCTTGGAGCGTTTCGCCTTCGACAACGTCTGCAAGGTGTCCTTCAACGAGGACACCGGCCGCCTCTCGGGCGACGACACTATCGAGGGCCGCGAGTTCGCCCGCGCATTCGAGCAAGCCAGCGAGCTTATCGTCGGCCGCTACAAGCACCCCTTCCTCCTCTCCTGGAAGCTCATGCGTTTTTTCAACATCGGCGACGAGCGCCGCCTCAAAGAGAAAATCGCGACCGTCCACCGGTTCGCCACCTCGGTCATCCGCCGCCGCAAGTCCGCAGCGTCCCTGGGCGACGACCTCCTATCCCGGTTCATCGCGGAGGCGGACTACCCGGACGAGTTCCTCCGCGACATCATCATCAGCTTCGTGCTGGCTGGACGGGACACCACGTCGGCCACGCTGACGTGGTTCTTCTGGTTGGCCTCGACTCGGCCGGAGGTCCTGGCCCGGGTCGAGGCCGAGGTGAACGCCGTCCGCCGCAAGAACGGGACCCGTGCGGGGGAGATGTTCACCTTGGAGGAAGTCCGGGAGATGGACTTCCTCCACGCTGCGCTTTCCGAAGCGCTGCGTCTATATCCGCCAGTGCCGCTGCAGACGCGAGCGTGCCACGAGGCGGACGAGTTCCCGGACGGAACCAAAGTCCGTCCGGGAACTACAGTGATGTATAACTCGTACGCGATGGGACGGATGAAGAGCATATGGGGGGAGGACTACACGGAGTTTCGTCCGGAACGGTGGTTGGACAAGGGGGGTGGGTTCCAGCCGAAGAGTCCGTTCCGGTTCCCGGTGTTTCACGCGGGGCCGAGGATGTGTTTGGGGAAGGAGATGGCGTATATTCAGATGAAGGCGGTGGCGGCGAGCGTGGTGGAGAGGTTCGAGGTGGAGGTGGTGGACAAGGAGAGAGTGAGGGAGAAGGACTACACTATGATTCTTAGGGTCAAAGGGGGACTTCCGGTCCGGCTCAAGGAGAAATCGGTTGCCGCAGGTTAATTGGGGTTCGGTTCGGTTTGTTGAACCCGATTTGGCTTTTATTTTGCAGTTGACGTTTTCACGTTTTGTTTGTCTTTCTGATATTTTCAATAAACATCATGAAAGTTGGCTAAGTTCGGTCACTCACATACTAATAAGAGTTGGGGTTGTTTGGCTGGCC

>F01_transcript/44765 full_length_coverage=13;length=1670;num_subreads=60

GATAATACTATAGTTTGGTAACATATACTACAATGTCTCCCACAGCCATCCAATTCTTCTTTCAGCCATACTTGCTTCCCCTTCTTTGCTTGTTCCCTCTCTTTCTACTGATTACGATAATAAAAAGCTACTCTAATAAGACCAAATTCCCTCCATCCCCGTTGAAGCTTCCCATAATCGGGAATCTCCACCAGCTTGATCGACCCCCCCACCGTTCTCTCCGTGGCCTCTCGGAGAAGCATGGAAATCTCATGCTTCTCTACTTAGGTCGCGTGCCGACTCTGGTGGTCTCGTCTGCAGACATGGCCCGTGAAATCATGAAAACTCATGATCTCACCTTCGCAAGTCGGCCTCCCTCAAAGGCCATCAGGATACTACATTCTGGAGCCATGGGTTTGGGCTTTGTGCCCTACGGCGAGCACTGGAGACAAATGAGGAAGCTCACCATCACCCACCTATTGAGCACCAAGATGGTGCAGTCCTTCGGGCTTGCTCGCGAGGAGGAAGTGGAGTTAATGATCGACAAGATTTCGTCCCGATCGGCATCGGTGGTGGATATGAGCGACGTGCTTTACTCGTTCACGAACGACATCATCTGCAGGGTGGTTTCCGGAAAGTTCTTTAGGAAAGAGGGAAGGAATGAGATGTTCCGAAGATTGACCGGGGAGAACTCTGCTTTGTTAGGGAAGTTTCATGTTGAGGACTTCCTCCCATGGCTTTCATGGCTTGACGTGTTCTTTGGCTTGAATGCGAGAGCAGAGAGGATCTCAAAGGAATGGGATGATGTTTTTGAAGAGACTATTCGTGATCATGCCGAGCGAGCAAAAGGCAATGAGCATGAGGGCGACGACTTCATTGATGTGTTGCTCTCCCTTCAGAAAAATCCCGACATGGACTATATCCTCAACAAGGGCAATATGAAAGCACTCCTAATGGGTATGTTTTCAGCGGGGACCGATACATCATACATACTACTCGAATGGGCCATGGCCGAGCTTCTTCAGAAGCCGGAAGCAATGAAGAAACTGCGAGACGAGGCACACAGACTATCAGGCGGACAATCCATAATCCGAGAGGAGAATTTGAATGAAATGAGCTACTTAAAAGCTGTGGTGAAAGAAGTCCTTCGACTTCACCCTCCAGCTCCATTGCTCCTTCCGCATGAGTCGAGGGAGGAATGCGAGATACAGGGCTATAAAATCCCCAAAGGAGCAAGAGTCATCATCAATGCTTGGGCAATCGGAAGGGACCCCAAAAGTTGGGATTCACCGGAGGAATTCTGGCCTGAGAGGTTCCTTGTTAGTGGCTCACCGGATTTCAAAGGGAATGATTTTCAGTTCATCCCTTTTGGGGCAGGGCGACGGATTTGTCCGGGGCTGAATTTTTCTGTCATCACGGCAGAGCTTGCCCTGGCAAATCTCGTGCATCGGTTCCATTGGGAGCTGCCGGCAGGCATGAGGAGCAAGGGTCTTGACATGACCGAGAACGAAGGATTAACAACCCCCAAGAAACAAAGACTTTGTCTCATTATGAAGCCTTTATTAATTTAGAAGGAAGGAAGGGCGTTGAAATGGATGGTTAATCGAGTTGGCAAGTTAATATTATCTTGAACTGTTAAAATGAATAATATTATGTATGATCACTACAAAAAATAATATTTTTGTCCACGT

>F01_transcript/45025 full_length_coverage=12;length=1635;num_subreads=60

GAAAAAAAAGAAGAGGAGTATGGAGCTGCTAGCACAGTGGAGTCTACTATCATGGATCATCCCATTAGGCATCATAGTTCTCTATATTCTAATCCAGGCAAGAGTAGTGAGATCCACCAAAAGACTTCCACCCGGCCCTTGGCAGCTCCCTATCGTCGGAAACCTATTTTCTCTCGGCAACAACCCCCACCGCTCCGCTGCCCATCTTGCTAATGTCTATGGACCCCTCGTGACCCTCAAACTCGGCCAAGTCACCACCATCGTCATCTCCTCCCCTGAAATGGCCCGCGAAATCCTCCATAATAAGGGACCAAGCCTTCTCTTCTCACTCGGCCCCTGAATCGGTCCGGGTTGACGGCCACCACGAGGCGTCCATGCCGTGGCTGCCCGTTTCACCTCTGTGGAAGAAGCTCCGCAAGATCTACGCCGTCCCACCTATTTGCGCCTGGAAAGATGCAGCTCAACAAAGGCCTTCGGGTCGACAAGGCGAGGGAGCTCATGAGGTACGTAGGCCGACAATGCGGTGCTCCAGTGGACATCGGGAAGGCCGCATTTATGACGTCACTGAATTTGATCTCGAACATATTGTTCTCGATCGATGTGGTCGATCCGCTGGCTGTCGGGGATTACGGGGGTCTGGATTTTAAAGGGGGTCGTGTGGGGGATCATGATAGAAGCCGGGAGACCGAACTTGTCGGACTTCTTCCCGGCGTTGGCGGGGCTCGACCTCCAAGGTAGGAGGCGCCGCGTTCAATCACTCTTTAAGCAAATGGATGGGGTGTTTGATGAATTGATAGCCAAGAGATTGCAAAGCTCGGAAACATACAATGACTTGTTGGATTGGCTCATCAAGTGTCAAGACAAAGAAGATGGATTGGATCTCCACGGAGTCATCCTAAAATCTTTTCTTAAGGACATCTTCGTTGCCGGGAGCGAGTCGATCTCAAGCACGGTGGAATGGGCAATGGCGGAGCTGCTCGGCAACCCGAGGTCGATGACAAAGGCACGCTCTGAGTTAATCGACGTAATCACGGACGGCGACGAATTAGAGGAGCCCATCATCTTGCGTCTTCCGTACCTTAAAGCAGTCGTGAAAGAGACCCTGCGGCTCCATCCCCCGCTCCCCTTCCTGCTTCCTCACAAGGCTGAGACAGATGTGGAGCTACACGGCTACATTGTTCCCCAGAATGCTAGGGTGCTGGTGAACGTTTGGGCTATGGGCCGAGACGAACGAGTTTGGGAGGATCCAGATAAGTTCATGCCCGAGAGGTTTCTAGATAAGGATGTCGACTTCAAGGGTGCGGATTTTGAGCTGATTCCATTTGGTTCGGGACGGAGGATTTGCCCCGGGATGCCCCTGGGGTCTCGAATGGTGCACCTTATGTTGGCTTCGTTGCTGCATTCGTTCGAGTGGAAGATGCCAGACGGTGTGGATGTTGACATGAGGGAGAAGTTCGGCCTCACATTGGCTAAGGCTGTGCCGCTTCAAGCCATCCCCGCCTCGAGGGGGTTGGCTTCCTAGAAACATTATTGTTTTATATGGACATCAAATCCTTTTTTTTTATGTGCTCTATATAGTTCTTTCATAGGACGAATAATAACCCAAGTAAATTTAAATGGCCTTGAGCCTGCATC

>F01_transcript/45078 full_length_coverage=3;length=1685;num_subreads=60

GAACACACTGTCCACCAGCAAGCCTAGAATGGTGTTGGAGCTCGCTGCCGTCGTCGCCGTATTCTTGGTTTCCTGTCGTCTTCCTTCTCGGATTCCGGCGAGGTGTCGGCGGGGCAGTCGAGAATGCGCCGCCGGGATCGTTCGGGTTTCCAGGTGGTGGGGGAGTCGTGGAGCTTCGTCCGGGCCCTGAAGCAGGACCGGGGCGGCGAGTGGGTTCGGAAGAGGGCGGATATGTACGGCCCGGTCTTCAAGACCCACCTCATGGGTTGTCCCACCGTCGTCGTCACCTGGCCGCGCGGGAAACAGGTTCGTCTTCAGCACCGACCACGAGATCCTCAGCGTGAAGCAGCCACCCTCGATCGCCAGGATCGGGGGCAAGCACAACATCTTCGAGCTCACCGGAGAAAGGTATAAGCTAGTGAAGGGTGCATTGGTGAGCTTTCTAAAACCCGAAAGCCTTCAAAATCATATCGGACCCATGGATGCTATAGTGAGAAGCCTTATTCTTAGCGAAACAAAAGGGAAGGATCAAATTCTTGCAGTCCCGTTCATGAAGAGGATCACGTTCAATCTGACATGTGCCCTACTATACGGACTGCATGATGATCAAACCAAAGAGGTTCTCTTGAAGGATTTTTCTGAGGCCTTCAAAGGAATGTGGTCCTTTCCTCTTAACCTGCCTGGTACGGCCTACCGCAGAGCTTTGAATGCGAGGTCAAGAATTGATAAGTACGTGCTGGCACTCTTGCGGAGCAAAAGAGAGCGTCTGCTCGAGGGGAGTGCAAGTGTGAAAGACGACGTCGTCTGTTGCCTGCTCGCATTGAGAGACGAACATGATGAGCTCATTGACGAACAGACGATAGTGGATAACTTCTTTTCTTTAACGATTGCCAGTCATGATACATCGGCAATTCTATTGAGTTTGATGGTGTGGAAATTGGCTAGAGATCCCATGATTTACCAAAAGGTTTTAGAAGAGCACCTGGGAATCATAAGAGAGAGGAGTGAAGGGGCACAAGGGAAGCTTACATGGAGTGAAGTACATAAGATGAAATATACTTGGAGAGTTGCGCAAGAGCTGATGAGGCTGATCCCTCCGGCATTCGGTAACTTCAGGAAAGCTCTGAAATCCGTCAAGTATGGAGGTTATGAGATTCCAGAGGGATGGCAGGTGCTATGGGAGACTAGCGGAACCCATCACGACAAGGACATCTTCGAGGATCCTACTTCCTTCGATCCATCTCGTTTCGAGAATCCGTCGAAGCCGATTCCACCCTTCGCCTACATCCCTTTCGGAGCAGGGCAGCGAATGTGTATAGGCAATGAGTTTGCGAGAGTTGAAGCCTTGACTGTGGTGCATCACCTAGTTACAAACTTTGAGTGGTCTCAGGTGTACCCTGATGAGCCGATCACACGCGATCCCATGGCCTATCCTTCCAAGGGTCTGCCAATCAAGCTCAAACCAAGAAAACTTTGAGAGTTTATATATTGAGTTCATGTGTTAAGTATGTAATGCATTGGTTTCTTGTTCAGCACTGATGAATAAGTTGCTCTTAGTTCAGCAAGGATCCAATAGCAATGGGTTGGTTGTAATTGTCTTTGAATGGATGAAGCTCGATGATGAATTTCTGAAGCAGGAGGTTGGACGCCAATTAAATACGTGGATTATTATTATTGTCTAAACT

>F01_transcript/45178 full_length_coverage=13;length=1704;num_subreads=60

GATAGAAGCAACACCATATTAAAACCTAGCGACTCTCTCTCTCTCTCTCTCTCTCTCTCTCATTCTCATTCTCATGGATGCAATAATCTATCTAGTTCTCTTCATTGCCTTCATACCAATCTTACTTCTCCTCACAATACGAACAAAAGGATCGGCACCGACTAACAAGTTGCCACCTGGATCCTTGGGGATCCCCGTCGTCGGACAGAGCCTCGGTCTCCTCCGCGCCATGCGGTCGAACACCGCCGAGGAATGGCTCGAGCAACGGATAAACAAGTACGGACCCATATCTAAACTGTCACTCTTCGGGGTGCCGACAGTGTTCCTCACAGGGCCGGCGGCAAACAAGTTCATATTCACCACCGAGTGGCTGCCGATGAATCAGCCGGGGTCCATCACACGGATAATGGGGCGACGGAACATGATGGAGCTGCATGGGGACGACCACAAGCGAGTCAGGGGGGCCGTCAGCCGATTCCTGAGGCCCGAAGTGCTGAAGCAGTATGTTGCTAAGATGGACTGGAGAGGTCAGACATCACCTTAACATGAACTGGATCGGTCGACAGACGGTCACGGTTTTGCCATTGATGAAAAGGCTAACATTTGACATAATCAGCTCCCTAATATTTGGGCTGGAGAAAGGACCCATTCGGGAGAGACTGGGCAGTCATTTTCAGGAGATGATCGAGGGGATGTGGGCAGTTCCGGTGAACTTACCGCTCACACGTTTCAACAAGAGCCTGAAGGCGAGTCGGAAAGCGCGGGAGGTGATCGTCGAGATCATCGATCAGAAAAAGGCGGCACTGAAGAATGGTAGCTGTGGGCAATACGACGATCTCATCACCTACTTGCTAAGCTTGAGTGTTGACAAAGAGACCACAATGACAGAGGAGGAGATCATCGACAATGCGGTGCTTGTCATGGTTGCCGGGCACGACACCTCGTCTATTCTCATGACCTTCATGGTCCGACGCCTCGCTAAAGATCATGCAACATTGGCTGCCGTCGTTCATGAACAAGAGGAGATATCCAGGGCCAAGGCCCCAGAGGAACCCCTTATGTGGGATGACCTTACTAAAATGAAGTACACATGGCGTGTCGCGATGGAGACACTAAGGATGATCCCTCCAGTGTTTGGGAGCTTCAGAAGAGCACTCAAAGATTTGGAGTTTGGTGGATACTTCATACCAAAAGGGTGGCAAGTGTTCTGGGCTTCCAGCATTACACAAATGGATGCGAATCTCTTCCATGATCCCCAAGTATTTGAACCAATGCGGTTCGAGAGCCAATCATCAGTTCCACCATATTGCTTTGTGGCATTTGGAGGAGGGCCGCGGATTTGCCCGGGCAATGATTTCGCAAGGATAGAAGTACTGGTCACCATGCACCATTTGGTGACACAATTCAGATGGAAATTGTGTTGCAAAGATGATGGGTTCTGCAGGGACCCATTACCATCGCCTTCGCAAGGACTTCCCATCCAACTTGAACTCAAGGCCGATCCTTTGGCAGCTTGTTCGATCTTAGAGTTTAAGCAAGTAAATTGCGTTATGGGATTCTTCTTAAGAACCTTTAAAATAATAAGTGAAGAATCTCGGTTTGTTGTTTCTAGCTTCTTGTGGTGTTATTGATTATATAAAGGCTTGTTAAAGGCACTGGTTTTAATTCCACGTCGCATAGGGCATGACGTTCTCTTCGTTTTAG

>F01_transcript/45501 full_length_coverage=21;length=1699;num_subreads=60

ATAGAAGGCTTGCTCTTACTAACAACCACAGCCAAAGATTGAAAAAAAGATGATCGCTCTTCTTCCCCAACTGCTGCTCCACTATTTTCAACATCATGCCATCCTCATAGCATGCCTCTCCCTCCTCACCATGTCATTCTCTCTCTACAAACTAGGCCGACGCAATAAACCATCCAAGAAACTGAACCTACCGCCTTCTCCCCCAAAGCTCCCGCTCATCGGGAACCTCCACCAGCTCGGTTCCATCCCCCACCAAACCCTTCGTACCCTCTCCAACAAATACGGCCCGCTGATGCTCCTCTACTTGGGACGCATCCCCACCGTCGTCATCTCTTCCACCGCTGTGGCCGAAGAGATGACGAAGACCCACGACCTTGTCTTCTCGAGCCGTCCCTCATCAAGCATATACAACAGGCTCTTCTACAACTGCCAAGACGTCGGTTTTGCGCCCTATGGCGAGTATTGGCGTCAGGTAAGAAAGATCTGCGTCCTCCACCTCTTGACCCTCAAGAGGGTCCAATCTTTCCGATCTGTGAGAGAGGAGGAGGTCACGATCATGGTAGAGAAAATACGGAGCAGCTCGGCTTTGGGGCCGGTCGACATGACGGCGACGTTGATCACGCTTACATATGATATAATATGCAGGGTGGTGCTTGGGAAGAAGTACCCGAGCACCGGAGCCCGCAAGATGCTGGAGGAGGTCATGGTCCTGATGGGGGAATCCCCTGTTGGTGACTTCATACCAGGATTATCGTGGTTGGATCGGTGGAGCGGGTTGGATGCGAGGGTGGGGAAGACCTTCAGTGAGGTTGATGCCTTCTTGGATAAAGTGGTGGAGGACCATATCCATGGTGAGAAGTCTCATGATGGTGAGGACTTTGTTGACGTTCTTCTCTCATTGGATGCCAAGGGATCAGATGATATGGGTGGCTTCTCTCTCACCAGAGACTGTATTAAGGCTCTGATTCTGGACATGCTTGTTGCTGGAACCGATAGCACCTTCACGGCGTTGGAATGGACAATGACAGAGCTCATCAGACACCCAGAATCAATGAAGCGTGTACAGGAGGAGGTACGTAGTGTTGAAGTGATAAAAGAGAAAGAGTTGGATCACATGAGATATTTAAAGGCGGTAATAAAAGAAGTCCTCAGGCTACACCCTCCGGGAACTCTTCTTATTCCTCGTGAAACCACAGAAGATGTTCGGTTGTATGGTTATGATATTCCTGCAAGGACGAGGGTCTTAGTGAATGCTTGGGCGATCGGAAGGGATCCCAAATCATGGGATCTACCCGAAGAGTTTCGACCGGATCGATTTATGGACAGTTCAGTTGATTTCAAAGGTCATGACTTTCAATTAATACCGTTTGGTGTAGGGCGAAGAGGTTGCCCTGGAATTGAGTTCACCGTCCCCACGCTCGAGCTTACACTTGCCACTCTCCTTCGTCATTTTGATTGGACTTTACCCAGTGGAACAAATATGGAGACTATGGACATTACTGAAAAATTTGGATTAACAACACGTAAAAAATCTAACTTGATTCTTGTACCGAAACTTATTTGACGCATGTAGTTTGATATTAATATTGTAATAAAGTGTTGACATATGAAAATGGTCAAGACCATGGTGGTGAAATCTATATGACGCATGTAGTATGATATTGATGTTGCAATAAGATGTTCACTTATGAAAATGGTC

>F01_transcript/45551 full_length_coverage=9;length=1693;num_subreads=60

GATCATCACATTACATTATCGGCCATGAACATCACCTTCTACTTCCTCTCCTGCGCCCTCGCAACCCTCCTCTTCTTCATCATCCAGCTCATATCCCCTTCATCCTCCACCAAGATCAAGAACCCGCCCCCCAGCCCCCTTTCGATCCCCTTCATTGGCCACCTCCACCTCATCTCGAGCCCCCTCCACCATGTCCTCTCCCGCCTCTCGGCCCGCCACGGCCCCATACTCCACCTCCGCCTCGGTTCCTGTCCCACACTCGTCGTCACCTCCCACGCCCTCGCCGAGGAGTGCCTCGTTGCCAACGACCTTGCCTTCGCCAACCGCCCCCGTCTCGCCACGTGGCGCCACCTCACTTACGACTACACTCACATAGGTACCGCCAACTATGGCCCTCTCTTGCGCGACATGCGCCGAATCGCTGCAGTCGAGCTGCTCTCGCCGCACTGCCTCAATTCCTTCACCAGTGCCCGCGAGGTGGAGGCACGGGCAATGGCGCGAAGACTCTTCAGAGAGGCGGCGGACGGGTTCACGAGGGTGGAGATGAAGTCGAGGCTGTTTGGCCTGACAATGAACTCGACAACGGTGATGCTCGTCGGAAAAAGGTACTACGACTACGAGGCAGGAGACACAGCAGAGGCACGAAGTGTACAGAAATTGCTGGAAGAAAACTTTTCCCTTCGGGCATTATCCAACCTCCGGGGCTTCCTACCGGTGTTAAGGTGGTTGGATCTGGGAGCCATGGAGAAGAAACTAGTAATGTACCGAAAAATTATGGACACTTTCTTCCAAGGTGTCGTTGATGAGCACCGGAGAGATATTGACAAAAAGAATGAGGAGAAAGAGGCTATTATGATCAATGCAATTTTGTCCTTGCAGAAGTCAGACCCGGAGCGCTACACCGACAATTTCATCAAGCCCCTTATCATGAATTTGTGGACGGCGGGAACCCACACTTCAGGGAATACATTGGAATGGGCTTTGTCCCTCCTCCTAAACAATCCACGTGTAATGAAGAAGGCACGCGCAGAGATCGACGATCATGTCGGAAATGGGCGCTTGCTACAAGAATCAGACCTCTCTAATCTTCCATATATCCATTGCATCATCACTGAAACCCTTCGATTGTATCCGGCAGCCCCACTCCTCGTCCCGCATGTATCATCCAAGGAGTGTGTCATCGGGGGCTTCAACATTCCACGCGGAATGATGCTATTGGTCAATGCCTACGCCATCCACAGAGACCCCAACGAGTGGGTGGAGCCCACCAAGTTCATCCCCGAGAGGTTCGAGAATGTTGAAGGGTTTGGTCGAGGTCGCCCGATGATTCCATTTGGGATGGGAAGGCGTCGATGCCCAGGGGACAGGCTCGCGATGCGAGTGATGGGCCTAGTGTTAGGTATTTTGATACAATGCTTCGAGTGGGAAAGGGTCGGCGAGGAAGCGGTGGACATGACCGAGGCTTCATCACTACTTTTACCAAAGGCCATCCCGTTAGAGGCCATGTATAGACCCCGTCCGACCATGGTGGATGTTCTTTCGTCGCTTTAGATGAATATCAAAAGGAAGTCAAGAATTTCAATCCATAATTGAAATATATATGATGTATTATGAAAAATTATTTCCAAAAATTTCGATATATATTTGAAATATATTTGATGTATTAAGAAAATATATTTTTGCAAGTTTGCCT

>F01_transcript/45715 full_length_coverage=2;length=1677;num_subreads=60

GAACAACATACTAGCAGCCTAGCATAATGTCCCCAACAATCCTCCAATGCATCTTCCACCCATACTTTCTCCCTCTTCTCTTCTTGTTTCCTCTCTTTCTACTTATAACAATAATAAGAAGCTTCTCCACTAAGATCAATCTCCCTCCATCACCACCAAAGCTTCCCATAATCGGGAATCTCCACCAGCTCGATCGACTCCCCCATCACTCTTTGCGTGCTCTCTCGGAGAAGCATGGGAACCTCATGCTTCTCTACTTAGGTCGCGTACCAACTCTGGTGGCCTCGTCTGCAGACGTGGCTCGTGAGATCCTGAAAACTCATGATCTCACCTTCGCCGGTCGGCCTCCCATAAAAGCCATTAGGGTAATACATTCTGGAGGCATGGGTTTGGGCTTCGTGCCCTACGGGGAGCATTGGAGGCAAATGAGGAAGCTCACCATCACCCACCTATTGAGCACCAAGATGGTGCAGTCCTTTCAGCTCGCTCGTGAGGAAGAAGTGGAGTTCATGATCGACAAGATTTTATCCCGATCGTCGTCGATGGTCAATATGAGCGACGTGTTGTACTCGTTCACGAACGACGTCATCTGCAGGGTGGTTTCCGGGAAGTTCTTTAGGAAAGAGGGAAGGAATAACATGTTTCGACGATTGATAGAGGAGTACGAAGTTTTGCTTGGGAAGTTCCATGTGGAGGACTTCCTCCCATGGCTTTCATGGCTTGATGTATACTTCGGCTTGAATTCAAGAGTCGAGAGGAACTCCAAGGATTGGGATGATGTTCTTGAAGAAACTATTAGGGATCACGCCGAGCGAGCAAAAGGCAATGAGTATGAGGGCGACCACTTCGTGGATGTGTTGCTCTCACTTCAGAAAAATCCCAACATGGGCGACATCCTCAACAAGGGCAACATGAAAGCACTCCTAATAGATATGTTTGCCGCGGGAACCGATACATCATACATACTTCTCGAATGGGCGATGGCTGAGCTTCTTCATAAGCCGAAAGCAATGAAGAAACTGCGAGACGAGGCACAAAGAATATCAAACGGAAAATTGGTAATCCAAGAGGAAAATCTGAATGAAATGAGCTACTTAAAAGCTGTGGTGAAAGAAGTCCTTCGACTTCACCCTCCAGCTCCATTGCTCCTGCCGCATGAGTCGAGGGAGGAATGCAAGATACAGGGCTATAAAATCCCCAAAGGAGCAAGAGTCATCATCAATGCTTGGGCAATCGGAAGGGACCCCAAAAGTTGGGATTCACCGGAGGAATTCCTGCCTGAGAGGTTCCTTGTTAGTGGCTCGCCGGATTTCAAAGGGAATGATTTTCAGTTCATCCCTTTTGGGGTGGGGCGGCGGATTTGCCCAGGAATGAACTTTGCTGCCACCACTGCAGAGCTTGCGCTGGCAAATCTTGTGCATCGGTTCGATTGGGAGCTGCCTGTTGGCACTGCGAGCGAAGTGCTTGACATGGCCGAGGGTGAAGGGATAACAACCAGGAAGAAAGAGAGACTTCACCTCATTATGAAGCCTTTGTTAACTTAGTAGTTTGAATGGATTGTTAGTCTTAAGGCCTAAGAATTGGCAAGCTAATAATATCTCGACTTGGTAGTTTGTAAAAAAAAATTATTATTATATATGTGTGCTAATATTTGAATGTGTGAGCCTGAGATTCAGC

>F01_transcript/46486 full_length_coverage=2;length=1651;num_subreads=60

ACCCTTTACAGAGAGAGATTAGAGAGAGAGAAGAGAGAGAGAGAGAGAGAGAGAGAGGGGTAAGTCTCTCAAAGGGGAGTGCTCCTACTCCTACCATGGCGTTGGAGTTGATCCTGGTTCTGTCTTCCTTGATCGTGATTCTCATCATCTTCTTCAGCTTCAAGAGTAATGGGAAGAGTGAGAGCAAATTGGCCAAGCTCCCACCGGGCCAAATGGGCTGGCCCTTCATCGGCCAAACCATCCCCTTCATGCAGCCACACTCCTCCGCCTCCCTCGGCCTCTTCATGGACCAAAACATAGCCAAGTATGGGAGGATCTTCCGGACGAACTTGCTGGCGAAGCCGACCATCGTGTCAGCGGACCCGGACTTCAACCGGTACATCCTGCAGAACGAAGGGCGGCTGTTCGAGAACAGCTGCCCCACGAGCATCAAGGAGATCATGGGGCCGTGGTCGATGCTCGCCCTCGCGGGCGACATCCACCGCGAGATGCGATCCATCGCGGTGAATTTCATGAGCAACGTCAAGCTCCGAACCTACTTCCTCCCTGACATCGAGCAGCAGGCCATAAAGGTCCTCGCCAGTTGGGAGAACACCCCCGAGGCCTTCTCGGCCCAAGAACAAGGAAAGAAGTTTGCGTTCAATCTGATGGTGAAGCATCTGATGAGCATGGATCCCGGGATGCCGGAGACTGAGAAGCTGCGGACGGAGTACCACGCATTCATGAAGGGGATGGCTTCAATTCCTTTGAATTTGCCCGGAACGGCTTATAGAAAGGCCTTGCAGTCGAGGTCAATAATTTTGAAGATCATGGGGCAAAAGCTAGACGAAAGAATCCGACAGGTACGTGACGGGTGCGAGGGATTGGAGCAGGATGATCTCTTGGCGTCCGTCTCCAAGCACCCACACCTCACAAAGGAGCAGATTCTCGATCTCATACTCAGCATGCTCTTCGCCGGTCACGAGACTTCCTCGGCTGCCATCGCCCTAGCTATCTATTTTCTCGATTCGTGCCCTAAAGCCGCCCAACAATTGAGGGAGGAGCACGTGGAGATCGCGCGACAAAAGGCTGAGCGAGGCGAGACGGGGCTCAATTGGGACGACTACAAGCAAATGGAGTTCACCCATTGTGTCATAAATGAAACCCTAAGGCTCGGAAACATTGTCAAGTTCTTGCATAGGAAAAACTCTTAAAGATGTCCAATTCAAAGGGTATGACATTCCATGTGGGTGGGAGGTGGTGACGATCATCTCAGCCGCTCATTTGGATCCGTCCGTGTATGATGAACCTCAGCGTTACAATCCATGGAGATGGCAGGTTAGTGACTCCATAAACTCCATACACATTCATCGCATGAGAACAGCATCATCAATTAGATAGGAGGGTTTTTGGCGTGGACCCCTACCAGAAATAATTTTTTATTTTTATTTTTTATTTTTATTTTTTTTAATCATGTGAAGATTTTACCAGCTGAACTAGCTGACATCTCCAACTTCACGTGACTACAAGTCTACAATTAGCTCCTAAATATTTTGACGGCCCTGATCTAATTTATAAAGTCAAGCAATGCAAATTCATTTATTTGGGTTTTATTTAAGTTCACGGTATTTTTATATACACCCCACAAACAAATAAAGTATTGATAGTTTTC

>F01_transcript/47119 full_length_coverage=2;length=1630;num_subreads=60

ATACTCTCCTCATGCAAGCTAGAGTATTAAGATCCACCAACAGACTTCCACCTGGCCCTCGGCCACTCCCCATTGTCGGAAACCTGTTCTTGCTCGGCCAAACCCCCCACCGCTCCCTTGCCCGACTCGCTGACACATTCGGCCCCTTAATGTCCCTCAGGCTCGGTCAAGTCATCACTGTCGTCGTTTCATCACCCGAAATGGCCCGTGAAGTCCTCCACAATAGGGACCAAGCCTTCTCATCCCGCTCGGTCCCCGACGCCGTCCGGATCAATGGCTACCACGAGGTATCCATCGCGTGGCTGCCTGTTTCACCACTATGGCGGAAGCTCCGGAAGATCTACGCTTTCCACCTCTTCTCCCCTCGGAAGATGGAGCTCAACGAAAGCCTCCGGATCGAGAAGTTGAAGGAGCTGGTGAGTTACGTGCACCAACGCTGCGGTGCTCCAGTGGACATTGGGCAAGCCGCGTTTATGACTTCCCTGAATTTGATATGGAATATGCTGTTCTCCATTGATGTCATCGATTCAGTGGTTGGATATTCAAATTTTAGAGGCGTGGTGTGGGGGGTAGTGAAAGAGGCCGGGAGGCCAAACTTGTCGGATTTCTTCCCGGCGATGGCGGGGCTGGACCTCCAAGGTAGGAGGCGCCGGGTTGGCAAACTCTCTAAAAATATGGATGGGGTGTTTGAGGAATTTATAGCACAGAGATTGAAGAGTTCCACAACATATAATGACTTCTTGGATTGGCTCATCAAGTGTCAACACGAAGAAGATGGATTGGAGCTGCATGGAGGCATTGTCAAATCTTTCCTTAAGGATATTTTCATCGCTGGAAGCGAGTCGACCTCTAGCACAGTGGTATGGGCAATGGCAGAGCTGCTCCGCAACCCGAGATCGATGGCAAAGGCACGCTCCGAGCTTATCGATGCCATCGGGGACGGGAACGAAATAGAGGAGCCCATCATCTTGCATCTTCGGTACCTCAAAGCAGTCGTGAAAGAGACGCTCCGTCTCCACCCCCCGCTCCCCTTCCTTCTACCGCGTAGGGCCGAGACTGACATCGAGCTAAATGGCTATATCATACCCAAACATTCTCAGGTACTGGTGAATGTTTGGGCTATGGGCCGAGACCATCGAGTTTGGGACGATCCCGACAAGTTCTTGCCTGAAAGGTTCCTCGATAAGGATGTGAACTTCAATGGTGGAAATTTTGAGCTGATACCATTTGGTGCGGGGCGGAGGATTTGTCCTGCGTTAGGAATGGGGTCTCGGATGGTTCATCTTATGTTGGCTTCGCTGCTATATCCATTCGAGTGGAAGGTGCCGGACGGGGTCCATGTTGATATGAAGGAGAAGTTCGGCCTCACATTGGCTAAGGCTGTGCCTCTTCGAGCTGTCCCCATCTAGAGAGCGTCCTCATCATAAAAACGGCCTTTCTGGCACCTTCATTATTACTTTGTATAATAGCCAACTCAGCAAAAGGATTGGTATCAATAATTGCTTATTACGAGGGCCCTATACGAGCACTCGGAATGGGGACTCTTGGTAATAAAAACTTAGTAAAACTTTAAAATTTTGCTTGTAAAATTTTATTGTCGTGAGTATCATTTACTTTTATTTTTGTGCAG

>F01_transcript/47155 full_length_coverage=5;length=1633;num_subreads=60

GATAATGCAATAGGTTGGTAGCAACATACTAGCAGCCTAGCATAATGTCCTCAACAATCCTCAATGGTCTTCCGCCCATACTTTCTCCCTCTTCTCTTTTTGTTTCCTCTCTTTCTACTTATAACAATAATAAAATGCTTCTCCACTAAGATCAATCTCCTTCCATCGCCACCAAAGCTTCCCATAATTGGGAACCTCCACCAGCTTGATCGACTCCCCCACCGCTCTCTGCATGCCCTCTCGGAGAAGCATGGGAATCTCATGCTTCTCTACTTAGGTCGCGTACCGACTCTGGTGGCCTCATCTGCAGACGTGGCTCGTGAGATTCTTAAAACTCATGATCTCACCTTCGCCAATCGGCCTCCCTCAAAAGCCATCAGGGTAATATATTCTGGAGGAATGGGTTTGGGTTTCGTGCCCTACGTGGAGCACTGGAGGCAAATGAAGAAGCTCACCATCACTCACCTATTGAGCACCAAGATGGTGCAGTCCTTTCAGCTCGCTCGTGAGGAAGAAGTGGAGTTCATGATCGACAAGATTTTGTCTCGATCGTCGTCGATGGTCGATATGAGCGACGCGTTGTACTCGTTCGCAAACGACGTCATCTGTAGGGTGGTTTCCGGAAAATTCTTTAGGAAAGAGGGAAGGAATAACATGTTTCGACGATTGATAGAGGAGTACGAAGTTTTGCTTGGGAAGTTCCATGTGGAGGACTTCCTCCCATGGCTTTCATGGCTTGATGTGTTCTTCGGCTTGAATTCAAGAGCTGAGAGGAACGCCAAAAAATGGGATGATGTTCTTGAAGAGATTATTAGGGATCATGCCGAGCGAGCAAGAGATAATGAGCACGAGGGCGACCACTTCGTGGATGTGTTGCTCTCGCTTCAAAAAAATCCCGACATGGGCTACATCCTCAACAAGGGAAACATGAAAGCACTCCTAATAGATATGTTTACCGCGGGAACCGATACATCATACATACTTCTTGAATGGGCGATGGCCGAGCTTCTTCAAAATCCGAAAGCAATGAAGAAACTGCGAGACGAGGCACGAAGAATATCAAACGGAAAATTGGTAATCCAAGAGGAAAATCTGCATGAAATGAGTTACTTGAAGGCTGTGGTGAAAGAAGTCCTTCGACTTCACCCTCCTGCTCCATTGCTCCTTCCGCATGAGTCGAGGGAGGAATGTGAGATACAAGGCTATAAAATCCCTAAAGGAGCAAGAGTCATCATAAATGCTTGGGCAATCGGAAGGGACCCCAAAAGTTGGGAATCACCGGAGGAATTCCTGCCTGAGAGGTTCCTTGTTAGTGGCTCGCCGGATTTCAAAGGGATTGATTTTCAGTTCATCCCTTTTGGGGCGGGGCGGCGGATTTGTCCAGGAATGAACTTTGCTGTCTCTACTGCAGAGCTTGCGCTAGCAAATCTTGTGCATCGATTCGATTGGGAGCTGCCTGTTGGCTCGGCGAGCCAAGGGCTTGACATGGCCGAGGGTGAAGGGATAACAACTAGAAAGAAAGAGAAACTTCACCTCATTATGAAGCCTTTGTTAACTTTGTAGTTTGAATGGATTGTTAGTCTTAAGGCCTATGAATTGGCAAGTTAATAATATCTTGACTTGGTAGTTTGTT

>F01_transcript/47351 full_length_coverage=2;length=1622;num_subreads=50

CTGGCCGCGGTCCGCCACTACCACCGGGACCTATAGGATGGCCGATCCTCGGCAACCTCCTGCAACTCGGGACGAAGCCTCATCACACCCTCGCCGCCATGGCCAAAACCTATGGCCCGCTCTTCCGCCTCCGACTCGGATCCGTCGAGGCCGTGGTCGCGTCGTCTGCCGCCGTGGCGGAGGAGTTCCTCCGCACCCACGACGGGAACTTCTGCAACCGACCACCCAACGCTGGTGCCGAGCACATCGCATATAATTACCAAGACCTCGTCTTCGCGCCCTACGGCCCAAGGTGGCGGTCCCTCCGCAAGCTCTGTGCTGTCCACCTCTTCTCTCAGAAGGCCCTCGACTCCCTCCGCCCCATCCGTGAGTCGGAGGTCCACCGCCTCGTCTCCCACCTCCGCTCCCTCGCTGCACAGAAAATCAACCTCGGGGCGGCGCTCAACGCCTGCGCCACCGACGCCCTCGGCCGCGCCACCGTCGGGAAGCGGGTTTTCGGCGACGAGGCGGCGGTGGAGGAGTTCAAGGGGATGGTGGTGGAACTGATGAGGCTCGCGGGGGTGTTCAACATCGGGGATTTCGTGAAGGGATTAGCGTGGCTCGACTTACAAGGGGTTGTCGGGAAAATGAAGCGGTTGCATCAACGGTACGACGTATTTCTCGATAGGATTCTCGAGGAGCATCGAGCGGCGGGGGGCGAGGGAGGAGGGGACCTCCTGAGTGTCCTCATTGGGTTGAAGGAGGACGTCGGCGGAGATGCGGTTAAGCTCACTGACACTGAAATCAAAGCCTTGTTTCTTAACTTGTTCACGGCCGGGACCGACACGACCTCTAGCACCGTGGAGTGGGCTCTGGCCGAGCTGATCCGGCACCCGGACATCCTAAAGCAAGCCCAGCTCGAGATCGACTCGATCGTAGGTCGGGCCCGGCCCGTTTCAGACTCGGACCTCGCAAATCTCCCTCTAGTGACGGCCATCGTGAAGGAAACATTCCGGCTCCATCCGTCGACCCCCCTCTCCCTACCACGAGTGGCCTCTGAAGCGTGCGAGGTCGGGGGCTACCGTGTCCCCAAGAACGCCACCCTTCTCGTCAATGTGTGGGCCATCGGGCGCGACCCGGCCTCTTGGCCCGACCCGTTAGAATTCCACCCGGCCCGATTCCTCCCTGGTGGGGATCACGAGGGGGCCGACGTCAGAGGGAACGACTTCGACGTCATACCCTTCGGGGCCGGCCGGAGGATCTGCGCCGGCATGAGCTTGGGCCTGCGGATGGTTCAGCTCATGACAGCGGCCTTGATCCACGCCTTCGATTGGACCCTCCCGGATGGCCAGGTTGTAGAAAAACTCGACATGGAGGAAGCCTACGGGCTTACGTTGCAAAGGGCCGTTCCTCTAGAGCTCAGGCCCATTCCGAGATTGGCCCAACAAGTATATGTGAAAGATTGTCAGGAGGCATAACAGTGTTAGAGCACTCTGTTGCTCTTCTCTCCAATGGTTAGCTTGATGGCTAAAGTATCACAGTACGTCAATATTTTGAGACGAGACTTTTGCCCTGTATCCCTATATTTAGTCTATATTTTTGTTATATTCTTAAGCTTTTTAATATTGCATTCACATCATATT

>F01_transcript/48619 full_length_coverage=5;length=1606;num_subreads=60

GGTGTGTGAGAGAGAGAGAGAGAGAGAGAGAGAGAGAGAGAGAGAGATGGAGGTGATAATCGTTGTTGTGTTAAGTTTGGTAATCCCGGTGGTGATTATGATAGTGTCCCAACAACAGTCGTAAGAGGAAGTTATCCAAGCTACCACCCGGAAACATGGGTTGGCCAATCATCGGAGAGGGCCTCTCTCCTCTCAAGGAAGAACCAAATCATATCTTGGGAGAATATTTGGAGACCCACATATCGAGGTATGGAAAGTTATTCACCACCAACATATTCGGATGGTTGGCGATCGTCTCAGTAGATGTTGAGCTAAAGCGCTTCATGTTGCAAAACGAAATGAAGCTATTTACTCATGCATGGCCGAGACATTTCAAGGTGTTGATTGGGGAATCCGCAACTATGTTCTTGACCGGAGATGCCCACAAGCACTCGAGGTCTAACTTTATCAAATTCTTCGCTGGGGCAAAGCTCCATAGCCCAATCTTGAGCCAAGTCGAGCAACTTGCAACAATGGCTATGAGCTCCTGGGAGAATAACTCCATCATACTTGCTAAAAATGAGATACTCAAGTTTGCCTTCAATACGATTGCAAAGGCGTCCATGGGCATGCGTGCTGAGGATCCTAAAACTGAGAAGCTGAGGCAACAATTCGACGTCTTTGTCGGGGGAATGTTTGCTGTAATTCCTTTCAAGTTCCCTGGGAATGCTTACTGGAAGGCTCTAAAGTCGAAAGACTACATTACAGAAGCTCTTGGCAAAGTAGTTGAAGAAAGAATTCAAATGTCTGCAGCAGTTGGAGTTGATGAGAATCTAGAAGAAGAGGACCTCCTTACTTGCCTATTGAATGACACTCGTTATACAAAGGAGCAAGTCATCCAAAATTTGCAGTTTGCGATTCTTGGCAATTACCTTACGACACCGAAAACAATGGCTACTCTAATCCATCAACTCGGAAAATGCCCTAAAGCAGTCGAACAACTTCGGGAGGAACAACTTGAAGCAATCAGGAACAAAGAGAAACAAGGGGATAGCAAATTAACGTGGTCAGACTACAAGAACATGGAGTTCACACAATGTGTTATAAATGAAGCACTTCGAGTTGCAAGCGCTGGAGTTATTCCCAAGAAGGCTATTGCAGATGTCAAATATAAAGGTTATATCATCCCGCAGGGCTGTCTTATCCTATCCCACAATGCAGCAATGAGTCATGATCCTGACTTGTACCATGATCCTAAGCATTTCAATCCCTGGAGATGGATTGGTGATGAATCAGGAGCAGATAAACCAAAGGCCACGTTCTTTGGTGGAGGTCCCAGGCAGTGCCCTGGTTCAGAGTTTGCGAGGTTGGAGATGACGGTCTTCCTTCATCATCTCATCGCAAAGTACAACTGGAAGCCGGTATCGCCTAACTACAATATTACGATAGCTAATGTCGACTCCCCTAAAGGTCTACCAATCAAGATTGAAGCATTAGTGTAAGCTCGGTCACCATTTCCAAGTTTATTGTAATCAGAGAGCAGCAGGCATGGAACTGAGCCTTTTGGGTCTTCAAATATTATCTATCGTCATTTTACATCAAAAAGCATCCCAAATATTTTGTGAGC

>F01_transcript/4993 full_length_coverage=2;length=3468;num_subreads=60

GGACACTCCAAACTGTCCAAACCCAACGTTGATTGCGTTGAGGCTTCATCTCCTCGCCTTCCTCTCATTCCTCTTCTCCCCCATTCTGCTGCAACTCATCAATGGCGTCCGCTGCCGTCACCGCCGCCGTCTCCCCTTCGCCCTGCCCCCCTCAGCTCCCGTCCAAATCCCATCTCCGGCGAGCCGCCGCCCTCTCCATCCTCTCTCGCGGCGGCGCCGTACGCTGCTCCGCCTCATCCAACGGCCGAAACCCTATCCCCGGCGGCGATCAGTCCTCCAAGGACGCCGATCGCCTCATCGAGAAGAAGCGCCGCTCCGACCTCGCCGACCGCATCGCCTCCGACGAGTTCACCGTCCAGCAATCCAGGTTGGTTTCTCTGCTGCGGAGGCTTGGACCGCCGGGGGAATTCTTGGCGGAGCTTCTGTCGAGGTCGGAGATCCCTCAGGCGAGGGGAGAGATCAGCTCCGTTGGCCGGACGGCCTTCTTCATCCCCCTTTACGAGCTTTTCCTCAGATACGGGGGAATCTTCCGCCTCACCTTCGGCCCCAAGTCCTTTCTGATTGTTTCCGATCCTGCTATCGCCAAGCACATACTCAAGGACAACTCCAAGGCTTATTCTAAGGGTATCCTAGCAGAAATTCTCGAGTTTGTTATGGGAAAGGGTTTGATCCCAGCCGATGGTGAAATCTGGCGTGTCCGAAGACGGGCGATTGTCCCGGCATTGCATCAGAAGTACGTGTCTGCCATGATCGGACTCTTTGGAAAAGCTTCATATCGGCTATGTGAGAAGTTAGATGCCGCAGCAGCCGATGGAGAGGATGTCGAGATGGAATCCCTCTTCTCGCGGTTGACACTTGATATCATCGGCAAAGCCGTCTTTAATTATGATTTTGATTCCTTATCACATGATAATGGAATAGTCGAGGCAGTTTACACTGTATTGCGGGAGGCAGAGCAACGGAGTACTTCTCCAATACCAACTTGGGAAATTCCTTTATGGAAGGATATATCTCCAAGGCAGAAGAAGGTCAATGTAGCTCTTAAGTTGATAAATGACACCCTTGATGATTTGATTGCTATCTGCAAGAGAATAGTAGATCAAGAGGAGTTGCAATTTCATGAAGAGTACATGAATGAGCAAGACCCAAGCATTCTTCACTTTTTATTGGCATCAGGGGATGACGTTTCTAGCAAGCAGCTTCGTGATGATTTGATGACTATGCTTATAGCCGGCCATGAAACATCTGCAGCGGTGCTAACATGGACCTTTTATCTTCTTTCTAAGGAACCAAGAGTCATGGCCAAGCTCCAAGATGAGGTTGACTCCGTTCTAGGAGACAGATTTCCAACCATTGAAGACGTCAAGAAACTGAAGTATACTACTCGAGTGATCAACGAATCACTGAGACTTTACCCACAACCACCAGTCTTAATTCGCCGTTCTCTTGAGAATGATGTGCTCGGGAAGTACCCTATCAAAAGGTAATGATTCTCTATTATATCTTCGTAAATTTATATCTGTTTGGAGTTTAACATTAACTTTATCCAATCTTCTATCTGAATGAAGAATAGCATTTCTACATATGTTGCAATTACATAAAAAATAAAAACCAGCTGACTATAGTCATCTGTTGCATAGTTATTTGTTGCAGAATTCCGGGGTTTTTTTTCCAATCTATTACCGGATTGGTTATGGAGCTTAATTGTGTACTTTGCACGAAGAAGATTCCTATCTGATTTTTCAATCATTATTATTTCGGTGTATCACGTGCTGCTTGATATTGGAGCTGTGTTCAGTTAATTCCTCAGAAAGGAGTGATGAAGGCTCTGATGAGATCAGAAAGGCAATGATTGTTTGCGTTCTTGAAATCTTTCTTATATTTCAGAGTTGTTACTGCGACATGTTTTATATTGTTCTAATGTGATGTTAGGCTTCATTGTGTGCTACAATGATCTTAACATGTATCCTTTACAAGCAATGAACACCCCACCATCTCCCACTCAAACCCAAGCTTCCGCTCCCGTGACATTGGACGCTGTGCAGCGCATGATCACTGAAAGCATGAGAGCCTACCATATGCAAAGCCAATATGCAATGAGGCCAGGCTACATGAAACCTTATCCTCCTGAGGTTGATCTGGTACTCTTTCCACCAAATTACCGACAACCCCAGTTCACTAAATGAAATGGTAATGGATCCCCGCATGAGCATCCACCATCTTTCTCATTTGGTACCAGCAGGTGGCAAAACGGTCATGAGATTTTGGTAAACTTCTAACATTATCTTCAATATATACGTGATGCCATGCGTTAAGCATCTAAAGTATTTCAACAGACCTGGGAAGATGATTTTTGGGTCTCAGTCAATGTGAATTGTTGGAAACTTGTGAAACACAATAAATCTACTATTCTATGGGAAACTATACCAATCGTATAATACTTGTTCCTAGCATATGACATAAAATATAATAAAGAATGGTCACCTGATGCACCCTTTTTAGTCATAGACCTAGTTATAAATTTTTGGTTCTAAGTGCAAATATATAGGTTAATATGGAACATGTGAGCTGGAAAGTTTTGGCTTCAACTTTACTTGATTATGTAGAGAATATCATTGATAATCATGGCTATTGACAAATGTTTGATTATTTCTTCATATGAAGCTCTTGTTCTTGTAGGCACATCATACAAATATGTTATATAGCATTTTACGGTTTTACCTACATTCTGTCTGCCCATCACAATATGCACATAGCACAGTATATTTCTTTCACCGTAAGATTAGAAGTAGAACCATCCTATGATATCATGAAAGAAATAATTTATGGCTTCATTGACATCTATACTGGCTTTAAGAGTATAAGTCTTTGTTCAGAATCTTGAGATTGTTCTTGTGTGGTTGACAGGGGTGAAGATATTTTCATTTCTTTATGGAACCTCCATCGTTGTCCTGATCATTGGGTTGATGCCGAGGGTTTCAACCCTGAAAGATGGCCTCTAGATGGACCAAATCCAAATGAGACTAATCAGAATTTCAGCTATTTACCTTTCGGTGGTGGGCCAAGGAAATGCGTTGGAGACATGTTTGCTTCATTTGAGACTGTGGTGGCGACATCGATGTTGGTGAGACGATTCAACTTCCAAATAGCCATCACGGCGCCTCCTGTGGAAATGACGACAGGAGCAACCATTCACACGACGGAGGGGTTGGTGATGACAGTAACACCAAGGATGCAGCCTCCGATAATTCCCTCGCTGGGGTTGAAGCGAGTGGTAACAGTAGAAAGTGATCAGCCTCAAAACTTGTCTCCCCCAGCTCCATTTCGAATCGGATAGCAGCCGCCTCATTAGATGCACTTGCCATTCATCTCTGGCAATGTAAATTTTCTCGCAACTTGTCTTTGTTATATTTCATGGAGAAATTAGCAAGCATGATGATCAATTCCACTCCAGAGAAATTTCTTTCC

>F01_transcript/22692 full_length_coverage=4;length=2423;num_subreads=60

GACATGATAGCCTCCATTTTCTCGCCCCCTTATGAGTTGCTCTCTAGAGTCCTCTCCAAGAAGAAATTATCTCCTCTCTTCTTCCATGAACGTTGGTCATTCTCCATCTTTATCTTCTCGAGAGCGACGTCATCGTCGATCCATCACACGAGGCATCGAGATAGCAAATAGAGAGATCCCTCTTTTTCTTCCTATGGATCTGACAATTGGATCGAGGTCTTCTTTCTCAATCTCGACGACTAAAAGCGACGTACGGTAAGGGGTGGCGGATCGAGGTCTCCTTTCTTAATCTGTGTTCCCTGGTGGCACATAACAAAGGGGCGACAGATCTAGCCACCCTTTCTTCTTGGGCAGCGAGATCGAGAGGGCCTTCTCTCGATCTGCGATCCAGATACAAAAGAGGGTCAATCACCCCCCTTCTTCCTCTTGTGTGGCGAGATCCTCGTAGGCGACGGGGGTCGTTTTCAGGATTCGCGCTTCCTCCACGCGGCACAAGAAATCGGCGGCAAGGTCGTCAGTTCTGCAGGCGAATGAGGTTGCTACCATGTGACCCATCCTCCCTTTTCAGCGACTAAGGTGGCGAGGAAGAGAACTACCCCCTTTTGTCTTCAATCCGATAGCCCGCCTGCCCGAGGTCAAATTTCGATGGGTGGTCGACCACCTATGGTCGATGGCGACCAGCAAAGAGAGAGGGGAGGAGGTGCCTCCCCTTCCTCGGTCCTCCTTCGACGGCGATGAGCACCTTGACCCCGAGCTCTCTCTCTCCCTTTCTCTTTCACACTCTCTCTCTCTCTCTCTCTCTCTCTCTCTCTCTCTCTCTCTCTCACACTCTCATCTCTCATGATGGCGTCACCAAGGATATCTCCAATCGCAGCGAGCGGATCGTCGCCATCCGCAGCGCATGGGTTGTCGCTGCCCGCGTTACACGGATCAAGACGATCGTTCATGGCGGGAATCAAAACTTGAGCTCAAAACAAGAGGTCAAGCCCAAGGACCGAATCTCGACGGATCGATCACAAGGGTGAGCGCCATAGTAGCGTCCTGATGGATCTCTTTGGTGAAGGGATATTCGCGGTGGACGGGGAGAAGTGGCAACATCAGAGAAAGCTTGCGAGCTACGAGTTCTCCACTAGAATCCTTCGAGAATTCAGCAGCGTCGTGTTTCGCTCCAATGCTGCAAAGCTTGCGAAGAAGATTGCAGGTTCTGCGAGCACTGGAGCTGTGATAGACATGCAGGAACTTTTTATGAAATCAACATTGGATTCCATATTTAAGGTCGGATTCGGGGTCGAGCTTGACACACTTTCTGAATCCGAAGATGTCGGAACTCGATTCAGCAAAGCCTTCGACAATGCAAACTGGATCACTTGCCACAGGTATATAGATATACTCTGGAGGCTCAAAAGGCGTCTCAACATTGGGTTAGAGTCCAAACTGAAAGAGGACATCAAGGTGATTGATGACTTTGTCTTCCAATTGATACACCACAAGAGGGAGCAGATGAAAAATGGGACCGAGGAAAAGGGCAAAGAAGATATTTTGTCGAGATTTATACTCGCGAGTGAGGAGGACCCAGAGCATATGACTGATCAATATCTGCGAGACATAATCCTCAATTTCTTGATCGCCGGGAAGGACACAACAGCAAACACTCTAACATGGTTCTTCTACTTGCTCTGCAAGAACCCCATAGTGCAGGAAAAGATCGTCAGCGAGATCAGAGACGCTGTGACATCGGAAGGTCAAAGCGCCGACATAATAGAATTTGCGTCCCAATTAACACAACCAGTGCTCGACAAGATGCAATATCTCCACGCTGCGCTCACAGAAACTCTAAGGATCTATCCCGCCATTCCATCTGAGGGAAAAACTGTGGAGGAGGACGATGTCCTTCCGGGTGGATTTAAGGTGAAGAAAGGAGATGGCGTGAGTTATATGCCATACGCCATGGGGAGGATGACATATCTCTGGGGTGAGGATGCCGAGGAATTCAGGCCGGAGCGGTGGCTCGAGAATGGGACCTTCCGGCCAGAGAGCCCTTTCAAATTCATCACGTTTAATGCGGGGCCTCGTATGTGCTTAGGGAAGGATTTTGCTTACAAGCAAATGAAGATTTCGGCGGCCCTCCTCCTCCATTTCTTCAAATTCAAACTTGAGGATGAGTCAAAGACAGCTACATATAAAACCATGTTGACCCTACATATGGATAAAGGCCTTCATCTTATTGCATCTTGTCGATGATGATATTAATATCGAAAGTCATAATGTATTTCTGTTAAGTGTTGTTTTATTTTGAAGATCAATGATAATGTGTGGTCTGTGAAATGGTACATTGCAAAACAAGTACTCATATGCAGTACAGTGGAATTATAGTTATGAATTGTGGTGCAATTCTTTAAGGTTGAGATGCAACTTTTATATCC

>F01_transcript/22874 full_length_coverage=3;length=2425;num_subreads=60

GTCTGCCATCTTTCTGCTCTTTTGTGAATGGCTTCAGATGACCCGTCTCTCTAACATTTCGATCCCAATGGATTCACGGGTCTTAGGGGCGCTGGCGGCGCTGCTGGCGGCGGCGGCCGCGTGGGTTATGAGAGCGGCGGCCGAGTGGTTGTGGTGGCAGCCGCGTCGGCTGGAGCGAGCCCTCCGGTCGCAAGGAGTTGAGGGAAATCCCTATCGTTTCCTCAATGGAGACTTGAAGGAGACGGTACGGCTCACCAAGGAAGCCCGGGCTCAACCCATCTCCCCACCATCTCACCGCTTCCTCCATCGTATCAACCCCCTCCTCCTACGCGCCATCTCCCATCACGGTAAATTTGCATTAACTTGGATAGGGCCCACTCCCCGAGTGAGCATTATGGACCCAGAGTTGGTACGAGAAGTTCTATCAAACAAGTTTGGTCATTTTGCGAAACCTAAGGCTGCCCCGATTGTCAAGTTGTTAGCTACAGGACTCGCAAACTATGAAGGTGAAAAATGGGTCAGGCACAGGAGAATTATTAACCCTGCTTTCCATCTCGAGAAGTTGAAGCGCATGTTGCCAGCATTTTTTTCTTGTTGCAGTGAATTGATAGGCAGGTGGGAGAGTTTGGTTGGCTGTGATGAATCCCGTGAAGTAGATGTTTGGCCTGAATTACAAAACCTCACAGGAGATGTCATATCTAGGACTGCATTCGGCAGCAGCTATGCCGAAGGGCGACGGATATTTCAACTCCAATCGGAGCAAGCTGAACTTCTTATTCAAGCCGTTCAAACTGTATACATACCGGGTTACAGGTTTTTGCCGACTCCGAAGAATATTAGAAGAACAAAAATTGACAAAGAGGTTCGAGCACTCTTGCGAAGCATAATTGAGAAGAGAGAAAATGCCATGAAAATGGGGGATGTACATGATGACTTGTTGGGTTTGTTAATGGAAAACAATTTGAAAGAAACAGAACATGGCAACTCTAAGAACACTGGGATGACTACTGAAGACGTGATTGAAGAATGTAAGTTGTTCTACTTTGCAGGGCAAGAGACGACATCAGTTTTGCTCACATGGACGATGATTTTGTTGGGTATGCATCCAAGCTGGCAGGATCGTGCAAGAGAAGAGGTTTCACAAGTATTTGGAAAGAGCAAACCAGACTTTGATGGCTTAAGCCGCTTGAAGACTGTGACCATGATCCTATACGAGGTTCTCAGGCTGTACCCCCCTTTTATTTTCCTCACCCGGAGAACATACAAGCCGATGAAGCTCGGAGGCATCACTTTTCCACCTGAAGTACAGCTCGCCCTGCCAATAATTTTCATTCACCATGATCCAGAATTCTGGGGTGAAGACGCCGAAGAATTCAATCCCGACAGATTTGCTGATGGTGTCTCAAAGGCATCGAAAAATCAGATGGCCTTCTTTCCGTTTGGTTGGGGTCCTCGTATCTGCATCGGCCAGGGCTTTGCAATGCTAGAAGCTAAGATGGGGCTGAGCATGATTCTTCAACGGTTCTCCTTCGAGCTCTCGCCAAACTACAGTCATGCTCCTCACACTAAGATTACTCTCCAGCCCCAGCATGGAGCTCAAATGGTGCTGCATAGGCTCTAAAACCTGCTGTTGATTGTTCCATGACTTTGAATTTGAAGACAGAATTGTGAAATCTGTATAGTTGTGCAGTGTATTTTACTATGTAGTGATAATTCGAATGAGTATTGGCAGTGTTCGCATTTCTCGACCTCGAACCCTCCCAGTCTGACGGTCCAGAGGCCTTAGTATTTTGGACAATAAATAAAAAATATCTGAGACTGGGAGGCGGATGCTCTCCATAGTTGGATTCTCGGTGTTGGGAATTGGAGATACTGTTTATTCCAATCAAATGCATTACAGACACACAAAGGGTTATTATTGGGTCTTATCATAGATAGTTGTTGAACCAGGTGCTAAGTATGCGAGGAGATGTACAAGCGGGCTGAAGAAGCAGGGAGCAAAGAGGGTATCAAGTTGAGCTCTATCCGTGCCTTTGTGCTGCTGGCGGCCGTGACCGACTATGGCAGCCGCGGTCGGAACCCGTATAGTCAGAGCCTGACTAGCCGTCCAGGCATCAGGCGGCCGCGGCCGGCTGGACGCGTATTGCCCAGTAACGCTTAGGCGTTGGGTGGTTTTAATTCGTTTCTTTTGGGAATATTTTTAAAATTATTGTACCCATTATATAAGAGAACGTAGGAACTTTATTAGGATTAAAAAATTTGATTAGATTGTAACCACTCTTTTCCCTCTTTCTAATCTTCAAGAAGAAGAAGAGATATCCACGAGGAATCTCTCGACATTGTAATCTTCATTGATTTCATTGAAGAAAGAGAAGCATCGTCTCGTCTCGTGGGCGTAGGTCAGAGAATAAGACCGAACCACAGT

UGT:

>F01_transcript/33044 full_length_coverage=2;length=2086;num_subreads=45

GCCTCATCAAGGCTTTTTCCATCCATCAAACACGATGACCAACCAAACATCCATCAATGGCTTCGGCCATCCGACCCCTCACTTCGTGTTGGTCCCTCTGATGGCCCAAGGCCACATGATCCCCATGGTCGACATGGCCCGCCTCCTCGCCGATCGCGGTGCCCTCGTCAGCTACATCACCACTCCCGTCAATGCCGCCCGAATCAAGCCCACCATCGAGAGCTGTCGCCTCCCCATTCGACTCGTCGAGCTCCCTTTCCCCACGGCCGAGGTCGGCTTGCCCGAGGGCTGCGAGAATGTCGATCTCATCCACTCCGGCGAACACATGATGCCCTTCTTCGAAGCCACCCGATTGCTCGCGGATCCCATGGAGCTCTATCTCCGCCAACAGCCTCAAGGCCCGACTTGCATCATCTCCGACTGGTGCAACTCGTGGACCGCCGAGGTCGCTCGGAAGCTCGGTGTGCCTCGCATCATCTTCCATGGCCCCTCTTCTTTCTACCTTCTGTGTGTTTATAATATCCGGAAGAATGGAATCTACGATCGGATCACCGATCCGTTCGAACCATTCGTGGTGCCCGACTTGCCACACGAGGGCTCGGTGGTGGTGACTACCGCACAAGCTCTTGGATTCTTGGACATACCGGGGTGGGAAAAGCTTCGGGATGAGGCATTGGAAGCAGAAGCGACGGCGGACGGGTTGGTGTTGAATTCTTTCGATGGATTGGAGCATTCATTCATCGAGAGCTACGCGAAGGCAATGGGTTCGAAGATCTGGGCGATAGGTCCCATGTCTTTACATAACAAGGATACCTCCTATAAGGCTACAAGAGGGAACAAGGCGGTGGTCGACCAGCATCGAGTTCTCACTTGGCTTGATTCTAAGAGCACTGGGTCGGTTATCTATGTTAGCTTCGGTTCTTTAGCACGAAATGACAAGTCGCAGATGATCGAGATCGGGCTAGGGTTGGAGGCATCGGATTGCCCCTTCATTTGGGTGGTCAAGGAAGTCGAGAAGTCTCCGGAGGTGGACAAATGGTTGTCGGGATTCGAGGAGAGGACGATGGAGAGGGGTCTTGTGATAAAGGGCTGGGCTCCGCAGGTGGTCATATTGTCGCACCCTGCGGTTGGAGGATTCGTGACGCACTGCGGGTGGAATTCGATGCTGGAGGCGGTGTGTGCCGGCGTGCCGGTGATGACTTGGCCGCACTTTGCCGACCAATTTCTCAATGAGAAGTTGATGGTGGAGGTTTTGAGGATCGGAGTTTCTATCGGGGCGAAGATTCCCAGTTACCCTCTACAATCGAAGGAGGGTGGGGAAATGCGGCTGGTGCAGAGGGAGGATGTAGAGAAGGCAGTGGTGAGTTTGATGGAGGAAGGGGAGGAAGGGAAGGAGAGGAGGGAGAGGGCTAAGGTGTTTGGTGTGAAGGCCAAGGAGGCAATGGAAGAAGGGGGCTCATCTTACAGGAGTTTGACACACTTGATCCAATATGTCATGGAGCGTTAAAACAAGTTCAAATTTTAAATATTTTGATTTTTATTTTGAACTTTTAGCTCCTGTTCTCTTCGGAGAAATGAGGCGGAAAAAAAAAATTTTCTGCTTATTTTTTTTTGTTCTTTTCGTTCTTATCACCCGGAAATTTTTTTAACTGTTTCCTAAATTTTAGGAAGACCCGGAAATTCATCATTTCCATTTCCGTTTCAAACAAGATGGAAATTGCTGATTTCCGGACGAAAATCTGATGTGGCGGACTGCCTTCCACATGATAGGGAAGATCAGAAAAAAAAAAAAACTGTTAGTGTGGGCGCCATATTTTTCTCACTTCCTATTTTTAAGCGATTTTTTTGTTTCCCCATCTCATTTCCGCAGATTTTTTCCGCCTCCCATTTCCCCTTATTTCTCAATTCTGTGAAGAGAACAGCACCTTATTGTAGTGAGGGGAGAGAGACAGGGAGATGGAGGGATTCCAAATTTCTATCCAATTTTTTAGTGTCTTGATTTTTATTTTTTAGCTTTTATTTTCATTCTTAGTTTATGTTGTAGTGAATATTTAAAATATGGATTCTAATATGTGTAAATTTTTGC

>F01_transcript/38535 full_length_coverage=14;length=1897;num_subreads=60

GAAAGAGCAATGCTCCGTAGAGAGATTTTTTCATCATCTTAATACCTGTTTTTATCATGGTATCAAAGCCCTAAGACTTGCTGAATTGAAATCTCATTTTGTTTGAGAGAGAGAGAGAGAGAGAGAGAATTCCTCACCAGTCAAATACAGAAGATAGGATTGAAGGCAGTAGAGTAGCAGCAGCCCAACCTCCCAAGCATCATGTGCTCTCCACCGCCCCACGCCCTCGTTCTTCCCTGCCCGGGGCAGGGCCACGTCATCCCCTTCCTCGAGCTCGCCCACTGCTTGGTCGACTGTGGCTTCGAGATCACATTCGTCAACACCGAATTCAACCACCACCGCATTCTAGCCGCCCTTTCCGACCCTCACTGTGAAAACAGTGACCTAATACATTTCGTCTCGATCCCCGATGGCCTGGCTCCCGACGACGACCGTAACGACCTCAAAAAGATAACAAACAGCATCTTGACAGTCATGCCAGGCTTTCTTGAGGACTTAATCCGATCCACCAACGAGTTCCGCGAGGATGGCAACAAAATCACGTGTATAATCGTAGATTACTTTATGGCATGGGCCCTCGCTATCGCCCGGAAGATTGGCCTCCGGTCTACCGTCTTCTTTCCGGCTGCAGCTGGAATGTTCCTAACGCTGGCGAGAATCTCGAAGCTGACCGATGACGGAATTATCGATGCTGCTGACGGATCCGTGAAGAGACACGAGATGATCCAGCTGACACCTGGAATGCCCCCGAGTTGCCCGAACCGCCTCCCATGGAATTGTATTGGCGATTCGGAGACTCGAAAAACGATGTTCCACTCCTTTGCCTCTTATAATCGTGCGACCGAGACGGCGGATTTCATTGCCTGCAACTCGTTCCTGGAAATTGAGGAACCGGTTTTCAGCTACGCCCCGAAATTGCTCCCCATTGGGCCTTTGCAGACCGGACTTCGGACTGGAAAACCGGTCGGGCATTTTCGGCGGGAGGATTCGAGCACGGTGGCGTGGCTCGACGAGCAGCCATCTGGTTCGGTGGTTTATGCGGCATTCGGAAGCTTCACTATTTTCGACCGACGGCAATTCCAAGAGCTCGCCCTCGGGCTCGAGCACACGGGCCGTCGGTTCCTATGGGTCGTCAGGTCCGACCTAACGGACGGACCGAGCGACCCATATCCGGACGGGTTCGGGCTAAGGGTGGCGAACCGTGGGAAGATGGTGGGCTGGGCCCCACAACACAGGGTGTTGGCCCACCCATCCATTGCTTGCTTTGTGAGCCACTGTGGTTGGAACTCAACTATGGAAGGGGTGAGGAATGGGCTGCGGTTCCTCTGCTGGCCTTACTTTGCTGACCAATTCTTCAACGAGAGTTACATTTGTGATGTGTGGAGGACTGGGTTGAGGGATGAAGGAAGATGAGAGGGGTATTGTCACCAAGGAACGGATCATATCAAAGGTGGATGAATTGATGGGCGACGACGAGATGAAAGCGAGAGCAATGGTTTTAAAGGTATTGGCCGATCAAAATGTCAACGAAGGCGGAGTTTCTTTCAAAAATTTGAATAGCTTTGCGGATGCCATGAAATGATGAGAGAGTTCTCTCTCTCTCATTTTCTTTCTATTTTCTGCAAAGGGAGAGCGGTAGTTTTCTCTGGTCATTATTAGTGCTTGAATCCACCCACAGAAGTATCACTGATGTTGGAGCTTGATCTCATCTGAAATTGATCTCATCCCGACCAACTTCAACCCAATAAATTTCTCCCAACATATTTCATTTAAAATTTTATTTGCTGCTTTAGTTTGTTAGTTCGTTTTAATTTATTTTTCCTCTTTTCTGTAAATTGAGATGTTATGTATTTTTAATGTATTGTTAATTTTATGAACGAAATTTCACTTTTTCATG

>F01_transcript/39103 full_length_coverage=8;length=1897;num_subreads=60

GAGTTTCAAGTAGAGAGAGAGAGAGAGAGAGAGAGAGAGAGAAAACAAACACTGATATCACCCACATCTCAATGGAGTCCCAACCAGAGCTCCTCCACCTAGTTTTCTTCCCCTTCCTCGCTCGCAGCCACATGATCCCCATGCTGGAGACCGCCCGCCTCGCCGTCGAGCGCGGCGTCAAAACCACCCTCGTCACCACCCCTGCCAACGCCCACCTCATCCACCCCGTCCTCCACCGCTCCAACTCATCTCTCCTCCCCTCCCATCCGCCGATGCAGCTCCAACTCATTCCCTTCCCCTCGGTCGAGTTCGGCATCCCCGAGGGGTGCGAGAACCTCACCTCTATCCCCCTCCCCCTCGTCGCGGCCTTCTTCAAAGCCATCTTCGCCCTGCGGGCCCCGCTCGGCGCGCTGTTGCGGGAGCTCGGCGCCCACGCCCTCGTCGCCGACGCGCTGTTCCCGTGGGCCACGGGGCTGGCGGCCGAGATGGGGATCCCGAGGCTTATCTTCCAGGTCATGGGTCTGTTCCCGCTCTGCGGTGCCCACGATCTCGATACCCATCGGCCGCACGAGGCCGTTGGTGGGGATGACGAGGAGTTTACCATCCCGGGGTTTCCGGACCCGGTGAAGCTTACCAGGGGACAGGTCCCCGAGGTCTTCAGGCACGATTTCATGCTGGCCCTCCTCCGCGACGCAGAGTTCACCAGCTACGGCGTGATAGTGAACAGCTTCTACGCCCTCGAGCCCAGTTACGCGGAGCACTACTACAAGGTGGCCCCCCGGAAGGTCTTCCTCCTCGGCCCGGTCGCCCTCGCCGGCTCCAATCCCTCGCCACTGCCATTGGAGAGTGGCGACCCCTGCATCACCTGGCTCGACTCCAAGCCCGACAACTCGGTCCTTTACCTGAGCTTCGGCACCACCTGCCGATTCAGCGACGAGCAGCTCGTCGAGCTCGCCGAGGGCCTCGCGTCCTCCGGCCACAACTTTGTGTGGGTCGTGGCCCTCCCCGAGAGCAGCGGCTCGACCAGCGAGGAGTGGCTTCCGGAAGGCTACGAGCGTAATGTGGCGGGTCGGGGACTTCTCGTAAATGGTTGGGCTCCGCAGACCGCGATCCTGAACCACCGGGCAGTGGGTGGGTTCGTGAGCCAATGCGGGTGGAACGCCGTCATGGAGGCGGTGGCGGCAGAGGTACCCATGATCACGTGGCCGCTTCATTCGGACCATTTCATCACCGAGAAGCTGTTCTGCGATGTGCTGCACGTGGCGGTGCCAATGTGGGAGGGGCGGAAAAGCATCTGGGATGATCAGAAGGAGGTGGTGCGGGCGAAAACAGTGGCAGCGTCGGTGAAGTGGCTTATGGGGGGCGGGGACGAGATCGAGGCGATGCGAAGGAGAGTGAGGGAGCTCGGGGAATTAGGACGGGCCGCGGTGGCAGAAGGCGGATCGTCCCACTCCGACATGGGCCGTCTTATCGACGTGCTCACGGAGGAGCGAAACAAGGCTAAGGAAATGATAGACAAGAACAGTGTTAGTTATGATTGTTCTGGTCGCAATTGATTATTTCTCGTCAAGAACTGATAGGAGTGTTTTTTGATAAACATGTTATGTTGGCGATCTCTAGTCACCACTTTTGAGACAGGAAGAATGAACTTGAGTTTTGGAGGGATTGAGACCATCATCTCTATTTGGAGTAATTTCTTCTGATTTTGTATTTGTGCCATCATCAATAAATTGGAAGACTTTGTCGTCGTGAATGTAGCCTGAGTGAAGAGGTGAACCACATATCTCTTTGTGTTATGTTATGTTTGCTTATTTGTTTTATTTTCTTGGTGAATGAGTTTTTAGATTATTTTGTTCCGTATTGATAATTGAATGGAGGAGTAATTATCCGTCGATCT

>F01_transcript/39110 full_length_coverage=65;length=1785;num_subreads=60

GACTCAGTGCCTGCGCCTTCCGTTAAAAGCCTCTCCTTCTCCGTTGTTTTCTCAGCACAAACCCATACGAATAATCTCTCAGACTGCTCTCTCTCTCTACCCTTCACTCTCTTTCTTCTCCGATCCCCCTTTTCTCTCTCTACATCTCTCGCCATGGGCTCCGACGCGAAGAATCCCCTCCACATCGTCTTCTTCCCCTTGATGAGCCCGGGCCACATGATCCCCATGGTGGACATGGCCCGCCTCTTCGCCCGCCGCGGCGTTAAGTCCACCCTCGTCACTACCCCCGCCAACGCCGCCAACATCCGCGCTGCAGTCGACCTCGACTCCGCCGCCGGCCGCCCCATCGCCCTCCACGTCATCCCCTTCCCCCCGCCGGAATCCACCAAGCTCGCCGCCGGTCAGGAGAACCTCGCCGCCGTCTCCACCGCCCAGTTCACCGATTTCATCAACGCCCTCTTCCTCTTCCGCCCCTCGATCGAATCCCTCCTCCTCGACCTCCATCCCGACTGCCTCGTCTCCGACTCGTTGTTCCCGTGGACAGCCGACCTCGGCACCCGGCGGATCATCTTCCATGGCGCCGGGGCATTCCCGATGTACGTGCTTGGCAAGATTTTTCAGTATTTTCCCCTCGAAAAAGAGGAGTTCACGATGGAGGGGCAGCCGGACGAGATCAAGCTCTATAAGAAAGGGTTGCCGGAGATCGTGGATAATCTGATGATGCTGCAGCTTTTGGGGGTGGCCGAGGCCAAGAGCTACGGCGTGGTGGTCAACACCTATAGAGAGATGGAGCCGACCTACGTGGACTACTACAAGGGGACGAAGCGGGCATGGTGCGTGGGTCCCGTATCTCTACTAGCCAACGACGGGTCGACGGATGTCGAGAGGGGGGAGGCGTCGACGGACGGTGAGCGAATCATGGGGTGGCTTGGGGGGCAGGCGGCGGCCTCGGTGGTTTACATGTGCTTCGGAAGCTTGTGCCACTTCAGCGGGGCGCAGCTGCGGGAGATCGCGGTTGGGCTCGAGGCGTCGGGGTTGCCGTTTCTTTGGGTGGTGAGGAAGGAGGCGGACGCGGACGAGATTGTGGAGAAGGATTGGATGCCGGAGGGATTCGAGCAGCGTGTGGAAGGTCGGGGGTTGGTGGTGCGGGGATGGGCCCCACAGACGGCGGTGCTGGCCCACGGGTCGGTGGGGTGGTTCGTGACCCACTGCGGGTGGAACTCCCTGCAGGAGGCGGCGGTGGCCGGAGTGCCGATGGTGACGTGGCCGCTGTTCCATGAGCAGTTCCTCAACCAGGAGCTGGTCGTCGAGGTCATGGGCGTCGGAGTGAGAATGTGGGAGGGCTTCAGGCGCAACAGTCTCGGTGACGACAAGGCCCTCGTCACCTCCGACGAGATCTCTGCCGTTCTGTCAAAGCTCGCTGCCGGCGGGGAGGAGGCGGAGGCGATGCGCCGGAAGGCGGGAGAGTACTCTGTCGCCGCCAAGAAGGCGGTTGCGGAGGGCGGCTCCACCCACACAGACATCACCAACCTCATCGAGGAGCTCGAGGGGCTTCGGAAGGAAAAGTAGACCGGGCCGGACTTAAATGGTCGGACCAGTGAAAAACCGACCGATTATCTCTCGGGTTTTCCTTCTCGAAGGGAAGACTTTATCACCTCTTGTACTTTTTTTTTTCTTTTCGCTAGAGTGAGTTGTTTCTGGATGGAATGAGTGCAGTTGCAATTGCAAAGGTGGTTGTTTGATTAAGAGTGATTTTGATGATGATTTTCATTATTTTTTACGCGT

>F01_transcript/40869 full_length_coverage=10;length=1839;num_subreads=60

GAGACACCAGAGCTCTCTCTCTCTCTCTCTCTCTCTCTCTCTCTCTCTCTCTCTCTCTCTCTCTCTCTCTCTCTCTCTCTCTCTCTCTCTCGTCATGGATCGAGAAAAGCTACACATAGTTGTGTTCCCGTGGCTAGCCTTTGGGCACATGTTGCCTTTTCTTCAGCTCTCCAAATCTTTGGCCACAAGGGGTCACCGAATCTCCTTTATCTCCACCACAAGAAACATTAAAAAGCTCCAGCCCAAAGTGCCACCCACGCTCTCCCAACTCATTCAATTCATCCCTCTACCTCTCCCCTCGGTCGACTGTTTACCGGCTACCGCCGAGTCTGTCTACGACATCCCTCTAAACCTCAGCCTTCACCTCCAGAAGGCGTTCGACGGCCTCGAGCACCCATTGGCCCAATTCCTCACTTCATCGTCTCCAAAGCCTGATTGGATCATCGTGGACTACGCCTCCGATTGGCTCCCGCCACTCGCATCAAAATTCAATGTGCCGTGTGTCCACTTCTGCACCAACATGGCCTCAACCATGGCGTTCATAGGACCCGTGTCAGAAGTGGACAACATAGACTCCATCACCGTTGAAAATGTGATCGTTCCGCGCAAATGGATGACATTCCCTACAAATGTAGCCCTTCACCCTTATGAGGCTCGACAGGTTCTTGATTTGTCCAGTTCAGATAATGCTTGGGATGGCTACCGCTACTTGCTCACCATCAGAGGTTGCAAGGCCGTAGCCATCCGAAGTTGCAACGAGTTCACGCCCAAGTGGTTGCCCCTCCTTCGAGATATCTACAAGAAACCCGTCATTCAGGTTGGCATGCTTTCCTCGATGCTGGTCGAAAATGGAGATTATGACAACAACGAGCTCAATGTGATAAAGTGGCTGGATCGACGATCTCCGAGTTCTGTCGTGTATGTAGCCTTTGGTAGCCAAGCCGAACTCAGCATCGAGCTCTTACACGAGCTAGCGCTTGGGCTTGAGCTCTCTAACTTGCCTTTTCTTTGGGCTTTTAGAAAGCCGACCACCAACGAAAACGAAGAGATTTTGCCCATAGGATTTCAAGAACGGACAAAGGACCGAGGACTCGTCGTCATGGGTTGGGTTCCCCAGCTCAGGGTCTTGGCCCATGAGTCGGTGGGTGGATTCATGACGCACTGTGGTTTTAGTTCGATCATCGAGAGCCTCCATTTTGGTCGTCCCCTAATCCTCTTCCCTTTATTCGGGGATCAAGGGCTCAATGCTCGGACGATGGTGGAGGAGAAGATCGGATTCGAGGTGAAGCGGGATGAGGAGGATGGGTCACTAACAAAGGAAGCGGTGGCCGAGGCATTGAGGTTGGTTATCGTCGAGACAGAGGGGGAACCTTTCAGGAAAAAAGCCAAAGAGATGATGGGGGTTTTCACGGACAAGAAATTTAAATGAGAGATATCTGGATGATTTCACTCAGTATCTAATGGACCATAAAGATAGCGAATAAAAACAGCTCAACCGAGGAGTCAGGCAAGAGGAAGGGTAGTGCAAACCAGTGCCACTGCTAAGGATATCTCCTTATATTTTTAGGATGATGATAACCTATATCTAGTGTGTGTTGTATGTGTTTGGGACACCAGCTTATAATATTAAAAAATGTGATCTAATTTAGAAAAAATAAAGCTATAAGATGTTGTAATAAGAGGAAACATGTAATATTATGTTTTAGAATAAATATGAATGGCTTGGACTGGCTCAAGGCTCCCTTCGCTCGTGGAAGAAGAAAATTTCTCAAAAGAAAATTGGTTGTAGTTTTTGCTTGGTAATAATGAGTTTTTACTTCACTTCAATCTTATCCTGAC

>F01_transcript/41117 full_length_coverage=6;length=1817;num_subreads=60

GACACAAGAGCTACCTCTCTCTCTCACTGTCTCTCACTCTCTACTCTCTCTCTCTCGTCATGGATGGAGAAAAGCTACACATAGTTGTGTTCCCATGGCTAGCCTTCGGGCACATGTTGCCCTATCTTGAGCTCTCCAAATCTTTGGCCGCAAGGGGCCACCAAATCTCCTTCATCTCTACCACAAGAAACATAGAAAAGCTCCAGCCCAAAGTGCCACCCACGCTCTCCCAACTCATTCAATTCATCCCTCTACCTCTCCCCCCGGTCGACGGCTTACCGGCTGCCGCCGAGTCCGTCTACGACATCCCTCTAAACCTCAGCCGTCACCTGGAGAAGGCGTTTGACGGCCTCGAGCGCCCCTTGGCTCAATTCCTCAGTTCGTCGTCCCCAAAGCCTGATTGGATCATCGTGGACTATGCCTCCGACTGGCTCCCGCCGCTCGCTTCCAAATTCAATGTGCCATGTGTCCACTTTTGCATCTTGATGGCCCCAGCCATGGCGTTCATGGGACCCGTGTCAGAAGTGGACAACATAGACTCCATCACGCTTGAAAAAGTGGTGGTTCCGCGCGAATGGATGACATTTCCTACAAATGTTGCCTTCCGCCCTTATGAGGCTCGACAGGTCCTTGATTTCTTAAGTTCAGACAATGCTTTGGATGCCTACCGCAACTTGCTCACCATCGGAGGTTGCAAGGCCGTGGCTCTCCGAGGCTGCAACGAGTTCATGCCGAAGTGGGTGCCCCTCATTCGAGATCTCTACAAGAAACCCGTCATCCAAGTTGGCATGCTTTCCCCGATGCCTGTCGAAAATGGAGATTATGATACCAGCGAACTCAAGATCATGGAGTGGTTGGATCGACGATCGCCGAGTTCTGTCATATATGTAGCCTTTGGGAGCCAAGCCGACCTCAGCATCGAGCTCTTACACGAGCTAGCGCTTGGGCTTGAGCTCTCTAACTTATCTTTTCTTTGGGCTCTTAGAAAGTCGACCACCAGCGAAAATGAAGTGATTTTGCCCGTAGGGTTTCAAGAACGGACAAAGGACCGAGGATTTGTCATCATGGGTTGGGTTTCCCAGCTCAAGGTCTTGGCCCACGAGTCGGTGGGAGGATTCATGACGCACTGCGGTTTCAGTTCGATCATCGAGAGCCTCCAATTTGGTCGTCCTCTAATCCTTTTCCCTTTGTTCGGGGATCAAGGGCTCAATGCTCGGGCGATGGTGGAGGAAAAGATCGGATTTGAGGTGAGGCGGGGAGAAGAGGATGGGTCACTAACAAAGGAAGCAGTGGCCGAGGCATTGAGGCTGGTTGTCGTCGAGGCTGAGGGGAAACCTTTCAGGAAAAAAGCCAAAGAGATGATGGAGGTTTTCGCGGACAAGAATTTAAATGAGAGATATATGGATGATTTCACTCAGTATCTAATGGACCATAAAGATAGCAAATGAAAGTAGCTCGGTTGAGACGCCAGGCAAGAGGAAGGGTGGTCCAAAGCAGTGCCATTGCTAAGGAGATCTCCTTCTATTTCTGGGATGAAGAGAATCTATATCTAGTGTGTGCTGTATGTGTTAGGGACACCAGCTTATAATATCTAAAAATGTGATGTAATTTAAGAAAAATAAAGCTGTAAGATGATATAATAAGAGGAACAATGTAATATTGCTTTTAGAATAAATATGAATGGCATGGACCGGTTCAAAGCTCCATTCACTTGTGGAAGAAGAAAAGTTCTCAAAAAAAAATGTTGGTTGTAATTTTTGTTTGCCTAATAATCATTTTTGTTTTGCTTCAATTCTATCCTGATCCGATCTTCCTGC

>F01_transcript/41118 full_length_coverage=2;length=1819;num_subreads=60

GGAACTGTGCTTTACTTCTGCGGGATCTGGATCGCCAGTTCTCCAATTTTGATCGGAGCAATCATTGCTCTCTCTAGCTCTGCAGCCTGAAACCAAGAGACTCCCTTCCGATGGATCTTCACGTCTTCTTCTTGCCCTTTCTTGCTTCGGGACACATGATCCCCATGGTGGATATCGCCCGCCTCCTCGCTGACCGCGGCGTCAAATCCACCATCGTCACCACCGCCGGGAACATCCCCCGAATCCAGCCCACCATCCAACGCTTCAACTCCATTTGTTCCTCCGACAAACCTCCAATTGAACTCCTCACCATCCCCTTCCCCTCATCGGTCCCTGACGACTTCTCCCTCCTCCCGACACTCGACTTCACCCCCGAATTCGCCGCCGCCATCAGTGGTCTCCGCCAACCCTTTGCCGAGCTCCTCGAGTCCCACCGCCCTGATTGCATCATCTCGGACTTCCTCTATGCGTGGACCAACGACCTCGGATACCCGAGGATAGTCTTCCACGCCGAGGGCTTCTTCTCCTGCGTCGCTCCCGGGGCCGTGGCATCTCAAAAGCTACACGAATCCGTTACCAGCGACGATGAACCCTTCGTCGTGACCGGCCTCCCGCACAGGATCGAGATGACTCGTTCACAGCTCCCGAACTTCTTCTTCTTCAAGTCGGATGATGACGTCGGGTTGAGAACGGGGGTCGTCGAGTCACTCTGCAACAGCTACGGAGCTGTGATGAACAGCTTCTACGAGCTGGAGCCCGAGTACGCCGACATTATGAAGAAGAACGCCACATTCAAGATCTGGCACATCGGACCTGTGTCGCTGTCCCGCAGGAACACTGCCGGTGAATCGGTCGAATCCGGCATAATTCGTAGCTGGCTCGACGGCAAGAACCCGAGCTCGGTTCTTTACGTCTGCTTCGGGAGCTTGGTGAAGTTCACGACACCCCAGTTCCGAGAGATCGCGTCTGGCCTCGAGGATTCCGGCCACCCGTTTATATGGGTGGTGAAGAAGAACGACGGAGATATTCTACCCGAGGGATTCGAGGAGAGGGTGAAGGGAAAAGGACTGCTCATAAAAGGTTGGGCGCCGCAAGTGATGATACTGAACCACCCGGCTATAGGTGGGTTCATGACGCACTGCGGTTGGAACTCCTGTTTGGAGGGGGTGAGCGCGGGGTTGCCGATGATCACATGGCCGATGTATACCGAGCAGTTCTTCAATGAAAAGCTGATCGTCAATGTGTTAGGGGTCGGAATTGCGATTGGGGCGAAGATTTGCAGTCTACATGAGGAAGAGAGGACGGCTGTGAAGGGAGAGGTTGTAGCGAAGGCGGTGAGGGAATTGTTGGGCAGTGGAGAGGAGGCCGAGGGGAGGAGGAGGAGAGCCAGGAAGGTCGGAGAGAATGCAAAGAGAGCAGTGGAGGAGGGCGGGTCGTCGTACAGTGAAATGGACCATCTTGTAGAGGATCTGATCAATTTGAAAACCGGTCGGTTCGGGTCCAATTCCATCACCCCAAATAAGTGATTAAATTTTGTGTGGATTTACTATCCTCTCTTTCAATAATTTTAGGTGTAGTTTGGATGCCCAATAAATTAAGAAAAAAAAAAATTGGATGGATCCTTGATGTACACCCGTGATAAGTTTTACGGATACTTATCCTTCCAATTTGTGAAAAAAAAAATGCTACTATTGCTCACAATTTTTATTTCCATTTGAGTACTCCATTATGTGGTCACTCAAAAGACTCACTCTAGATGAGAGAGGATCCAAAAAGATGATGTGACATGTATTTGACATGATGGATAACAATGATTATC

>F01_transcript/42184 full_length_coverage=4;length=1716;num_subreads=35

GACACCCGAGCTCACAGAAGTCTCTCTCTCTCTCTCTCTCTCTCTCTCTCTCTCTCTCTCTCTCTCTCTCTCTCTCGCATCATGGATGGAGGAAGCATGCACATAGTTGCGTTCCCAGCCCTTGCCTATGGGCACATGATGCCCTATCTTGAGCTCTCCAAATCTTTGGCCACAAGGGGCCACCAAATCTCCTTCATCTCCACCACAACAAACATAAAAAAGCTCCAGCCAAAAGTGCCACCCGCACTCTCCCAACTCATTCAATTCATCCCTCTAACTCTCCCGCCGGTCGACGGCTTACCGGCCACCGTCGAGTCCTTCACGGACATCCCCCAAAGCCTCGCCCATTACTTGAAGAAGGCGTTGAAAGGCCTCGACATCCCCTTCGCCCAATTCCTCAGTTCATCGTCGCCAAAGCCTGATTGGATCATCGTAGACTACATCTACAACTGGGTCCCGCGGCTCGCTTCCGAATTAAATGTGCCGTGTGTCCAATACAGCGTCTTTTTTCCCTCGACCTTGGCATTCATGGGACCCGTGTCAGAAGTGGACAACATAGACTCCCTCACGGTTGAAAAATTGATCGTTCCGCGCGAATGGATGACATTCCCTACAAATGTAGCCTTTCGCCCTTATGAGGCTCGAGAGATCCTTGATAGCTTCAGTTCAGACAATGCTTGGGATGGCCAAAGCAACTTGCTCACCATCGAAGGTTGCAAGGCCGTGGCCATCCGAAGCTGCAATGAGTTCATGCCCAAGTGGTTACCCCTCATTCGAGATCTCTACAAGAAACCCGTCATCCAAGTTGGCATGCCTCTCCCCTCGCTGGTCGAAAATAGAGATTATGACACTAGCGAACTCAAGGTCATGGAGTGGCTAGATCGATGTTCTCCGAGTTCTGTCGTATATGTGGCCTTTGGGAGCGAAGCCGAAGTCAGCATCGAGCTCTTACATGAGCTAGCACTTGGGCTCGAGCTCTCTAACTTGCCTTTCCTTTGGGCTTTTAGAAAGTCGACCACCTACGAAAATAAAGAGATTTTTCCAGAAGGGTTCCAAGAACGAACAAAGGACAGAGGATTCGTCGTCATGGGTTGGGTTCCCCAGCTCAAGGTCTTGGCCCACGAGTCGGTGGGAGGATTTTTTACGCATTGCGGTAGTAGTTCGATCATCGAGAGCCTCCATTTTGGTCATCCCCTAATCCTCTTTCCTATATTCGGGGATCAACCACTCAGTGCTCGAGCGATACAGGAGGAGATGATCGGATTCGAGGTGAAGCGGGGTGAGGAGGATGGGTCACTAACAAAGGAAGCGGTAGCCGAGGCATTGAGATTGATTGTCATCGAGGCAGAGGGGGATCCTTTTAGGCAAAAAGCGAAAGAGATAATGGAGGTTTTCACGAGAAAAAATTTCAATGAGAAATATATAGATGATTTCACTCAGTATCTAATGGAGCATAAGGATAACAAATGAAAGTACCTCGTTCAAGGATCTAGACAAGAGGAAGGGCAGTGCATAACAGGGCCACAACTAAGGGTGTCTTCTATTTCTATGATGAAGGGAACATATATGTAGTGTGTTGCAGATGTTGGAATACCAACTTATAATATCCAAAAATATAATGTAATTTAGGGAAAATAAAGCTGTAAGATAATGCAATAAGACGGAACATGTAATATGATTTTTATATTTAATAAATTTGAATGTTCTGCAACGGCT

>F01_transcript/42313 full_length_coverage=3;length=1782;num_subreads=60

GCCGTGCAAACTTGGTAGCGTAGGCAACCTCAGCTTCACGCCCAATGTTCTTCATCCTCTATATATAAATCCCACCCCAACCATAAACAAATGGGAACAAAGCAAAGCAAAGAAAAGATGTCCGCCGGCAAGATCATCGTCGTCCCTTTCCCCCACCCGGGCCACATATTTCCCGCCACTGAGCTCTGTCGCCACCTCGCCGCCCGTAATTACAGCATCACCCTCTTCCTCCCCTCCACCTCATCCTCATCCTCATCCTCATCGTCCCTCCACCCTCTCATCACTACCGTCGAATTCCCCGTCCCCCACCCCACGAACTTCCCGCCCCATCTTCTAGAAGACACCCTCCCCGATCTCCTCGCCCGCCATATCGACCCCCCGCCCGTCTGCGCCATTGTTGACGAGATGATCGGCTGGCTAATCGACGCCTGCGCATCACTCAACATCCCCGCCGTCAGCTTCTTCACCTCCAGCGCCTGCTCCTCCGCCATCCAGCACGCCACGTCATCCATCAACCCCTCCGACCTGCTCCCCGGGCAGCCCCTCCTCCTGCCCGGCCTGCCCGACGACCTCTCCCTCACCGCCTCCGACCTCCCCCCGCCGGATCATGGTCCCGGTCGCCGCGGCGGTGGTCCCCACCGCGGACCGAGGGTGAGGGGGATGGCGTCGCTCGCCGGTGCGAAGGCGATCCTGGTGAACACGTGCGACGATCTGGAGCGTCCGTTCGTGGAGTACATGGCGCGGGTGGCCGGGAAGCCGGTGTGGGGCGTGGGCCCGCTGCTCCCGGCCCAGTTCTGGTCCTCCACGGCGTCGCACGTCCGGGACAGGGAGGTCCGGTCGCGGCGGGAATCGAGCGTCAGCGAAGGTGACGTCATCCGGTGGCTGGACTCCCGTCCCCGGGGGTCTGTGGTCTACGTGTCGTTCGGCAGCCTAGTGGTCCCCTCGGAAGAGGAGCTGGGCCAGCTGGCGGACGGACTGAGGGAGTCCGGGCGGCCTTTCATCTGGGCCGTACAGCCCGACGCCAGGAGGCCCGGCCCGGACGGCCCGATCGCCGCCGGAGTCTCGGCCGCTGAGCTGGCGCGCCGAACCGGATCAATGGGACTGGTGGTGGAGGGGTGGGCCCCGCAGCTGGCAATACTGGGCCACCCGTCGACGGGGGGGTTCGTGTGCCACTGCGGGTGGAACTCAACGGTGGAGGCGCTGGCGTTCGGAATGCCGATGCTGACGTGGCCGGTGAGAGGGGACCAGTACCACAACGCCAAGCTGGTGACGAACCGGCTCGGGACCGGTTGGGCCGTGAGGGGAGCCGGACCGGTCGTGGGAAAGGGAGACGTGGCGCGGGGGATCGAGAAGGTGATGGGGGACGAGGAAATCAAGCGCCGGGCGGCGGCGGTGGGGGAGATGCTTTCCGCCGGATTCCCTCCGAGCTCGTCGGATTCGGTGGACGAATTTCTCGATTTCGTCAGCAATGATGCTCGAAAGAATTGACAAGTCCAGTCCTTTACTGTGTCACTTTCTTTATACCATAATAATTCGGGCACTCTTATTCTGTGTCGGCAATCGATCCAGCTTATCCACAAAAAACATGCGGTTCGTCGGGATTCTGAGTAGTCGTCCGTCTCTTTACTTTTATTCCTTCTTGTGTGGAAAGAGCATCTTTCATGGTTCCCTTGGCTAGGGATCGTCCAATAATACGATTCTTCATATATGTATCAGAGTTCACAATCTTCAGTGGTTTTTTTTTTTTGGCGAACAATAAGCTTCAGTATTCATTTGTTTCCT

>F01_transcript/42413 full_length_coverage=5;length=1792;num_subreads=60

ATCAACGCAGAGTACATGGGGACACCAGAAGAGCTCTCTCTCTCTCTCTCTCTCTCTCTCTCTCTCTCTCTCTCTCTCTCTCGTCATGGATGGAGGAAACCTGCACATAGTTGCGTTCCCTGTGCTAGCCTATGGGCACATGATGCCCTATCTTGAGCTCTCCAAATTGTTGGCCACAAGGGGCCACCAAATCTCCTTCATCTCCACCACAAAAAACATAAAAAAGCTCCAGCCCAAAGTGCCACCCACACTCTCCCAACTCATTCAATTCATCCCTCTAACTCTCCCGCCAGTCGACGGCTTACCGCCTACTGCCGAGTCCTACACGGACATCCCCCAAAGCCTCGTCCAATACCTCAAGAAGGCATTCGACGGCCTCGAGGGCCCCTTCGCCCAATTCCTCAGTTCATCGTCCCCAAAGCCTGATTGGATCATCGTAGACTACCTCTACCCCTGGGTCCCTCGGGTCGCTTCCGAATTCAATGTGCCGTGTGTCCACTTCAGCATCGTAGCGGCCACGGCCCTGGCATTCTTCGGACCCGTGTCAGAAGTGAACAACATGGACTCCCTCACAGTTGAAAAATTAATCGTTAAGCGCGAATGGGTGACATTCCCTTCAAATGTGGCCTTACGTACTTATGAGGCCCGAGGGGTCCTTGATTGCTTCAATTCAGACAATGCTTGGGATGCCCACAGCGAATTGCTCACTATCGGAGGTTGCAAGGCCGTGGCCATCCGAAGCTGCAACGAGTTCGTGCCCAAGTGGTTATCCCTCATTCAGGATCTCTACAAGAAACCCGTCATCCAGGTTGGCATGCTTTTTCCAACGCTGGTCGAAAATGGCGATTATGACACCGGTGAACTCAAGATCATGGAGTGGCTGGACCGACGATCTCCAGGTTCTGTCGTATATGTAGCCTTTGGGAGTGAAGCCGAACTCAACATTGAGATCTTACATGAGCTAGCGCTTGGGATCGAGCTCTCTAACTTGCCTTTCCTTTGGGCTTTTAGAAAATCAACCACCTATGAAAATGAAGATATTTTGCCCAAAGGGTTCCTAGAACGAACAAAGGATCGAGGATTCGTCGTCATGGGTTGGGTTCCCCAGCCTAAGATTTTGGCCCACAAGTCAGTGGGAGGATTCCTGACGCACTGCGGATATAGTTCCGTCATCGAGGGTCTCAATTTTGGTCACCCCCTAATTCTCTTCCCTATATTCGGGGATCAACCACTTAATGCTCGAATGATGCACGAGGAGATGATCGGATTCGAGGTGAAGAGGGATGAGGAGGATGGGTCACTAACAAAGGAAGCGGTAGCTGAGGCATTGAGGTTGGTTGTCGTCGAGGCAGAGGGGGATCCTTTCAGGCAAAAAGCCAAAGAGATGATGGAGGCTTTCACAAGAAAGAGTTGCAATGAGAAATATATAGATGATTTTATTCAGTATCTAATGGGCCATAAGGATAGCAAATGAAAGTAGCTTACCCGAGGCAGAAGACAAGAGGAAGGACAGTGCATAGTAGGGCACAACTAAATGCATCTCCTTCATTTCTGGGATAAAGGGAACCTATATGTATTGTGTGTTGTAGGTGTTGGAATACCAACTTATAATATCAAAAAATATAATGTAATTTAGGTGAAATAAAGTTGTAAGATGGAATAATAAGACGAAACATGTAAAATGTCTTTTATATTGATTAAATATGAATGTCCTGCAACTGCTCAAGTCATCCTTCCCTCATGAAAGACAAAATTTCTAAATAAAAAGCTAGTTGTAGTTTATGTTTGGTAC

>F01_transcript/43033 full_length_coverage=13;length=1769;num_subreads=60

GAGTTTCAAGTAGAGAGAGAGAAGTAGAGAGAGAGAAAACAAACACTGATATCACCCACGTCTCAATGGAGTCCCAACCAGAGCTCCTCCACCTAGTTTTCTTCCCCTTCCTCGCTCGCAGCCACATGATCCCCATGCTGGAGACCGCCCGCCTCGCCGTCGAGCGCGGTGTCAAAACCACCCTCGTCACCACCCCTGCCAACGCCCACCTCATCCGCCCCGCCCTCCACCGCTCCAACTCCTCTCTCCTCCCCTCCCATCCCCCGATGCAGCTCCAACTCATTCCCTTCCCCTCGGCCGAGTTCGGCATCCCCGAGGGGTGCGAGAACCTCACCTCTCTCCCCCTCCCCCTCGCCGCCGCCTTCTTCAACGCCATCTTCGCCCTGCGGGCCCCGCTCGGCGCGCTGTTGCGGGAGCTCAGCGCTCACGCCCTGGTCGCCGACGCGCTGTTCCCGTGGGCCACGGGGCTGGCGGCCGAGATGGGGATCCCGAGGCTTATCTTCCAGGTCACGGGTCTGTTCCCGCTCTGCGCTGCCCACGATCTCGATGCACATCGGCCGCACGAGGCCGTTGGTGGGGATGACGAGGAGTTTACCATCCCGGGGTTTCCGGACCCGGTGACGCTTACCAGGGGGACAGATCCCCGAGGTCTTCAGGCACAATTTCATGCTGGCCCTCCTCCGCGACGCAGAGCTCACCAGCTACGGCGTGATAGTGAACAGCTTCTACGCCCTCGAGCCCAGCTACGCGGAGCACTACTACAAGGTGGCCCCCCGGAAGGTCTTCCTCCTCGGCCCGGTCGCCCTCGCCGGCTCCAATCCCTCGCCGCCGCCATTGGAGAGTGGCGACCCCTGCATCACCTGGCTCGACTCCAAGCCCGACGACTCGGTCCTTTACCTGAGCTTCGGCACCCTCTGCCGATTCAGCAACGAGCAGCTCGTCGAGCTCGCTGAGGGCCTCGCGTCCTCCGGCCACAACTTTGTGTGGGTCGTGGCCCGCCCCGAGAGCAGCGGTGGGCCCGGCGAGGAGTGGCTTCCGGAAGGCTACGAGCGTAATGTGGCGGGCCGGGGGCTTCTCGTAAGTGGTTGGGCTCCGCAGACCGCGATCCTGAACCACCGGGCGGTGGGTGGGTTCGTGATCCACTGCGGGTGGAACTCCGTCATGGAGGCGGTGGCGGCGGAGGTGCCCATGGCCACGTGGCCGCTTCACTCAGAGCAATTCGTCATTGAGAAGCTGCTCTGCGATGTGCTGCACGTGGCGGTGCCAATGTGGGAGGGGTGGAAAAGCATCTGGGACGATCAGAAGGAGGTGGTGCGGGCGGGAACAGTGGCAGCGTCGGTGAAGCGGCTTATGGGGGGCGGGGACGAGGTGGAGGCGATGCGAAGGAGAGTGAGGGAGCTCGGGGAATTAGGACGGGCCGCGGTGGCAGAAAGCGGATCATCCCACTCCGACATGGGCCGTCTTATCGACGTGTTGACGGAGGAGCGAAGCAAGGCTAAGGAAATGATAGACAAGAAGGGTGTTAGTTATGATTGTTCTGGTCGCAACTGATTATTTCTCGCCAAGAACTAGCAGTGTTTTTTGATAAACATGTTATGTTGGCGATCTCACTTGGAACTCAACACATGGCGAGGTATTTCATCGAGCACTGCGTACATTATTTTGGACATGTCTCTAGTCTAGATGAATTGGCTCCAACACATGATTTTTTTGTCATTTTTTTGCTTATTTTGCTCTCTAAGCTAAGTGATAAGTTGGGAGCTTTTTTG

>F01_transcript/43260 full_length_coverage=5;length=1757;num_subreads=60

GGAAGAAATCCAGACATTCTTCCTCCATTGCATATTTTCACTCACAGCGAGCACTAAAATGACGAGCTCGACTGAGTGGGTAGTCCTCTACCCGTCACCGGGGATGGGCCACCTCGTCTCCATGGTCGAGCTTGCGAAGGTCCTCCTCGGAAACGGGCTCTCGATCACCATCCTCATGGTCGAGCCCCACTACAACACGGGCGCCACCGCTCCCTTCATCTCCGCCGTCTCCGCCGCTCACCCCTCCATTGCCTTCCACCACCTCCCCTCCCTCTCCCCACCCCCCAACTCCTCCCCACACCGTGAGGCCCAGGCCATGGACCTCCTCCGCCTCTCCAACCCCAACCTCCAATCCTTCCTCCAATCCTCATCCCTCTCTCCCGCCGCCCTGATCATCGACTTCTTCTGCGGTTTCGCTCTCAATGTATCCTCCCATCTCCGTATCCCAACCTACTGCTTCTTCACCTCGGGCGCATGCTTCCTCGCCGCCTTCCTCAAGTTACCCAACATCCACTCCCGATTTCCGGTCAGTTTCCGAGAGCTCGGCTCGACCCTTATCGACGTCCCCGGTATCCCTTCCATCCCTGCCGATCACATGCCGCTGCTACTGCTTGATCGGGACGGCGAAGCTTATAAGGGGTTTTATCACCTATCGGAGCGATGTTCTGAGTCGGACGGGATAATCGTCAACACGTTTGATGCGCTCGAGCCAAGGGCGATCGAGGCCATCTCGAGAGATGGAATGAAACTGTACTGTACCGGGCCGCTGATCACCGAGGAGAGGGACATGACGAACGGGGCGGATTGCCTTGAGTGGCTCGATGGGCAACCGAGGGGGAGCGTGGTGTTCCTTTGCTTTGGGAGTCTTGGACTTTTCTCAGTTGAGCAGCTGAGGGAGATCGCCATCGGGTTGGAGCGGAGCGGGCAGCGATTTCTGTGGGTGGTGCGGAGCCCGCCGAGCGACGATCCGGCCAAACGGTTCGCTAAGCCGGCTGAGCCGCTCCTCCACGTGCTGCTCCCCGATGGGTTCCTCAACCGAACAGGCGACAGGGGAATGGTAGTGAAGTCGTGGGCCCCACAGACGGCCGTTCTGCGCCACCCGTCAGTGGGCGGCTTCGTGACGCACTGCGGGTGGAACTCCATCTTGGAGGCGATATGCACGGGGGTCCCCATGGTCGGGTGGCCACTCTACGCTGAGCAGCGGGTGAACATAATTTTTCTCGTCGAGGAGATGAAGCTGGCCGCGGCGATGGAGGGCTACGATTCGCCCCTCGTGCCGGCGGAGGAGGTGGAGACACGTGTGGGGTGGCTGATGGACTCGGACGGCGGGCGGGAGCTGAGATTGCGTACGGAAGCGGCGAAGAAGGCTGCGGCCACGGCGCTGCAGGAGGGCGGGTCGTCACGTGAGGCGTTGGTGGAGCTGGTGGCGCGGCTGAAACGGAGGTGATGGAGGCGGGTGTCCTTATCGACCGTGTGGGTGCCGTGACGAGATTAAATTCAAGAGCTGGTAATAAAACGAAGCAAAACAAAATGGGAAACTTTGGGGTCTGGAATTCTATTTGTTGCTTTGGTGCCGTGACGACATTAAATTATGATTGTCGTCTTTGTATTTTTTTTTTTTCTTCTCAGTTCAGCGTCTCCTGTAACAATGTTTTCAAAATCGGACCGATGAAATATAAATACATTGGTCGAACCAATTCAGTGTGTAAATGCTTTTTATTAGTAAAATGAAGTGAGCTAAATATTGGCCGGTAGAT

>F01_transcript/44192 full_length_coverage=9;length=1714;num_subreads=60

GGAACAACATAACCTTGGCAGTATCAATCTCCTGTTAACTCTCTCATCCCCATGGGTTCCGAAGCTCAACAACTCCACATCCTCTTCTTCCCCATGATGGCCGGCCGGCCACATGATCCCCATGCTCGACATGGCCCGCCTCTTCGCCTCCCGGGGCGTCAAGGCCACCCTCGTCATCACCCCGGCCAACTACGACGCCATCCGATCCACCGTCGACCGTGCCAAAACACCAAATATAGAAGTCCTCTCCATTCCCTTCCCCTTCCAGAACCTCTCCACCAATCCCTCGCCGGAGTTCAAGGCCAAGTTCCTCCAGGCCATCGAGATGCTTCGCGACCCCTTCGACCGAATCCTCTCCGAGTTCACACCCGACTGCGTCGTCACCGACATGTTCTTCCCCTGGTCCGCCGACGTTGCCGCCGAACACGACGTGCCGAGGCTCGTGTTCCACGGATCGTGCTGCTTCTCGCAGTGCGCCATGGACGCGATATTTCGTTACGACGCAGTGGAGAGCTCACCGGTCGATGTCGAGTCCTTTGTGGTTCCGGGTTTACCCCATAGGATCGAGATGCTCAAGTCACAGGTCATGGATTTTAAATCAATGCCCGCCATGGGGGAACTCTTTCGTAGAGTGAAGGAGTCGAAGGAGCGGACCTACGGCGTGGTGGTCAACAGCTTCCACGAGCTCGAGCCGGAGTATGTTCGGCACTACCGCGAGGTGATGGGGTGGAAGACGTGGCAGGTCGGACCAGTGTCACTGTCCAACACAAACAATGCAGACATGTTGGGGAGAGGGGGTAATATAATCTCCTCACATGCTTGCTTGGAGTGGCTCGATGGTAAACAACCGGGCTCGGTATTGTACGTCTGCTTCGGGAGTCTTGGCGAATTCACTGCGGCCCAGCTCATCGAACTCGCGGTTGGGCTCGAGACGTCCGGCGTTCCATTTGTTTGGGTGGTGAGGAAGGGCGCGACCGAGCTTTTGCCGGCGGGGTATGTGGAGAGAGTAAAAGAAAGGGGGTTGGTGGTGGAGGGGTGGGTCCCGCAGATGTTGATACTGAACCACCGAGCGATAGGAGGGTTCCTGACACACTGCGGGTGGAACTCGAGCATCGAGGGGATCTCCGCCGGCTTGCCGATGGCCACATGGCCGCTGCATGCGGAGCAATTCTACAACGAGCAGCTATTGGTCAAGGTGTTGGGGGTCGGTGTGGCGGTGGGGATAGAGAAGTGGATAATGTTGCCTGAGGACAGGCCCGTTGTGGGGTCGGCGTCCGTGGAGAGGGCTGTGGCGAGGCTGATGGGAGGCGGGGAGGAGGCGCAGAGGCGGCGACGGAGAGCGAGGGAGTTATGTGATATGGCAAAAGCGGCCGTGTCGGACGGCGGATCATCGTTCTTGGACATGGGCAATCTAATCAAGGAATTGACGGACATGCGAAATTTGACTTAGAAAAATAAAAAATGAGTCTCGCTCAAAATGCAATAGTAATCGTTAATTAAGCTCTCTGCTTATAGTAAGTTTCGTTGAGGAAACGTTGATCTAGTTATATGTTGTGAAACACATAATATTATATATACAACATGTTTTTTTGTTTAATTTTTTTTTTTTACTATGATGATGATAAATGTCCATCGATGATGAATTATGTAGATCAACTCACTAGACACAACATAAAGAGCAAGAGAGACAAATTTTTTAGATAGTTCACTCGCG

>F01_transcript/44385 full_length_coverage=28;length=1703;num_subreads=60

GAGACACCAGAGCTCTCTCTCTCTCTCTCTCTCTCTCTCTCTCTCTCTCTCTCTCTCTCTCTCTCGTCATGGATCGAGAAAAGCTACACATAGTTGTGTTCCCGTGGCTAGCCTTTGGGCACATGTTGCCTTTTCTTCAGCTCTCCAAATCTTTGGCCACAAGGGGTCATCGAATCTCCTTTATCTCCACCACAAGAAACATTAAAAAGCTCCAGCCCAAAGTGCCACCCACGCTCTCCCAACTCATTCAATTCATCCCTCTACCTCTCCCCTCGGTCGACTGTTTACCGGCTACCGCCGAGTCTGTCTACGACATCCCTCTAAACCTCAGCCTTCACCTCCAGAAGGCGTTCGACGGCCTCGAGCACCCATTGGCCCAATTCCTCACTTCATCGTCTCCAAAGCCTGATTGGATCATCGTGGACTACGCCTCCGATTGGCTCCCGCCACTTGCATCAAAATTCAATGTGCCGTGTGTCCACTTCTGCACCAACATGGCCTCAACCATGGCGTTCATAGGACCCGTGTCAGAAGTGGACAACATAGACTCCATCACCGTTGAAAATGTGATCGTTCCGCGCAAATGGATGACATTCCCTACAAATGTAGCCCTTCACCCTTATGAGGCTCGACAGGTTCTTGATTTGTCCAGTTCAGATAATGCTTGGGATGGCTACCGCTACTTGCTCACCATCAGAGGTTGCAAGGCCGTAGCCATCCGAAGTTGCAACGAGTTCACGCCCAAGTGGTTGCCCCTCCTTCGAGATATCTACAAGAAACCCGTCATTCAGGTTGGCATGCTTTCCTCGATGTTGGTCGAAAATGGAGATTATGACAACAACGAGCTCAATGTGATAAAGTGGCTGGATCGACGATCTCCGAGTTCTGTCGTGTATGTAGCCTTTGGTAGCCAAGCCGAACTCAGCATCGAGCTCTTACACGAGCTAGCGCTTGGGCTTGAGCTCTCTAACTTGCCTTTTCTTTGGGCTTTTAGAAAGCCGACCACCAACGAAAACGAAGAGATTTTGCCCATAGGATTTCAAGAACGGACAAAGGACCGAGGACTCGTCGTCATGGGTTGGGTTCCCCAGCTCAGGGTCTTGGCCCATGAGTCGGTGGGTGGATTCATGACGCACTGTGGTTTTAGTTCGATCATCGAGAGCCTCCATTTTGGTCGTCCCCTAATCCTCTTCCCTTTATTCGGGGATCAAGGGCTCAATGCTCGGACGATGGTGGAGGAGAAGATCGGATTCGAGGTGAAGCGGGATGAGGAGGATGGGTCACTAACAAAGGAAGCGGTGGCCGAGGCATTGAGGTTGGTTATCGTCGAGACAGAGGGGGAACCTTTCAGGAAAAAAGCCAAAGAGATGATGGGGGTTTTCACGGACAAGAATTTAAATGAGAGATATCTGGATGATTTCACTCAGTATCTAATGGACCATAAAGATAGCGAATAAAAACAGCTCAACCGAGGAGTTCAGGCAAGAGGAAGGGTAGTGCAAACCAGTGCCACTGCTAAGGATATCTCCTTATATTTTTAGGATGATGATAACCTATATCTAGTGTGTGTTGTATGTGTTTGGGACACCAGCTTATAATATTCAAAAATGTGATCTAATTTAGAAAAAATAAAGCTATAAGATGTTGTAATAAGAGGAAACATGTAATATTATGTTTTAGAAATAAATATGAATATGGCTTGG

>F01_transcript/44405 full_length_coverage=10;length=1764;num_subreads=60

GAACATCATCTGTGATCATCAAAATAAAAAGTTGATGGTAATATCAACCCCCATGAATCCCCCCATCTCCCCTCAAAATCCATTCGTCCTCCACACTCCCCAAATCCCAACCCATCTCCGCCATGGGCTCCGACGATCGTCAACCTCTGACGGCGTTCTTCATCCCCTACTTCGCGACGGGGCACATGATCCCCCTCGTCGACATCGCCCGCCTCCTCGCCGCCCGTGGCGTCGACTCCACCGTCCTCGCCACCCCTGCCAACGCCGCCCTCATCCAACGCACCGTCGACTACGCCGCAGCTACCGGCCTCCCGATTCGAACCCTCACCTACCCCTTCCCCGCGGCGGAATCCGGCCTCCCCCCCGGCGTCGAGAACATCAGCCGCCTCCCTCCCGCTGACGCCCACCGGATCGACGACGCCACTTGGCACGTGCGGCCCTCGCACGAGCGCCTCATTCGGGCCCACCGACCAAACGCCGTCTTCTCCGACATCCACTTCCCCTGGACCACAGCCATGGCGAAAGAGGTCGGGGCGGTGAGGATCTCCTTCGACGCCCTCGGGCTGTTCCCCATATCGGTTATGAACGTCCTGTTCGGAAAGCTACCACACCTAGCCGTCACAAGGGACGACGAGCAGTTCCTCGTACCGGGGCTGCCGCACCCGATTCACATGGTCCGGCCCGAGCTGCCAGATTTTCTCCGCGGCAAGACGCACATCACAGGTCTCATGGAGAGCATTGGGGAGGCGGAGGCCGGCAGCCTTGGCGTGGTCGTCAACAGCTTTTTAGAGATGGAGTCGGAGTACGCTGACTACTACTACAAGGCCACCAACCGGAAGTTATGGTGTGTGGGCCCCGTCTCCCTCGCCTACACTGCCGGCGATGATGTGGCAACTCGTGGCAGCGACAACCCTGCGGCCGCAGCCAATCGCCGGCAGTGCTTCGACTGGCTCGAGGGGAAGGAGGCTGGGTCGGTGGTCTACGTGGCATTTGGGAGCTGGTCTCACTTTTCAGACGAGCAGCTCCGTGAAATGGCGGCAGGGCTCGAGTTTTCCGGCCACCCGTTCATCTGGGTAGTACGAGACGATGGGGCCACGTGGATGCCCGCAGGGTTCGAGGAGCAGCTGGCAGGGCGGGGGCTGGTGGTCCGGGGGTGGGCCCCGCAGGTGTCCGGTGCTGGGCCACGCGGCAGTGGGCGGCTTCGTGACCCACTGCGGGTGGAACTCGGTGCTGGAGGGCGTCACGTCCGGCCTCCCGATGGCCACGTTGCCCCTCTCCACGGAGCAGTTCATCAACGAGAAGCTCGTCGTCGATGTCCTCGGCGTCGCCGTCAGGATCTCCGAGGCCCCACGTGGTACAGCGGACGGCGACGGTGCCATCGTGCCCGCTGCGGACGTGACGCGCGCCGTCGACCGGATCATGGCCAGCGACGATGAGGCGACGACCATGCGGGCGGCGGTGCGGCGGCTCGGGGCGGCGGCGAGGGTGGCGGTGAAAGAGGGCGGGTCGTCTTATGTTGGTCTCACTCGTCTCATCGATGAGATTCGGGCGGGATCTAGAAAGGAATCGAAAGTCTGATCTATTTGCTTTGAAACTCGGATCAGCCATAATAAGAGTCCAACTTGTAAGGACTCTCTCTTATCTCACTCACAAATATGTATGTTTTTGTGTATTCTTTGCTTGGAACTAAAATGAACTGCAAAATGTAATGTCCTTTTTTCCAAGTATTTGAATAAGAAGACGGGGAATGTTTTTTATTTTTCT

>F01_transcript/44442 full_length_coverage=90;length=1690;num_subreads=60

GACACCAGAAGAGCCTCTCTCTCTCTCTCTCTCTCTCTCTCTCTCTCTCTCTCTCTCTCTCTCTCTCGTCATGGATGGAGGAAACCTGCACATAGTTGCGTTCCCAGTGCTAGCCTATGGGCACATGATGCCCTATCTTGAGCTCTCCAAATTGTTGGCCACAAGGGGCCACCAAATCTCCTTCATCTCCACCACAAAAAACATAAAAAAGCTCCAGCCCAAAGTGCCACCCACACTCTCTCAACTCATTCAATTCATCCCTCTAACTCTCCCGCCGGTCGACGGCTTACCGCCTACTGCCGAGTCCTACACGGACATCCCCCAAAGCCTCGTCCAATACCTCAAGAAGGCATTCGACGGCCTCGAGGGCCCCTTCGCCCAATTCCTCAGTTCATCGTCCCCAAAACCTGATTGGATCATCGTAGACTACCTCTACCCCTGGGTCCCACGGGTCGCTTCCGAATTCAATGTGCCGTGTGTCCACTTCAGCATCGTAGCGGCCACGGCCCTGGCATTCTTCGGACCCGTGTCAGAAGTGAACAACATGGACTCCCTCACAGTTGAAAAATTAATCGTTAAGCGCGAATGGGTGACATTCCCTTCAAATGTGGCCTTACGTACTTATGAGGCCCGAGGGGTACTTGATTGCTTCAATTCAGACAATGCTTGGGATGCCCACAGCGAATTGCTCACCATCGGAGGTTGCAAGGCCGTGGCCATCCGAAGCTGCAACGAGTTCGTGCCCAAGTGGTTATCCCTCATTCAGGATCTCTACAAGAAACCCGTCATCCAGGTTGGCATGCTTTTTCCAACGTCTGGTCGAAAAATGGCGATTATGACACCAGTGAACTCAAGATCATGGAGTGGTTAGACCGACGATCTCCAGGTTCTGTCGTATATGTAGCCTTTGGGAGTGAAGCTGAACTCAACATTGAGATCTTACATGAGCTAGCGCTTGGGATCGAGCTCTCTAACTTGCCTTTCCTTTGGGCTTTTAGAAAATCAACCACCTATGAAAATGAAGATATTTTGCCCGAAGGGTTCCTAGAACGAACAAAGGATCGAGGATTCGTCGTCATGGGTTGGGTTCCCCAGCCTAAGATTTTGGCCCACGAGTCAGTGGGAGGATTCTTGACGCACTGCGGATATAGTTCCGTCATCGAGGGCCTCAATTTTGGTCACCCCCTAATTCTCTTCCCTATATTCGGGGATCAACCGCTTAATGCTCGAATGATGCACGAGGAGATGATCGGATTCGAGGTGAAGAGGGACGAGGAGGATGGGTCACTAACAAAGGAAGCGGTAGCTGAGGCATTGAGGTTGGTTGTCGTCGAGGCAGAGGGGGATCCTTTCAGGAAAAAAGCCAAAGAGATGATGGAGGCTTTCACAAGAAAGAGTTGCAATGAGAAATATATAGATGATTTCATTCAGTATTTAATGGGCCATAAGGATAGCAAATGAAAGTAGCTTACCCGAGGCATCAGACAAGAGGAAGGGCAGTGCATAGTAGGGTCACAACTAAATGCATCTCCTTCCATTTCTAGGATGAAGGGAACCTATATGTATTGTGTGTTGTAGGTGTTGGAATACCAACTTATAATATAAAAAAATATAATGTAATTTAGGGGAAATAAAGTTGTAAGATGGAATAATAAGACGAAACATATAAAATGGCTTTTATATTGATTAG

>F01_transcript/44445 full_length_coverage=19;length=1719;num_subreads=60

GAGTTTCAAGTAGAGAGAGACAGAGAGAGAGAGAGAAAACAAACACTGATATCACCCACATCTCAATGGAGTCCCAACCAGAGCTCCTCCACCTAGTTTTCTTCCCCTTCCTCGCTCGCAGCCACATGATCCCCATGCTGGAGACCGCCCGCCTCGCCGTCGAGCGCGGCGTCAAAACCACCCTCGTCACCACCCCTGCCAACGCCCACCTCATCCACCCCGTCCTCCACCGCTCCAACTCATCTCTCCTCCCCTCCCATCCGCCGATGCAGCTCCAACTCATTCCCTTCCCCTCGGCCGAGTTCGGCATCCCCGAGGGGTGCGAGAACCTCACCTCTATCCCCCTCCCCCTCGTCGCCGCCTTCTTCAAAGCCATCTTCGCCCTGCGGGCCCCGCTCGCCGCGCTGTTGCGGGAGCTCGGCGCCCACGCCCTCGTCGCCGATGCGCTGTTCCCGTGGGCCACGGGGCTGGCGGCCGAGATGGGGATCCCGAGGCTTATCTTCCAGGTCATGGGTCTGTTCCCGCTCTGCGGTGCCCACGATCTCGATACCCATCGGCCGCACGAGGCCGTTGGTGGGGATGACGAGGAGTTTACCATCCCGGGGTTTCCGGACCCGGTGAAGCTTACCAGGGGACAGGTCCCCGAGGTCTTCAGGCACGATTTCATGCTGGCCCTCCTCCGCGACGCAGAGTTCACCAGCTACGGCGTGATAGTGAACAGCTTCTACGCCCTCGAGCCCAGTTACGCCGAGCACTACTACAAGGTGGCCCCCCGGAAGGTCTTCCTCCTCGGCCCGGTCTTCATCGCCGGCTCCAATCCCTCGCCGCTGCCATTGGAGAGAGGCGACCCCTGCATCACCTGGCTCGACTCCAAGCCCGACAACTCGACCCTTTACGTGAGCTTCGGCAGCCTCTGCCGATTCAGCGACGAGCAACTCGTCGAGCTCGCCGAGGGCCTCGCGTCCTCCGGCCACAACTTTGTGTGGGTCGTGGCCCTCCCCGAGAGCAGTGGCTCGACCGGCGAGGAGTGGCTTCCGGAAGGCTACGAACGTAATGTGGCGGGCCGGGGGCTTCTCGTAAGTGGTTGGGCTCCGCAGACAGTGATCCTGAACCACCGAGCGGTGGGTGGGTTCGTGACCCAATGCGGGTGGAACGCCGTCATGGAGGGGATGGCGGCAGAGGTACCCATGGCCACGTGGCCGCTTCATTCAGACCATTTCATCACCGAGAAGCTGCTCTGCGATGTGCTGCACGTGGCGGTGCCAATGTGGGAGGGGAGGAAAAGCATCTGGGACGATCAGAAGGAGGTGGTACGGGCGGGAACAGTGGCAGCGTCGGTGAAGTGGCTTATGGGAGGCGGGGACGAGATCGAGGCGATGCGAAGGAGAGTGAGGGAGCTCGGGGAATTAGGACGGGCCGCGGTGGCAGAAGGCGGATCATCTCACTCCGACATGGGTCGTCTTATCGACGTGCTCACGGAGGAGCGAAGCAAGGCTAAGGAAATGATAGACAAGAACGGTGTTAGTTATGATTGTTCTGGTCGCAATTGATTATTTCTCGCCAAAAACTGATAGCAGTGTTTTTTGATAAACATGTTATGTTGGCTATCTCACTTGGAACCCAACACATTGCGAGGTATTTCATGGAGCACTTGTGTACTTTATTTTGGACACGTGTCTACTCTAGATGAATTGGCTCCAACACCTGATTTTTTTTGCC

>F01_transcript/44734 full_length_coverage=2;length=1710;num_subreads=60

ATCCAAGGCATTCTTCCTCCGTTGCATACTCTTACACTCACAGCGAGCACCAAAATGAAGAGATCGGCTGAGTCGGTAGTCCTCTACCCGTCACCCGGAATGGGCCACCTTGTCTCCATGGTCGAGCTTGCGAAGCTCCTCCTCGGAAACGGCCTCTCGGTCACCATCCTCATCGTCGAGCCCCACTACAACACGGGTGCCACCACCCCCTTCATCTCCGCCGTCTCCACAGCCCACCCCTCCATCGCCTTCCACCGCCTCCCCTCCGTCTCCCCTCCCCCTAACCCCTCCCCCCATCACGAAGCCCTCGCCTTCGACCTCCTCCGCCTCTCCAACCCTAACCTCCGCTCCTTCCTCCAATCCACATCCCCCTCTCCCGCTGCACTCATCATCGACTTCTTTTGCGGTTTTGCTCTCCATGTATCCTCCGATCTTTGTATCCCAACCTACTACTTCTACACCTCGGGTGCATGCGTCCTCGCCACCTTCTTAAAGTTACCCAACATCCACTCCCGATTTTCCTCCAGTTTCCGAGAGCTCGGCTCCACCCGTATCGATGTCCCTGGCATCCCTCCCATCCCTGCCGATCACATGCCGCTGCCGCTGCTAGATCGAGACGACGAAGCTTATAAGGGATTTTATTATCTATCGGAGCGATTTTCTGAGTCGGACGGGACAATCATCAACACGTTCGATGCGCTCGAGCCAAGGGCGATCGAGGCGATCTCGAGGGATGGAATGAAGCTGTACTGTATCGGACCGCTGATCACCGAGGAGAGGGACAGGACGAAAGGGGCGGATTGCCTTGAGTGGCTCGATGGGCAACCGAGGGGGAGCGTGGTGGTCCTTTGCTTTGGGAGTCTTGGACTTTTCTCAGTTGAGCAGCTGAGAGAGATCGCCATTGGGTTGGAGCGGAGCGGGCAACGATTTCTGTGGGTGGTGCGGAGCCCGCCAAGCGACGATCCGGCCGAACGGTTCGCTAAGCCGGCTGAGCCGGACCTCCACGTGCTGCTCCCTGATGGGTTCCTCAACAGAACAGGACGACCGGGGAATGGTGGTGAAGTCGTGGGCCCCGCAGACGGCCGTTCTGCGCCATCCGTCAGTGGGCGGCTTCGTGACGCACTGCGGGTGGAACTCCATCTTGGAGGCGATATGCATGGGGGTCCCCATGATTGGGTGGCCAATGTACGCTGAGCAGCGGATGAACAAAGTCTTTCTGGTCGAGGAGATGAAGCTGGCCGTGGCGATGGAGGGCTACGATTCGCCCCTCGTGCGGGCGGAGGAGGTGGAGACACGTGTGGGGTGGCTGATGGAGTCGGACGGCGGGCGGGAGCTGAGGCGGCGCACGGAAGCGGCGAAGGAGGCTGCGGCGGCAGCGCTGATGGAGGTCGGGTCGTCACGTGCGGCGCTGGTGGAGTTTGTGGGGCGGCTGAAACGGAGGTGATGGGTGCGGGTGTCCTTATCGAACGAGTGGGTGCCGTGACGAAATTAAATTCAGGTTCTTCTAATAAAGCCAAGCAAAAAAAATGGGGAGCTTTCGGAAGTGGAATCCGTTTGGTGCAGTGATGAAATTAAATAATGATTGGCCTCTTTGTTTTTTTCGATTTGAATATGACCTGTCAGCTTTCCCGTTTTAAAATCATCTCCTCGCTTTGAATGGAGGATCGCATTTACCTAATATTGATTTATATAAACAATAGATTAAGGCCC

>F01_transcript/44939 full_length_coverage=3;length=1703;num_subreads=60

GGAAAAAATAAAAAACTATCATTCTCCGGCGAGAATGGGGAGTGAGAGCAGGCAGAAGCACTTCGTGCTGGTTCCGTGGCTGTCCCACGGGCACGTGATCCCCATGATGGACATGGCCCGCCTCCTCGCCGAGCGGGACGGCATCCACGTGACCGTGGCCATCTCCCCCGTGGGCGCAGAACGCATCCGTAGCTGCTTCATAGAGCCCGTGGCCGCCGCCAAGCTCCCCATCTCCTTCGTCGACCTCCCCTTCCCCTGCGCCGAAGCCGGCCTCCCTGACGGCGTGGAGACCATCGAGCAAATCCAGGACCCCTCCCTGTTCCCCAAGATGCACGTGGCAGCCGGCCTCCTCCGCAAACCACTCGAGTCCATACTCCGAGAGCTCCCCCGCAAGCCCTCCGTCATCCTCGCTGACCTCTACCACCCGTGGGCCCGGGAAGTCGCCGCCGACCTTGGCGTCCCGCTGCTGCTCTACTACGTGTTTCCCTGCTTCACCATCCTCGTCTACCGAAGTTTGAGACAGCATGGTATCTACGATGACGGCGCGGCAGATGCGAGTCGGATGTTCCCGGTGCCCGACGCCCCGGAGTACATGGTCAGCCGGGCACAGGCGCCGGGGACCTTCGACAGGCCCGGATGGGAGTGGCTTCGCGAGGAGGCTATTGCAGCCGAGTCCGCCGCCGCCGGGGTTATTTTTCACAGCTTCGACCAGCTCGAGCCCAATTTCCTCCCCAAGTTCCAGGAGATCATGGGTGGCCTGAAGACGTGGGCCATCGGCCCGCTGTCCCTCAGCCACAAGAACGTGCTGGCAGAGCGCGGGAGCGCAAATGAAGTGGCAGCCGACAGCTGCCTCACCTGGCTCGACGCCAACGCCCCCGCCTCGGTCATCTACGTCTGCTTTGGCACCAACACATACTGGACCCCTCAGCAGATCATTGAGGTCGGGTCCGGGATAGAGAGCTCGGGCCACCCCTTCATCTGGGTGCTGAAGAAGCGGGAGCTGACGCCCGAGGTGGAGGAGTTCCTGTCGGGAGAGTTCGAGGAGCGGGTGCAGGACCGAGGCCTGCTCATCAGGGGCTGGGCCCCTCAGGCGGCCATACTGACCCATAAGTCAATCGGGGGATTCATGACGCATGGCGGGTGGAACTCGTCGATCGAGGGGGTGGCGGCCGGGGTGCCAATGCTGACGTGGCCGCACTTCGAGGACCAGTTCTTGCACCAGATGATCATCGTTCAGGTGCTAGGGATGGGGATTGGAGTCGGGGTGCGGGCGCAGGAGGACTACATTCGCGCAGGTGATGGACACCATCAAGCGGGAGCAGGTCGAGAAGGCGGTGAGGGAGCTGATGGGAGGAGGGGAGGAAGCCGACGCGAGGAGGAGGAAGGCGAAGGAGTACGGGGAGAAGGCGAGGAATGCCATGGAGGTCGGGGGGTCGTCGTACGTGAACCTGACCGAAGTGATCGACTCCGTTCCGTTCGTCGCCGCCACCGAGAATGGTGGCGGTGACTAACTACTTTCTTCATTCATCTTCTTGGTGCCGAAATCAAATTGCCGCAATTTCAAATTCCTCTACCTATCTCTGTGCGTCGACTGTAATAAAATAGTACTATTGAATCAGTGATTAGTATTTTTTCTTTTGTCGCTTAATTTTTGGTTGGATTAGGCGACCAATGCATCTTGAGCGAGTGTCTTTTTTTTTTGG

>F01_transcript/45099 full_length_coverage=16;length=1716;num_subreads=60

GCCCATGGAAACGGAATGGGCCCAACTAGAGAGGTGCTCGCAGACCCCCCGTCTTATCCTCCTCCGCGGCTTTAATTCTGACGGTGGAGGCTGAATCCAGCACGAAGGGATCGATGGAAGCTCAGAGGGAAACGGTGACGCACGTGCTTGTGTTCCCCTTCCCGATCCAGGGTCACGTGGCCAGCATGCTGAAGCTCGCTGAGCTGTTGTCCCTCGCCAGAATCCACGTCACCTTCCTCAACACTGACGGCAACCACCAGCGCCTCCGCAGCTTCTCCGGCTCTTACGCACGGCTCGACCAGAGAACCAGTTTTTATTTCCGGTCCATCCCCGACGGTTTGCCGGAGGACCATCCCCGGTCCGGCGGCTCACAAGTGTTTGACCTCGTCCACTCGCTGAGGGCCCGGTCCATGGCCCCGTTTCGGGAGCTGCTGATCGCCGGCGGATCAGAGGGGAGGCCGCCGATCACGTGCGCGGTGGTGGACGCTCTGATGAACTTCGCGGTGGAGACGGCGGAGGCTGCGGGGATCCCGACGGTTGCTTTTCGTACAATCAGCGCCTGTAGCTTCTGGGCGTATGCCTGTATCCCCAACCTCATACAGAGTGGGGAAATCCCGTTGCCAGAGGGAGCGAACATGGATGAGCCGATCCAAAATATCCCCGGGATGGAGCATTTTCTCAGACGCCGAGACCTCCCCAGCTTTCTTCGACAAGTGAAAGACATAACGGACCCCAACTTCCAGTTCCTCGCCGACGCCACATCTGACACGAGCCGAGCGAGAGGGACTCATCCTCAACACCCCAGAATTCTTGGAGGGATCAGTGCTCGTCCGAATCAAAACCCTTTGCCCCATCACATACGCCCTCGGGCCCCTTCACTCCCTCCTCACAACTCACGCCTTCACCTCCGGTGTTGAAGCTGGCTTCGACATCTCTGGCAACTTGATGCAAGTGGACTTGACCTGTATGGAGTGGCTCGACTCCCAGCCTGATCGGTCCGTGGTCTACGTTAGCTTTGGTAGCCTCACAGTGGTCTCGAGCGAGGAGCTTTCAGAGTTCTGGCACGGTCTGATCGACAGCGGTCATCGGTTCTTGTGGGTAGTGCGATCAGACCTCGTCAGTGATCATGAGTCGGTGTCGATGTTGCTGGATGATGCGGCGACAGCGACAAGGGAGAGGGGATGCATTGTGGGGTGGGCGCCGCAAGGGGAGGTGTTGGCCCACAGATCGGTGGGTTGTTTCCTGACGCACAGTGGGTGGAATTCGACGTTGGAGAGCGTGGTGGCGGGGGTGCCGATGATCTGCTGGCCATTTTTCGCGGACCAGCAGATCAACAGCAGGTACGTGAGCGAGGTGTGGAGGGTGGGTCTCGACATGAAGGACATTTGCAAGAGGGACATGGTGGAGAGGATGGTGAGGGATGCGATGGAGGGAGAGAAATCGGAGGAGTTGAGGAGGTCGTCGAGGGAGATGGCTGAGATGGCACAAAGGACTATCGAGGAAGGTGGCTCGTCATATAATGACTTTTGGCGTCTCGTCAAGGATATCAAGTCCTTGAGCTCACTCAAATCTTGAGATCATGTGGTGTTATGAGTTTTTACTATAATGGCAACTTTTCGTTAGTCATTTTCTTATGATGCCTTTGTGTATTGTTCTACAAAATATTGGCGTTTGAACTCTAATAAGATTGAACAAATTTTAAAAGAATTTTTCATG

>F01_transcript/45112 full_length_coverage=4;length=1710;num_subreads=60

GATGACCAAACAAACACATCCAAAAATGACCAACCGAACAATCATGGATGCCTTTGGTCATCCCCGTCCTCACTTCGTGTTGGTTCCTCTAATGGCCCAAGGCCACATGATCCCCATGCTGGACATGGCCTACCTCCTCGCCGACCGTGGTTCCCTCGTCAGTTTCATCACCACTCCTGTTAACGCTGCACGAATCAAGTCCACCATCGATCGTGTCAAGGACTCTCGCCTCCCAATCCGATTCATCGAGCTACCATTCCGCTCCTCTGAAGCTGGATTGCCCGACGGATGTGAGAATGTCGACCTCGTTGAATCTCTCGGCTTCCTCAAATCCTTCTTTGAGGGCACACACATTCTCTCAGAACCCTTAGAGCTCTATCTCCAAAAACAACAGCACGGCCCAATTTGCATGATCTCCGATTTTAGCCTCCCGTGGACCGCAGAGCTCGCTCGGAAGCTTAATATACCCAGGTTGGTATTCCATGGACCCTCATGCTTCTACCTTCTCAGTGTTCATAATATTCAGAAACATAGAATCTCCGGCCGCATCACTCATCAATTTGAGCCATTCCTTGTGCCCGACTTGCCACAACAGTGGTCCATAGTGGTCACGAGAGTACAATCACTGGGGTTCTTTGAGATTCCAGGATGGGAGGAGTTCCATAAGAAGATACTAGAGTCGGAAGCGGCAGCGGAAGGAGTTGTCATGAATTCATTCGAAGAATTGGAGGATTGTTTCATTGAGAGCTATCGAAAGGTTATGGGGAAGAAGGTTTGGACAATTGGGCCTTTGTGTCTCCACAACAAGGATAACTCCTATAAGGCCGCGAGAGGGAATAAAGCTGTTGTTGACGAACATCGAGTTCTCAATTGGCTCGATTCTATGAGCAGAGATTCAGTTATCTACATTAGCTTTGGTAGTCTAGCGCGCAATCGCGATTCACAGATGATAGAGATAGGACTGGGATTGGAGGCATCGAACAGACCGTTCATTTGGGTGATCAAGGAACGTGAGAACTCGTCGGAGGTGAACAAATGGTTGTCCGAGGGGTTCGAGGAGAGGATCAGGGAGAGGGGTCTTATAATAAAAGGTTGGGCACCGCAACTGATGATTCTGTCACACCCTGCTGTTGGAGGATTCTTAACACACTGCGGTTGGAATTCGATACTGGAGGCCGTGTCTGTTGGCTTGCCAGTGATCACGTGGCCACACTTTAGCGATCAATTTCTCAATGAGAAGTTGATGGTGGAGGTACTGAGGATTGGGGTTTCTATCGGGGCGAAGATTCCGACATACTATCTAAATCCTGAGGATAGTGAAGAGCGACTGGTGACCAAAGAAGAAGTGGAGAAAGCTGTGGCGAGTTTGATGGACGAAGGGGAGGAAGGAAAGGAGAGAAGGCAGAGGGCTAAGGTGCTTGCAATGAAGGCTAAGGAGGCAATGGAAGTAGGGGGTTCATCTTACAAGAACTTGACACACATGATCCAATATGTGTTGGGGCATCAGAATAACAACTTACCAGAATAGAGGATCTGAGAGAAAGAGAGATTCCAATGTCTTTGATTTTTACATTTAGATTTTGTACTAGTAAAAATTTGCAGCATGCAATAAGTGGAAGAGAGAATCCATTCTCTCTAGATTTTGACAATTATTGTGGGATACTCTTTCTCTTAGAAAAAATCAATGGTTTTAGCCAAACATGGAACCGT

>F01_transcript/45482 full_length_coverage=22;length=1656;num_subreads=60

GGACTTCTACGGGATCTGGATCGCCAGTTCTTCAACTTTGATCGAAGCAATCATTGCTCTCTCTAGCTCTGCAGCCTGAAGCCAAGAGGCTCCCTTCCGATGGATCTTCACGTCTTCTTCTTGCCCTTTCTTGCTTCGGGACACATGATCCCCATGGTGGATATCGCCCGCCTCCTCAGTGACCGCGGCGTCAAATCCACCATCGTCACCACCGCCGGGAACATCCCACGAATCCAGCCCACCATCCAACGCTTCAACTCCATTTGTTCCTCCGACAAACCTCCAATCGAACTCCTCACCATCCCCTTCCCCTCCTCCGTCCCCGACAACATCTCCCTCCTCCCGACACCCGACCTCACCCCCGAGTTCGCCGCCGCAATCTCCAACCTCCGCCAACCCTTCGCCAAACTCCTCGACGCCCACCGCCCCGACTTCATCGTCACCGACCTCTTCTACCCATGGTCCGCCGATCTCGGCTACACCAGAATCGAATTCCACGTCTCCAGCAGCTTCTCGTCCGTCGTCCCCGGCTTCATCTCCCGCCAGAAGCTGCTTGAATCTGTTTCTGGCGATTTGGACCCCTTCGTCGTGACCGGACTTCCTCACAGGGTCGAGATGTCCCGGTCTCAGCTCCCCACGTTCGTTACCGATCCCGAAGAGGTGATAGCGCAGATAGTTGCAGCTCAGTCGAAGAGCTATGGCATGGTGATGAACAGCTTCTACGAGCTGGAGGCCGAGTACGTCAACATCCTGAAGAGCGCTGCTCCCATAAAGGTCTGGCATGTCGGTCCGGTATGGATGTCCGGAAGCGAGACGGTCGATCCATCAGTGAGAGGAAACAAATTGACTATCGAGTCAGGCATAATTCGTAGCTGGCTCGATAGCAAGAGCCCTGGCTCAGTTCTTTACGTCTGTTTCGGGAGTTTGGGGAAGTTCACAACCTCCCATCTGCATGAGATCGCGTCGGGTCTCGAGGATTCGGGGCACCCGTTTATATGGGCGGTAAGTAAGAATGGTGGAGATCTCCCCAAAGGATTCGAGGAGAGAATGGCGGCAGTGGGGAAGGGGATGGTGATAAGGGGTTGGGTGCCGCAAGTGATGATTTTGAACCATCTGGCGGTGGGTGGGTTCATGACGCACTGCGGGTGGAACTCGTGCTTGGAGGGAGCGAGCGCAGGGGTGCCGATGATCACGTGGCCAATGTTTTCAGAGCAGTTTTTTAACGAGAAGCTTATGGTTGATGTGTTGGAGATCGGGATTGCCATTGGGGTGAAGATTTGCAGCGCAAATGAGGAGAAGAGGACGCCGGTGAAGGGAGAGGATGTGACCAAGGCGGTGAGGGGATTGATGGGTGAAGGTGAGAATGTCGAGGGGAGGAGAAGGAGGGCGAGGGAGGTCGGAGAGAAGGCAAGGAGGGCAGTGGAGGAGGGTGGGTCGTCGTACCGTGAAATGGATCGTCTCATAGAGGATCTCATCAATTTGAAGGCGGCTAAGCTTGGGTCCGGGTCTAGCAAGGACCAGGTCTGATTTGCATTTGTTTGGGTCGAACCCTTTTTATTTTATTGGGCGGGCGCGAGGTTTGTATCATCTTAATTTGAGCCTCTTATTTACTTTACGAGGAATGCAATAAATAATAAGTTTGTGTCATCCACGTCC

>F01_transcript/45842 full_length_coverage=2;length=1674;num_subreads=60

ACCCACTCTCTCGCCCTTATCCTGCGCCGCTGCGCGTCTCCTCCCTGCTAACGTGCGCCCCCACCCCCATGGCGCCGCACGTGGCGATGTTGGCGTTCCCGTTCGGCACGCACGCGGCGCCACTCTACTCCCTGGCTCGGAGCCTCGCCGCCGCTGCACCGGGCGCCGTCTTCTCCTTCCTCAACTCGGCGAGGTCCAACGCCGCCCTCGCGCGCGCCTTCTCCGGCTCCGCGCCGCCGAACCTGCGGTCATACGACGTGGACGACGGCGGGTCGGCGGTGGCAGCGGCGGGGCCGGAGGAGGAGGTAGGGTTGTTCCTTGGAGTGTCGCCGGAGAACTTCGAGGCGGCGTTGGAAATGGCGGTGGAGGGAAGGGAGAAGGTGAGCTGCCTTGTGAGCGACGCCTTCCTTTGGTTCGCGAGGGCGATGGCGGAGAAGATGGGGGCGCCGTGGGTCACCCTCTGGACCGGCGGGCCCGTCAGCCTCGTCGCGCACCTGTACACCGACGAGCTCCGACGAACCAACGGCATCGGGGAACCAGATGGGCACCTGGACTTCATCCCCTACATGTCTCCATTGCGGGTTCAGGACCTACCTGAAGGGGTCGTCTCGGGCAATCTGGAATCGGTCTTCGCTCGCATGCTCCACCGCATGGCTGACGAGCTCCCCAATGCCGCCGCCGTCGCCCTCAACACTTTCCGCGGCCTTGATCCCGACCTCGACCATGAGTTCGAGTCCAAATTCAAGCGCGCTCTCTATGTTGGCCCATTGAACCTCCTCGTCCCGCAGCCGGCCGGGCAGGACGATCACAACTGCGTCCCCTGGCTCGACTCGCAGGAGCCAGCCACCGTGGCTTACATCAGCTTCGGCACGCTTATTTGCCCTCCGCCAATGGAGCTGGCGGAGCTAGCGGCGGGGCTGGAGGCTAGCGGTGCACCATTCCTCTGGTCGCTGAAGGATGGGGCGAGGGAGCACCTACCCGCGGGGTTCTTGGACAGGACCAAGGACCGAGGGGTCGTGGTGCCATGGGCGCCCCAGGTGCTTGTACTGGGACACAAGGCGGCCGGTGTCTTCGTGACGCACTGCGGGTGGAACTCAGTGATGGAGAGCGTGACGGCCGGAGTGCCAATGCTGTGCCGCCCATTCCTCGGGGACCAGAGGCTGAATGCGGGAGTGGTGTCGCACGTGTGGAGGATCGGGGCTGGGTTTGAGGCCGGGGTGGTGACACGCGATGCTATGTCGACTGCTCTGAGCTCGGTGCTGCAGGGGGAAGAAGGGAAGAGGATGAGGGCGAGGATCAGGGACATCAGGGAGAAGGCCACACTTGCTGTGGCCCCCCAAGGAGGATCGACGCATGATTTTACTTCTCTGTTGGAGATTGTTTCTGGGTGCTACATCCGGGAGAAGGCGACACTTGCGGTTGCCCCCGAAGGAGGCTCGACGCATGATTTCATTTCTCAGTTGGAGATTGTTTCTGGGTGCTGAAAAGATCAAGATGGTATTTTCTAGAGTCCTTCAAAGGTCTTTATTATTGTCAAGAGTATTTATTGTTATGCCACTAGAGTAATATTTTGTTGTTAGTTTTTTTATTATGTCTTAATGTAATGTTTTACATAATGTTGACTTTTGAACCCAAATAAGATATAACAAATCCTAACGGGATTCTCCTCGTGTG

>F01_transcript/46036 full_length_coverage=6;length=1633;num_subreads=60

GGGAAATACTTGCCGTATATCAAAGTGAAGATCTCTCTCTAGCTGACGGGGTAATTTTCCGAATCCAATGGGCTCCGTTCCGATAGATCTTCACGTCTTCTTTTTGCCCTTTCTTGCTTCGGGACACATGATCCCCATGGTGGACATCGCTCGCCTCCTCGCCGACCGAGGCGTCAAATGCTACCGTCGTCACCACCGCGGGGAACATCCCCCGAATCCAGCCCACCATCCAACATTTCAACTCCATTTGTTCCTCCGACAAACCTCCAATCGAACTCCTCACCATCCCCTTCCCCTCCTCCGTCCCCGACAACGTTTCCGGCCTCCCGACCCCCGACCTCACTCCCGAATTCGCCGCCGCCATCTGTGACCTCCGCCAACCCTTTGTCGAGCTCCTCGAAATCCCACCGCCCCGATTGCATCATCTCCGACATCTTCTACCCGTGGACCAACGACCTCGGATACCCGAGGATTGCTTTCCACGGCATCGGCTTCTTCTCCTGCGTCGTTCCCGGCACCCTCGCATATCAAAAGCTACACGAATCCGTTACCGCCGATGATGAACCATTCGTCGTGGGTGGCCTCCCGCATAGTATCGAGATGACCCGCTCCCAGCTCCCGGGCTTCTTCATGTCGCCTGACAGCTTGTTGATATCGGGTGTCGTCGACTGGCATCACAATTGCTACGGGGTTGTGATGAACAGCTTCTACGAGCTGGAGCCCGAGTACGCCGACATCATGAAGATGAACGCCAACTTTAAGATCTGGCATATCGGGCCTGTGTCACTGTCTGGCAGCAACACTGCCGGTCGACTGGTCGAATCCGACCTTATTGGTAGCTGGCTCAATGGCAAAAACCCGAGCTCGGTTCTTTACGTCTGCTTCGGGAGCTCGGGGAAGTTCACGACGCCCCAGCTCCGTGAGATCGCATCCGGCCTCGAGGATTCCGGCCACCCGTTTATATGGGCGGTGAACAAGAGCGACGAAAATCTGCCCGAGGGGTTCGAGGAGAGGGTGAAGGGAAAAGGACTGGTGATAAAAGGGTGGGCGCCGCAGGTGATGATATTGAACCACCAGGCTGTCGGTGGGTTCATGACGCACTGCGGCTGGAATTCGTGTTTGGAGGGAGCAAGCGCCGGGTTGCCGATGATCACGTGGCCGATGTTTGGTGACCAGTTCGTCAACGAGAGACTGATTGTCGATGTCGTGGGGATGGGGATAGGAATCGGGACGAAGGTTTGTAGCGCACATGAGGAAGAGAGGACGGTGGTGAAGGGGGAGGATGTGGCCAAGGCGGTGAGGGGATTGATGGGCGGCGATGAGGCCGAGAGGAGGAGGAAGAGGGCTAGGGAGGTCAGAGAGAAGGCGAGGAGAGCAGTGGAGGAGGGCGGCTCGTCGTACAGCGAAACGGACCGTCTCGTAGAGGATATAATCAATTTGAAAACGGCTCGGGTCCAGTTCGATCACCCCAAATAAGACTTAGATTTTGTGTGGATCCATTATCCTCTGCTAAATTTATTGGGTTTGTATTTCTGCCATTTATCAGCTCACTCACTGTAGTCATGTGTTGTTTTCATTGGGATTGGATCTATGCCATATATAGCTCTACGATTGAGGTTTTTAGTCTTTCTTC

>F01_transcript/46097 full_length_coverage=2;length=1677;num_subreads=60

GATCTCTCTCTCTCTCTCTCTCTCTCTCTCTCTCTCTCTCTCTCTCTCTCACTCTCTCTCTCTGGTCATGGATGGAGAAAAGCTACACATTGTTGTGTTCCCATGGCTAGCCTTTGGGCACATGTTGCCCTATCTTGAGCTCTCCAAATCTTTGGCCGCAAGGGGCCACCAAATCTCCTTCATCTCTACCACAAGAAACATAAAAAAGCTCCAGCCCAAAGTTCCACCCACGCTCTCCCAACTCATTCAATTCATCGCTCTACCTCTCCCTCCGGTCGACGGCTTACCGGCTGCCGCCGAGTCCGTCTTCGACATCCCTCTAAACCTCAGCCGTCACCTCGAGAAGGCGTTCGACGGCCTCGAGCGCCCCTTGGCCCAATTCCTCAGTTCGTCGTCCCCAAAGCCTGATTGGATCATCGTGGACTATGCCTCCGACTGGCTCCCGTCGCTCGCTTCCAAATTCAATGTGCCATGTGTCCACTTCTGCATCTTGATGGCCCCAGCCATGGCGTTCATGGGACCCGTGTCAGAAGTGGACAACATAGACTCCATCACGCTTGAAAAAGTGGTGGTTCCGCGCGAATGGATGACATTTCCTACAAATGTAGCCTTCCGCCCTTATGAGGCTCGACAGGTCCTTGATCTCTTAAGTTCAGACAATGCTTTGGATGCCTACCGCAACTTGCTCACCATCGGAGGTTGCAAGGCCGTGGCTCTCCGAGGCTGCAACGAGTTCATGCCGAAGTGGGTGCCCCTCATTCGAGATCTCTACAAGAAACCCGTCATCCAAGTTGGCATGCTTTCCCCGATGCCTGTCGACAATGGAGATTATGATACCAGCGAACTCAAGATCATGGAGTGGTTGGATCGACGATCGCCGAGTTCTGTCATATATGTAGCCTTTGGGAGCCAAGCCGACCTCAGCATCGAGCTCTTACACGAGCTAGCGCTTGGGCTTGAGCTCTCTAACTTGTCTTTTCTTTGGGCTCTTAGAAAGTCGACCACTAGCGAAAATAAAGTGATTTTGCCCGTAGGGTTCCAAGAACGGACAAAGGATCGAGGATTTGTCATCATGGGTTGGGTTTCCCAGCTCAAGGTCTTGGCACACGAGTCGGTGGGAGGATTCATGACGCACTGCGGTTTCAGTTCGATCATCGAGAGCCTCCAATTTGGTCGTCCTCTAATTCTTTTCCCTTTGTTCGGGGATCAAGGGCTCAATGCTCGGGCGATGGTGGAGGAGAAGATCGGATTTGAGGTGAGGCGGGGCGAGGAGGATGGGTCACTAACAAAGGAAGCAGTGGCAGAGGCATTGAGGCTGGTTGTCGTCGAGGCAGAGGGGGAACCTTTCAGGAAAAAAGCTAAAGAGATGATGGAGGTTTTTGCGGACAAGAATTTAAATGAGAGATATATGGATGATTTCACTCAGTATCTAATGGACCATAAAGATAGCAAATGAAAGTAGCTCGATTGAGGCGTCAGGCAAGAGGAAGGGCGGTCCAAAGTAGTGCCACTGCTAAGGGCATCTCCTTTTATTTCTGGGATGAAGAGAATCTATATCTAGTGTGTGTGGTATGTGTTAGGGACACCAGCTTATAATATCTAAAAATGTGATGTAATTTAAGAAAAATAAAGTTGTAAGATGATGTAATAAGAGGAACAATATAATATTGCTTTTAG

>F01_transcript/46105 full_length_coverage=4;length=1660;num_subreads=60

GGAACAACATAACATTCACAATCTCTCAGCAATCACTGCTGTCAATCTCCTGTTGACTCACTCCTCCCCCATGGGTTCCGAATCTCAGCCAATACTCCACGTCCTCTTCTTCCCCATGATGGCCGCCGGCCACATGATCCCCATGGTGGACATGGCCCGCCTCTTTGCGGCCCGTGGCGTCAAGGCCACCATCGTCATCACCCCCGCCAACTCCGACGCCATCCGATCCACCGTCGACCGCGCCAAAACCCCAAATATAGAATTCCTCATCGTCCCCTTTCCCTTCGAGAACCTCTCCGCCGTCTCCTCCCTGGAGTTCAGGGCCAAGTTCCTCCAGGCGATGGAGATGCTCCGCGACCCCTTCGACCGAATCCTCGCTGACCACCTCCCCGACTGCGTCATCACCGACATGTTCTTCCCGTGGTCCGCCGACGTGGCCGCCAAACACAACGTACCGAGGCTCGTGTTCCACGGGTCGTGCTGCTTCTCGCATTGCGCCATGGACGCGATAGTTCGTTACAACATAGTGGAGAGCTCGCCAACCGATGTCGAATCCTTTGTGGTTCCGGGTTTACCGCATAGGATCGGGATGCTCAAGTCGCAGATCCCGGATCATAATAAATCCATGCCCGCCCTCGGGGAACTGCTTCGTAGGACGAAGGAGTCGGAGGAGCGGAGCTACGGCGTGGTAGTGAACAGCTTCTACGAGCTCGAGCCCGACTATGTTCGGCACTACCGCGAGGTGATGGGGCGGAGAGCGTGGCTGGTGGGACCGGTGTCACTCTCCAACGCCAACGAGGCCGACATGTTGGGGAGAGGGGGTAACAAGATCTCCTCCCATGCTTGCTTGGATTGGCTTGACGGTAAAGAACCGGGCTCGGTATTGTACGTCTGTTTTGGGAGTCTTGGCGACTTCACTGCTGCCCAGCTCAACGAGCTCGCGGTTGGGCTCGACGCGTCGGGCGTTCCGTTTGTTTGGGTGGTGAGGAAGGGCGCGACAGAGCTTTTGCCAGCGAGTTATGAGGAGAGGGTGGAGGGAAGGGGGATGGTGGTGGAGGGGTGGGCTCCGCAGATATCAATACTGAACCACCGGGCGGTGGGGGGGTTCCTGACGCACTGCGGGTGGAACTCCAGCATCGAGGGGATCTCCGCCGGGCTGCCGATGGCGACTTGGCCGCTGTACGCGGAGCAGTTCTACAACGAGCGGCTACTGGTCGAGGTGTTGGAGGTCGGTGTGGCAGTGGGGATCGAGAAGTGCATAATGCTTCCCGAAGACAGGCCGGTGGTGGGGGCGGCCTCTGTCGAGAGGGCTGTGGGGAGGCTGATGGGAGGCGGGGAGGAGACGGAGAGGCGGAGACAGCGGGCGAGGGAGTTTGGTGATATGGCAAAGGCGGCGGTGGCGGACGGCGGATCATCCTTCGTGGAGATGGGCAATCTGATCAAGGAGTTGACCGGCATGCGGAATTTGACTACGAAAACTGAAGCCTGACTCATGTTCACAATGTAATAATAACTGTTAATTAAGCTCTCTGCTGACAGTGAGCTTTATCTCCGCTTTAGTTTTGACCAACATGCTAAATTTGTCTTGGCTTTTGTATTTTTCAATGTTGCACAATTAAATGTCAGGGGTTCTCCATGAGTGGGGTACACAATTAATCAT

>F01_transcript/46190 full_length_coverage=9;length=1658;num_subreads=60

GGATCATCATATGAATCAGAGCCCCTTGTTATCTGAACCTCATCATATCCTCGTCCACAAAGATGGGGGAGCAACCACTGAATACCATGAACGCTCCGCACATAGTGATGATCCCATTGATGGCCCAAGGCCACATGATTCCCATGGTGGACATTGCCGTCCTCCTCGCAGGCCGCGGGGTCACTGTCACCTTTGTGGTGACCCCCCTTAATGCTGCCCGAATCCAGCACACCATCGATCGGTCCCGAGAATTCGGCCTCCCGATCGACTTCCTCATCCTCCCATTCCCCTGTGCGGCCGTCGGCCTCCCCGAAGGCTACGAGAATGCCGACACGCTTCCCTCCCACGACCTCATGCCCCAATTCTTCGCCGCCGCCGCCCTCTTGCGTCCCCCCCTTACCCTCCACCTGCGGTCGCACGTCCCCCTCGTGTGTCATCTCCGACATGTGCCACCCTTGGACCCACGAGCTTGCCCGCGAGTTCGAAGTCCCCAGGCTTGTCTTTCACGGCTTCGGACTCTTTGCACTCCTTTGCTTCTATAACATCCGGCACAATGACGACGCCATCAAGTCGAGACTAGAAGAGGAGGATGTCAATAGGCCCTTTGTGGTGCCTGGATTGCCTGATCGGATCGAGCTCACTAGGGCCCAGGTCCCCGGGTTCAAGGCCCCTCCGAACATGCAGTTGTTCAGGGAGGAGATTTTTCGGACCGAGGAAGCGGCCGATGGGGTCGTGGTGAACAGCTCTGACGATCTCGAGCCGCCCTACAAAGAATGGTACGAGAAAACTATAGGGAAGAAGGTGTGGACAATTGGGCCGATGTCAATGTACAACAAAAACACCTCAGAGATGGCACTGAGGGGGAACAAGGCTGCCATCGACAAGGACCGGTGCCTAAGCTGGCTCGACTCCATGACTCCTAGCTCCGTCCTTTATGTCAGCTTCGGGAGCATCGTGCGCACCACAATCACGCAGCTGATCGAGACCGGTCTCGGCCTGGAGGCTTCGAGCCACCCCTTCATCTGGGTGATCAAAGGCAGCGACAAAATGTCGGAGGTGGAGGAATGGTTGAAGGAGTTCGAGTCGAGGACGAGCGAGAGGGGACTGATAATCAAGGGGTGGGCACCGCAGGTGATGATACTGTCGCACCAGGCAGTGGGAGGGTTTATGACGCACTGCGGTTGGAACTCGACGCTGGAGGGAATGTCGGCGGGGGTGCCTATGATAACATGGCCCCACTTTGCGGAGCAGTTCCTGAACGAGAGGTTCGTGGTCGACGTTGCCAAGGTCGGGGTTCCAGTGGGGGTGAAGTCGCCTACCACGTGGGGGTTGGATACGGGCGAGGTGATGGTGAAGAGGGAAGGAGTCGAGAGGGCAGTGAGGGTGTTGATGGACGGAGGAAATGAGGGAGAAGAGAGAAGGGCGAGGGCAAAGGCGCTGGAGGCGAAGGCGAGAAAGACGATCGAGGAGGGGGGATCGTCTTTCCGTAATACTACACTACTGATTCAGCACATAGCTGCTATGGCAACAAAGATTGTATAGAGGAAAAGCAACACTTTTTGGCCTCTTAAACTATAAATTTTTAATATTTCATGTTTTCATGGATTATGTTATAAAGTGGCAGTTAACAAATTGATTGTCCACTTTTGATCGGTCAC

>F01_transcript/46229 full_length_coverage=4;length=1670;num_subreads=60

GAGAGTGGTGATCAGAAAGAGAAGGAGAGTTTGAAGGATGGTGAAACCCGGAGTTGTGTTCATCCCCTGTCCGGCCATGGGCCACTTCATCTCAGCGGTGGAGTTCGCGAAGAAGCTCGAGAGCCGCTTCGCCGTCACCATCCTCCAAATTGCCCATATTCCCCCGGCATGGTCTTCCGCCATCAAGGCTTATGTGGAGTCGGTCGTGTCCTCCGGCTTCGACATCCGATTCGAAGAGCTCCCTCACATTGTTCCTCCTCATGCGGAGAAGCAAAGACCAGAGACCTTCGCCTCTCGTTTAATGGAGGTCAACAAATCGCCGGCAAGAGACGCCATCTCTCTGATTCCTAACAATAATATCGCGGCAATAATTCTCGATCTGTTCGGCAGCTCCATGGTGGACGTCGCCGATGAGCTCGGCATCCCGGCCTACATCTACTTCACCTCGAACGCGGCCTTGCTCGGGTCCATGCTCTACCTCCCCACCCTCCATGTAAAGGTCCCGTGCGAATTGATGGAGTTCGGAGGGCACATCTCGATGCCCGGACTCCCTCCCGTGCCGCCCCATTGCATGCCATCGTTCGCGATGGACAAGAAGGACGAGGCCTACACCACGTTCATCAACCACGGATTGTGCTTTCGAAAGGCTAAAGGCCTAATCATTAACACATTTGAAGATATCGAGCCGAAGACCCTGCAAGCATTGGCGGACGGCGAGTACCTCCCAGACCACACTATGCCACCGGTCTTTCCGTCGGACCGGTGCTGGCGATCAAGCCGAAGAATGATCAGCCGCATGATTGCATCGCATGGCTCGACAAGCAACCTGTGGCTTCGGTGGTGTTCTTGTGCTTTGGAAGCATGGGATGCTTCGAAGCACCCCAGGTGAAGGAGATGGCGGTGGGGCTGGAACGGAGCGGCCACCGGTTCTTGTGGGCTCTTCGTAGCCCTAACACCACCGATAGCTTCCGGTTTCCGACCGACTCCGATCTCGACGAGGTGCTGCCCGAAGGATTCTTGGAGAGGACTGAAGGGAGGGGGCTGGTGTTGCCGTCTTGGGTTCCGCAAACTGAGATACTGTCCCACGAGGCGGTGGGGGGATTCGTCACGCACTGCGGGTGGAACTCAGTGATGGAGGCGGTGTGGCTCGGGGTGCCGATGCTCGGGTGGCCGCTATTTGGGGACCAGCACTTGAATTGTTTGGCGATGGTGAAACAGTCGGGCGTGGCTCTCGAGCTCAAGTTGGACTACAACAATGGCGGGTTCGTGAGTTCGGGGGAGCTGGAAAAGGGAGTTAGGTCCCTGATGGGGGACCTCCAGGATTCGAGGAATGTGAGAGGTAAGGTGAAGCAGATGAAAGTCGACAGCCGGAGAACCGTGATGGAAGGGGGTTCATCCAATGTCGCATTGGAACAGTTGATTTGTGAATTAAACAAGAAGGTGGCCGTTGCCGTTACTGTTGATAAGAATAATAATTGCTATGTGACTCCTAAGAAGATTGTTGCTACGTGATGAGCTATGTTCGTTGGACAGCTCATGCTTCAAGTCCAACTTAAAATATTTTCTAATAACATATTATCTTTCCATTGCCATGTTAAAAAGAATACTCCTCTAAATCTTTTTCTTTTTTATTACAATTTATAAGTGGTCCAAGATGGAATCTTCATTAC

>F01_transcript/46554 full_length_coverage=13;length=1648;num_subreads=60

GGACCAAACAAACACATCCAAAAATGGACCAACCGAACAATCATGGATGCCATTGGCCATCCCCGTCCTCACTTCGTGTTGGTCCCTCTAATGGCCCAAGGCCACATGATCCCCATGGTGGACATGGCCTACCTCCTCGCCGACCGTGGTTCCCTCGTCAGTTTCATCACCACTCCTGTTAACGCTGCACGAATCAAGTCCACCATCGATCGTGTCAAGGACTCTCGCCTCCCAATCCGATTCATCGAGCTACCATTCCGCTCCTCTGAAGCTGGATTGCCCGAAGGATGTGAGAATGTTGACCTCGTTGAATCTCTCGGCTTCCTCAAATCCTTCTTTGAGGGCACACACATTCTCTCAGAACCCTTAGAGCTCTATCTCCAAAAACAACAGCACAGCCCAATCTGCATGATCTCCGATTTTAGCCTCCCGTGGACCGCAGAGCTCGCTCGGAAGCTTAATATACCCAGGTTGGTATTCCATGGACCCTCATGCTTCTACCTTCTCAGTGTTCATAATATTCAGAAACATAGAATCTCCGACCGCATCACTCATCAATTTGAGCCATTCCTTGTGCCCGACTTGCCACAACAGTGGTCCATAGTGGTCACGAGAGTACAATCACTGGGGTTCTTTGAGATTCCAGGATGGGAGGAGTTCCATAAGAAGATACTAGAGTCGGAAGCGGCAGCGGAAGGAGTTGTCATGAATTCATTCGAAGAATTGGAGGATTGTTTCATTGAGAGCTATCGAAAGGTTATGGAGAAGAAGGTTTGGACAATTGGGCCTTTGTGTCTCCACAACAAGGATAACTCCTATAAGGCCGCGAGAGGGAATAAAGCTGTTGTTGACGAACATCGAGTTCTCAATTGGCTCGATTCTATGAGCAGAGATTCAGTTATCTACATTAGCTTTGGTAGTCTAGCGCGCAATCGCGATTCACAGATGATAGAGATAGGACTGGGATTGGAGGCATCGAACAGACCGTTCATTTGGGTGATCAAGGAACGTGAGAACTCGTCGGAGGTGAACAAATGGTTGTCCGAGGGGTTCGAGGAGAGGATCAGGGAGAGGGGTCTTATAATAAAAGGTTGGGCACCGCAACTGATGATTCTGTCACACCCTGCTGTTGGAGGATTCTTAACACACTGCGGTTGGAATTCGATACTGGAGGCCGTGTCTGTTGGCTTGCCAGTGATCACGTGGCCACACTTCAGCGATCAATTTCTCAATGAGAAGTTGATGGTGGAGGTACTGAGGATTGGGGTTTCTATCGGGGCGAAGATTCCGACATACTATCTAAATCCTGAGGATAGTGAAGAGCGACTGGTGACCAAAGAAGAAGTGGAGAAAGCTGTGGCGAGTTTGATGGACGAAGGGGAGGAAGGAAAGGAGAGAAGGCAGAGGGCTAAGGTGCTTGCGATGAAGGCTAAGGAGGCAATGGAAGTAGGGGGTTCATCTTACAAGAACTTGACACACATGATCCAATATGTGTTGGGGCATCAGAATAACAACTTACCAGAATAGAGGATCTGAGAGAAAGAGAGATTCCAATGTCTTTGATTTTTGCATTTAGATTTTGTAGCTAGTAAAAATTTGCAGCATGCAATAAGTGGAAGAGAGAATCCATTCTCTCTAGATTTTGACG

>F01_transcript/46557 full_length_coverage=7;length=1659;num_subreads=60

GGAGACACGAGAATCATTAAACAAAGGTTGGACTATGGGGTCACAAACTCAACCCCTCCACATGCTCTTCTTCCCTTTCATGGCCGCCAGCCACATGATCCCAATGGTCGACATGGCGAAGCTCTTTGCCGCACGAGGAGTGAAGGTAACCATCCTCACAACTCCATCAAACGCTGAAACAATAAAATCCGCCATTGAACCCGGCCACCAAGCCATCGCCCTCGCGCTCATCCCCTTCCCCTCGGTGGCCGTCGGCTTGCCAGAAGGCTGCGAGAACCTCTCCGCTGTGCCCACTATCGAGCTACAGAACAGATTCTGCAGAGCACTGGCAATGCTCCGCCAGCCGTTCGATCGAGTCCTCATCGACCTCCTCCCTGACTGCGTCGTCTCCGATTCGTTCCTCCCGTGGACCTGCGAAATCGCCGCCAATTATAATGTGCCGAGGCTCGTGTTTCACGGCACGAGCCACTTCTCTATGTGCATCTCGGACAGCTTCGATCGTCACGTGCCGGTCGTGAGCTTGGCACCGGATGGCCACGAGTGTTTGATAGCCCTGGGTCTACCTCACCGGATCGAGATGCCCCGGTCCCAGATGCCCGACCTCACCGGATCCGGATCCGATTTCCCGCAACTTCTGGCAGAGGCACAGGAGGCGAATCTGAAGAGCTACGGCATGGTGATGAATAGCTTCTACGAGCTGGAGCCGGAATACGTCAGTCACTTCCAAAAAGTCCTCGACCGGAAATCGTGGAATGTGGGTCCCGTGTCCCTATACAACAAAGACACAATAGAGATGCGATCGAGGGGACACACAAGTGTAGTCGACGTCAACAATTGCTTGAAATGGCTCGACGGCAAGAGCTCGGGCTCGGTCGTGTACATCGCATTCGGTACTCTAGGCGAGATGGGGATGGATCAAGTCCGCGAGATAGCTCTCGGGCTCGAGGCTTCGGGCCGACCTTTTATATGGGTGGTGAGGAATGTTGATTGGCTGCCGGAGGATTACGAGGACATGGTCGGGGATCGGGGGATGGTGATAAGGGGTTGGGCACCGCAGGTGCTGATCCTAAGCCATCAGGCGGTGGGGGGAATGGTGACACACTGCGGGTGGAACACGTGCTTGGAAGGGGTGTCGGCCGGGGTGCCGATGGTGACGTGGCCGTTCTTTGTGGAGCATTTCTTTAACGAGCGGTTGATCGTGGAGTTGCTGGGGATCGGGGTGAGCTTGGGGGTGAAGGAGTGTGCGATAAAGAAAGAGGAGATGCCGGTGGTGGATTCGGCGAAGATAGAAAGGGCGGTGAATCGGCTGATGAGTGAAGGAGACGAGGCGGAGGTAAGGAGGAGGAGGGCGAGGGAGCTCGGAGAGAAGGCAAAGAGGGCAATAGAGGAGGGTGGCTCGTCGTACATTGATGTTGGGAATTTAATCGCTGAGTTGAGTTGCAGGCCGAGACCAGTCCACAACAAGATACCAGCAGTTTCACCCTAAAGCACACATAAGCATTATGTATCTAAAGCGCAATCTCTACAACAGAACATGCTTTTAACCGCCCACAAAAGTATGCCCTAAATAATTTTTCATGTACTCAATGTTATAAAAAGTCCTTACCTATTAATGGCATGCTTAGCTCACGTTTAATAATTTAAATAAACGTCGTGACT

>F01_transcript/46696 full_length_coverage=4;length=1661;num_subreads=60

GGACTTGGTGAGTAAGCAGTGTCGCCTTGCTTATGGACCTCTGTAGCTCTCAGTATATTTCATCTCTATATACTAGCACAACTCCGGAGACACGAGAATCATTAAACAAAGGTTGGACTATGGCGTCACAAACTCAACCCCTTCACATGCTCTTCTTCCCTTTCATGACCGCCAGCCACATGATCCCAATGGTGGACATGGCGAAGCTCTTTGCCGCACGAGGAGTGAAGGTAACCATCCTCACAACTCCATCAAACGCTGAAACAATAAAATCCGCCATTGAATCCGGCCACCAAGCCATCGCCCTCGCGCTCATCCCCTTCCCCTCGGTGGCCGTCGGCTTGCCAGAAGGCTGCGAGAACCTCTCCGCTGTGCCCACTATCGAGCTACAGAACAGATTCTGCAGAGCACTGGCAATGCTCCGCCAGCCGTTCGATCGAGTCCTCATCGACCTCCTCCCTGACTGCGTCGTCTCCGATTCGTTCCTCCCGTGGACCTACGAAATCGCCGCCAAATATAATGTGCCGAGGCTCGTGTTTCACGGCACGAGCCACCTCTCTATTTGCATCTCGGACAGCTTCGCTCGTCACGTGCCGGTCGTGAGCTTGGCACCGGATGGCCACGAGTGTTTGATAGCCCCGGGTCTACCTCACCGGATCAAGATGCTCCGGTCCCAGATGCCCGACCTCACCGGTTCCGGATCCGATTTACCGCAACTTCTGGCAGAGGAGGCGAATCTGAAGAGCTACGGCCTGGTGATGAATAGTTTCTACGAGCTGGAGCCGGAATACGTCCGTCACTTCCAAAAAGTCCTCGACCGAAAATCGTGGAATGTGGGTCCTGTGTCCCTATACAACAAAGACACAATAGAGATGCGATTGAGGGGACACACAAGTGTAGTCGACGTCAACAATTGCTTGGAATGGCTCGACGGCAAGAGCTCGGGCTCGGTTGTGTACATCGCATTCGGAACCCTAGGCGAGATGGGGATGGATCAAGTCCGCGAGATAGCTCTCGGGCTCGAGGCTTCAGGCCGACCTTTTATATGGGTGGTGAGGAATGTTGATTGGCTGCCGGAGGATTACGAGGACATGGTCGGGGATCGGGGGATGCTGATAAGGGGTTGGGCACCGCAGGTGCTGATCCTAAGCCATCAGGCGGTGGGGGGAATGGTGACACACTGCGGGTGGAACACGTGCTTGGAAGGGGTATCGGCCGGGGTGCCGATGGTCACGTGGCCGTTCTTTGAGGAGCATTTCTTTAACGAGTTGTTGATCGTGGAGTTGCTGGGGATCGGGATAAGCTTGGGGGTGAAGGAGTGTGCGATAAAGAAAGAGAGGCCGGTGGTGGATTCGGCGAAGATAGAAAGGGCGGTGAATCAGCTGATGAGTGAAGGAGACGAAGCGGAGGTAAGGAGGAGGAGGGCGAGGGAGCTCGGAGAGAAGGCAAAGAGGGCAATAGATGAGGGTGGCTCGTCGTACATTGATGTTGGGAATTTAATTGCTGAGTTGAGTTGCAGGCCGAGACCAGTCCACAACAAGATACCAGCAGTTTCATCCTAAAGCACACATAAGCATTATGTATCTAAAGCGCAATCTCTACAACAGAACATGCTTTTAACTGCCCACAAAAGTATGCCCTAAATAATTTTTCATGTACTC

>F01_transcript/46707 full_length_coverage=32;length=1644;num_subreads=60

GGAATCAGAGCCCCTTGATATCTGAACCTTATCATATCCTCCTCCACAAAGATGGGAGAGCAACCACTGAATGCCATGAACCCTCCCCACATAGTGATGATCCCACTGATGGCCCAAGGCCACATGATTCCCATGGTGGACATTGCCGTCCTCCTCGCAGGCCGCGGGGTCACTGTCACCTTTGTGGTGACCCCCCTCAATGCTGCCCGAATCCAGCCCACCATTGACCGGTCCCGAGAATTCGGCCTCCCGATCGAATTCCTCGTCCTCCCATTCCCCTGTGCGGCCGTCGGCCTCCCCGAAGGCTACGAGAATGCCGACACGCTTCCCTCCCGTGACCTCGCACCCCAATTCTTCGCTGCCGCCGCCCTCTTGCGTCCCCCCCTTACCCTCCACCTGCGGTCGCACGCCCCCTCGTGTGTCATCTCCGACATGTGCCACCCTTGGACCCACGAGCTCGCCCGCGAGTTCCAAGTCCCCAGGCTCGTCTTTCACGGCTTCGGACTCTTTGCACTCCTTTGCTTCTACAACATCCGGCACAATGACTGACGCCATCAAGTCGAGACTAGAAGAGGAGGATGTCAATAGGCCCTTTGTGGTGCCTGGATTGCCTGATCGGATCGAGCTCACGAGGGCCCAGGTCCCCGGGTTCATGCTCACTCCGAACATGCAGATGTTCAGGGAGGAGATGTTTCGGACCGAGGAAGCAGCCGATGGGGTTGTGGTGAACAGCTCTGACGATCTGGAGCCGCCCTACAAAGAATGGTACGAGAAAACTATAGGGAAGAAGGTGTGGACAATTGGGCCGATGTCAATGTACAACAGAAACGCCTCAGAGATGGCTCTGAGGGGGAACAAGGCTGCCATCGACAACGACCGGTGTCTAAGCTGGCTGGACTCCATGACCCCTAGCTCCGTCCTTTATGTCAGCTTCGGGAGCCTCGTGCGCACCACAATCACGCAGCTGATCGAGATCGGTCTCGGCCTGGAAGCTTCGGGCCACCCCTTCATCTGGGTGATCAAAGGCAGCGAGAAAATGTCGGAGGTGGAGGAATGGTTGAAGGAGTTCGAGTCGAGGACGAGCGAGAGGGGACTGATAATCAAGGGGTGGGCACCGCAGGTGATGATACTGTCGCACCAGGCAGTGGGGGGGTTCATGACGCACTGCGGTTGGAACTCGACGTTGGAGGGATTGTCGGCGGGGGTGCCTATGATAACATGGCCCCACTTTGCGGAGCAATTCCTGAACGAGAGGTTGGTGATCGACGTTGCCAAGGTCGGAGTTCCAGTGGGGGTGAAGTCGCCCACCACGTGGGCGTTGGATACGGGCGAGGTGATGGTGAAGAGGGAAGAAGTCGAGAGGGCAGTGAGGGTGTTGATGGACGGAGGGAAGGAGGGAGAAGAGAGGAGGGCAAGGGCAAAGGAGATGGAGGCGAAGGCGAGGAAGACGATCGAGGAGGGGGGATCGTCTTTCCGTAATACTACACTACTGATTCAGCACATAGCTGCTATGGAAACAAAGATTGTATAGAGGAAAGCAGCACTTTTTGGCCTCTTAAACTATAAATTTTTAATATTTCATGTTTTCACTGATTATGTTATAAAGTGGCAGTTAACATATTGATTGTCCACTTTTGAATAGTC

>F01_transcript/46898 full_length_coverage=3;length=1654;num_subreads=60

ATGGTGAGACACGCTAGTGGCATATTCCTCTCTCTTGGTTGTACTCTTTTGCTGATTCCAGTGTCTATTGATTCGAGCTCCATGGGCTCTCAAGCCCAACCCCTCCGCATCGTCTTCCTCCCCATGATGGCCGCCGGCCACATGATCCCCATGGTCGACACAGCTCGTCTCTTAGCCGCCCGTGGCGTCAAGTGCACCATCGTCACCACGCCCGCCAACGCCCCCGCCATGGACCGCGCCAAACATCAGAACAACATCGAGATCCTCCTCATCCCCTTCCCCTCGGCCGCCGTCGGGCTCCCCGAGGGCATGGAGAACTTATCCGCCTCCTCCTCCCCGGAGTTCAAGATCAAGTTCTTCGAAGCCATCGATATGCTCCGCCACCCGTTTGACAGAATCCTCACCGACCTCCGTCCCGACTGCATCGTCACCGACATGTTCTTCATCTGGTCTGCCGACGTCGCCGCCGAACACGGCGTGCCGAGGCTCGCCTTCTTCGTCTCCTGCTTCTTCGCGCAGTGCGCCTCGGACGCGATATATCGCCACAACGTATTGGAGAGCTCGCCGGCGGATGTGGAGTCTTTTGTGGTTCCGGGTCTGCCGGATAGGATCGAGATGCTCAAGTCCCAGATCCCGAATCCTAAGTTTGTGGTGGGCATGGAAGGGAAGATGCGACAACTGAAGGAGATGGTGGAGCGGAGCAACGGCGTTGTGGTGAACAGCTTCTACGAGCTGGAGCCGGAGTATGTCCGGCACTGCCGGGAGGTGGTGGGGTGGAGGGCGTGGCACGTGGGCCCCGTGTCGCTCTGCAATGCGGAGGAAGCCGACAAGCTCGGGAGAGGGGGGGACAAGAACAAGATCTCCTCCGAGTCTTGCTTGGAATGGCTTGACGGTAAAGAACCGGGCTCGGTGTTGTACGTTTGCTTCGGGAGCGAGGGCGAGTTCACTGCGGCTCAGCTCCACGAGCTCGCGGTGGGGATGGAGGCCTCCGGATACCCTTTTATGTGGGTGGTGAGGAACGGAGTGTTGCCGGCGGGGTTCGAGGAGAGGGTGGAGGGGAGAGGGATGGTGGTGAGGAGTTGGGCCCCGCAGATCTCCATACTGAGCCACCGGGCGACCGGGGGGTTCCTCACGCACTGCGGGTGGAACTCGAGCATGGAGGGGATAGTGGCGGGTCTGCCGATGGCCACGTGGCCGTTGGCAGCGGAGCAGTTCTACAACGAGCGGCTGCTGGTGGACGTGCTGGGGGTCGGTGTGGCGGTGGGGGTCGAGAAATTCTCGTTCACGTATGAGGACCGGCTGCTGGTGAGGTCGGCCGCAGTGGAGAGGGCGGTGGGGAGGGTGATGGCAAGCGGAGAGGAGGCGGATAACCGGCGCCGACGGGCGAGGGAGCTGAGTGATCTGGCGAAGGGGGCGGTGCTGGACGACGGATCATCGTTTGTGGAGATGGGGAATCTGATAAAGGAATTGACGGACATGCGGAATTTGGCCACTAAAAATCAATGAGTTGGCGAATTTTTTCTCTTTGGTTTGGAGTTTGTTCTCTCTTTGGTTTGGAGATTGTCTAGAAGGTTATAATTTAAGCTCTTGCAACTAAAGATGACAAAAAATGTATTTACTAAAATTATAATTTTATCGAAAATATTTGGGATTG

>F01_transcript/46915 full_length_coverage=2;length=1648;num_subreads=24

GATCACTGCTGCTTTTTGCCAATCGATCTGCGAAGCAATCGAATCGTCCTCTACCCCGCCGATCCGCCCCCATGGATCCCCTCCACGTGGTGGCGCTCCCTTACCCGGGCCGAGGCCACATCAACCCCATGATGGTCCTCTGCCGCCTCCTCGCCGACCGCGGCCTCCTCGTCACCTTCGTTGTCACCGAGGAGTGGCTCGCCCTCCTCGCCTCCTCGTCGCCGCCGCCGCCGCCGAACGTCCGGCTCCGCGCCATCCCCAACGTCATCCCTTCCGAGTCCGTGCGCGGTGCCGACTTCATCGGCTTTCTCGAAGCCGTCTACACCAAGATGCGGGAGCCCTTCGAGCGACTCCTCGACCGCCTCGAGCCGCCGCCCGTGGCGATCGTGGCGGACACATACATGCCGTGGGCGGCGGCGGCGGCGGGGCGGCGAGGGATTCCGGCGTGCTCGGTGTTCACCATGTCCGCTGCATTCTTCTCCGTCCTCTGCCAGTTCGACCGGATCGGGGACGCCTCAGAAGAAGCTGACGACCCTTTGGAGAAGTATGTCCCTGGTCTCACTTCTGTACGTCTATCCGAGTTACACTCCATCTTTGCTGGCATGGAGAACCCAAAGAAGAAAGCCATGGAAGCCATATCTTGGGCGAGGAAGGCACATTGCATCCTCTTCACTTCCTTCTACGAACTCGAGCCCTGCGTCATCGATGCACTACGATCCGAGCTCTCATGCCCTTTGTACCCAGTTGGTCCTTCCATTCCTTATATGATGCTCCAAGAGAATCCAATCAAACCTCTCCAACACGACGACAACCCTGACTATTATGAGTGGCTCGACTCTCAGCCTCAGTGCTCCGTGCTGTACATCTCATTAGGCAGCTTCCTATCGGTCTCGGCTCTACAAATGGACGAGATCGCAGCAGGCCTGATGGCGAGTGGGGCCCGGTTCTTGTGGGTCGCTCGTGAGTGCTCGTCTCGGATGCAGGAGATGAGCAGTAGAGCGGGATTAGTTTTGCCATGGTGCGACCAGTTGAGGGTCCTTTGCCATCCTTCGATCGGAGGGTTCTTGACACACTGCGGGTGGAATTCGACGCTGGAGGCTGTCTTTTCCGGGGTTCCGATGATTGTTTTCCCCATCTTTGGGATCAAGTTATGGATGGCCGGCTCGTGGTAGATGAGTGGAAGGTTGGCCTAAGAGTGAGGGAAGGGAAAGAATTTGTTGGGAGAGCAGAAATTGCCAAGGTTGTGAAGAGGTTGATGGAAGGATGAGGAGATGAGGAAGAGAAGTATGGAGCTCCGTGATGTTTGTCGTGGTGCGATCGGAGTTGGTGGATCCTTGCATAGCAATCTCGATTCTTTTGTTAGAGATCTCATTAAAGGTCATGGTTGTTAGGTAATTGAATGTGTAGCTTAATATCAAGGCGACTTTGATAGGAAGTTTATGTTATTTACAAGCTATTAGTAGCTCAAATGATGTGTTGGAATGTAGTAACTTGTTGGAGGGAAATTAATAAGCTACCTTCAATTGGTAAGATTCCCTTATTATCTCCAGAACATTTGATGGACTGCAAAACATTTGTGCCTAAGATGATATGGACAATTTATTATCTTAAGTTTTGTGTTGGAATAAAGTAAATTCTATTTTGTGGC

>F01_transcript/47116 full_length_coverage=3;length=1633;num_subreads=60

GGGAGAGATAAGGAGAGAGAGCGTTAACGCTCCGTTATCTAAGAATGGAAACTCGCCATTCCAACTAGCCTTTCAATCAAAATGGATCATTTGAACCCTGTCAACAGACCACCTCATCCCTTACGTCCCCCTCTTTGTCCCCATCCAGTGATCTGAATGCTTTTAAGCTCACAAGCTCCATCAGTGAGAAGCAAACATGGATGCTGGAAATGGACGACGGAGATCCGGCCACGTCCTCCTCCTCCCATTCCCGGCCCAGGGCCACATCAACCCCCTCCTCCAGTTTGGCAAGCGCCTTGCCTCGCACGGCGCCGCCACCACCATCGCCATCACCCGCTTCTTCCGAAAAACCTCGAAGCCCGCCGAAACTGTGGGCCCAGTCGCCCTCGCTGCCATTTCCGACGGCTTCGATGAAGGTGGCTTCAATGCGGCCGACAGCGCCCGGTCTTACCTCGACCGGCTCGAGTTAGCCGGGTCCGAGACATTAGCTGAGCTCATCGAGTCTGAGAAAGCTGAGGGCCGGCCAGTCGACGTCTTGGTCTACGACGCCTTCTTGCCGTGGGCCCTCGACGTGGCGAAGCAGCACGGAGTCGCGTCAGCTGTGTTCTTCACGCAGTCGTGCGCAGTGGATCTGGTGTTCTTTCACGCGTCGAAGGGGCGGGTGGCTCTGCCGACGATGGAGGCGGTGTCGCTTCCAGGGTTGCCGTGGTTGGATCTGCGGGACCTGCCGTCGTTCTTGACCGATCCGCTGCCGGGTCCCTACCCGGCTTACAGGGACGTGCTGTTGGGCCAGTTCAAAAACATCGACGGGGCCGACTTCGTCCTTATGAGCACTTTTTCCGAACTGGAGCATCAGGCAGTAGAGTGGATGCGATCATTGTGGCCTATAAAGACGATTGGCCCCACAGTCCCTTCGTCCTATCTCGACAACCGCTTGTGTGACGACACCCACTACGGCTTCCACTTCTTTGATCCCGATACCGGACGATGCATGTCATGGCTCGACTCGAAGCTCCCCAACTCCGTCATCTACGTGTCCTTCGGTAGCTTCTCCTCCGTCTGTGAAGAACAGATGACTGAGGTCGCGGCAGCCCTCACCAAAGCCGGAAACCACTTCCTCTGGGTGGTGAGATCCTCGGAGATGTCAAAACTCCCAGATGGATTTGTTCGAGATTTGTCGGAGATCGGTCTGGTGGTAAGCTGGAGTCCACAGCTGGAGGTGCTGGCCCATGAGGCAGTGGGGTGCTTCGTGACCCACTGCGGGTGGAACTCGACGCTGGAGGGGCTCAGCCTCGGGGTCCCGATGGTTGCGGTGCCCCTGTGGACGGACCAACCGACCAACGCCAAGTTTGTGGCAGACGTGTGGGGGGTCGGGGTGAGAGCGGTGACCGACGAGCATGGGATTATTGGGAGGGAGGAGATGGCGAGGTGTGTGAAAGCAGTGATGGATGGAGGTGAAAGGAGCGAGCAGATGAGGATGAATGCGAGGAAGTGGAAGGAGGCGGCAAAGGTGGCGGTCAATGCGGGCGGAAGTTCGGATAGGAACATTGTAGAGTTCGTTAAGAAATTTTGCTCATGAGTAGGAAATAAATTGAGACTGTTGACTGGTCCTTTTCCATCCAAATCCTTTTAT

>F01_transcript/47598 full_length_coverage=21;length=1612;num_subreads=60

GAGAGTGGTGATCAGAAAGAGAAGGAGAGTTTGAAGGATGGTGAAACCCGGAGTTGTGTTCATCCCCTGTCCGGCCATGGGCCACTTCATCTCAGCGGTGGAGTTCGCCAAGAAGCTCGAGAGCCGCTTCGCCGTCACCATCCTCCAAATTGCCCATATTCCCCCGGCATGGTCTTCCGCCATCAAGGCTTATGTGGAGTCGGTCGTGTCCTCCGGCTTCGACATCCGATTCGAAGAGCTCCCTCACATTGTTCCTCCTCATGCGGAGAAGCAAAGACCAGAGACCTTCGCCTCTCGTTTAATGGAGGTCAACAAATCGCCGGCAAGAGACGCCATCTCGCTGATTCCTAACAATAATATCGCGGCAATAATCCTCGATCTGTTCGGCAGCTCCATGGTGGACGTCGCCGATGAGCTCGGCATCCCGGCTTACATCTACTTCACCTCGAACGCGGCCTTGCTCGGGTCCATGCTCTACCTCCCCACCCTCCATGTAAAGGTCCCGTGCGAATTGATGGAGTTCGGAGGGCACATCTCGATGCCCGGACTCCCTCCCGTGCCATCCCATTGCATGCCATCGTTCGCGATGGACAAGAAGGACGAGGCCTACACCACGTTCATCAACCACGGATTGTGCTTTCGAAAGGCTAAAGGCCTAATCATTAACACATTTGAAGATATCGAGCCGAAGACCCTGCAAGCATTGGCGGACGGCGAGTACCTCCCCGACCACACTATGCCACCGGTCTTTCCCGTCGGACCGGTGCTGGCGATCAAGCCGAAGAGTGATCAGCCGCATGATTGCATCGCATGGCTCGACAAGCAACCTGTGGCTTCGGTGGTGTTCTTGTGCTTTGGAAGCATGGGATGCTTCGAAGCACCCCAGGTGAAGGAGATGGCGGTGGGGCTGGAACGGAGCGGCCACCGGTTCTTGTGGGCTCTTCGTAGCCCTAACACCATCGATAGCTTCCGGTTCCCGACCGACTCCGATCTCGACGAGGTGCTGCCCGGAAGGATTCTTGGAGAGGACTGAAGGGAGGGGGCTGGTGTCGCCGTCTTGGGTTCCACAAACTGAGATACTGTCCCACGAGGCGGTGGGGGGATTCGTCACGCACTGCGGGTGGAACTCAGTGCTGGAGGCGATGTGGCTCGGGGTGCCCATGCTCGGGTGGCCGCTATTTGGGGACCAGCACTTGAATTGTTTGGCGATGGTGAAACAGTCGGGCGTGGCTCTCGAGCTCAAGTTGGACTACAACAATGGCGGGTTCGTGAGTTCGGGGGAGCTGGAAAAGGGAGTTAGGTCCCTGATGGGGGACCTCCAGGATTCGAGGAAAGTGAGAGGTAAGGTGAATCAGATGAAAGTCGACAGCCGGAGAACCGTGATGGACGGGGGTTCATCCAATGTCGCATTGGAACAGTTGATTTGTGAATTAAACAAGAAGGTGGCGGTTGCCGTTACTGTTGATAAGAATAATAATTGCTATGTGACTCCTAAGAAGATTGTTGCTACGTGATGAGCTATGTTCGTTGGACAGCTCATGCTTCAAGTCCAACTTAAAATATTTTCTAATAACATATTATCTTTCCATTGCCATGTTAAAAAGAATACTCC

>F01_transcript/48138 full_length_coverage=2;length=1702;num_subreads=50

GAGAAGAAAGCAAGGCTATTACTTACATCCCATGCGATACTTTCACGCACAGCGAGCACTAAATGACGAGCTCGACTGTAGTGGGTAGTCCTCTACCCGTCACCGGGGATGGGCCACCTCGTCTCCATGGTCGAGCTTGCGAAGGTCCTCCTCGGAAACGGGCTCTCGATCACCATCCTCATGGTCGAGCCCCACTACAACACGGGCGCCACCGCTCCCTTCATCTCCGCCGTCTCCGCCGCTCACCCCTCCATTGCCTTCCACCACCTCCCCTCCCTCTCCCCACCCCCCAACTCCTCCCCACACCGTGAGGCCCAGGCCATGGACCTCCTCCGCCTCTCCAACCCCAACCTCCAATCCTTCCTCCAATCCTCATCCCCCTCTCCCGCCGCCCTGATCATCGACTTCTTCTGCGGTTTCGCTCTCAATGTATCCTCCCATCTCCGTATCCCAACCTACTGCTTCTTCACCTCGGGCGCATGCTTCCTCGCCGTCTTCCTCAAGTTACCCAACATCCACTCCCGATTTTCGGTCAGTTTTCGGGAGCTCGGCTCGACCCTTATCGACGTCCCCGGTATCCCTTCCATCCCTGCCGATCACATGCCGCTGGTACTGCTTGATCGGGACGACGAAGCTTATAAGGGGTTTTATTACCTATCGGAGCGATGTTCTGAGTCGGACGGGATAATCGTCAACACGTTCGATGCGCTCGAGCCAAGGACGATCGAGGCCATCTCGAGAGATGGAATGAAACTGTACTGTACCGGGCCGCTGATCACCGAGGAGAGGGACATGACGAACGGGGCGGATTGCCTTGAGTGGCTCGATGGGCAACCGAGGGGGAGCGTCGTGTTCCTTTGCTTTGGGAGTCTTGGACTTTTCTCAGTTGAGCAGCTGAGAGAGATCGCCATCGGGTTGGAGCGGAGCGGGCAGCGATTTCTGTGGGTGGTGCGGAGCCCGCCGAGCGACGATCCGGCCAAACGGTTCGCTAAGCCGGCTGAGGCGGTCCTCCACGTGCTGCTCCCCGATGGGTTCCTCAACCGAACAGGCGACAGGGGAATGGTAGTGAAGTCGTGGGCCCCACAGACGGCCGTTCTGCGCCACCCGTCAGTGGGCGGCTTCGTGACGCACTGCGGGTGGAACTCCATCTTGGAGGCGATATGCACGGGGGTCCCCATGGTCGGGTGGCCACTCTACGCTGAGCAGCGGGTGAACATAATTTTTCTCGTCGAGGAGATGAAGCTGGCCGCGGCGATGGAGGGCTACGATTCGCCCCTCGTGTCGGCGGAGGAGGTGGAGACACGTGTGGGGTGGCTGATGGACTCGGACGGCGGGCGGGAGCTGAGGTTGCGTACGGAAGCGGCGAAGAAGGCTGCGGCCACGGCGCTGCAGGAGGGTGGGTCGTCACGTGAGGCGTTGGTGGAGCTGGTGGCGCGGCTGAAACGGAGGTGATGGAGGCGGGTGTCCTTATCGACCGTGTGGGTGCCGTGACGAGATTAAATTCAGGAGCTGGTAATAAAACGAAACAAAACAAAATGGGGAACTTTGGGGTCTGGAATCCTATTCGTTGCTTTGGTGCCGTGACGGCATTAAATTATGATTGTCGTCTTTGTATTTTTTTTTTTTCCTTCTCAGTTCAGCGTCTCCTGTAACAATGTTTTCAAAATCGGACCGATCACATATATGACGTTTTAAATCATCC

>F01_transcript/48305 full_length_coverage=4;length=1586;num_subreads=60

ATGGTGACTCATTTCCTGATTTCAGTGGCTATTGACTCATGAGCTCCCTCCGCATCGTCTTCCTCCCCATGTTGGCCGCAGGCCACATGATCCCCATGGTCGACACGGCTCGTCTCTTAGCCGCCCGTGGCGTCAAGTGCACCATTGTCACCACGCCCGGCAACGCCCCCGTCATGGACCGCGTCAAACATCAGAACAACATCGAGATCCTCCTCATCCCCTTCCCCTCGGCCGCCGTCGGGCTCCCCGACGGCATGGAGAACTTATCCGCCTCCTCCTCCCCGGAGTTCAAGATCAAGTTATTCCAAGCCCTCGATATGCTCCGCCACCCGTTTGACAGAATCCTCACCGACCTCCGCCCCGACTGCATCGTCACCGACATGTTCTTCATTTGGTCTGCCGACGTCGCCGCCGAACACGGCGTGCCGAGGCTCGCCTTCTTCACCTCCAGCTTCTTCTCGCAGTGCGCCCAGGACGCGATATTTCGCCACAACGTATTGGAGAGCTCGCCAGCGGATGTGGAGTCTTTTGTGGTTCCGGGCCTGCCGGATAGGATCGAGATGCTCAAGTCCCAAATCCCGAATCCTAAGTTTGTGGTGGGCATGGAAAGGATGATGCGACAACTGAAGGAGACGGTGGAGCAGAGCTATGGCGTGTTGGTGAACAGCTTCTATGAGCTGGAGCCGGAGTATGTCCGGCACTGCCGGGAGGTGGTGGGGTGGAGGGCGTGGCACGTGGGCCCCGTGTCGCTCTGCAATGTGGAGGAAGCCGACAAGGTCGGGAGAGGGGGGGACAAGAACAAAATCTCCTCCGAGTCTTGCTTGGAATGGCTTGACGGTAAAGAACCGGGCTCGGTGTTGTACATTTGCTTCGGGAGCGAGGGCGACTTCACTGCAGCTCAGCTCAACGAGCTCGCGGTGGGGCTCGAGGCCTCCGGATGCCCTTTTATGTGGGTGGTGAGGAGCGGAGTGTTGCCGGCGGGGTTCGAGGAGAGGGTGGAGGGGAGAGGGATGGTGGCGAGGGGTTGGGCCCCGCAGATCTCCATACTGAGCCACCGGGCGACCGGGGGGTTCCTCACGCACTGCGGGTGGAACTCCAGCATGGAGGGGATAGTGGCGGGTCTACCGATGGCCACGTGGCCGTTGACAGCGGATCATTTCTACCACGAGCGGCTGCTGGTGGACGTGCTGGGGGTCGGCGTGGCGGTGGGGGCCGAGAACTTCTCGTTCATGTATGAGGACCGGCTGCTGGTGGGGTCGGCCACAGTGGAGAGGGCGGTGGGGAGGGTGATGGCAACCGGAGAGGAGGCAAATAACCGGCGGCGACGGGCGAGGGAGCTGAGCGATCTGGCGAAGGGGGCTGTGCTGGACGGCGGATCATCGTTTGTAGAGATGGGGAATCTGATCAAGGAATTGACGGACATGCGGAATTTGGCCACTAAAAATCAATGAGTTGGCGAATTTTTTTTTCTCTTTGGTTTGGAGTATGTTTTCTCTTTGATTTGGAGATTGTCTACAAGTTTATAATTTAAGCTCTTGCAACTAAAGATGACAAAAAATACATTTATTAAAATTATAATTTTATCT

>F01_transcript/48334 full_length_coverage=7;length=1593;num_subreads=60

GCTCTCTCTACGTTGTATACTCATTTCCTGATTTCAGTGTCTATTGACTCATGAGCTCCCTCCGCATCGTCTTCCTCCCCATGTTGGCCGCCGGCCACATGATCCCCATGGTCGACACGGCTCGTCTCTTTGCCGCCCGTGGCGTCAAGTGCACCATCGTCACCACACCCGCCAACGCCCCCGTCATGGACCGCGCCAAACGTCAGAACAACATCGAGATCCTCCTCATCCCCTTCCCCTCGGCCGCCGTCGGGATCCCCGAGGGCATGGAGAACTTTTCCGCCTCCTAACTCCCCGGAGTTCAAGATAAAGTTCTTCGAAGCCCTCGATATGCTCCGCCACCCGTTTGACAGAATCCTCACCGACCTCCGCCCAGACTGCATCGTCACCGACATGTTGTTCATCTGGTCTGCCGACGTCGCCGCCGAACACGGCGTGCCGAGACTCGCCTTCTTCACATCCAGCTTCTTCTCGCTGTGCGCCCCGGACGCGATATATCGCCACAACGTATTGGAGAAGCTCGCCGGCGGATGTGGAGTCTTTTGTGGTTCCGGGTCTGCCGGATAGGATCGAGATGCTCAAGTCCCAAATCCTGAATCCTAAGTTTGTGGTGGGCATGGAAGGGATGATGCGACAACTGAAGGAGACGGTGGAGCAGAGCTACGGCGTGTTGGTGAACAGCTTCTACGAGCTGGAGCCGGAGTATGTCCGGCACTGCCGGGAGGTGGTGGGGTGGAGGGCGTGGCACGTGGGCCCCGTGTCGCTCTGCAATGTGGAGGAAGCCGACAAGGTCGGGAGAGGGGGGGACAAGAACAAGATCTCCTCCGAGTCTTGCTTGGAATGGCTTGACGGTAAAGAACCGGGCTCGGTGTTGTACGTTTGCTTCGGGAGCGAGGGCGAGTTCACTGCAGCTCAGCTCGACGAGCTCGCGGTGGGGCTCGAGGCCTCCGGATGCCCTTTTATGTGGGTGGTGAGGAGCGAAGTGTTGCCGGCGGGGTTCGAGGAGAAGGTGGAGGGGAGAGGGATGGTGGCGAGGGGTTGGGCCCCGCAGATCTCCATACTGAGCCACCGGGCGACCGGGGGGTTCCTCACGCACTGCGGGTGGAACTCGAGCATGGAGGGGTTAGTGGCGGGTCTGCCGATGGCCACAGTGGCCGTTGGCAGCGGAGCAGTTCTACAACGAGCGGCTGCTGGTGGACGTGCTGGGGGTCGGCGTGGCGGTGGGGGTCGAGAAGACCTCGTTCACGGTATGAGGACCGGCTGCTGGTGGGGTCGGCCGCAGTGGAGAGGGCGGTGGGGAGGGTGATGGCAAGCGGAGAGGAGGCGGATAACCGGCGACGACGGGCGAGGGAAGCTGAGTGATCTGGCGAAGGGGGCGGTGCTGGACGGCGGATCATCGTTTATGGAGATGAAAATCTGATAAAGGAATTGACGGACATGCGGAATTTGGTCACTAAAAATCAATGAGTTAGCCAATTTTTTCTCTTTGGTTTGGAGTTTGTTCTCTTTGATTTGAAGATTGTCTAGAAGGTTATAATTTAAGCTCTTGCAATTAAAGATGACAAAAAATGTATTTACTAAAATTATAATTTT

>F01_transcript/48407 full_length_coverage=3;length=1589;num_subreads=45

GATTTAAACAAAGGTTGGACTATGGCGTCACAAACTCAACCCCTTCACATGCTCTTCTTCCCTTTCATGACCGCCAGCCACATGATCCCAATGGTGGACATGGCGAAGCTCTTTGCCGCACGAGGAGTGAAGGTAACCATCCTCACAACTCCATCAAACGCTGAAACAATAAAATCCGCCATTGAATCCGGCCACCAAGCCATCGCCCTCGCGCTCATCCCCTTCCCCTCGGTGGCCGTCGGCTTGCCAGAAGGCTGCGAGAACCTCTCCGCTGTGCCCACTATCGAGCTACAGAACAGATTCTGCAGAGCACTGGCAATGCTCCGCCAGCCGTTCGATCGAGTCCTCATCGACCTCCTCCCTGACTGCGTCGTCTCCGATTCGTTCCTCCCGTGGACCTACGAAATCGCCGCCAAATATAATGTGCCGAGGCTCGTGTTTCACGGCACGAGCCACCTCTCTATTTGCATCTCGGACAGCTTCGCTCGTCACGTGCCGGTCGTGAGCTTGGCACCGGATGGCCACGAGTGTTTGATAGCCCCGGGTCTACCTCACCGGATCAAGATGCTCCGGTCCCAGATGCCCGACCTCACCTGTTCCGGATCCGATTTACCGCAACTTCTGGCAGAGGAGGCGAATCTGAAGAGCTACGGCCTGGTGATGAATAGTTTCTACGAGCTGGAGCCGGAATACGTCCGTCACTTCCAAAAAGTCCTCGACCGAAAATCGTGGAATGTGGGTCCTGTGTCCCTATACAACAAAGACACAATAGAGATGCGATTGAGGGGACACACAAGTGTAGTCGACGTCAACAATTGCTTGGAATGGCTCGACGGCAAGAGCTCGGGCTCGGTTGTGTACATCGCATTCGGAACCCTAGGCGAGATGGGGATGGATCAAGTCCGCGAGATAGCTCTCGGGCTCGAGGCTTCAGGCCGACCTTTTATATGGGTGGTGAGGAATGTTGATTGGCTGCCGGAGGATTACGAGGACATGGTCGGGGATCGGGGGATGCTGATAAGGGGTTGGGCACCGCAGGTGCTGATCCTAAGCCATCAGGCGGTGGGGGGAATGGTGACACACTGCGGGTGGAACACGTGCTTGGAAGGGGTGTCGGCCGGGGTGCCGATGGTGACGTGGCCGTTCTTTGAGGAGCATTTCTTTAACGAGTTATTGATCGTGGAGTTGCTGGGGATCGGGATAAGCTTGGGGGTGAAGGAGTGTGCGATAAAGAAAGAGAGGCCGGTGGTGGATTCGGCGAAGATAGAAAGGGCGGTGAATCAGCTGATGAGTGAAGGAGACGAAGCGGAGGTAAGGAGGAGGAGGGCGAGGGAGCTCGGAGAGAAGGCAAAGAGGGCAATAGATGAGGGTGGCTCGTCGTACATTGATGTTGGGAATTTAATCGCTGAGTTGAGTTGCAGGCCGAGACCAGTCCACAACAAGATACCAGCAGTTTCATCCTAAAGCACACATAAGCATTATGTATCTAAAGCGCAATCTCTACAACAGAACATGCTTTTAACCGCCCACAAAAGTATGTCCTAAATAATTTTTCATATACTCAATGTTATAAAAAGTCCTTACCTATT

>F01_transcript/48910 full_length_coverage=10;length=1558;num_subreads=60

GGCCCACTCAGTTTCTGTGAGGCCTGCGAGCAACGAATCAATAATCTCCGCCGCCATGGAATCCGGCCACGAAACCCGCCACATCGTGGCAATCCCCTACCCGGCTCGAGGCCACATCAACCCCATGATGGCTCTCTGCCATCTTCTGGCCGCCCGAGACGTCCACGTCACCGTCACCGTCGTCGTCACTGACGAGTGGCGCCAGCTTCTCACCTCCGGCGAGCCAACTTCTCACCTCCGGCTCCACGCCATTCCCAACGTCCTGCCCTCCGAGAAGACACGCGGAAATGACTACCGGGCCTTCACGGATGCCGTCTTTTCCAAGATGGAGGGGCCTGTCGAGGCCGCGATCGAAGGGATCGATCCACCGGTGGAAGCCGTCATCACCGACTCACTGCTGCTGTGGGGGGCGGCGATTGCCGGACGGAGGAACGTTCCGGTGTGGTGGCTGTGGACGCACTCCACCACCGTGTTCCGGGCGTTGCATGAATTCGACAAGCTCGACAAAGCCGGCCGATTGCCGACAGACATACCGGCCAAAGGAGATGATCCATTAGCCTACCTCCCAAGACCCTTCATCCCACGTTCCGCCGACATCACCACCAATACCTCCAACATAGTTAGTTTTCAGAATTTTGTATCCCTTTTCTCCTGGTTTCCAAGAGCTCAAGGCCTCATCTTCACCTCCGTCTATGAGCTGGAAAGCCAAGCCATTGATACTCTACAATCGGAGATCCAGGTTCCATTTTACACATTTGGCCCTCCCATCCCTTACATGTCACATCATGATGATTTTGAAGCTACAAGGCCCGACTACATCAAATGGTTGGACTCTCAGCCCGTAAGCTCTGTGCTCTACATCTCGTTGGGGAGCTTCTTACCGATCTCGGGCCTACAGATTGAGGAGTTGGGAATTGGGCTCCATGAGAGCGGGGTTCGCTTCCTGTGGATATCCCGTGGAGAACCAGAATACACCAAACATACGAGCAGGAGCGCAGGCCAGGTGGTGAGTTGGTGTGACCAGCTTAGAGTACTGTCTCATCCTTCAGTCGGAGGATTCTTGACACACTGTGGTTGGAACTCCACTCTTGAGGGGGTCTTTACTGGGGTTCCAATGCTTACGTTCCCTCTGATCTGGGATCAGTACCCGAACGCTAAGTCAGTCGTTGATGACTGGAAGGTCGGCCTCAGGTTGAAGGATGAGGGGGAAGAGGAGAGGATCGTAGGTCAGGAGAAGATTGTGAGGATGGTGAAAAAACTGATGGCCTTGGATGATGAGGAGAGCGAGGGATTGAGGAAAAGGAGTGCGGAGCTCAAGAGCAAGTGTCAGAAGGCATTGGAGGAGCATGGATCATCAACTGCCAACCTCGATGCTTTATTCCAAGAGATTATCGGTAAGAAAAGCAGTAGGAGTGATATTTGAAAATGCTCTTTCTAATGGTATTTGTTTAAGTCATGATACCAGATGCTTGAATAATTCTCTCATTTAAACAGTTCATCCACCTATGTTCTTCAAACATGAATATGATTTAGGAGTGAGCTTGTATTGCGCAAATCT

>F01_transcript/49173 full_length_coverage=4;length=1563;num_subreads=60

GAAAGTGAAGATCTCTCTCTAGCGACGGGGTAATTTTCCGAATCATGGGCTCCGTTCCGATAGATCTTCACGTCTTCTTTTTGCCCTTTCTTGCTTCGGGACACATGATCCCCATGGTGGACATCGCTCGCCTCCTCGCCGACCGAGGCGTCAAATCCACCGTGGTCACCACCGCGGGGAACATCCCCCGAATCCAGCCCACCATCCAACATTTCAACTCCATTTGTTCCTCCGACAAACCTCCAATCGAACTCCTCACCATCCCCTTCCCCTCCTCCGTCCCCGACAACGTTTCCGGCCTCCCGACCCCCGACCTCACTCCCGAATTCGCCGCCGCCATCTGTGACCTCCGCCAACCCTTTGTCGAGCTCCTCGAATCCCACCGCCCCGATTGCATCATCTCCGACATCTTCTACCCGTGGACCAACGACCTCGGATACCCGAGGATTGCTTTCCACGGCATCGGCTTCTTCTCCTGTGTCGTTCCCGGCACCCTCGCATATCAAAAGCTACACGAATCCGTTACCGCCGATGATGAACCATTCGTCGTGGGTGGCCTCCCGCATAGGATCGAGATGACCCGCTCCCAGCTCCCGGGCTTCTTCATGTCGCCTGACAGCTTGTTGATATCGGGTGTCGTCGACTGGCATCACAATTGCTACGGGGTTGTGATGAACAGCTTCTACGAGCTGGAGCCCGAGTACGCCGACATCATGAAGATGAACGCCAGCTTTAAGATCTGGCATGTCGGGCCTGTGTCACTCTCTGGCAGCAACACTGCCGGTCGACTGGTCGAATCCGACCTTATTGGTAGCTGGCTCAATGGCAAAAACCCGAGCTCGGTTCTTTACGTCTGCTTCGGGAGCTCGGGGAAGTTCACGACGCCCCAGCTCCGTGAGATCGCATCCGGCCTCGAGGATTCCGGCCACCCGTTTATATGGGCGGTGAACAAGAGCGACGAAAATCTGCCCGAGGGGTTCGAGGAGAGGGTGAAGAGAAAAGGACTGGTGATAAAAGGGTGGGCGCCGCAGGTGATGATATTGAACCACCAGGCTGTCGGTGGGTTCATGACGCACTGCGGCTGGAATTCGTGTTTGGAGGGAGCAAGCGCCGGGTTGCCGATGATCACGTGGCCGATGTTTGGTGACCAGTTTGTCAACGAGAGACTGATTGTCGATGTCGTGGGGATGGGGATAGGAATCGGGACGAAGGTTTGTAGCGCACATGAGGAAGAGAGGACGGTGGTGAAGGGGGAGGATGTGGCCAAGGCGGTGAGGGGATTGATGGGCGGCGATGAGGCGGAGAGGAGGAGGAAGAGGGCTAGGGAGGTCAGAGAGAAGGCGAGGAGAGCAGTGGAGGAGGGCGGCTCGTCGTACAGCGAAACGGACCGTCTCGTAGAGGATATAATCAATTTGAAAACGGCTCGGGTCCAGTTCGATCACCCCAAATAAGACTTAGATTTTGTGTGGATCCATTATCCTCTGCTAAATTCATGGATTTTGTGTTGTTTTATTGGGATTGGATCTATGCCATATATTGCTCTACGATTGAGGTTTTTTAGTT

>F01_transcript/50365 full_length_coverage=3;length=1509;num_subreads=48

GACACCAGAGCTCACAGAAGAGTCTCTCTCTCTCTCTCTCTCTCTCTCTCTCTCTCGCTCGTCATGGATGGAGGAAACCTGCACATAGTCGTGTTCCCATGGCTAGCCTATGGCCACATGATGCCCTTTCTTGAACTCTCCAAATCTTTGGCCATAAGGGGTCACCAAATCTCCTTCATCTCCACCACAAAAAACATAAAGAAGCTCCAACCCAAAGTCCCACAAAATCTCTCCCAACTCATCCAATTCATCCCTCTCCTTCTTCCGCCAACCGACGGCTTACCGGCCACAGCTGAGGCCACCTCCGACATCCCTCCAAACCTCGTCCAATACCTCAAGAAGGCATTCGACTGTCTCGATCGTCCCTTCGCCCAATTCCTCAGTTCATCGTCCCCAAAGCCGGATTGGATCATTCTAGACTTCGCCTCCTACTGGCTCCCGCCACTCGCTTCCAAATTCAATGTGCCATGTGTCTACTTTTGCATCTTCTTCCCCTCAGCCCTGGTATTCGTTGGACCCATGTTAGAAGTCGACAACATAGGCTCCGTCACGGCTGAACAATTGATCGTTCCGCGCAAATGGATCACATTCCCTACCAACATGGCCTACCACCCTTATGAGGCCCGAGAGGCCGTTGAGTTCTTCAAATCAAGCAATGCATCGGGAGTGTCCGATGCTCACCGCTTCTTGCTCACCATCAAAGGTTGCAAGGCCGTGGCCATCCGGAGCTGCAATGAGTTCATGCCCCAGTGGTTGTCCCTCCTCCGGGATCTATATAAGAAAGCCATCATCCCGGTTGGCATGCTTCTCCCGTCGCTGGCTGAAAATGAAGATTCTAACACCAGCGAACTCAGAGTCATGGAGTGGCTAGACCAACAATCTTTGAGTTCTGTCGTGTATGTAGCCTTTGGGAGTGAAGCCAAAGTCAGCATCGAGCTATTACATGAGCTAGCGCTTGGGCTCGAACTCTCTAACTTTCCTTTCCTTTGGGCACTTAGAAAGTCGGCCACCGACGAAAGGGAAAACATTTTGCCTGAAGGATTCGAAGAACGAACAAAGAACCGAGGATTTGTGTCCATGCATTGGGTTCCCCAGCTCAGGGTCTTGGCCCACAATTCAGTGGGGGGATTCTTGACACACTGCGGTAGCAGTTCGATCATCGAGAGCCTCCATTTTGGGCGCCCTCTAATCCTCTTCCCTATATTCTTGGATCAAGGGCTTAATGCTCGGGTGATGGAGGAAGAGAAGATCGGATTCGAGGTGAAGCGGGGCGAGGAGGATGGGTCACTAAGAAAGGAAGTGGTAGCCAAGGCATTGAGATTGATTGTCATCGACGTAGAGGGAGAACCTTTCAGGAAAAATACCAAAGAGATGATGAGAGTTTTGGTGGAAAAAAAATTTCATGAGAGATATGTGGATGATCTCACTCGGTATCTAATGGACCATAAGGATTGTAAATGAATGCTGGTTGTAGTTTATGTTTAGTAATAATCATTTTTTATTCTCTCC

>F01_transcript/50724 full_length_coverage=2;length=1482;num_subreads=60

GTCTCTACGTTGTATACTCATTTCCTGATTTCAGTGTCTATTGACTCATGAGCTCCCTCCGCATCGTCTTCCTCCCCATGTTGGCCGCCGGCCACATGATCCCCATGGTCGACACGGCTCGTCTCTTAGCTGCCCGTGGCGTCAAGTGCACCATCGTCACCACGCCCGCCAACGCCCCCGTCATGGACCGCGCCAAACGTCAGAACAACATCGAGATCCTCCTCATTCCCTTCCCCTCGGCCGCCGTCGGGATCCCCGAGGGCATGGAGAACTTTTCCGCCTCCAACTCCCCGGAGTTCAAGATAAAGTTCTTCGAAGCCCTCGATATGCTCCGTCACCCGTTTGACAGAATCCTCACCGACCTCCGCCCAGACTGCATCGTCACCGACATGTTGTTCATCTGGTCTGCCGACGTCGCCGCCGAACACCGCGTGCCGAGGCTCGCCTTCTTCACCTCCAGCTTCTTCTCGCAGTGCGCCCCGGACGCGATATATCGCCACAACTTATTGGAGAGCTCGCCGGCGGATGTGGAGTCTTTTGTGGTTCCGGGTCTGCCGGATAGGATCGAGATGCTCAAGTCCCAAATCCTGAATCCTAAGTTTGTCGTGGGCATGGAAGGGATGATGCGACAACTGAAGGAGACGGTGGAGCAGAGCTACGGCGTGTTGGTGAACAGCTTCTACGAGCTGGAGCCGGAGTATGTCCGGCACTGCCGGGAGGTGGTGGGGTGGAGGGCGTGGCACGTGGGCCCCGTGTCGCTCTGCAATGTGGAGGAAGCCGACAAGGTCGGGAGAGGGGGGGACAAGAACAAGATCTCCTCCGAGTCTTGCTTGGAATGGCTTGACGGTAAAGAACCGGGCTCGGTGTTGTACGTTTGCTTCGGGAGCGAGGGCGAGTTCACTGCAGCTCAGCTCGACGAGCTCGCGGTGGGGCTCGAGGCCTCCGGATGCCCTTTTATGTGGGTGGTGAGGAGCGAAGTTTTGCCGGCGGGGTTCGAGGAGAAGGTGGAGGGGAGAGGGATGGTGGCGAGGGGTTGGGCCCCGCAGATCTCCATACTGAGCCACCGGGCGACCGGGGGGTTCCTCACGCACTGCGGGTGGAACTCGAGCATGGAGGGGTTAGTGGCGGGTCTGCCGATGGTCACGTGGCCGTTGGCAGCGGAGCAGTTCTACAACGAGCGGCTGCTGGTGGACGTGCTGGGGGTCGGCGTGGCGGTGGGGGTCGAGAAGACCTCGTTCACGTATGAGGACCGGCTGCTGGTGGGGTCGGCCGCAGTGGAGAGGGCGGTGGGGAGGGTGATGGCAAGCGGAGAGGAGGCAGATAACCGGCGACGACGGGCGAGGGAGCTGAGTGATCTGGCGAAGGGGGCGGTGCTGGACGGCGGATCATCGTTTGTGGAGATAGGAAATCTGATAAAGGAATTGACGGACATGCGAAATTTGGTCACTAAAAATCAATGAGTTGGCAAATTTTTTTCTCTCC

>F01_transcript/52887 full_length_coverage=3;length=1389;num_subreads=60

ACCTCCTCCGCCTCTCCAACCCCAACCTCCAATCCTTCCTCCAATCCTCATGCCTCTCTCCCGCCGCCCTGATCATCGACTTCTTCTGCGGTTTCGCTCTCAATGTATCCTCCCATCTCCGTATCCCAACCTACTGCTTCTTCACCTCGGGCGCATGCTTCCTCGCCGCCTTCCTCAAGTTACCCAACATCCACTCCCGATTTCCGGTCAGTTTCCGGGAGCTCGGCTCGACCCTTATCGACGTCCCCGGTATCCCTTCCATCCCTGCCGATCACATGCCGCTGCTACTGCTTGATCGGGACGACGAAGCTTATAAGGGGTTTTATCACCTATCGGAGCGATGTTCTGAGTCGGACGGGATAATCGTCAACACGTTTGATGCGCTCGAGCCAAGGGCGATCGAGGCCATCTCGAGAGATGGAATGAAACTGTACTGTACCGGGCCGCTGATCACCGAGGAGAGGGACATGACGAACGGGGCGGATTGCCTTGAGTGGCTCGATGGGCAACCGAGGGGGAGCGTGGTGTTCCTTTGCTTTGGGAGTCTTGGACTTTTCTCAGTTGAGCAGCTGAGGGAGATCGCCATCGGGTTGGAGCGGAGCGGGCAGCGGTTTCTGTGGGTGGTGCGGAGCCCGCCGAGCGACGATCCGGCCAAACGGTTCGCTAAGCCGGCAGAGCCGATCCTCCACGTGCTGCTCCCCGATGGGTTCCTCAACCGAACAGGCGACAGGGGAATGGTAGTGAAGTCGTGGGCCCCACAGACGGCCGTTCTGCGCCACCCGTCAGTGGGCGGCTTCGTGACGCACTGCGGGTGGAACTCCATCTTGGAGGCGATATGCACGGGGGTCCCCATGGTCGGGTGGCCACTCTACGCTGAGCAGCGGGTGAACATAATTTTTCTCGTCGAGGAGATGAAGCTGGCCGCGGCGATGGAGGGCTACGATTCGCCCCTCGTGTCGGCGGAGGAGGTGGAGACACGTGTGGGGTGGTTGATGGACTCGGACGGCGGGCGGGAGCTGAGGTTGCGTACGGAAACGGCGAAGAAGGCTGCGGCCACGGCGCTGCAGGAGGGCGGGTCGTCACGTGAGGCGTTGGTGGAGCTAGTGGCGCGGCTGAAACGGAGGTGATGGAGGCGGGTGTCCTTATCGACCGTGTGGGTGTCGTGACGAGATTAAATTCAGGAGCTGGTAATAAAACGAAGCAAAACAAAATGAGAAACTTTGGGGTCTGGAATCCTATTAGTTGCTTTGGTGCTCTGACGACATTAAATTATGATTGTCGTCTTTGTATTTTTTTCTTTCTCTTTTTCTTTTTTTTTCTCAGTTCAACGTCTCCTGTAACAATGTTTTCAAAATCGGACCGATGAAATATAAATACATTGGTTGAACC

>F01_transcript/12407 full_length_coverage=3;length=2842;num_subreads=60

AAGGAAAAAATAAAAAACTATCATTCTCCGGCGAGAATGGAGAGTGAGAGCAGCCAGAGGCACTTCGTGCTAGTCCCGTGGCTGTCCCACGGGCACGTGATCCCCATGATGGACATGGCCCGGCTCCTCGCCGGGCGGGACGGCATCCACGTGACCGTGGCCATCTCCCCCGTGGGTGCGGAACGCATCCGGAGCTGCTTCATAGAGCCCGTGGCCGCCGCGAAGCTCCCCATCTCCTTCGTCGAGCTCCCCTTCCCCTGCGCCGAAGCCGGTCTCCCTGAAGGCGTGGAGACCATCGAGCAAATCCAAGACCCCTCCCTCTTCCCCAAGATGCACGTGGCCGCCGGCCTCCTCCGCAAACCACTCGAGTCCAAACTCCGGGAGCTCCCCCGCAAGCCCTCCGTCATCCTCGCTGACCTCTACCACCCGTGGGCGCGGGAAGTCGCCGCCGACCTCGGCGTCCCGCTGCTGCTCTACTACGTGTTCCCCTGCTTCACCATCCTCGTCTACCGCAGTCTGAGACAGCATGGTATCTACGATGACGGCGCGGCGGACGCGAGTCGGATGTTCCCGGTGCCCGACGCCCCGGAGTACATGGTCAGCCGGGCGCAGGCGCCGGGGACCTTCGACAGGCCCGGGTGGGAGTGGATTCGCGAGGAGGCTATTGCAGCCGAGTCCGCCGCCGCCGGGGTTATTTTTCACAGCTTCGACCAGCTCGAGCCCAATTTCCTCCCCAAGTTCCAGGTACTCTCTCCCCCGTTGTGCTCTATATGGTGCTTCAAAATGAGAAAATGCATCAATGATCCTCTATATTCCAAGAATCATTGTAGCTCTTTCAAACTTTATGTTCATCCTTTTCAACCCTTTGTCTTATTTTTTTTCATTAAAATATTAAAAATTTTAAATCACCTCATGTCATTTAGTTATCAATAAAATATTATATTTTATTATTATACAGCATCAATGATTAAAGTTTGCTTCCTCTATATCACATATTTCGATAAAATGTAATTTCAAAATGAAGGCAACTATTCTAAATGCTCCGGGTCATCTCCGGACCTCAGTAGAAACTTGACTAAATTTAAATTTTGAGTTGAAAAAAAACACAAAAGACAAACTCTATGTTTACTCAGAATACAGGGAGAAGTTGAGCGACTCTAACGGGTGGCCATCCTAGTAGGTGGCATTCATTTTCCCAACCTAAAATAACTATTAGGGTTCACTGTTTTGACTCTCACGTGGGTTCCATAAGAGCAAATCCAACTGGAATATGAAAACGAGTATGAAAAATGATGGGTCCCATGCATATTTTTATACCCATTTGGTGGGCCATGCATTTTTACATACTCAGCAGATTTTAGACTTAGGTATGAAAATTTTGAAATGATGAGAGTAATGAATAAAATATAATAAAATATATTAAATAATTAACTATTGACTGAGTCAATTGAATAAAATATAATAGATCCTCACATGGTTAATTTTTCAGACTCAACAGTACCGTCTGAAGATTTCCGGGTTTTCATACCCCCTTGAGTTGCAAGGGCAAAAAAAAAAGATTCAGGAGAGAGAAAGTTCCAAACTCAAGAGAGAGAATGTGATAGTTGGCCTTTTTATTTTTAATTAATTTTTTATGGTCTCCTTTACTTTTCCCCATGTGGTGTTCTTCCTCACCATCTCCATTTAAATAAAATATGATGTGAGGTCTATCTTCATTATTTTAGAGATTAAAATATTTTAAAATATGTATGGAGATGCAAGATGAATATTTAGGTTGGAGTGAATATTTTTTTACCAGAGAGGAGCATCTTAGCGGAGACCGAGGTGAAACTCGGGGTGCCTGAATTATATTAAAAAAAGTATATCTAGTTTATATATATTTTTTAAAATTCCGTGCATTGTATGTGCATTGGATGGCTAAATTTTCTACTGCCTCCAGGAGATCATGGGTGGCCTGAAGACGTGGACCATCGGCCCGCTGTCCCTCAGCCACAAGAACGTGCTGGCAGAGCGCGGGAGCGCAAATGCAGTAGCCGCCGACAGCTGCCTCACCTGGCTCGACGCCAACGCCCCCGCCTCCGTCATCTACGTCTGCTTCGGCACCAACACATACTGGACCCCTCAGCAGATCATTGAGGTCGGGTCCGGGATAGAGAGCTCGGGCCACCCCTTCATCTGGGTGCTGAAGAAGCGGGAGCTGACGCCCGAGGTGGAGGAGTTCCTGTCGGGAGGGTTCGAGGAGCGGGTGCAGGACCGAGGCCTGCTCATCAGGGGCTGGGCCCCTCAGGCGGCCATACTGACTCATAAGTCAATCGGGGGATTCATGACGCATGGCGGGTGGAACTCGTCGATCGAGGGGGTGGCGGCCGGGGTGCCAATGCTGATGTGGCCGCACTTCGAGGACCAGTTCCTGCACCAGATGATCATCGTTCAGGTGCTAGGGATGGGGATCGGAGTCGGGGTGCGGGCGCAGGAGGACTACATCGCGCAGGTGATGGACACCATCAAGCGGGAGCAGGTCGAGAAGGCGGTGAGAGAGCTGATGGGAGGAGGGGAGGAAGCCGACATGAGGAGGAGGAAGGCGAAGGAGTACGGGGAGAAGGCGAGGAATGCCATGGAGGTCGGGGGGTCGTCGTATGTGAACCTGACCGAAGTGATCGACTCCGTTCCGTTCGTCGCCGCCACCGAGAATGGTGGCGGTGACTAACTATTTTCTTCATTTATCTTCTTGGTGCCGAAACCAAATTGCCGTCATTTCAAATTCCTCTACCTATCTCTGTTCGTCGACTGTAATAAAATAGTACTATTGAATCAGTGATTAGTAGTTTTTCTTTTGCCGCTT

>F01_transcript/23323 full_length_coverage=2;length=2388;num_subreads=60

GAGTTTCAAGTAGAGAGAGAGAGAGAAAACGAACACTGATATCACCCACATCTCAATGGAGTCCCAACCAGAGCCCCTCCACCTAGTTTTCTTCCCCTTCCTCGCTCGCAGCCACATGATCCCAATGCTGGAGACCGCCCGCCTCGCCGTCGAGCGCGGCGTCAAAACCACCCTCGTCACCACCCCTGCCAACGCCCACCTCATCCACCCCGTCCTCCACCGCTCCAACTCATCTCTCCTCCCCTCCCATCCGCCGATGCAGCTCCAACTCATTCCCTTCCCCTCGGCCGAGTTCGGCATCCCCGAGGGGTGCGAGAACCTCACCTCTATCCCCCTCCCCCTCGTCGCCGCCTTCTTCAACGCCATCTTCACCCTGCGGGCCCCGCTCGGCGCGCTGTTGCGGGAGCTCGGCGCCCACGCCCTCGTCGCCGACGCGCTATTCCCGTGGGCCACGGGGCTGGCGGCCGAGATGGGGATCCCGAGGCTTATCTTCCAGGTCATGGGTCTGTTCCCGCTCTGCGGTGCCCACGATCTCGATACCCATCGGCCGCACGAGGCCGTTGGTGGGGATGACGAGGAGTTTACCATCCCGGGGTTTCCGGACCCGGTGAAGCTTACCAGGGGACAGGTCCCCGAGGTCTTCAGGTAAGCTTTTGGAGGGAAACAGGGCAGGCTCAGGGGGGGCAGCAACCGAAGCGGCTGCTTTGGGCCCCCGAGTTAAGGTCCCCTCGATCTTAAATTTAGTGTTGAAGGTAGCAGTAAGTTGATGACTTGGGAGGGCAGCTCTCTCTTCTCTACTGCCATTGGACGCCTCTTAAAAAAGTTTTTTTAATTATAAAATTTCATAAGCCACCATATATTTTTTTTAAAAAAAATTTATAAAGCAAACATGGTAGGAGATTGAAATCTCTCAAATTTCAATCGACTCTAACAAAAAAAAAACTATAACACTCTCTCTTTGATTTTGGTTTGTTTTCTCTCTCTAATTTTAGAGTGCTATTTTGAAATTAAAAAAAATATTGTTAATTAATCCTGTTATCATCTTTCTTTTCGAATTATAATTTTAAATTTAATTTATTATATCCTTATTGTTACTTTAATATTGTTACTTTAAGATAATGAAGCTATGTTTTATTTTTTGGAGTTGTGGGTATATTTTTGTGCTCCTAATGTTAATTTAAACTCAACGATTTTAATAGCTAATTTAAACAATTATATGTGCAAGAAAATTTTTATTTATGGGTAATGGCAATAAAAAAAAATTATAAATAAAAAATAAATTGTAATAGCTCTGCATAAACATTGCAGATTTCTAGTTGTAGTATTTATAGAAGAGACCAGAACTCTCTAGCAAATTACTCTTTCCCACAATGCAGGCACGATTTCATGCTGGCCCTCCTCCGCGACGCAGAGTTCACCAGCTACGGCGTGATAGTGAACAGCTTCTACGCCCTCGAGCCCAGTTACGCGGAGCACTACTACAAGGTGGCCCCCCGGAAGGTCTTCCTCCTCGGCCCGGTCGCCCTCGCCGGCTCCAATCCCTCGCCGCTGTCATTGGAGAGTGGCGACCCCTGCATCACCTGGCTCGACTCCAAGCCCGACAACTCGGTCCTTTACCTGAGCTTCGGCACCACCTGCCGATTCAGCGACGAGCAGCTCGTCGAGCTCGCCGAGGGCCTCGCGTCCTCCGGCCACAACTTTGTGTGGGTCGTGGCCCTCCCCGAGAGCAGCGGCTCGACCGGCAAGGAGTGGCTTCCGGAAGGCTACGAGCATAATGTGGCGGGTCGGGGACTTCTCGTAAGTGGTTGGGCTCCGCAGACCGCGATCCTGAACCACCGGGCGGTGGGTGGGTTCGTGAGCCAATGCGGGTGGAACGCCGTCATGGAGGCGGTGGCGGCAGAGGTACCCATGATCACGTGGCCGCTTCATTCGGACCATTTCATCACCGAGAAGCTGTTCTGCGATGTGCTGCACGTGGCGGTGCCAATGTGGGAGGGGCGGAAAAGCATCTGGGATGATCAGAAGGAGGTGGTGCGGGCGAAAACAGTGGCAGCGTCGGTGAAGCGGCTTATGGGTGGCGGGAACGAGATCGAGGCGATGCGAAGGAGAATGAGGGAGCTCGGGGAATTAGGACGGGCCGCGGTGGCAGAAGGCGGATCGTCCCACTCCGACATGAGCCGTCTTATTCACGTGCTCACGGAGGAGCGAAGCAAGGCTAAGAAAATGATAGACAATAATGGTGTTAGTTATGATTGTTCTGGTCGCAATTGATTATTTCTCGTCAAGAAATGATGGCAGTGTTTTTTGATAAACATGTTTATGTTAGCAATCTCACTTGGAACCCAACACATTGCAAGCTATTTCATCGAGCACCGACATGTCTCTAGT

**Supplementary File 3**

**Pennogenin (1)**: white amorphous powder; 1H-NMR (500 MHz, CD3OD) δH 0.81 (3H, d, J = 6.0 Hz, H-27), 0.85 (3H, s, H-18), 0.90 (3H, d, J = 7.5 Hz, H-21), 1.05 (3H, s, H-19), 3.50 (1H, m, H-3), 4.02 (1H, dd, J = 7.5 Hz, 6.5 Hz, H-16), 5.36 (1H, m, H-6); 13C-NMR (125MHz, CD3OD) δC 38.5 (C-1), 33.2 (C-2), 72.4 (C-3), 43.0 (C-4), 142.3 (C-5), 122.2 (C-6), 32.3 (C-7), 32.9 (C-8), 51.4 (C-9), 37.8 (C-10), 21.7 (C-11), 32.5 (C-12), 45.5 (C-13), 53.9 (C-14), 31.3 (C-15), 90.6 (C-16), 91.3 (C-17), 17.49 (C-18), 19.9 (C-19), 45.8 (C-20), 9.1 (C-21), 110.9 (C-22), 32.1 (C-23), 29.4 (C-24), 33.3 (C-25), 67.7 (C-26), 17.52 (C-27).

**Floribundasaponin A (2)**: white amorphous powder; 1H-NMR (800 MHz, CD3OD) δH 0.81 (3H, d, J = 6.4 Hz, H-27), 0.85 (3H, s, H-18), 0.90 (3H, d, J = 7.2 Hz, H-21), 1.06 (3H, s, H-19), 3.16 (1H, t, J = 8.0 Hz, H-2′), 3.27 (1H, m, H-3), 3.66 (1H, dd, J = 11.2 Hz, 5.6 Hz, H-6′a), 3.86 (1H, brd, J = 11.2 Hz, 5.6 Hz, H-6′b), 4.02 (1H, t, J = 6.4 Hz, H-16), 4.39 (1H, d, J = 8.0 Hz, H-1′), 5.40 (1H, m, H-6); 13C-NMR (200 MHz, CD3OD) δC 38.5 (C-1), 33.3 (C-2), 77.9 (C-3), 39.7 (C-4), 142.0 (C-5), 122.5 (C-6), 32.1 (C-7), 32.9 (C-8), 51.5 (C-9), 38.0 (C-10), 21.7 (C-11), 32.5 (C-12), 45.9 (C-13), 53.9 (C-14), 31.3 (C-15), 90.6 (C-16), 91.3 (C-17), 17.49 (C-18), 19.8 (C-19), 45.5 (C-20), 9.1 (C-21), 111.0 (C-22), 32.1 (C-23), 29.4 (C-24), 30.7 (C-25), 67.7 (C-26), 17.52 (C-27), 102.5 (C-1′), 75.1 (C-2′), 79.8 (C-3′), 71.7 (C-4′), 78.1 (C-5′), 62.8 (C-6′).
